# Genomic and transcriptomic analyses of the subterranean termite *Reticulitermes speratus:* gene duplication facilitates social evolution

**DOI:** 10.1101/2021.07.11.451559

**Authors:** Shuji Shigenobu, Yoshinobu Hayashi, Dai Watanabe, Gaku Tokuda, Masaru Y Hojo, Kouhei Toga, Ryota Saiki, Hajime Yaguchi, Yudai Masuoka, Ryutaro Suzuki, Shogo Suzuki, Moe Kimura, Masatoshi Matsunami, Yasuhiro Sugime, Kohei Oguchi, Teruyuki Niimi, Hiroki Gotoh, Masaru K Hojo, Satoshi Miyazaki, Atsushi Toyoda, Toru Miura, Kiyoto Maekawa

## Abstract

Termites are model social organisms characterized by a polyphenic caste system. Subterranean termites (Rhinotermitidae) are ecologically and economically important species, including acting as destructive pests. Rhinotermitidae occupies an important evolutionary position within the clade representing an intermediate taxon between the higher (Termitidae) and lower (other families) termites. Here, we report the genome, transcriptome and methylome of the Japanese subterranean termite *Reticulitermes speratus*. The analyses highlight the significance of gene duplication in social evolution in this termite. Gene duplication associated with caste-biased gene expression is prevalent in the *R. speratus* genome. Such duplicated genes encompass diverse categories related to social functions, including lipocalins (chemical communication), cellulases (wood digestion and social interaction), lysozymes (social immunity), geranylgeranyl diphosphate synthase (social defense) and a novel class of termite lineage-specific genes with unknown functions. Paralogous genes were often observed in tandem in the genome, but the expression patterns were highly variable, exhibiting caste biases. Some duplicated genes assayed were expressed in caste-specific organs, such as the accessory glands of the queen ovary and frontal glands in soldier heads. We propose that gene duplication facilitates social evolution through regulatory diversification leading to caste-biased expression and subfunctionalization and/or neofunctionalization that confers caste-specialized functions.

**Significance Statement:** Termites are model social organisms characterized by a sophisticated caste system, where distinct castes arise from the same genome. Our genomics data of Japanese subterranean termite provides insights into the evolution of the social system, highlighting the significance of gene duplication. Gene duplication associated with caste-biased gene expression is prevalent in the termite genome. Many of the duplicated genes were related to social functions, such as chemical communication, social immunity and defense, and they often expressed in caste-specific organs. We propose that gene duplication facilitates social evolution through regulatory diversification leading to caste-biased expression and functional specialization. In addition, since subterranean termites are ecologically and economically important species including destructive pests in the world, our genomics data serves as a foundation for these studies.

## Introduction

The evolution of eusociality, i.e., animal societies defined by the reproductive division of labor, cooperative brood care and multiple overlapping generations, represents one of the major transitions in evolution, having increased the level of biological complexity (1). Eusocial insects such as bees, wasps, ants and termites show sophisticated systems based on the division of labor among castes, which is one of the pinnacles of eusocial evolution (2). Recent advances in molecular biological technologies and omics studies have revealed many molecular mechanisms underlying eusociality and have led to the establishment of a new field of study known as “sociogenomics” (3). The genomes of major eusocial hymenopteran lineages, i.e., ants, bees and wasps, have been sequenced, and the differences in gene expression, DNA methylation (4–6)(7) (8–11) and histone modification (12) (13) among castes have been explored. These sociogenomics studies in hymenopterans revealed some genetic bases of social evolution, including the co-option of genetic toolkits of conserved genes, changes in protein-coding genes, cis-regulatory evolution leading to genetic network reconstruction, epigenetic modifications and taxonomically restricted genes (TRG) (14, 15).

Isoptera (termites) is another representative insect lineage exhibiting highly sophisticated eusociality and a wide range of social complexities (16). Termites are hemimetabolous and diploid insects that are phylogenetically distant from hymenopterans with holometaboly and haplodiploidy. Termite societies are characterized by reproductives of both sexes, workers and soldiers. In the termite lineage,eusociality is thought to have evolved once, although the levels of social complexity features, such as colony size, feeding habitat, symbiosis with microorganisms and caste developmental pathways, diverged among termite species. These characteristics are especially different between the two major termite sublineages, i.e., the early-branching families (called “lower” termites) and the most apical family Termitidae (“higher” termites) (Fig. 1a). To date, based on the whole-genome sequences of a few termite species, the commonality and diversity of genetic repertoires between Isoptera and Hymenoptera or between termite lineages and their solitary outgroup (i.e., cockroach) have been investigated (17, 18). Additionally, in *Zootermopsis nevadensis*, clear differences in gene expression levels among castes (17) and in DNA methylation between alates and workers (19) were detected.

**Figure 1:**
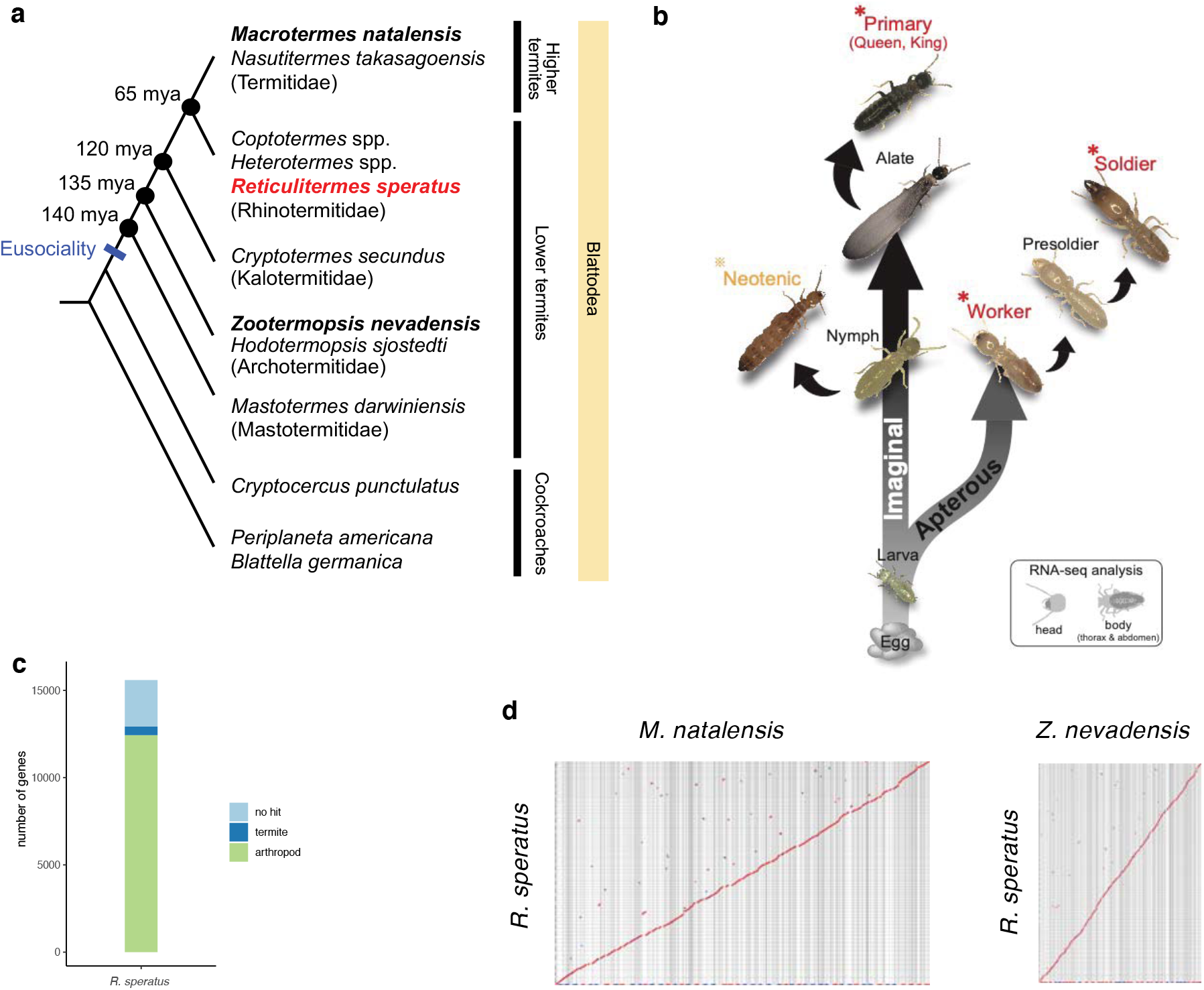
Phylogenetic position of *Reticulitermes speratus* in Blattodea, its developmental pathway, and evolution of the gene repertoire and genome structure. **(a)** Phylogenetic tree of termites and cockroaches. Estimated divergence dates (mya: million years ago) are based on Bucek et al. (80). *R. speratus* is marked in red, and two termites mainly compared in this study are marked with bold characters. **(b)** Developmental pathway of *R. speratus*. There are 2 larval stages before the molt into a nymph (with wing buds) or worker (no wing buds). There are 6 imaginal stages, and the 6th-stage nymphs molt into alates, which are primary reproductives (queen and king). Secondary reproductives (neotenics) differentiate from the 3rd- to 6th-stage nymphs. In the apterous line, there are at least 5 stages of workers. Some workers in the colony molt into presoldiers and soldiers. Female neotenics used for genome sequencing and 3 castes used for RNA-seq are marked with asterisks. **(c)** Gene repertoire of *R. speratus* categorized by orthology. *R. speratu*s genes were compared to those of 88 arthropods and grouped into three classes: orthologs shared with other arthropods (labeled ‘arthropod’), orthologs shared with other termites (*Z. nevadensis* and/or *M. natalensis*) but with no orthologs in other arthropods (labeled ‘termite’), and orphan genes unique to *R. speratus* (labeled ‘no hit’). **(d)** High conservation of synteny between termite genomes revealed by dot plots generated by comparing *R. speratus* with *Z. nevadensis* and *M. natalensis*. Scaffolds longer than 2.0 Mb in the *R. speratus* assembly are used for plotting. Forward alignments are plotted in red and reverse alignments are plotted in blue.

Among the more than 2900 extant species of termites (Isoptera) (20), subterranean termites (Rhinotermitidae), especially two genera, *Reticulitermes* and *Coptotermes*, occupy an important evolutionary position (Fig 1a). Recent phylogenetic studies showed that Rhinotermitidae is paraphyletic, and a clade including *Reticulitermes, Coptotermes* and *Heterotermes* was shown to be sister to Termitidae (21, 22). In particular, *Reticulitermes* exhibits intermediate social complexity between those of higher (Termitidae) and lower (all the other families) termites (23), for example, this genus displays primitive feeding ecology and gut symbiont features, a relatively complex colony structure and a caste development mode termed the bifurcated pathway (Fig. 1b). Moreover, *Reticulitermes* is the most common termite group in palearctic (24) and nearctic (25) regions and a major pest causing serious damage to human-made wooden structures (26). For these reasons, members of this genus are probably among the most studied termites (16). Nevertheless, despite their evolutionary, ecological and economic relevance, subterranean termites remain an understudied group in terms of both genetics and genomics.

In this study, we targeted the Japanese subterranean termite *Reticulitermes speratus*. We conducted whole-genome sequencing, caste-specific RNA-seq analysis and whole-genome bisulfite sequencing of *R. speratus* to understand the genomic, transcriptomic and epigenetic bases of the social life of this termite species. *R. speratus* nymphoids are almost exclusively produced parthenogenetically by automixis with terminal fusion in primary queens, such that the genome should be homozygous at most loci (27), which provides an advantage in genome sequencing. We also compared the omics data of *R. speratus* with those of sequenced higher and lower termites (Fig. 1a). Our integrative analyses of the genome and transcriptome of *R. speratus* and other termites revealed that gene duplications are often associated with caste-biased gene expression and caste-specific functions, which highlights the significant role of gene duplication in eusocial evolution in the termite lineage.

## Results and Discussion

### Genomic features of *Reticulitermes speratus*

Genome sequencing of *R. speratus* was performed with genomic DNA isolated from female secondary reproductives (nymphoids) [Fig. 1b]. *R. speratus* nymphoids are almost exclusively produced parthenogenetically by automixis with terminal fusion in primary queens, such that the genome should be homozygous at most loci (27) and thus ease de Bruijn-graph-based genome assembly. We generated a total of 86 Gb of Illumina HiSeq sequence data and assembled them *de novo* into 5817 scaffolds with an N50 of 1.97 Mb and total size of 881 Mb [Table 1], covering 88% of the genome based on the genome size (1.0 Gb) estimated by flow cytometry (28). The assembled *R. speratus* genome has high coverage of coding regions, capturing 99.2% (98.5% complete; 0.7% fragmented) of 1367 Insecta benchmarking universal single-copy orthologs (BUSCOs) (29) [Table 1]. The *R. speratus* genome is rich in repetitive elements, which make up 40.4% of the genome. A total of 15,591 protein-coding genes were predicted by combining the reference-guided assembly of RNA-seq reads (36 libraries derived from different castes, sexes and body parts; see below for details) and homology-based gene prediction followed by manual curation of gene families of interest [Fig. 1c]. Whole-genome bisulfite sequencing revealed extensive gene body methylation of the *R. speratus* genome, amounting to 8.8% of methylated cytosines in the CG context [Supplementary Fig. 1]. The genome-wide DNA methylation landscape was similar to that of a dampwood termite *Z. nevadensis* (12%) (19). These omics data and a genome browser are available at http://www.termite.nibb.info/retsp/.

**Table 1.**
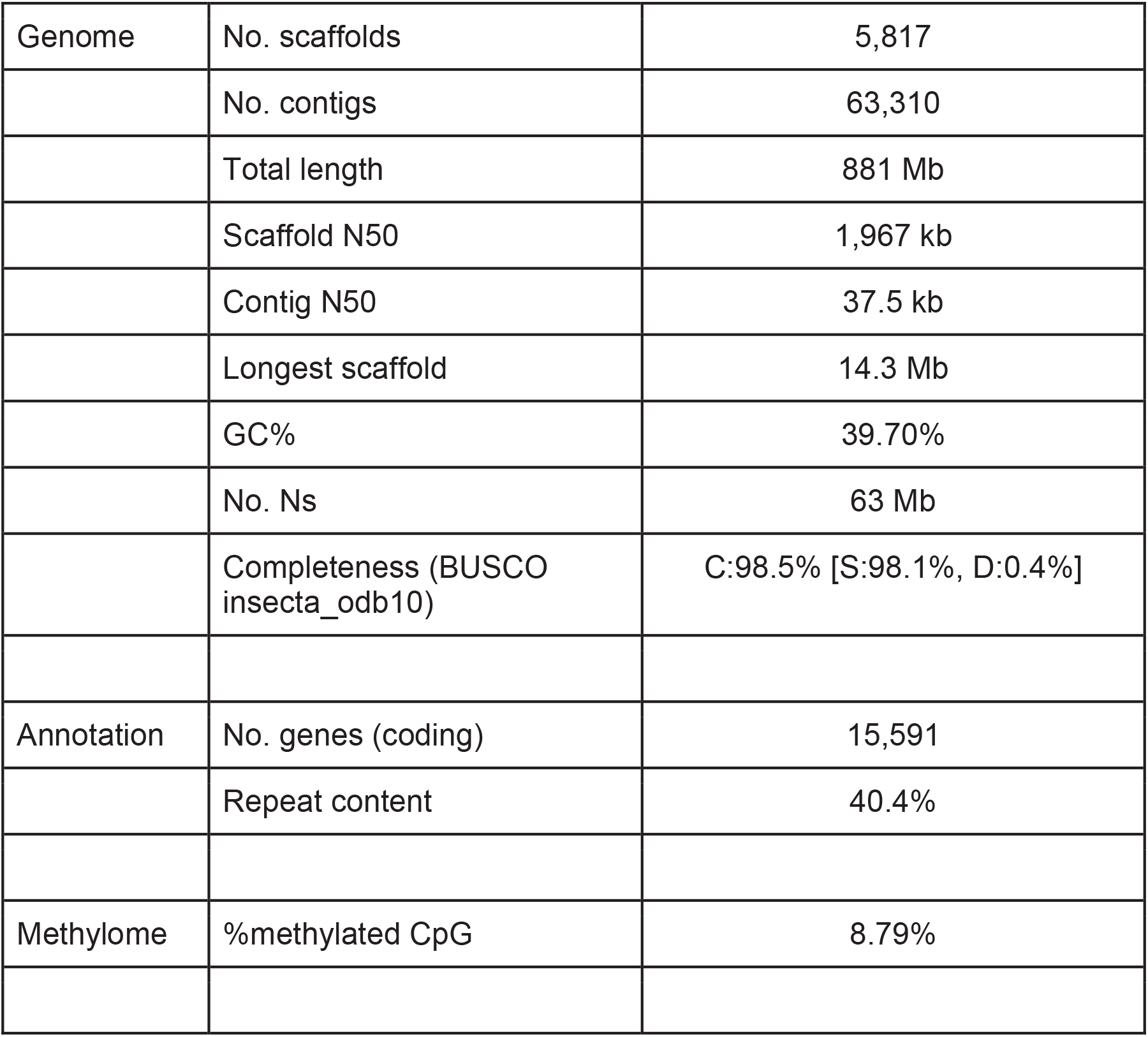
Summary of *Reticulitermes speratus* genome assembly, annotation and methylome.

|  |  |  |
| --- | --- | --- |
| Genome | No. scaffolds | 5,817 |
|  | No. contigs | 63,310 |
|  | Total length | 881 Mb |
|  | Scaffold N50 | 1,967 kb |
|  | Contig N50 | 37.5 kb |
|  | Longest scaffold | 14.3 Mb |
|  | GC% | 39.70% |
|  | No. Ns | 63 Mb |
|  | Completeness (BUSCO<br>insecta_odb10) | C:98.5% [S:98.1%, D:0.4%] |
| Annotation | No. genes (coding) | 15,591 |
|  | Repeat content | 40.4% |
| Methylome | %methylated CpG | 8.79% |

We compared the *R. speratus* gene repertoire with those of 88 other arthropods, including the two termites *Z. nevadensis* and *Macrotermes natalensis* (17, 30). Ortholog analysis showed that 12,032 (82.9%) of the 15,591 genes in *R. speratus* genes were shared with other arthropods, and 1773 were taxonomically restricted (TRGs) to Isoptera, among which 430 were shared with the other two termites and 1343 were unique to *R. speratus* [Fig. 1c]. Whole-genome comparison with two sequenced termites, *M. natalensis* and *Z. nevadensis*, showed a high degree of synteny conservation [Fig. 1d]. We identified 2799 syntenic blocks (N50: 858.4 kb) shared with *M. natalensis* that covered 95.1% of the *R. speratus* genome where 560.4 Mb of nucleotides was aligned, while 3650 syntenic blocks (N50: 591.1 kb) shared with *Z. nevadensis* covered 72.1% of the *R. speratus* genome where 116.7 Mb was aligned. Only a few cases of large genomic rearrangements were found between termite genomes, at least, at the contiguity level of the current assemblies, suggesting overall conservation of genome architecture in the lineage of termites over 135 MY (Fig. 1a). Interestingly, despite such high conservation of macrosynteny, interruptions or breaks in local synteny were observed and often associated with tandem gene duplications. For example, when we examined regions containing large tandem gene duplications (> 5-gene tandem duplications), synteny between the *R. speratus* and *M. natalensis* genomes was interrupted in 10 of 21 regions (examples shown in Supplementary Fig 3).

### Transcriptome differentiation among castes

Distinct castes arise from the same genome, a phenomenon called caste polyphenism which is a distinctive hallmark in social insects (31, 32). To elucidate caste-biased gene expression in order to understand the mechanism underlying the caste-specific phenotypes, we compared the transcriptomes of three castes (primary reproductives, workers and soldiers) in *R. speratus* [Fig. 1b; Supplementary Table 2]. We sequenced 36 RNA-seq libraries, representing three biological replicates of both sexes and two body parts (“head” and “thorax + abdomen”) for each of the three castes.

The results clearly showed that termite castes were distinctively differentiated at the gene expression level. The multidimensional scaling (MDS) plot depicted the three castes as clearly distinct transcriptomic clusters for both the head and thorax + abdomen transcriptomes [Fig. 2a]. However, little sexual difference was detected within each caste, although reproductives showed substantial transcriptomic differences in thorax + abdomen samples between queens and kings, probably due to the difference in the reproductive organs [Fig. 2a]. Using a generalized linear model (GLM) with caste and sex as explanatory variables, we identified 1579 and 2076 genes differentially expressed among castes in head and thorax + abdomen samples, respectively, with the criteria of a false discovery rate (FDR)-corrected P < 0.01, while we identified only 6 and 79 genes that were differentially expressed between sexes in the head and thorax + abdomen samples, respectively, with the same criteria. We focused on the genes that were differentially expressed among castes (caste-DEGs) and further classified them into three categories of caste-biased genes (i.e., reproductive-, worker-, and soldier-biased genes), with a criterion of >2-fold higher expression relative to that in the other two castes. These caste-biased genes should account for the specialized functions of each caste. Soldier samples exhibited the highest number of caste-specific genes, suggesting the highly specialized functions of the soldier caste. This is consistent with the finding of a previous RNA-seq analysis of the Eastern subterranean termite *Reticulitermes flavipes*, reporting that a majority of DEGs were soldier-specific (33); 73 of the 93 DEGs identified were up- or downregulated specifically in the soldier caste. In addition to these soldier-specific *R. flavipes* genes (e.g., *troponin C* and fatty *acyl-CoA reductase*), the caste-biased genes identified in our transcriptome analysis of *R. speratus* included genes previously reported as caste-biased genes in other termites (34–37), e.g., *vitellogenin* (reproductives), *geranylgeranyl pyrophosphate synthase* (soldiers), and *beta-glucosidase* (probably associated with cellulase; workers). This consistency between transcriptome analyses of different termite species indicates that the RNA-seq analysis in this study is reliable and that the regulation and perhaps the functions of these caste-biased genes are conserved across the termite lineage.

**Figure 2:**
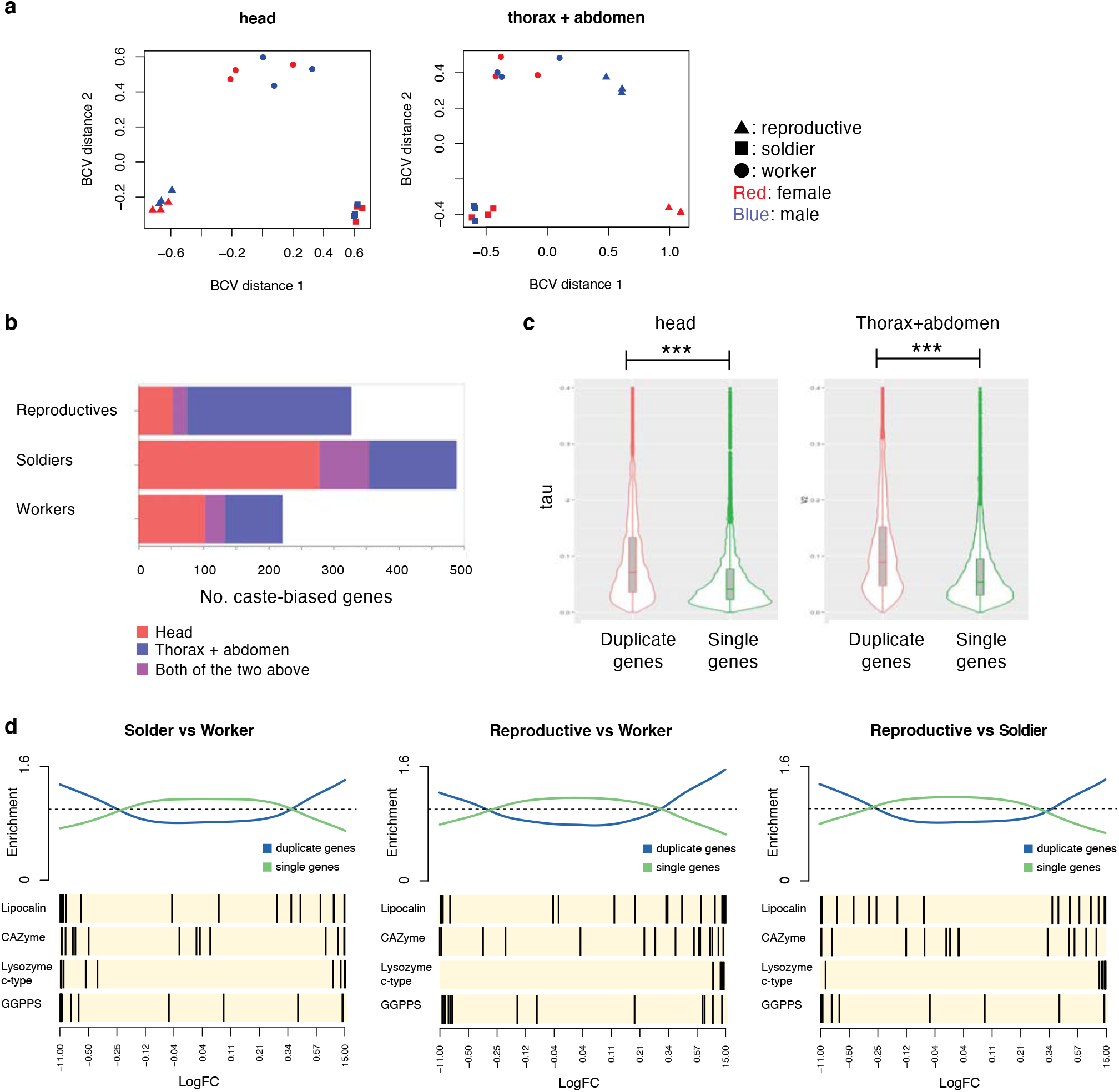
Caste-specific transcriptome analysis and the enrichment of duplicate genes for caste-biased genes. **(a)** Multidimensional scaling (MDS) plot of RNA-seq data showing relatedness between the expression profiles of different castes (reproductive, soldier and worker) and sexes (male and female). The left panel plots RNA-seq data from head samples, and the right panel plots data from thorax + abdomen samples. Three biological replicates were analyzed for each condition and plotted individually. **(b)** Numbers of caste-biased genes with >2-fold higher expression levels than the other two castes. Colours in each bar indicate the differences of RNA-seq data obtained. **(c)** Violin plots showing the distribution of tau indexes of duplicate genes and single genes. Tau values range between 0 and 1, with low values indicating invariable and constitutive expression between castes and higher values supporting caste specificity. In both body part samples, the tau values of duplicate genes were significantly greater than those of single genes (p < 2.2e-16, Wilcoxon rank sum test). **(d)** Enrichment of caste-DEGs (differentially expressed genes among castes) for duplicate genes. In each comparison between castes (soldier vs worker, reproductive vs worker,and reproductive vs soldier), all genes are ranked and ordered by log-fold-change value along the horizontal axis. Black bars mark the positions of genes. Genes of sociality-related functions highlighted in the text are selected and plotted in the lower panels. Curved lines in the upper panel show relative enrichment of the duplicate genes (blue line) or single genes (green line) relative to uniform ordering.

The caste-DEGs were enriched for Gene Ontology terms related to a wide array of functions [Supplementary Table 4], such as hormone metabolism, chitin metabolism, hydrolase activity, oxidoreductase activity, lipid metabolism, signaling and lysozyme activity. Protein motifs enriched in the caste-DEGs were also identified, including cytochrome P450, lipocalin, lysozyme, glycosyl hydrolase family, TGF-beta and trypsin [Supplementary Table 5]. Among the 1773 taxonomically restricted genes (TRGs) that were restricted to Isoptera (see above), the termite-shared TRGs showed strong enrichment for caste-DEG (Fisher’s exact test, P < 1.0e-7 for head samples, P < 1.0e-10 for thorax+abdomen samples), while the TRGs found only in *R. speratus* (orphan genes) did not (P = 0.99 and P = 0.97, respectively).

To investigate the relationship between caste-biased gene expression and DNA methylation, we analyzed differential methylation levels among three castes (reproductives, workers, and soldiers). The BS-seq data showed that the global CpG methylation patterns were very similar among the castes [Supplementary Fig. 2ab], in contrast to the methylation pattern of *Z. nevadensis*, in which DNA methylation differed strongly between castes (winged adults vs. final-instar larvae) and was strongly linked to caste-specific splicing (19). Instead, gene body DNA methylation of *R. speratus* seems to be important for the expression of housekeeping genes, as reported in the drywood termite *Cryptotermes secundus* (18). Housekeeping genes exhibited a high degree of gene body methylation in all castes of *R. speratus*, while caste-biased genes showed a significantly lower level DNA methylation [Supplementary Fig. 2cd].

### Gene duplication and caste-biased gene expression

Evolutionary novelties are often brought about by gene duplications (38) (reviewed in (39)), and the transition to eusociality in Hymenoptera has been associated with gene family expansion (18, 40, 41). Our ortholog analysis comparing the *R. speratus* gene repertoire with those of 88 other arthropods identified 1396 multigene families duplicated in the *R. speratus* genome. Interestingly, compared to the genome as a whole, the set of caste-DEGs identified above was significantly enriched for genes in multigene families (X-squared = 218.62, df = 1, p-value < 2.2e-16). We also calculated the tau score as a proxy of caste specificity of gene expression for all genes and found that duplicated genes were significantly more caste-specific than single-copy genes (p < 2.2e-16, Wilcoxon rank sum test) in both transcriptome data sets (head and thorax+abdomen) [Fig. 2b]. Additionally, gene set tests showed that sets of duplicated genes were differentially expressed in all pairwise comparisons between castes [Fig. 2c]. These data highlight the important roles of gene duplication in the caste evolution of termites.

### Multigene families related to caste-specific traits in *R. speratus*

Caste-biased multigene families were associated with diverse functional categories, some of which were strongly related to caste-specific behaviors and tasks. Here, we highlight five families, namely, lipocalins (protein transporters for social communication and physiological signaling), cellulases (carbohydrate-active enzymes for worker wood digestion), lysozymes (immune-related genes for social immunity), geranylgeranyl diphosphate synthases (metabolic enzymes for the production of soldier defensive chemicals), and a novel termite-specific gene family with unknown functions, as examples of multigene families relevant to termite sociality. Molecular evolution studies have shown that the redundancy caused by gene duplication may allow one paralog to acquire a new function (neofunctionalization) or divide the ancestral function among paralogs (subfunctionalization) (38, 39). We are particularly interested in the evolutionary impact of gene duplication on caste specialization through neo/subfunctionalization.

#### Lipocalins

Lipocalins belong to a family of proteins, with molecular recognition properties such as the ability to bind a range of small hydrophobic molecules (e.g. pheromones) and specific cell surface receptors, and to form complexes with soluble macromolecules (42). A previous study identified a gene of the lipocalin family, SOL1, that is exclusively expressed in the mandibular glands of mature soldiers of the rotten-wood termite *Hodotermopsis sjostedti* (43). SOL1 is thought to function as a signaling molecule for defensive social interactions among termite colony members (31). Moreover, RNA-seq analysis showed that a lipocalin gene, *Neural Lazarillo homolog 1* (*ZnNlaz1*), was specifically expressed in soldier-destined larvae in an incipient colony of *Z. nevadensis* (44). Gene function and protein localization analyses suggested that ZnNLaz1 was a crucial regulator of soldier differentiation through the regulation of trophallactic interactions with a queen. Thus, it was of interest that the lipocalin-related motif (Pfam PF00061; lipocalin/cytosolic fatty-acid binding protein family) was significantly enriched in the list of caste-DEGs (FDR < 0.05; Supplementary Table 5).

We identified 18 lipocalin family genes in the *R. speratus* genome [Fig. 3a-c; Supplementary Table 7]. The number of lipocalin genes was larger than those in other insects [Fig. 3c]. Phylogenetic analysis of lipocalin family genes identified in arthropods, including three termite species, revealed a highly dynamic evolutionary history of this protein family [Fig. 3a]. A couple of subfamilies, namely clades A and B, had experienced extensive expansion in the termite lineage. The most drastic expansion was found in clade A, which includes *H. sjostedti* SOL1. In this clade, 7, 9 and 5 genes were identified in *R. speratus, M. natalensis* and *Z. nevadensis*, respectively, and extensive and independent gene expansions occurred in each species. Clade B was also composed of genes with a termite lineage specific expansion. In many cases, these lipocalin genes were found in tandem arrays in the *R. speratu*s genome [Fig. 3b]. The inferred phylogenetic tree indicated that duplications in each clade occurred after the divergence of termites from a common ancestor.

**Figure 3:**
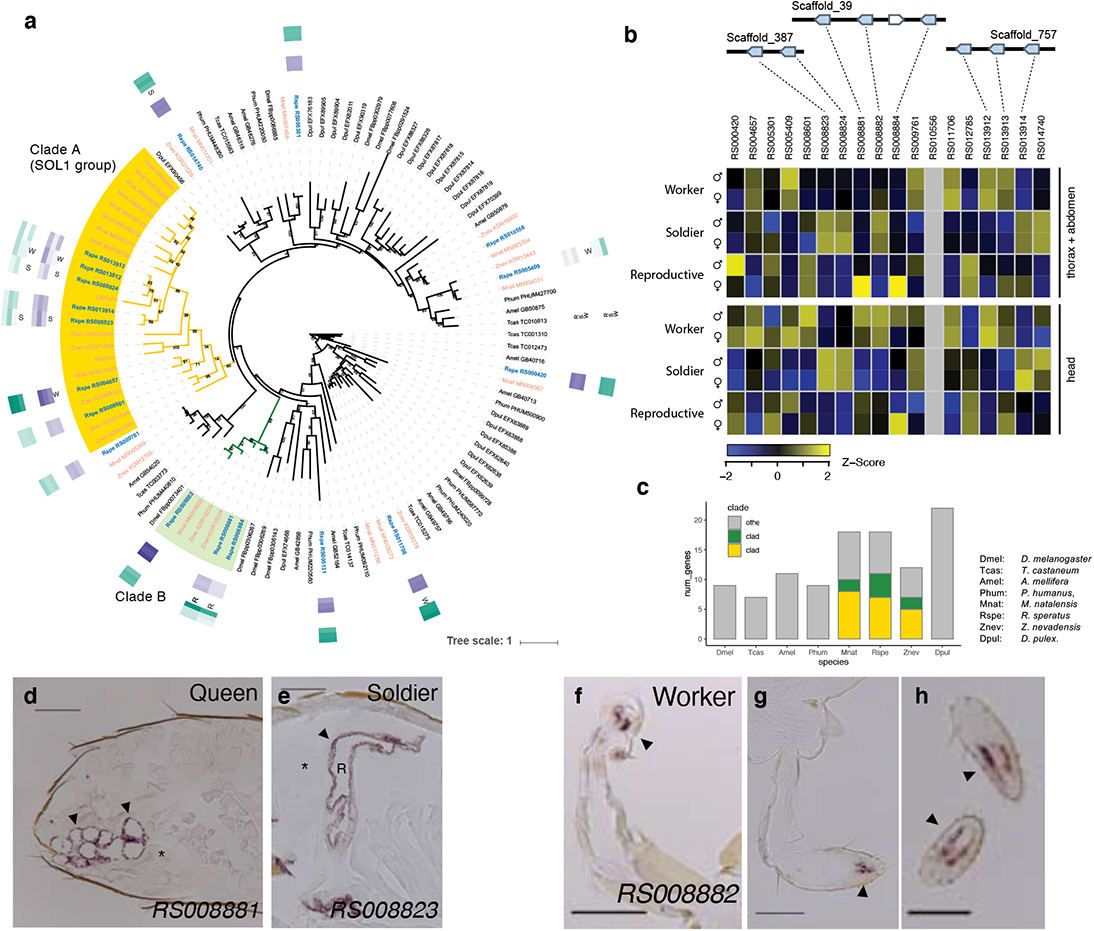
Lipocalin genes in *R. speratu*s. **(a)** Maximum likelihood (ML) tree of lipocalin homologs based on the amino acid sequences obtained with a log gamma (LG) model. Branches leading to clade A and clade B, which show gene family expansion specific to termite sublineages, are marked in yellow and green, respectively. Color gradients in the outer tracks show the expression levels as averaged log(RPKM+1) values in three castes (reproductive, soldier, and worker). Expression levels of head samples and thorax + abdomen samples are shown in purple and green, respectively. Caste-DEGs (differentially expressed genes among castes) are marked as R, S, or W beside the color gradients, indicating biases toward the reproductive, soldier, or worker caste, respectively. **(b)** Lipocalin multigene clusters in the *R. speratus* genome and their relative expression levels among castes. The heatmap shows the Z-scores of the log(RPKM+1) values in the caste-specific transcriptome. **(c)** Comparison of the number of lipocalin subclasses among representative arthropods. Note clades A and B are specific to termites. **(d)** Vertical cryosection of the queen abdomen subjected to *in situ* hybridization with an antisense DIG-labeled *RS008881* mRNA probe. The accessory gland cell layer is stained dark (arrowhead), in contrast to the other ovarian tissues, including the spermatica (asterisk). Bar = 0.2 mm. **(e)** Photographs of *in situ* hybridization for *RS008823* mRNA in the soldier head. The front of the head is on the left side. The gland cell layer surrounding the frontal gland reservoir (R) is stained dark (arrowhead). The asterisk indicates the brain. Bar = 0.1 mm. **(f, g, h)** Vertical cryosection of the worker antenna (f) and horizontal cryosections of the worker labial palp (g, right palp) and maxillary palp (h, the last segment of the left (upper) and right (lower) palp) subjected to *in situ* hybridization for *RS008882* mRNA. Tissues around some sensilla are stained dark (arrowhead). Bar = 0.1 mm. Photographs of cryosections hybridized with sense probes (negative controls) are shown in Supplementary Fig. 6a-c.

A comparison of the transcriptome among castes revealed that most lipocalin genes (15 of 18) showed caste-biased gene expression [Fig. 2, Fig. 3ab]. The caste specificity, however, varied among genes, regardless of sequence similarity and positional proximity on the genome. In particular, the expression levels of genes in clades A and B drastically changed among castes. For example, *RS008823* and *RS008824* displayed solder-specific expression, the expression of *RS013912* was biased toward workers, and *RS013913* was downregulated in soldiers. *RS008881* and *RS008884* were exclusively expressed in queen bodies (thorax + abdomen), while *RS008882*, a gene next to the aforementioned two genes, showed quite different expression patterns and high expression levels in heads, especially those of workers. These results indicate that termite lipocalin genes underwent dynamic expansion in terms of gene repertoire, and regulatory diversification of caste-biased expression. This gene expansion and regulatory diversification of lipocalins may facilitate the evolution of the molecules involved in signaling during caste development and among individuals through social interactions.

To address the caste-specific function of the lipocalin paralogs, the expression patterns of several selected caste-biased lipocalin genes were examined by *in situ* hybridization [Fig. 3d-h, Supplementary Fig. 6]. *RS008881*, a queen-biased lipocalin gene, was found to be expressed exclusively in the accessory glands of the ovary [Fig. 3d]. The next gene on the same scaffold, *RS008882*, was shown to be specifically expressed in worker antennae and maxillary/labial palps [Fig. 3f-h]. *RS008823*, a soldier-biased gene, was expressed exclusively in the frontal gland cells of the soldier heads [Fig. 3e]. Note that ovaries and frontal glands develop during postembryogenesis in a caste-specific manner (i.e., ovaries in queens and frontal glands in soldiers) in the pathway of caste differentiation in *R. speratus*. Antennae and maxillary/labial palps are not caste-specific but crucial sensory organs, especially for blind termite immatures, such as workers. Given that animal lipocalins generally work as carrier proteins (45), there is a possibility that focal termite lipocalins bind and convey some molecules to the targets from caste-specific organs (e.g., egg-recognition pheromone and soldier defensive and/or inhibiting substances; (46–48)), or participate in sensory reception, such as the role of odorant-binding proteins (49).

#### Cellulases

Lignocellulose degradation in termites is achieved by a diverse array of carbohydrate-active enzymes (CAZymes) produced by the host and their intestinal symbionts. The repertoire of CAZyme families in the genome of *R. speratus* did not show considerable differences from those of other nonxylophagous insects, such as a honeybee and a fruit fly (Supplementary Fig. 4). However, we found gene family expansion and expressional diversification for glycoside hydrolase family (GH) 1 and GH9 members. The majority of GH1 and GH9 members are β-glucosidase (BGs; EC 3.2.1.21) and endo-β-1,4-glucanases (EGs; EC 3.2.1.4), respectively, which are essential for cellulose digestion in termites (50).

We identified 16 GH1 paralogs [Supplementary Table 8]. Such gene expansion of GH1 was also observed in the genome of other termites, but the reason for the gene expansion remains elusive (51). Although the phylogenetic tree divided these GH1 paralogs into four distinct groups (clades A to D in Fig. 4a), most of them were tandemly located in the genome of *R. speratus* (Fig. 4bc). The predominantly expressed BG gene was *RS004136*, while the expression of this gene was clearly biased toward the bodies (thorax + abdomen) of reproductives and workers (Fig. 4b). This gene formed a rigid clade with *bona fide* BGs reported from the salivary glands or midgut of termites (clade A in Fig. 4a) (52), suggesting that this gene is involved in cellulose digestion in *R. speratus*. Indeed, *in situ* hybridization analysis showed that *RS004136* was specifically expressed in the salivary glands of workers (Fig. 4de, Supplementary Fig. 7a). Other GH1 members showed a wide variety of expression patterns across castes and body parts [Fig. 4b]. Some of them might have diversified their functions, other than wood digestion, related to termite sociality, such as egg-recognition pheromones (53). A typical example displaying such diversification was *RS004624*, which was expressed specifically in the abdomens of queens [Fig. 4b]. The peptide sequence of this gene showed a monophyletic relationship with that of Neofem2 of *Cryptotermes secundus* (clade D in Fig. 4a), which is a queen recognition pheromone probably functioning in the suppression of reproductive emergence (54). *In situ* hybridization showed that *RS004624* was specifically expressed in the accessory glands of queen ovaries (Fig. 4fg, Supplementary Fig. 7b), suggesting that *RS004624* is involved in enzymatic activities in queen-specific glands. Together with the results for a queen-biased lipocalin (*RS008884*), this finding indicates that the queen accessory glands may produce some queen-specific pheromones. Like lipocalins, GH1 paralogs are also typical examples of multigene family members participating in caste-specific tasks, which may be acquired by gene duplication resulting in neo- or subfunctionalization.

**Figure 4:**
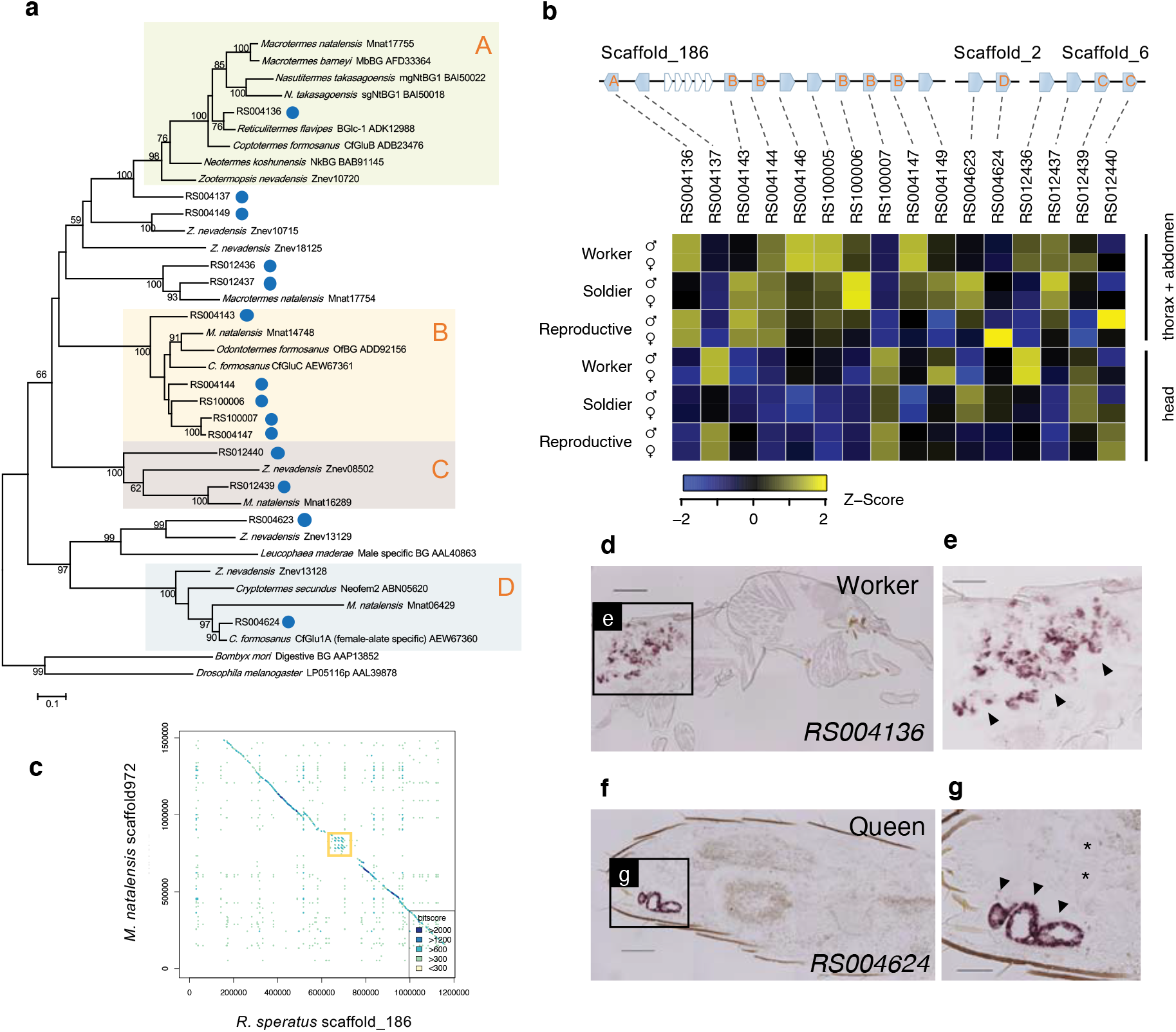
Glycoside hydrolase family (GH) 1 in the *R. speratus* genome. **(a)** ML tree of GH1 genes based on the amino acid sequences obtained with a LG+G+I model. Fourteen of 16 GH1 genes in *R. speratus* were used; two genes (*RS004146* and *RS100005*) were removed from the analysis due to incomplete retrieval of the coding sequences from gapped scaffolds. GH1 subclasses are colored and labeled A, B, C, and D. **(b)** GH1 multigene clusters in the *R. speratus* genome and their expression levels. Letters A-D on the gene structures represent GH1 subclasses categorized in the phylogenetic tree in (a). The heatmap shows the Z-scores of the log(RPKM+1) values in the caste-specific transcriptome. **(c)** Synteny comparison around the GH1 multigene cluster region (orange rectangle) between *R. speratus* and *M. natalensis* genomes. **(d)** Vertical cryosection of the worker thorax subjected to *in situ* hybridization with an antisense DIG-labeled *RS004136* mRNA probe. The head part is on the right side. Bar = 0.2 mm. **(e)** Magnified view of the worker thorax. The salivary gland cells are specifically stained dark (arrowhead). Bar = 0.1 mm. **(f)** Vertical cryosection of the queen abdomen subjected to *in situ* hybridization for RS004624 mRNA. Bar = 0.2 mm. **(g)** Magnified view of the queen ovary. The accessory gland cell layer is stained dark (arrowhead), in contrast to the other ovarian tissues, including ovarioles with two oocytes (asterisks). Bar = 0.1 mm. See Supplementary Fig. 7a-b for negative controls of the *in situ* hybridization experiments (d-g).

We found four paralogs of GH9 in *R. speratus* [Supplementary Fig. 5, Supplementary Table 8]. Although several insect GH9 EGs have acquired the ability to hydrolyze hemicellulose (55), neo- or subfunctionalization of termite EGs has yet to be clarified. Intriguingly, we found that the GH9 member *RS006396* was weakly but uniformly expressed across all termite body parts and castes. This result suggests that some GH9 members also perform a function other than that of cellulase, as is the case for GH1.

#### Lysozymes

The immune system of termites is of particular interest, because the group living of termites with nonsclerotized and nonpigmented epidermis and microbe-rich habitat puts them at high risk for pathogenic infections (56). Thus, defense against pathogenic microbes is important for termites. In the *R. speratus* genome we identified 251 immune-related genes [Supplementary Information 1.9, Supplementary Table 26]. The repertoire and number of immune-related genes of *R. speratus* showed no large differences compared to those of other insect species, but a notable exception was found for lysozymes [Supplementary Fig. 8].

Lysozymes are involved in bacteriolysis through hydrolysis of β-1,4-linkages in the peptidoglycans present in bacterial cell walls, and three distinct types of lysozymes, chicken- or conventional-type (c-type), goose-type (g-type), and invertebrate-type (i-type) lysozymes, have been found in animals (57). We identified 13 and 3 genes encoding c-type and i-type lysozymes, respectively, and the number of lysozyme genes was larger than those in other insects [Fig. 5, Supplementary Table 9]. Phylogenetic analysis revealed that c-type lysozymes underwent extensive gene duplications in the sublineage leading to *R. speratus* [Fig. 5a]. Seven c-type lysozymes formed a tandem array on scaffold_859 [Fig. 5b], probably generated by repeated tandem gene duplication events. Interestingly, most of the c-type lysozyme genes showed caste-biased expression. Three genes (*RS014698, RS100022*, and *RS100023*) exhibited high expression levels compared to those of other lysozyme genes and were expressed in a soldier-specific manner, while *RS100026* was expressed in a worker-specific manner and *RS100024* and *RS100025* were highly expressed in both workers and soldiers [Fig. 5b]. The differential expression patterns of the lysozyme genes in *R. speratus* may represent division of labor among castes in terms of colony-level immunity.

**Figure 5:**
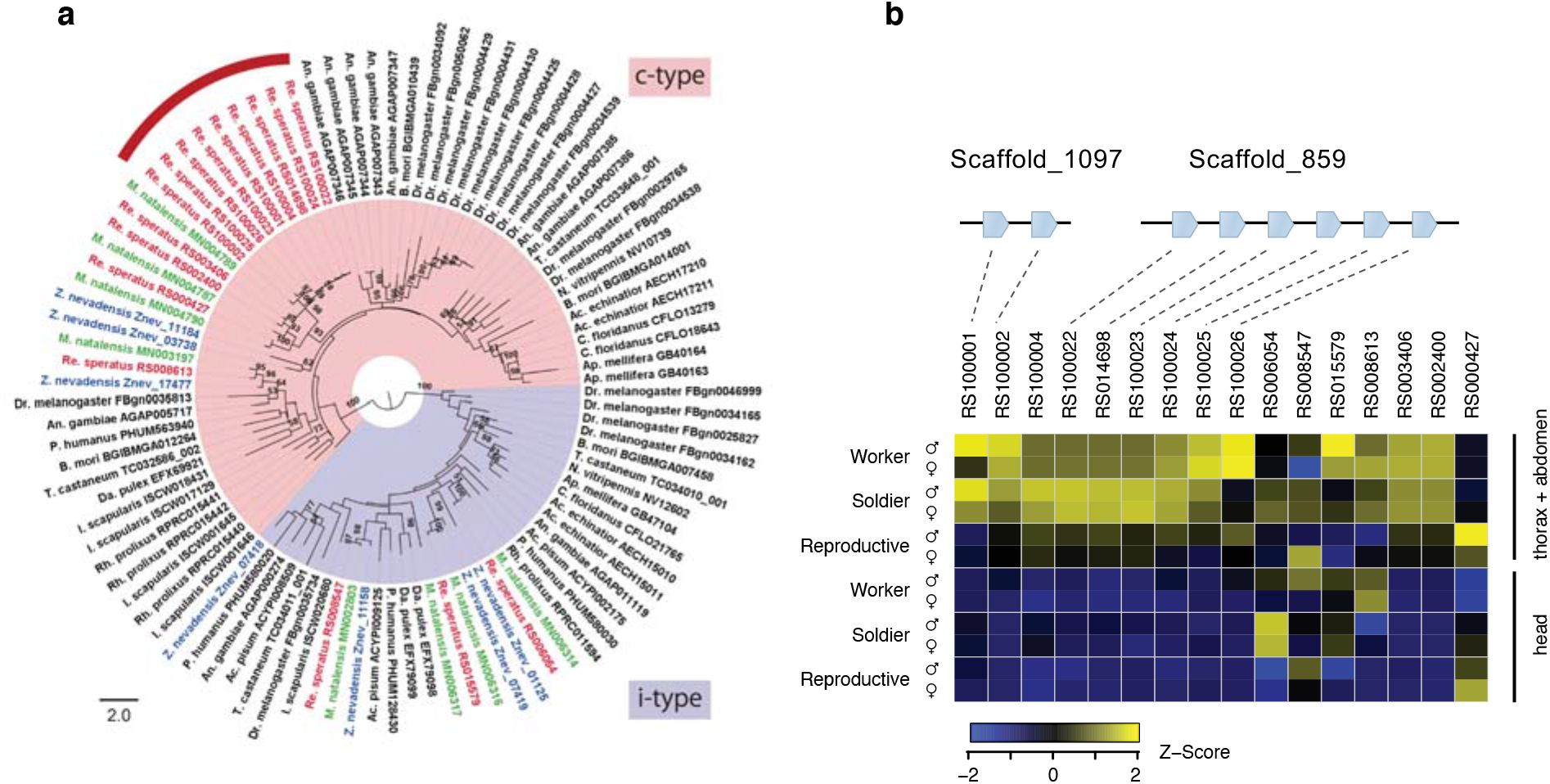
Lysozyme family in the *R. speratus* genome. **(a)** ML tree of lysozyme genes with a GTR+G model. The red curve indicates a lineage-specific gene expansion observed in the *R. speratus* genome for a c-type lysozyme. **(b)** Lysozyme multigene clusters in the *R. speratus* genome and their relative expression levels among castes. The heatmap shows the Z-scores of the log(RPKM+1) values in the caste-specific transcriptome.

It is also possible that duplicated lysozymes may have functions other than immunity. A previous study indicated that the salivary glands of *R. speratus* secrete c-type lysozymes to digest bacteria ingested by termites through social feeding behavior (58). The same lysozyme genes are also expressed in the queen ovaries and eggs and play a role in egg recognition as proteinaceous pheromones in *R. speratus* (48, 53). We could not find identical sequences of these lysozyme genes in our gene models, but these sequences were most closely related to *RS002400* with 88% nucleotide identity, which occupied the basal position of the lineage-specific gene expansion (Fig. 5a).

#### GGPP synthase

Whole-genome comparison of *R. speratus* with *Z. nevadensis* and *M. natalensis* revealed a 270 kb *R. speratus*-specific fragment in scaffold_31, while the rest of this scaffold showed very high syntenic conservation among the three termites [Fig 6a]. We found that the *R. speratus*-specific region was encompassed by a tandemly duplicated gene cluster composed of 13 genes encoding geranylgeranyl diphosphate (GGPP) synthase [Fig. 6b, Supplementary Table 10]. GGPP synthase catalyzes the consecutive condensation of an allylic diphosphate with three molecules of isopentenyl diphosphate to produce GGPP, an essential precursor for the biosynthesis of diterpenes, carotenoids and retinoids (59–61). The extensive duplication of GGPP synthase paralogs observed in *R. speratus* is unusual because the genomes of other insects surveyed have only a single copy of GGPS synthase gene. The phylogenetic analysis of GGPP synthase homologs revealed two clusters, a possibly ancestral group (including *RS007484*) and an apical group (including other paralogs identified) [Fig. 6c]. The latter cluster also contained some GGPP synthase paralogs obtained from the termitid *Nasutitermes takasagoensis* (34)

**Figure 6:**
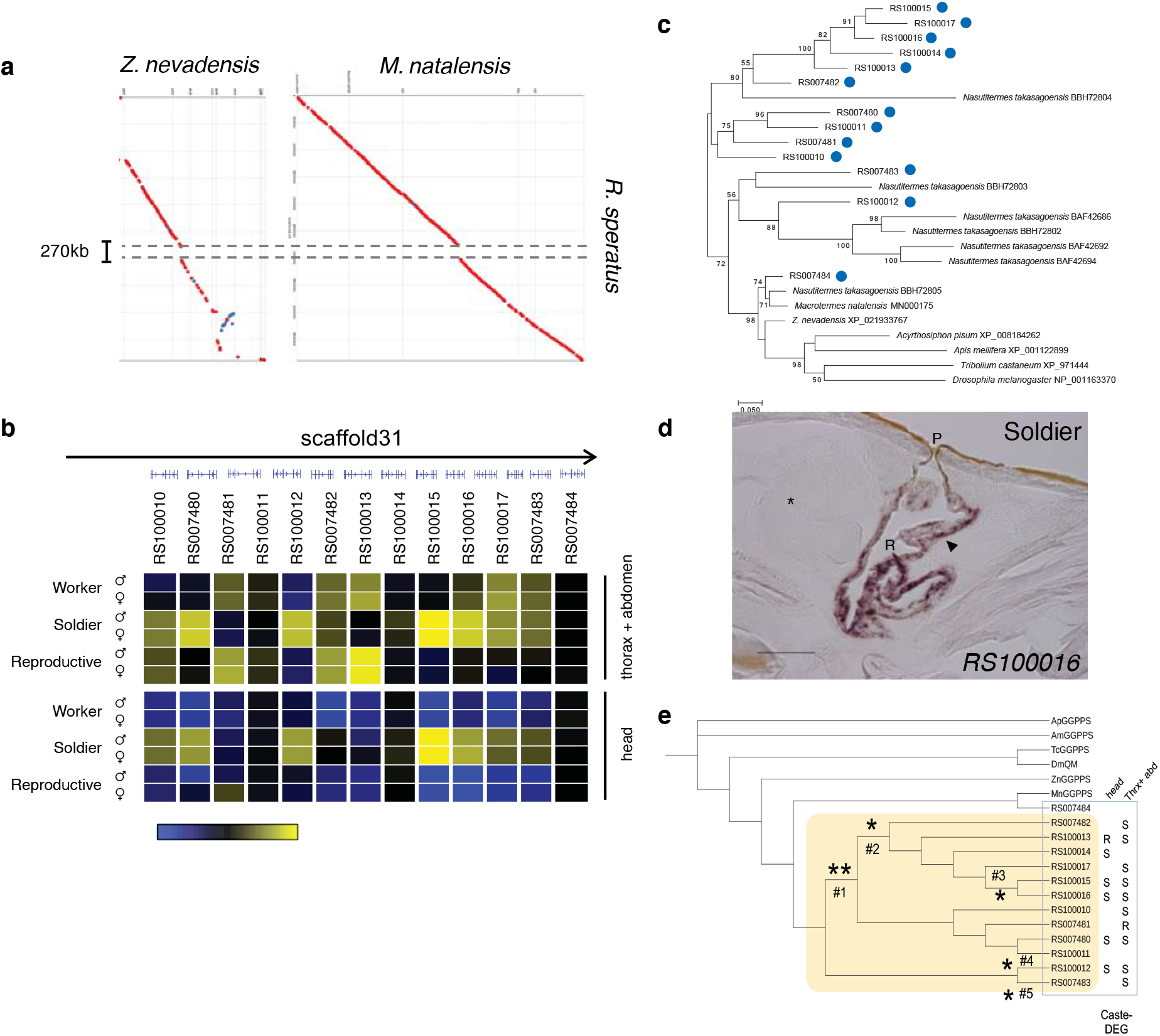
Geranylgeranyl diphosphate (GGPP) synthase homologs in the *R. speratus* genome. **(a)** Synteny comparison around GGPP synthase loci among three termites, *R. speratus, M. natalensis* and *Z. nevadensis. R. speratus*-specific insertions were found, where GGPP synthase paralogs were tandemly duplicated in the *R. speratus* genome. **(b)** Genomic location and gene expression of *R. speratus* GGPPS homologs. The heatmap shows the expression level calculated by mean-centered log(RPKM+1). Yellow indicates high expression, while blue denotes low expression. Black represents the mean level of expression among the castes. Note that the heatmap of *RS007484* is almost entirely black for all samples, which indicates that expression was invariable among castes, while most of the rest of the paralogs showed caste-biased expression. **(c)** ML tree of GGPP synthase homologs with a LG+G model. *R. speratus* genes are marked with blue circles. **(d)** Vertical cryosection of the soldier head subjected to *in situ* hybridization for *RS100016* mRNA. The front of the head is on the left side. The gland cell layer surrounding the frontal gland reservoir (R) is stained dark (arrowhead). The asterisk indicates the brain. The frontal pore (P) discharging frontal gland secretion is also observed. Bar = 0.1 mm. See Supplementary Fig. 7c for the negative control experiment. **(e)** Molecular evolutionary analysis of *R. speratus* GGPP synthase homologs by the PAML branch-site test. Detected positive selection is marked with a single asterisk * (p < 0.05) or double asterisks ** (p < 0.01) next to the corresponding branches.

Transcriptome data indicated that all of the GGPP synthase genes, except *RS007484* which was a member of the ancestral group in the phylogenetic tree, showed caste-biased expression, and caste specificity varied across the paralogs [Fig 6b]. Specifically, *RS100010, RS007480, RS100012, RS100015, RS100016, RS100017* and *RS007483* showed soldier-specific expression, while *RS007481, RS007482* and *RS100013* showed reproductive-specific expression [Fig 6b]. Several GGPP synthase genes have been identified in some termite species and are known to function in a caste-specific manner; for example, the soldiers of *N. takasagoensis* synthesize defensive polycyclic diterpenes by high expression of the GGPP synthase gene in the frontal gland to use chemical defense (62). It has been reported that the soldiers of *Reticulitermes* have a frontal gland in which diterpenes are synthesized, although the biological role is not fully understood (63–65). Consequently, it is possible that the soldier-specific GGPP synthases identified to date are involved in chemical defense. Indeed, *in situ* hybridization revealed that the soldier-specific GGPP synthase *RS100016* was expressed exclusively in the soldier frontal gland, as shown in a previous study (66) [Fig. 6d, Supplementary Fig. 7c]. It is also possible that reproductive-specific GGPP synthases are involved in the metabolism of other diterpenes, such as pheromone synthesis, especially *RS007481*, which shows strong queen-specific expression in the thorax and abdomen and may play a role in the synthesis of queen substances.

Under the branch-site (BS) model of codon substitutions (67), significant positive selection was detected on five branches of *R. speratus* GGPP synthase family tree [Fig. 6e]: ancestral branches #1 and #2, and the branches leading to RS100017 (branch #3), RS100012 (branch #4) and RS007483 (branch #5). These results suggest that all GGPP synthase paralogs of *R. speratus* except the ancestral type *RS007484* have experienced positive selection and finally acquired novel roles for the production of defensive and/or pheromonal substances.

### The TY family, a novel gene family restricted to termites

Numerous studies have shown that novel genes (e.g., TRG) play important roles in the evolution of novel social phenotypes in hymenopteran social insects (8, 68, 69). We found that termite-shared TRGs showed strong enrichment for caste-DEGs (see above). A striking example of caste-biased TRGs is a tandem array of three novel genes [Fig. 7a; Supplementary Table 11], *RS001196, RS001197* and *RS001198*, that have no significant homologs in any organisms outside termite clades. These three genes were expressed at extremely high levels (up to 250,000 RPKM), which constituted approximately 30% of the worker head transcriptome, and strongly biased across the three castes [Fig. 7a]. Each gene was composed of a single exon encoding a short peptide ∼60 aa in length that contained a secretion signal peptide in the N-terminal region followed by a middle part rich in charged amino acid residues and C-terminal part rich in polar amino acids with unusually high number of tyrosine residues [Fig. 7b]. Here, we named this novel class of peptides the termite-specific tyrosine-rich peptide family (TY family). The three TY genes showed modest sequence similarity with each other, suggesting that they are paralogs derived by tandem duplication. TY family orthologs were also found in the genomes of *Z. nevadensis* and *M. natalensis* [Fig. 7b]. We estimated pairwise evolutionary rates (the ratio of nonsynonymous to synonymous substitutions, i.e., dN/dS) between *R. speratus* and *Z. nevadensis* for these three peptides. The dN/dS for each gene ranged from 0.03 to 0.16 (Fig. 7c), indicating that they evolved under strong purifying selection and suggesting a conserved function in the termite lineage. Indeed, *Z. nevadensis* orthologs were also expressed at a high level in the soldier and worker castes in a pattern similar to that in *R. speratus*.

**Figure 7:**
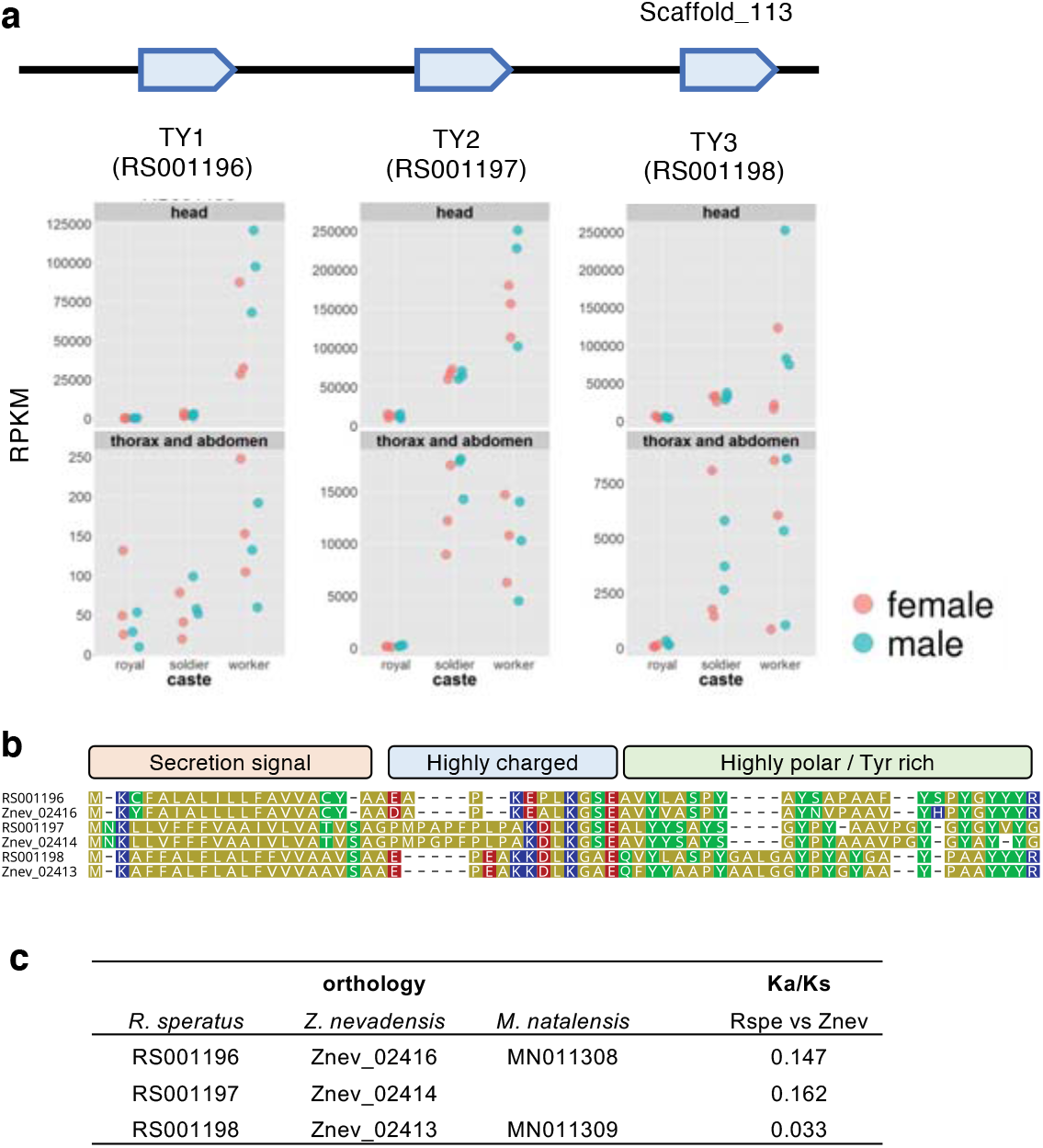
The TY family, a novel secretion gene family identified from termite taxonomically restricted genes. **(a)** Genomic locations and caste-biased expression patterns of TY family genes. **(b)** Multiple alignment of TY homologs of *R. speratus* and *Z. nevadensis*. Protein motifs and structural characteristics are represented. **(c)** Orthology of TY homologs in three termites and the results of the Ka/Ks analysis.

### Facilitation of caste specification by gene duplication

Recent advances in sociogenomics in different social insects are promoting our understanding of the genetic bases of social evolution, which include the co-option of genetic toolkits of conserved genes, changes in protein-coding genes, cis-regulatory evolution leading to genetic network reconstruction, epigenetic modifications and TRGs (15, 70). In addition to these components, our genomic and transcriptomic analyses in *R. speratus* highlighted the significance of gene duplication for caste specialization. Gene duplication is, in general, a key source of genetic innovation that plays a role in the evolution of phenotypic complexity; gene duplication allows for subsequent divergent evolution of the resultant gene copies, enabling evolutionary innovations in protein functions and/or expression patterns (71–73). Regarding eusocial evolution in insects, Gadagkar (74) first pointed out the importance of gene duplication; ‘genetic release followed by diversifying evolution’ made possible the appearance of multiple caste phenotypes in social insects. Many decades later, genomic analyses revealed gene family expansion, especially in relation to chemical communication, in both ants (odorant receptors; (75–77)) and termites (ionotropic receptors; (18)). Based on Godagkar’s hypothesis, duplicated genes can be released from the constraints of original selection, leading to new directional evolution, i.e., for caste-specific functions (e.g., queen- or worker-trait genes). However, the detailed roles and significance of gene duplication in social evolution have been elusive.

This study revealed that gene duplication associated with caste-biased gene expression is prevalent in the *R. speratus* genome. The list of duplicated genes encompasses a wide array of functional categories related to the social behaviors in termites as exemplified by transporters such as lipocalins (communication and physiological signaling; cf. (31, 44)), digestive enzymes such as carbohydrate-active enzymes, immune-related genes such as lysozymes (social immunity), and metabolic enzymes such as GGPP synthase (social defense). This study demonstrated that caste-specific expression patterns differed among in-paralogs.

Although such paralogous genes were often observed in tandem in the genome, the expression patterns were often independent from one another, showing differential caste biases in many cases. Additionally, discordant caste biases in transcriptional expression were observed among closely related paralogs with similar coding sequences, as represented by little correlation between phylogenetic position and caste specificity (Fig. 3a). Although the regulatory and evolutionary mechanisms underlying caste-biased expression patterns are elusive, these examples strongly suggest that gene duplications have facilitated caste specialization, leading to social evolution in termites.

After the gain of caste-biased gene regulation, subfunctionalization and/or neofunctionalization seems to have occurred, leading to caste-specific expression and caste-specialized functions. For example, in the case of lipocalin family, lipocalin paralogs were generated by lineage-specific functional expansion in caste-specific organs or tissues: a queen-specific lipocalin (*RS008881*) was expressed specifically in the ovarian accessory glands, while a soldier-biased lipocalin (*RS008823*), was expressed exclusively in the frontal glands in soldier heads [Fig. 3de]. Taken together, we hypothesize that, in termites, caste specification through gene duplication proceeds by the following three steps: 1) gene family expansion by tandem gene duplication, 2) regulatory diversification leading to an expression pattern restricted to a certain caste, and 3) subfunctionalization and/or neofunctionalization of the gene products conferring caste-specific functions. As an exaptation of these steps, the case in which one (or some) of the multiple functions of pleiotropic genes are allocated and specialized to a duplicated gene copy might have led to caste-specific subfunctionalization (38, 39).

Recently, it was suggested that the evolution of phenotypic differences among castes in the honey bee was associated with the gene duplication, by showing that duplicated genes had higher levels of caste-biased expression compared to singleton genes (78). It was also shown that the level of gene duplication was correlated with social complexity in bees (superfamily Apoidea) (78). Given the independent origin of eusociality in termites and honeybees, gene duplications might be a shared mechanism facilitating the evolution of caste systems in social insects.

## Materials and Methods

### Insects

All mature colonies of *Reticulitermes speratus* used for genome, RNA, and Bisulfite sequencing (BS-seq), were collected in Furudo, Toyama Prefecture, Japan [Supplementary Table 1]. Detailed sample information is described in *SI Appendix, Supplementary Methodology*.

### Sample collection, genome sequencing and assembly

All colonies of *Reticulitermes speratus* used for genome, RNA, and bisulfite sequencing were collected at Furudo, Toyama Prefecture, Japan.Detailed sample information is described in Supplementary Table 1 and *SI Appendix, Supplementary Methodology*.

We used female secondary reproductives (nymphoids I and II) for genome sequencing. We excluded the gut and ovaries of nymphoids to avoid contamination by DNAs from the king or other microorganisms. Genomic DNA was isolated from each individual using a Genomic-tip 20/G (Qiagen). We used 5 microsatellite loci (Rf6-1, Rf21-1, Rf24-2, Rs02, and Rs03) to confirm whether they were homozygous at these loci and shared the same genotype. The purified genomic DNA purified was fragmented with a Covaris S2 sonicator (Covaris), size-selected with BluePippin (Sage Science), and then used to create two pair-end libraries using a TruSeq DNA Sample Preparation Kit (Illumina) with insert sizes of ∼250 and ∼800 bp [Supplementary Table 3]. Four Mate-pair libraries with peaks at ∼3 kb, ∼5 kb, ∼8 kb and ∼10 kb, respectively, were also created using a Nextera Mate Pair Sample Preparation Kit (Illumina) [Supplementary Table 3].These libraries were sequenced using an Illumina HiSeq system with 2 × 151 bp paired-end sequencing protocol. Reads of the pair-end and mate-pair libraries were assembled using ALLPATHS-LG (build# 47878), with default parameters. BUSCO v4.0.6(29)(https://busco.ezlab.org/) was used in quantitative measuring for the assessment of genome assembly using insecta_odb10 as the lineage input. A genome browser was built using JBrowse (https://jbrowse.org/)

### Gene prediction

A protein-coding gene reference set was generated with two main sources of evidence, aligned *R. speratus* transcripts and aligned homologous proteins of other insects, and a set of *ab initio* gene predictions. RNA-seq reads were assembled *de novo* using Trinity, and then mapped to the genome using Exonerate. We processed homology evidence at the protein level using the reference proteomes of 7 sequenced insects including *Z. nevadensis* and Blattodea protein sequences predicted from RNA-seq of *Periplaneta americana* and *Nasutitermes takasagoensis*. These proteins were split-mapped to the *R. speratus* genome with Exonerate. These models were merged using the EvidenceModeler (EVM), which yielded 15584 gene models. Seventy-four genes were manually inspected and corrected. In particular, tandemly duplicated genes were liable to be incorrect gene prediction with erroneous exon–exon connections across homologs. The final set of 15591 genes was designated as Rspe OGS1.0 [Supplementary Data 2 (DOI:10.6084/m9.figshare.14267381)]. The quality of theOGS1.0 was evaluated by assessing two types of evidence, homology and expression. Among 15591 genes, 12996 (83.3%) showed any hits in the NCBI nr database, 10440 (70.0%) included known protein motifs defined in the Pfam database, and 14302 (91.7%) showed evidence of expression with a threshold of RPKM = 1.0 in any sample of caste-specific RNA-seq data. In sum, 15577 (99.9%) had evidence for the presence of homologs and/or expression.

### Orthology inference and gene duplication analysis

Orthology determination among three termites: Orthologous genes among the proteomes of three termite species, *R. speratus, Z. nevadensis*, and *M. natalensis* (gene models RspeOGS1.0, ZnevOGSv2.229, and MnatOGS3, respectively), were determined by pairwise comparisons with InParanoid v4.1 followed by three-species comparison with MultiParanoid. *M. natalensis* gene set, MnatOGS3, was built in this study using a similar pipeline as used for *R. speratus*.

Ortholog analysis with arthropod proteomes: Orthology relationships of *R. speratus* genes (OGS1.0) with other arthropod genes were analyzed by referring to the OrthoDB gene orthology database ver.8 (87 arthropod species) (https://www.orthodb.org/). We grouped *R. speratus* genes with the OrthoDB ortholog group using a two-step clustering procedure. For each *R. speratus* protein, BLASTP was used to find similar proteins among the arthropod proteins, and the ortholog group of the top hit was provisionally assigned to the query *R. speratus* gene. Then, the ortholog grouping was evaluated by comparing the similarity level (BLAST bit score) among members within the focal ortholog group. We keep the grouping if the BLAST bit score between the query *R. speratus* gene and top arthropod gene was higher than the minimal score within the original cluster members. Among 15591 *R. speratus* OGS1.0 genes, 12434 were clustered into 9033 OrthoDB Arthropod ortholog groups. Gene duplication was assessed based on this clustering. If two or more members of one species were included in a single ortholog group, they were regarded as a multigene family.

### RNA-seq

W4–5 workers (old workers) and soldiers were collected from each colony. To collect primary reproductives, dealated adults were chosen randomly from each colony in accordance with the method of the previous literature (79), and female– male pairs were mated (Supplementary Table 1). Kings and queens were sampled after 4 months. Each individual was divided into head and body parts (thorax + abdomen). We prepared RNA-seq libraries for 12 categories based on castes (reproductives, workers and soldiers), sexes (males and females) and body parts (head, and thorax + abdomen). Three biological replications of the 12 categories were made with three different field colonies totaling 36 RNA-seq libraries [Supplementary Table 2]. All Illumina libraries prepared using a TruSeq Stranded mRNA Library Prep kit were subjected to a single-end sequencing of 101 bp fragments on HiSeq 2500. The cleaned reads were mapped onto the genome with TopHat v2.1.0 guided by the OGS1.0 gene models. Transcript abundances were estimated using featureCounts and normalized with the trimmed mean of M-values (TMM) algorism in edgeR. Differentially expressed genes among castes and between sexes were detected in each body part (head / thorax and abdomen) using a generalized linear model with two factors, namely, caste and sex using edgeR with the conditions set as false discovery rate (FDR) < 0.01 and the log2 fold change of the expression level > 1.

## Data Availability

Data from whole-genome sequencing, transcriptome sequencing, and methylome sequencing have been deposited in the DDBJ database under BioProject accessions PRJDB2984, PRJDB5589 and PRJDB11323, respectively. The analyzed data including genome assembly, gene prediction, annotation, and gene expression are available through FigShare (https://doi.org/10.6084/m9.figshare.c.5483235). The *R. speratus* genome browser is available at http://www.termite.nibb.info/retsp/.

## Code availability

Custom R and Ruby scripts were deposited into Github (https://github.com/termiteg/retsp_genome_paper).

## Acknowledgments

We thank R. H. Suzuki and A. Karasawa for experimental support, T. Nishiyama and M. Hasebe for discussion on genome analyses, N. Kanasaki and K. Kai for rearing insects, T. Shibata, S. Ohi, T. Aizu, H. Ishizaki, H. Asao for next-generation sequencing (NGS), and K. Yamaguchi for NGS data management. Computations were partially performed on the supercomputers at the Data Integration and Analysis Facility, National Institute for Basic Biology. This study was funded by the JSPS/MEXT KAKENHI Grant Numbers 25128705, 24570022, 16K07511, JP19H03273, 22128008, 19K22294, 221S0002 and NIBB Collaborative Research Programs (20-323).

## Author contributions

S.S., Y.H., T.M., and K.M. designed and managed the project. D.W., K.T., R.S., H.Y., Y.M., R.S., and K.M. collected samples. D.W., R.S., Y.M., and R.S. performed the DNA extraction. S.S., and A.T. performed the library construction and genome sequencing. Y.H., D.W., K.T., R.S., H.Y., and Y.M. generated the RNA-Seq data. S.S., Y.H., and R.S. generated the BS-Seq data. S.S., Y.H., D.W., G.T., M.Y.H, K.T., M.M., Y.S., K.O., T.N., H.G., M.K.H., and S.M. contributed to the genome assembly and annotation. S.Su, and M.K. performed histological analyses. S.S., Y.H., G.T., T.M., and K.M. drafted the manuscript. All authors contributed to the final version of the manuscript.

