## Supplementary Information for "Genomic and transcriptomic analyses of the subterranean termite *Reticulitermes speratus:* gene duplication facilitates social evolution"

**This PDF file includes:**

Supplementary text
Figures S1 to S36
Tables S1 to S28
Legends for Datasets S1
SI References

**Other supplementary materials for this manuscript include the following:**

Datasets S1

### Supplementary Information Text

#### Supplementary Note: Genes involved in specific biological functions

Following the prediction of genes encoded in the *Reticulitermes speratus* genome, we manually annotated and investigated the genes of some functional categories that characterize the ecology, evolution, behavior, development and physiology of the subterranean termite. We analyzed 15 categories, among which lipocalin, glycoside hydrolase family, lysozyme family, geranylgeranyl diphosphate (GGPP) synthase and the novel secretion gene family TY are described in the main text. Here, the other 10 categories (sex determination; epigenetics; chemosensory genes; biogenic amines and neuropeptides; juvenile hormone-related genes; ecdysone-related genes; insulin/insulin-like signaling pathway; toolkit genes involved in wing formation; immunity; insecticide target and detoxification genes) are described. Finally, a report on a caste-specific expression of microRNAs (miRNAs) is introduced.

#### Sex determination

In insects, sex determination is cell-autonomously controlled by a cascade of RNA splicing, in which mRNAs are alternatively spliced in a sex-specific manner. Although the upstream genes in this cascade differ among taxa, the most downstream key gene *doublesex* (*dsx*) is conserved among insects, and also in some of crustaceans and chelicerates[1–4]. Since *dsx* encodes sex-specific transcription factors, the cascade results in sex-specific transcriptions of downstream genes that are responsible for sex differentiation. In contrast to holometabolous insects, the sex determination cascades in hemimetabolous insects including termites are scarcely understood[5,6]. Thus, our genome research in *Reticulitermes speratus* provides important information on gene repertories related to the sex determination cascades in hemimetabolous insects.

The sex determination cascade might also be responsible for social organization in termites, since some termite species show sex-specific or sex-biased caste ratio, suggesting that some regulatory factors for caste differentiation sexually differ[7]. In *R. speratus*, sex ratio of workers is known to be nearly equal, whereas those of nymph and soldier are female-biased[8]. For example, differentiation into female secondary reproductives was inhibited by the pheromone derived from female primary reproductives[9], suggesting that they have the reaction mechanism to sex-specific pheromones regulating the ergatoid differentiation. Therefore, the sex determination cascade is one of the most likely candidate regulatory mechanisms for the sex-specific caste differentiation. Thus, the annotation of sex-determination genes could also help to understand the regulatory mechanisms underlying the eusociality in termites. Here, orthologs of the genes reported as sex determination genes in other insect species were searched in the *R. speratus* genome, and compared the expression levels of those genes between sexes and among castes by transcriptomic analyses.

We selected the 27 candidate genes from the *Drosophila melanogaster* genes categorized as “sex determination (BSID: 492283)” in BioSystems database at NCBI (http://www.ncbi.nlm.nih.gov/biosystems/)[10] and selected 4 candidate genes based on previous studies on the sex determination in silkworm *Bombyx mori*[11–13]. Database searches were performed using full length of amino acid sequences of 27 *Drosophila* genes or 3 *Bombyx* genes against our *Reticulitermes* gene model database (RsGM8_pep) via BLASTP algorithm (See Supplementary Table 12 for the accession numbers of each query sequence). Searching for an ortholog of *Bombyx Feminizer*, which encodes microRNA, was performed via BLASTN algorithm. These analyses revealed that the *R. speratus* genome possessed orthologs of the 24 *Drosophila* genes and those of the 3 *Bombyx* genes (Supplementary Table 12). For the orthologs of major components of the sex determination cascade in *Drosophila*, orthologs of *Sex-lethal* (*Sxl*), *transformer* (*tra*), *transformer-2* (*tra2*) and *fruitless* (*fru*) were found, whereas, surprisingly, the *dsx* ortholog was not found. The BLAST search for *Drosophila dsx* as a query hit three orthologs of the *doublesex-mab3* related transcription factors genes (*Dmrt11B*, *Dmrt93B*, *Dmrt99B*), but not *dsx* ortholog. The insect *dsx* is a member of the *Dmrt* gene family that is conserved among a wide array of animal phyla[14]. Both *dsx* and *Dmrt* paralogs share a well conserved DNA-binding domain (DM domain), and some *Dmrt* genes also plays essential roles in gonad development and sexual differentiation outside Insecta[14]. The *dsx* ortholog would have been lost in *R. speratus* genome, and any other factors, e.g., *Dmrt* genes, might be substituted as the most downstream genes in their sex determination cascade. Alternatively, because the German cockroach *Blattella germanica* has the conserved *dsx* ortholog[6], the domain sequences may have diverged during the course of termite evolution.

The *R. speratus* orthologs of 3 regulatory genes (*deadpan*, *groucho*, *scute*) for splicing of the most upstream gene (*Sxl*) in the *Drosophila* cascade were duplicated, whereas no ortholog of two (*sisterless-A* and *degringolade*) of them were found. An ortholog of *stand still*, required for the *Drosophila* germline sex determination, was neither found. Additionally, the ortholog of *B. mori Feminizer* was not found, while the others coding proteins were found in the *R. speratus* genome. Although a primary signal for sex determination in *R. speratus* was unknown, the signal should be different from those in *D. melanogaster* (the dose of X-linked signal element[15]) and *B. mori* (*Feminizer* piRNA on the W chromosome[11]).

RNA-seq analysis revealed the expression patterns for 25 out of 30 candidate orthologs (Supplementary Table 12). These data were compared the expression levels between sexes and among three castes (primary reproductives, soldiers and workers) in two body parts [heads and the remaining parts (thorax + abdomen)] (biological triplicates; NCBI BioProject Accession No. PRJDB5589). Statistical analysis revealed that 14 out of 25 orthologs showed caste-biased expression patterns while none showed sex-biased (Supplementary Table 12). For example, the expressions of *Dmrt11* orthologs (RS007930) were higher in reproductive heads, moderate in worker heads, and lower in soldier heads (FDR = 7.85E-11, GLM, Supplementary Fig. 9), the orthologs of *outstreched* (*os* RS015475, FDR = 1.87E-08) and *ovarian tumor* (*out*, RS009292, 8.00E-07) were highly expressed in female reproductive bodies (Supplementary Fig. 9). It was suggested that these orthologs were involved in their sex determination or the downstream pathways of sex differentiation.

Our analyses revealed that most orthologs of sex determination genes identified in other insects, were conserved in the *R. speratus* genome, and that some of them were expressed in a cased-biased manner. However, it remains unknown which of them play roles in their sex determination cascade. In order to test their roles for sex determination, it should be examined whether these genes were spliced in a sex-specific manner, and whether these genes actually regulate their sex-specific trait expressions.

**Epigenetics: Histone modifying enzymes**

Histone posttranslational modifications (PTMs), one of the epigenetic mechanisms, play important roles in gene-expression regulations without genomic changes, resulting in alteration of development and behavior in various organisms[16–18]. Since such PTMs can be changed in response to environmental stimuli[16,19], polyphenic developments in insects, including caste differentiation in social insects, are considered to be affected by PTMs[20–23]. For example, when honeybee larvae were fed with royal jelly, which contain (E)-10-hydroxy-2-decenoic acid possessing histone deacetylase inhibitor activity, such larvae differentiate into queen bees[24]. Furthermore, an association between PTM (especially, histone acetylation and methylation) patterns and caste identities was shown in the carpenter ant *Camponotus floridanus*[25]. It suggests that histone acetylation and methylation have important roles in the regulation of caste differentiation in social hymenopterans. However, in termites, such roles of PTMs remain unknown. Here, as a first step of understanding the PTM roles in the caste differentiation in *R. speratus*, we searched four types of histone-modifying enzyme genes, i.e., genes encoding histone acetyltransferases (HATs), histone deacetylases (HDACs), histone methyltransferases and histone demethylase. In addition, we examined expression patterns of those genes between sexes and among castes by an RNA-seq analysis.

The *R. speratus* gene models were searched for histone acetyltransferases, deacetylases, methyltransferases, and demethylases with BLASTP algorithm (E-value cutoff of 1e-10)[26] using query protein sequences of those enzymes derived from *Zootermopsis nevadensis*[27] which belongs to Archotermopsidae and is phylogenetically distant to *R. speratus,* and *D. melanogaster.* We also investigated the expression levels of those genes by an RNA-seq analysis (for the method of the RNA-seq analysis, see main text).

We identified 69 histone-modifying enzyme genes in *R. speratus* (17 histone acetyltransferase genes, 13 histone deacetylase genes, 26 histone methyltransferase genes, 13 histone demethylase genes; Supplementary Table 13). Repertoire of these genes was almost identical between *R. speratus* and *Z. nevadensis* (Supplementary Table 13), suggesting that the repertoire have been highly conserved in the course of termite diversification.

Our RNA-seq analyses revealed that 47 of 69 genes showed caste-biased expression patterns while all of the 69 genes showed no significant differences in expression levels between sexes (FDR < 0.05, Supplementary Fig. 10 and Supplementary Table 13). Interestingly, *KDM1A* (*Lsd1*, coding histone demethylase gene) and *SETDB1* (*eggless*, coding histone methyltransferase gene), both of which regulate oogenesis in *Drosophila melanogaster*[28–30], were highly expressed in thorax and abdomen of queens (Supplementary Fig. 10). The ovarian development in queens is thus suggested to be regulated by PTMs, especially histone methylation and demethylation. Furthermore, while *DOT1L* (*grappa*, coding histone methyltransferase gene) is required for stress resistance in *D. melanogaster* (List et al. 2009), *grappa* expression levels were relatively high in soldiers of *R. speratus* throughout the body (Supplementary Fig. 10). It was suggested that the enhanced stress resistance might be required for the colony defense by soldiers.

Generally, aging is considered to be linked to the sirtuin lysine deacetylases enzymes (*SIRT*), and some genes encoding SIRT enzymes are highly expressed in long-lived reproductive castes of an ant and a termite (*SIRT1* and *SIRT6* in *Harpegnathos saltator*; *SIRT6* and *SIRT7* in *Z. nevadensis*)[27,31]. In *R. speratus*, expression levels of *SIRT6* and *SIRT7* were higher in the body (thorax + abdomen) of female primary reproductives (queens) than in those of male reproductives (kings), workers and soldiers (Supplementary Fig. 10). However, queens derived from alates (=winged imagos) are suggested to live rather shorter than kings[32]. Therefore, at least in *R. speratus*, *SIRT6* and *SIRT7* might contribute to egg-laying but not to the life-span elongation.

These results suggest that the caste-biased expressions of genes encoding histone modifying enzymes would determine the caste identities through caste-specific PTMs in *R. speratus*. Therefore, such caste-specific PTMs must be caused during the caste differentiation process in their postembryonic development. Elucidation of the regulatory mechanisms for caste-specific PTMs will help us to understand how social organizations of termites are maintained.

**Epigenetics: DNA methylation**

DNA methylation, the addition of methyl groups to DNA bases, is one of the most important regulatory mechanisms of gene expression. The regulation of gene expression by DNA methylation is considered to be a candidate for a proximate mechanism to produce insect phenotypic plasticity[33,34]. Indeed, in the honey bee *Apis mellifera*, regulation of DNA methylation has been shown to take an important role in the caste differentiation of queens and workers[35]. Because termites exhibit relatively high levels of DNA methylation among insects[36,37] (also see the main text), DNA methylation should get more attention in termite biology.

To date, some major factors regulating DNA methylation status have been reported. DNA (cytosine-5)-methyltransferase 1 (DNMT1) methylates the replicated DNA strand based on the methylation status of the template DNA[38,39] and DNA (cytosine-5)- methyltransferase 3 (DNMT3) adds a methyl group to the unmethylated DNA region[40,41]. Ten-eleven translocation methylcytosine dioxygenases (TETs) are involved in the oxidative demethylation of 5-methylcytosine, and thymine-DNA glycosylase (TDG) also contribute to DNA demethylation through the process of base excision repair[42]. Proteins of methyl-CpG binding domain (MBD) family are capable of binding to methylated CpGs, and involved in a gene expression regulation machinery[43,44]. It has been suggested based on transcriptome sequencing data that *DNMT1*, *DNMT3*, *MBD* and *TET* genes are present[45,46] in the genome of *Reticulitermes speratus*, but their copy numbers and gene expression levels in different castes still remain unknown. In addition to those gene, the copy number and expression level of the *TDG* gene should also be examined to accelerates the study on DNA methylation in *R. speratus*.

To identify those genes, we carried out BLASTP searches against the gene models and TBLASTN searches against the genome assembly of *R. speratus*. As query sequences for the BLAST searches, we used protein sequences of the DNMT1, DNMT3 and MBD genes identified in other insect species (listed in Hayashi et al.[45]) and those of the TET and TDG genes of *Drosophila melanogaster* (NCBI accession numbers of NP_001261344.1 and NP_651925.1, respectively). We applied an E-value threshold of 1e-10 for the BLAST searches. In addition to *R. speratus*, we performed the BLAST searches for those genes against genomic and protein sequence data of *Blattella germanica*[47], *Periplaneta americana*[48], *Zootermopsis nevadensis*[27], *Macrotermes natalensis* [49] and *Coptotermes formosanus*[50]. Furthermore, we also performed mapping of the query sequences of the BLAST searches onto the genome assemblies of the blattodean species using Exonerate[51].

We found one copy of each of those genes in *R. speratus* (Supplementary Table 14), meaning that *R. speratus* preserves the major component of DNA methylation and demethylation. The other termites examined in this study were also showed to possess all of the genes. However, *DNMT3* was not found from two cockroach species, *B. germanica* and *P. americana*. It has been known that lineage-specific duplications and deletions in *DNMT* have occurred independently in various taxa (reviewed in Glastad et al.[34]). *DNMT3* has been lost in some cockroach lineages, while preserved in termite genomes, suggesting that *DNMT3* preservation is important for the evolution and maintenance of social life. Further studies are required on the *DNMT3* functions to clarify this hypothesis. Our RNA-seq analysis revealed that *MBD-like* and *MBD-R2* differentially expressed among castes both in head and in thorax and abdomen (Supplementary Fig. 11). The expression level of *DNMT3* in head and that of *TDG* in thorax and abdomen were also significantly different among castes (Supplementary Fig. 11). Further researches are required to examine if expression differences in those genes contribute to maintaining the caste identity and caste differentiation.

**Chemosensory genes**

Social insects use diverse chemical signals to communicate various information and maintain their colonies. In insects, chemical compounds are detected by receptor proteins expressed in olfactory and gustatory receptor neurons. Three classes of chemosensory receptors are known in insects. Odorant receptors (ORs) and gustatory receptors (GRs) are seven transmembrane proteins and generally expressed in the chemosensory appendages[52,53]. The ionotropic receptors (IRs) were found most recently and are members of the ionotropic glutamate receptor family[54].

Sensory neurons are surrounded by a hydrophilic sensillar lymph. Many odorants are hydrophobic, so that these water-insoluble lipophilic compounds require water-soluble carrier proteins to access the membrane receptors of sensory neurons[55]. A variety of proteins in the sensillar lymph are known to involve in this process, including odorant binding proteins (OBPs) and chemosensory proteins (CSPs). These proteins help to solubilize hydrophobic chemicals and to transport specific ligands to receptor proteins involving odor detection, discrimination and coding[56,57]. Sensory neuron membrane proteins (SNMPs) are transmembrane proteins belonging to the CD36 protein family. SNMPs are expressed in olfactory sensory neurons, presumably supporting the ORs to capture odor molecules on the sensory neurons[58].

Recent comparative genomics of social insects reveals lineage specific expansion of chemosensory genes in social insects. In ants, ORs are diversified, showing largest repertoire in insects[59]. Subsets of OBPs and CSPs are specifically expressed in the antennae but others are expressed primarily in non-chemosensory tissues[60–65], suggesting that OBPs and CSPs are not restricted in chemosensory functions. In contrast to ants, the genome sequence of *Z. nevadensis* revealed chemoreceptor repertoire were expanded in IRs not in ORs, and almost all OBPs are expressed in antennae[27]. These results suggest that expansion of chemosensory gene family is crucial for their complex social lives, but interestingly, different classes of gene families are expanded in the independent social evolution in ants and termites. Here, we reported chemosensory gene repertoires of subterranean termites (*R. speratus*), as well as their gene expression patterns among castes (primary reproductives, soldiers, and workers) and between sexes (females and males) for CSPs and SNMP. Details of analyses for OBPs and receptor genes will be reported elsewhere.

We used blastp to search for models of chemosensory genes in *R. speratus* for using protein sequences of other insect species[27,66–72] as queries with an e-value cutoff of 1.0E−5. We also ran a HMM search using the OS-D superfamily (pfam03392), CD36 family (PF01130), 7tm Chemosensory receptor (pfam08395), Ligand-gated ion channel (pfam00060) and Ligated ion channel L-glutamate- and glycine-binding site (PF10613) as a query. For phylogenetic analyses, we produced an alignment with the E-INS-i strategy of MAFFT[73] using protein sequences of *D. melanogaster*, *A. mellifera*, *A. pisum*, *P. humanus*, *Z. nevadensis* and *R. speratus* for CSPs.

Ambiguous sections of the alignment were removed using trimAl (option-'gappyout')[74]. This alignment was used to produce a maximum likelihood phylogenetic tree using RAxML[75] with 100 bootstrap replicates.

For chemosensory receptor genes, we found 31 OR, 25 GR, and 92 IR candidates from automatically annotated gene models of *R. speratus* (Supplementary Table 15-17). For OR genes, the numbers were almost one-tenth of OR numbers in ant species and the about half of OR numbers reported in *Z. nevadensis*. As suggested by Terrapon et al.[27], OR gene repertoire might not be expanded as in ants. However, because of OR genes are difficult to be assembled and annotated automatically[69], it is possible that many OR as well as GR genes were overlooked in our gene models. Further intensive manual annotation is needed to confirm the OR gene repertoires in *Reticulitermes* termites. The IR family was most expanded receptor family in *R. speratus*. This is also true for *Z. nevadensis* genomes with two distinctive termite-specific expansion sub-families[27]. Although ligand-specificity is unknown in many IRs, this receptor family might have diverse function in termite society.

We found 5 SNMPs from *R. sepratus* and 3 SNMPs from *Z. nevadensis* gene models (Supplementary Table 18). RNA-seq analyses detected the expression of 3 SNMPs (*RspeSNMP1a*, *1b* and *2*), and they were expressed in both head and body (thorax + abdomen) parts (Supplementary Fig. 12). Although it is suggested that SNMPs were involved in sex pheromone reception[58], *R. speratus* SNMPs were not expressed differentially between sexes as well as among castes in the head part (Supplementary Fig. 13, FDR > 0.05). Therefore, SNMPs might be not involved in the sex pheromone perception in *R. speratus* termite.

For odorant carrier proteins, we found 10 CSPs from *R. speratus* gene models (Supplementary Table 19). We also found 10 CSPs from *Z. nevadensis* genome, and the numbers of *R. speratus* CSP genes are compatible with *Z. nevadensis*. Seven of 10 CSPs were completely modeled in *R. speratus*. In the ants CSPs are major antennal protein and there are number of CSP gene specifically expanded in ants[63,76]. The number of CSP genes was smaller in termite species than in ants and clear termite-specific expansion was not detected in the CSP phylogenetic trees (Supplementary Fig. 14). RNA-seq analyses revealed that 5 CSPs were mainly expressed in the head part but the others (2 CSPs) were mainly expressed in the body (thorax + abdomen) part (Supplementary Fig. 15), suggesting that the CSPs were not restricted to the peripheral chemosensory events. There were no sex-specific expression patterns of CSPs in the head parts. Among the 5 head-specific CSPs, 3 of them were differentially expressed among castes and were relatively highly expressed in the non-reproductive castes (soldiers and workers) compared with primary reproductives (Supplementary Fig. 16, FDR < 0.05). These 3 CSPs are likely to be involved in the communication related to social tasks such as foraging and colony defense.

**Biogenic amines and neuropeptides**

Biogenic amines, which include neurotransmitters, and neuropeptides regulates behaviors and physiological status of individuals. The genes encoding those peptides are widely conserved among insects[77,78]. The roles of those peptides in the regulation of behaviors and physiological status that are underpinned have been extensively examined in the honeybee *Apis mellifera* [79,80]. On the other hand, still remains much to be explored in termites, which have acquired eusociality independently of the honeybees. To further explore the roles of biogenic amines and neuropeptides in termites, we identified genes and analyzed gene expression patterns among castes in *R. speratus*.

Using the sequence information of biogenic amine-related genes[27] and *Z. nevadensis* neuropeptide genes[81] as queries, we carried out Blast searches for those genes against the gene models of *R. speratus*. As a result, there were basically no differences in gene repertoires and numbers between *Z. nevadensis* and *R. speratus* (Supplementary Table 20).

Expression levels of each gene were compared among three castes, two sexes, two body parts (head and thorax + abdomen) using RNA-seq data. The results showed that some identified genes were expressed in a caste-specific manner (Supplementary Fig. 17). Interestingly, high expression levels involved in dopamine biosynthesis, *Pale* (*RS010906*) and *Dopa decarboxylase* (*RS006642*), were observed in soldiers and workers than in primary reproductives (Supplementary Fig. 17). On the other hand, *Dopamine N acetyltransferase*

(*RS005696*), involved in the metabolism from dopamine to N-acetyldopamine[82], was highly expressed in soldiers (especially in heads; Supplementary Fig. 17). Consequently, intrinsic dopamine levels (probably in heads including brains) may be different among castes in *R. speratus*. Generally in insects, it is well known that species-specific behaviors, such as trophallactic behavior in ants and antipredator behavior in beetles, are affected by the dopamine levels[83–85]. In the yellow fever mosquito *Aedes aegypti*, dopamine was also involved in the cuticular sclerotization[86]. Indeed in termites, brain dopamine levels were significantly higher in soldiers (and soldier-destined individuals) than workers (and worker-destined individuals) in *Hodotermopsis sjostedti* and *Z. nevadensis*, respectively[87,88]. Consequently, there is a possibility that differences of dopamine levels are related to the caste-specific social behavior and morphology in *R. speratus*.

Next, three dopamine receptor genes (*Dop1-3*) were identified, among which different expression levels were observed only in *Dop1* (Supplementary Fig. 18). The expression level of *Dop1* in queens (thorax + abdomen) was much higher than that in other castes including kings (Supplementary Fig. 18). Because dopamine receptor genes were expressed in ovarian tissues in *A. mellifera*[89], *R. speratus Dop1* may also be expressed in queen ovaries and involved in the ovarian development. *Tyramine β hydroxylase* (*RS013347*) involved in octopamine biosynthesis was expressed a little bit higher in soldiers (thorax + abdomen) than the other castes (Supplementary Fig. 17). More clearly, higher expression levels of *Octopamine-Tyramine receptor* (*RS000810*) and *Octopamine receptor* (*RS008926*) were observed in soldier heads ( Supplementary Fig. 18). In *D. melanogaster*, RNAi of *Tyramine β hydroxylase* resulted in the decreases of aggression in both males and females[90] and locomotor speed induced by food deprivation[91]. In *H. sjostedti*, both octopamine and tyramine levels in brains/suboesophageal ganglion were higher in soldiers, and octopamine-tyramine neurons were specifically enlarged in soldiers[87]. Consequently, present results observed in *R. speratus* and previous studies performed in other species suggest that biogenic amines such as dopamine, octopamine and tyramine are crucial for termite soldier-specific roles.

Finally, we identified 30 neuropeptides-related genes in *R. speratus* genome. Expression levels of 15 genes were significantly different among castes (Supplementary Fig. 19). It should be noted that, in most cases, expression levels tended to be higher in heads than in thoraces and abdomens (refer to the RPKM values, Supplementary Fig. 19). Given that the neuropeptides have crucial roles for regulating a wide range of insect behaviors[92], these 15 genes identified are involved in caste-specific behaviors and physiological actions in *R. speratus*.

**Juvenile hormone-related genes**

Juvenile hormone (JH) is the central factor for polyphenisms seen in insects, including caste differentiation in social insects[93,94]. Since it has long been known that the transition of JH titer plays critical roles in the caste differentiation in termites[95], the factors up- and downstream of the JH action have been particularly focused, especially in lower termites (e.g. Miura and Maekawa[96]). For example, RNAi of a JH binding protein gene *Hexamerin* (*Hex*) promotes the presoldier molt in *Reticulitermes flavipes*, suggesting that the sequestration of JH is important for the soldier differentiation[97]. Moreover, expression patterns of JH biosynthesis genes have been elucidated during the presoldier molt under natural condition in *Z. nevadensis*[98]. The expression changes of two genes (*JHAMT* and *CYP15A1*), involved in the final steps of JH biosynthesis, are suggested to be crucial for the presoldier molt. RNAi for the receptor gene *Methoprene-torelant* (*Met*) was also investigated during soldier and neotenic differentiations in *Z. nevadensis* and *R. speratus*, respectively[99,100], suggesting that caste-specific morphogenesis (e.g., head and mandible enlargement in soldiers) and/or physiological changes (e.g., up-regulation of *Vitellogenin* in neotenics) are regulated under the JH action. Recently, based on the comparisons of expression patterns of JH-related genes between queens and workers in three species with genome information, Jongepier et al.[101] suggest that the JH action differs between lower and higher termites. However, the roles of JH-related genes have yet to be elucidated in higher termites. To clarify this issue, information of *R. speratus* is very important, because a sister group relationship between a clade containing *Reticulitermes* and the Termitidae (higher termites) was strongly supported by multiple previous studies on the molecular phylogeny[102–104]. Here, we identified JH-related genes in *R. speratus,* mainly based on the information of lower and higher termites (*Z. nevadensis* and *Macrotermes natalensis*)[27,49]. As the results of expression analyses of those genes among castes, we discuss about the roles of related genes and the diverse JH action in termites.

First, we identified 15 JH biosynthetic genes and 5 signaling genes (Supplementary Table 21). The gene repertoires and the numbers were conserved and essentially similar to those in *Z. nevadensis* and *M. natalensis*. As shown in other reports on hemimetabolous insects (Villalobos-Sambucaro et al. [105] 2015), two *Met* isoforms (*Met A* and *B*) were found in termites and cockroaches (Supplementary Fig. 20), although it was not clear whether there were any functional differences between the two isoforms. Our expression analyses using RNA-seq data showed that many genes in the early steps of the JH biosynthetic pathway (e.g. *HMGS1*, *HMGR*, *DD* and *IPPI*) were highly expressed in all the body parts of soldiers, i.e., in heads, thoraces and abdomens (Supplementary Fig. 21). Soldiers of this species have well-developed frontal glands in their heads and thoraces producing defensive substances including isoprenoids, which are synthesized via the early steps of the JH biosynthetic pathway (also known as the mevalonate pathway)[106]. Consequently, the gene up-regulations seen in soldiers (Supplementary Fig. 21) suggested to be responsible for the production of defensive chemicals in frontal glands. On the other hand, in the late steps of the JH biosynthetic pathway, *JHAMT* and *CYP15A1* were highly expressed in heads of primary reproductives (Supplementary Fig. 22). Up-regulation of these genes were reported to strongly correlate to JH levels in the locust *Schistocerca gregaria*[107]. According to the previous work[108], primary reproductives used for RNA-seq (4 months after colony foundation) may start to increase JH level for reproduction. The results suggest that high *JHAMT* and *CYP15A1* expression levels in heads are also related to these physiological changes in primary reproductives. Moreover, the expressions of most genes in the late steps, including *JHAMT* and *CYP15A1*, in workers tended to be higher than in soldiers, but the similar levels as those in primary reproductives (Supplementary Fig. 22). These tendencies in *R. speratus* are different from those in two lower termites, *Z. nevadensis* and *Cryptotermes secundus*, but similar to those in a higher termite *Macrotermes natalensis* [101]. Similar tendencies were also observed in the expression patterns of JH signaling genes (Supplementary Fig. 23). For example, up-regulations of *SRC* (also known as *taiman*) in soldiers and workers compared to primary reproductives were observed (Supplementary Fig. 23). Moreover, although relatively high expression levels of both *Kr-h1* and *Br-C* were observed in primary reproductives, *E93* was highly expressed in soldiers especially in heads (Supplementary Fig. 23). These tendencies were similar to those in the higher termite *M. natalensis*[101]. Overall, the similar expression patterns of JH-related genes between *R. speratus* and *M. natalensis* suggest that the change of JH signaling action is required for the evolution of complex caste system observed in both *Reticulitermes* spp. and higher termites, both of which possess the forked caste differentiation pathways[102–104].

Second, precursor and receptor of allatotropin/allatostatin were identified (Supplementary Table 21). High expression levels of queen *allatotropin receptor* and king *allatostatin precursor* were observed in thorax and abdomen samples (Supplementary Fig. 23). Both allatotropin and allatostatin are neuropeptides to regulate the JH biosynthesis in corpora allata, but shown to be highly expressed in pupal stage and tissue-specific patterns in the beetle *Tribolium castaneum*[109]. Further analyses should be performed to know biological significances on the up-regulations of allatotropin/allatostatin in termite reproductives.

Finally, genes for JH binding and degradation were identified (Supplementary Table 21). Both *Hex1* and *Hex2* were highly expressed in workers than those in primary reproductives and soldiers (Supplementary Fig. 24). These patterns are not contradict with those observed in the honeybee *Apis mellifera*, in which hexamerines were highly expressed in workers than queens[110]. Large numbers of *JH esterases* (*JHEs*) were obtained in *R. speratus* as in two other species with genome information. Molecular phylogenetic tree based on amino acid sequences of *JHEs* was constructed by MEGA7[111], using acethylcholin-esterase genes of *Drosophila melanogaster* and three termites as outgroups (Supplementary Fig. 25). The resultant phylogeny showed a specific clade containing termite JHEs. Generally in insects, JHEs are mainly produced in fat bodies and catalyze the hydrolysis of JH in hemolymph[112,113]. Consequently, different JHE actions in each individual are probably involved in JH titer changes that may lead caste differentiation. Interestingly, *RS001960*, *61*, *65-67* were included in the same scaffold 129, all of which were observed in the specific termite clade (Supplementary Fig. 25), were differently expressed among castes (Supplementary Fig. 24). Similarly, RS004712-13 (scaffold 20) and RS014537-38 (scaffold 83) were also differently expressed among castes (Supplementary Fig. 26). These may be reflected by gene duplication and neo-/sub-functionalization, as discussed in the main part of this paper. Further detailed expression and functional analyses should be required to clarify this possibility.

**Ecdysone-related genes**

Termite caste differentiation is deeply associated with molting events. Generally in insects, molting events are regulated by both juvenile hormone (JH) and 20-hydroxyecdysone (20E; active form of ecdysone)[114]. The gene responsible for the 20E biosynthesis and the signaling pathways have been well studied in some model insects (e.g. Niwa & Niwa[115]). Ecdysone is generally produced in the molt glands (also known as prothoracic glands) by a suite of enzymatic reactions, released into hemolymph and converted into 20E in peripheral target tissues.

Studies on the 20E roles in caste differentiation are very few in termites, compared to those of JH shown in the previous section. However, some recent literatures clarify the crucial roles of 20E signaling in the termite caste differentiation. For example, an artificial 20E application induced the worker-worker molt in *Reticulitermes speratus*[116], suggesting that there is a 20E signaling pathway to regulate the worker molt similar to the nymphal molt in other hemimetabolous insects. Moreover, for the soldier differentiation (worker-presoldier and presoldier-soldier molts), the ecdysone receptor (EcR) was shown to be activated in *Zootermopsis nevadensis*[117]. Based on the expression and function analyses of 20E signaling genes in *Z. nevadensis*, Masuoka et al.[118] suggest that there are two different 20E signaling pathways, one of which has a role for the molting to the next instar, and another provides a role for the soldier-specific morphogenesis. According to the reports on the expression patterns of some 20E-related genes selected in three termite species with genome information[47], similar queen-biased or worker-biased expression patterns were observed in the 20E biosynthesis genes. To know whether the 20E roles are common or diversified among termite species, especially between lower and higher termites, expression patterns of related genes should be clarified more in detail. Here, we identified 20E-related genes in *R. speratus* using the information of *Drosophila melanogaster* and *Bombyx mori*, and examined the expression patterns among castes.

We identified 20E biosynthesis (7), receptor (2) and signaling (11) genes from the genome of *R. speratus* (Supplementary Table 22). Although gene numbers and repertoires were essentially similar to those of *D. melanogaster* and *B. mori*, *Prothoracicotropic hormone* (*PTTH*), i.e., the conserved neuropeptide that activate molt glands, could not be observed in the current gene model (Rspe OGS1.0).

Expression analyses were performed using RNA-seq data. Huge differences of expression levels among three castes and two body parts (head and thorax + abdomen) were observed in many genes (Supplementary Fig. 27, 28). For the 20E biosynthesis genes, *neverland* (*RS010513*) and *shade* (*RS006327*) were highly expressed in thorax and abdomen, compared to those in head. The expression localization of *neverland* other than molt glands are completely unclear in termites. The final step of 20E biosynthesis (enzymatic activity of hydroxylation) is normally regulated by *shade* in the peripheral tissues including the fat body[115]. Up-regulation of *shade* were observed both in soldiers and workers, suggesting that 20E titer and signaling pathway activity are activated in those castes. Moreover, *phantom* (*RS002862*), *shadow* (*RS010451*) and *spook* (*RS010514*, primary reproductives and workers) were relatively highly expressed in the head part. These genes may be expressed mainly in the molt glands of termites and involved in the biosynthesis of ecdysone. Interestingly, these genes were also highly expressed in thorax and abdomen of queens (all three genes) and kings (*shadow*). It is known that ecdysone is produced in the reproductive organs, and involved in ovarian development, proliferation of spermatogonium and sperm formation in some insects[119,120]. Further expression and functional analyses should be performed to know whether *phantom*, *shadow* and *spook* are expressed in ovary and testis in termites.

Expression patterns of *EcR* (RS006194) and *USP* (RS005985) were essentially similar to each other, except for thorax and abdomen in primary reproductives. High expression levels of receptor genes in soldiers and workers (and *EcR* in queen thorax and abdomen) were not contradict with the results of 20E biosynthesis genes. Notable expression patterns of the 20E signaling genes were observed in *HR38* (*RS008487*) and *E93* (*RS003976*); both genes were highly expressed in soldiers (and also *HR38* in workers). In the honeybee *Apis mellifera*, up-regulations of 20E signaling genes were observed in workers (Kubo, 2012), and *HR38* was expressed in forager brains[121]. There is a possibility that *HR38*-related 20E activity is involved in caste-specific behaviors both in honeybees and termites. It should be noted that JH-Met-Kr-h1-E93 (MEKRE93) pathway has crucial roles in both hemi- and holometabolan metamorphoses[122–124]. Soldiers are the developmentally-terminal stage and cannot molt into the next instar. Further analyses should be performed to clarify the functional meanings of highly E93 expression in soldiers and the role of E93-related signalings on the soldier formation in termites.

**Insulin/insulin-like signaling pathway**

Insulin/insulin-like signaling (IIS) pathway is highly conserved from invertebrate to mammals. In insects, insulin-like peptides are involved in the pathway and serve as hormones and growth factors[125]. The insulin signaling plays an important role in caste differentiation of some social insects. In honeybees, the IIS pathway contributes to the caste differentiation and aging[126,127]. In a damp-wood termite *Hodotermopsis sjostedti*, this pathway is also shown to be involved in the soldier differentiation (Hattori et al[128]). Therefore, studies on IIS are thus fundamental to the better understanding on insect sociality. Here, we searched for the IIS genes[129] in the genome of *R. speratus,* and investigated the differential expression levels of those genes among castes and between sexesm based on the RNA-seq data (for details, see Methods in the main text).

We carried out BLASTP searches against the *R. speratus* gene models using the insulin signaling genes of *D. melanogaster*[129] (Supplementary Table 23), based on the sequence similarity (E-value cutoff of 1e-10) and phylogenetic relationships shown below. We identified all of the IIS components from the *R. speratus* genome, with some genes duplicated. We found that Ras85D (Ras1) genes were duplicated in termite/cockroach lineages (Supplementary Table 23, Supplementary Fig. 29). Except for termite/cockroach lineage, Ras1 gene duplication was observed only in the ponerine ant *Harpegnathos saltator* (Supplementary Fig. 29). We found three copies (*RS000922*, *RS007018*, and *RS007019*) of *InR* genes (Supplementary Fig. 30) in *R. speratus*. Three copies of *InR* are also found in *Z. nevadensis* (Xu and Zhang[130]). Increase of the copy number of *InR* might be involved in the social evolution in termites. On the other hand, the ImpL2 gene was absent in the genomes of *R. speratus* and cockroaches (Supplementary Table 23). The ImpL2 protein bind with the insulin-like peptides and inhibits the insulin signaling activity[131]. This suggests that the IIS regulation in termites and cockroachs differs from that in other insects.

Our RNA-seq analysis revealed that, including the above mentioned *Ras1* and *InR* genes, there were significant differences in gene expression levels among primary reproductives, workers and solders in heads (16 genes) and the thoraxes and abdomens (20 genes) (Supplementary Fig. 31). It is considered that most of the differentially expressed genes in the soldier heads are involved in the soldier-specific functions because the head of soldier exhibits the highly specialized external and internal morphology associated with defensive tasks with pheromonal exocrine glands[132,133]. The primary reproductives exhibited the highest expression levels in two copies (*RS000933* and *RS013615*) of *Ras1* in the thoraxes and abdomens (Supplementary Fig. 31), suggesting that these genes are involved in the gonad development. Two *InR* genes (*RS000922* and *RS007019*) were highly expressed in soldiers, whereas *RS007018* were highly expressed in workers (FDR < 0.05) (Supplementary Fig. 31). This means that different gene copies of *InR* could possess some caste-specific functions.

**Toolkit genes involved in wing formation**

The acquisition of wings is one of the largest events in the insect evolution, that have led the adaptive radiation in insects. To date, some genes involved in the wing formation have been identified in insects[134]. It has also been known that environmental factors can affect expression of those genes, contributing to the regulation of wing polyphenism (e.g. Hartfelder and Emlen[135], Xu et al.[136]). The wing polyphenism is thus a remarkable example of gene-environment interaction in development. In some insect species, for example ants[137] and aphids[138], association between expression of some homologs of *Drosophila* wing patterning genes and wing polyphenisms has been shown. Termites also exhibit wing polyphenism among castes; workers and soldiers do not have wings but nymphs, which develop into adults, possess wing buds[139].

To identify wing-development toolkit genes referred to in Abouheif and Wray[137] and Brisson et al.[138] in *R. speratus*, we performed BLASTP searches for those genes of *D. melanogaster* against the gene models of *R. speratus.* Gene orthology was confirmed by reciprocal BLAST between *R. speratus* and *D. melanogaster* genes, and by phylogenetic analyses. We found no duplications and losses in the wing-development toolkit genes of *R. speratus* (Supplementary Table 24).

Our RNA-seq data showed that the expression of *Daughters against dpp* (*Dad*) in both soldiers and workers were lower than that of primary reproductives (FDR < 0.05, Supplementary Fig. 32). *Dad* is induced by *Dpp*, and then act as the antagonist of Dpp52. Therefore, low expression of *Dad* in both soldiers and workers suggest the low activities of *Dpp* signaling. Although the *apterous* gene is suggested to be involved in aphid wing polyphenisms[138], no significant differences were observed among castes in *R. speratus*. Moreover, it is interesting to note that the wing-specific selector gene *vestigial* was significantly highly expressed in wing-less soldier heads (Supplementary Fig. 32). We further need to examine gene expression profiles and gene function during development of each caste on the wing primordia and wing buds to discuss the developmental mechanism of termite wing polyphenism. Identification of the wing developmental toolkit genes in this study facilitate those study.

**Immunity**

The group living of social insects enables task partitioning among individuals, i.e., division of labor, which is thought to realize higher productivity per individual than the solitary living does. On the other hand, group living is at high risk for infectious diseases because of high density of individuals and strong and frequent interactions among individuals. Moreover, many termite species including *R. speratus*, live in pathogenic microbe-rich environments, such as damp woods and soils[140]. Thus, pathogenic infections could have a large impact on fitness of termite colonies. Since, generally, immune system of individuals is the major defense mechanism against pathogens, and thus it is likely that termite immune systems have adaptively evolved. Identification of genes involved in the immune system gives us an opportunity to study how termites respond to pathogens.

We searched the *R. speratus* genome for the immune-related genes listed in ImmunoDB, which were classified into 27 categories based on gene function and pathway[141]. First, we downloaded protein sequences of the immune-related genes of *D. melanogaster*, *Anopheles gambiae*, and *Aedes aegypti* from ImmunoDB. Then, using those protein sequences as queries, BLASTP searches were carried out against the gene model RspeOGS1.0 to identify homologs of those genes in *R. speratus*. PfamScan was also performed for the categories with a specific protein domain.

We identified 251 immune-related genes from the *R. speratus* genome (Supplementary Table 25). Almost all genes involved in IMD/JNK, Toll and JAK/STAT signaling pathways were found in *R. speratus* as in the damp wood termite *Z. nevadensis* [27]. In all of the categories except for lysozymes, the numbers of genes are not remarkably increased or decreased in *R. speratus* compared to other insect species (Supplementary Fig. 8).

Eight genes encoding antimicrobial peptides (AMPs) were identified in the *R. speratus* genome (two defensins, termicin, crustin, locusin, prolixicin, two thaumatins; Supplementary Table 25). Most of them showed caste-biased expressions (Supplementary Fig. 33). One (RS002487) of the two defensin genes exhibited soldier-biased expressions while the termicin, crustin and one of the thaumatin genes worker-biased. Moreover, as mentioned in the main text, many of the lysozyme genes also exhibited caste-biased expressions.

Our results of the differential expressions of AMPs and lysozymes suggest possibility of division of labor among castes on colony-level immunity. AMPs and lysozymes are “effectors” of an immune system, which directly interact with microbes. Those effectors might have different target microbes (gram-positive, gram-negative bacteria, fungi, etc.). In addition, expression profiles of the effector genes were rather different among castes. Therefore, the differences of expression profiles may result in difference on immune potential against different microbes among castes. Division of labor on colony-level immunity among castes were also suggested in *R. speratus*[142]. Further studies are required to understand the adaptive evolution in termites, responding to pathogens.

**Insecticide target and detoxification genes**

*R. speratus* is a devastating pest species that can cause serious damages to wooden constructions and huge economic losses[143]. To prevent the damage by pest termites, use of Insecticides is efficient. To date, various kinds of insecticides were developed and are actually used for the termite control. However, development of new insecticides is still important to further reduce negative influence on environment and organisms other than termites[144]. The genome sequence information of pest species, especially sequences of insecticide target genes and detoxification-related genes, is useful to develop such insecticides and to study insecticide resistance mechanisms[145]. Here, we report some major insecticide target genes, namely, genes related to ion channels, chitin synthesis and muscle contraction. We also report the genes related to the detoxification of insecticides, that is, cytochrome P450 monooxygenases (CYP), glutathione S-transferases (GST), carboxylesterases (CCE). In insects, those three groups of enzymes play major roles in synthesis and degradation of endogenous substrates such as hormones and pheromones as well as in metabolization of exogenous substrates such as insecticides[146-148], and can be candidates of causal factor of insecticide resistance.

To identify the insecticide target genes, we performed BLASTP or TBLASTN searches against the gene models or the genome assembly of *R. speratus* using the amino acid sequences of those genes of *Drosophila melanogaster* as queries. Identification of the detoxification genes was carried out by profile hidden markov models (profile HMMs) search. Profile HMMs of CYP (PF00067), GST (PF00043 and PF02798), CCE (PF00135) that were retrieved from the Pfam database (http://pfam.xfam.org) were searched against the gene models of *R. speratus* using HMMER v3.1b1[149]. The profile HMM searches were also carried out against the gene models of *Z. nevadensis*[27] and *M. natalensis*[49] and the amino acid sequences obtained from transcriptome sequencing data (assembled contig sequences using Trinity[150]) of *Periplaneta americana*[151] and *Cryptocercus punctulatus*[152] (DRA001254 and DRA004598, respectively). To perform a phylogenetic analysis on the amino acid sequences with the profile HMM hits, those sequences were aligned with MAFFT[71], and the best models of amino acid replacements in the alignments were determined using ProtTest v3.4[153]. Then, a maximum likelihood-based phylogenetic trees were generated based on the alignments with the best replacement models using RAxML[73].

We identified 64 insecticide target genes in total (Supplementary Table 26). No remarkable gene family expansion was found in those genes. We found 106 CYP genes, 20 GST genes and 43 CCE genes in *R. speratus* (Supplementary Table 27). Numbers of genes identified in termites were smaller than those in 2 cockroaches, *P. americana* and *C. punctulatus* (Supplementary Table 27, 28). In the phylogenetic trees of the CYP genes, we found a clade containing only *R. speratus* genes with a high bootstrap support (Supplementary Fig. 34). In GST and CCE genes, we did not find and species-specific gene duplications in *R. speratus* (Supplementary Fig. 35, 36).

The identification of those genes could contribute to the development of species-specific control methods in *R. speratus*. The duplicated CYP genes could be new targets for the control methods. Examining functions of those genes are required for further development of the control methods. In the insecticide target genes and the GST and CCE genes, although no species-specific genes were found, there might be species-specific mutations resulting in changes in gene functions. Those mutations could be new targets of species-specific control methods. In this study, a large number of genes that could be targets of new control methods for *R. speratus* were identified, enabling new researches on termite controls.

**MicroRNAs (miRNAs)**

To evaluate the role of miRNAs for caste polyphenism, we have performed small RNA sequencing in workers and soldiers of *R. speratus*[154]. We identified eight miRNAs, which were differentially expressed in soldiers and workers.

### Supplementary Methodology

#### Insects

All mature colonies of *Reticulitermes speratus* used for genome, RNA, and Bisulfite sequencing (BS-seq), were collected in Furudo, Toyama Prefecture, Japan (colony #1-8) [Supplementary Table 1]. Pieces of logs were brought back to the laboratory and kept in plastic cases in constant darkness. For the extraction of genomic DNA, we used female secondary reproductives (nymphoids) in colony #1 (total of 2 individuals), collected in November 2013. For RNA sequencing (RNA-seq), workers and soldiers were sampled from colonies #2, #3, and #4 collected in September 2014. Primary reproductives (queen and king) were sampled from incipient colonies newly founded by alates (winged adults) that emerged from colonies #5, #6, and #7 collected in April–May 2014. For BS-seq, workers and soldiers were sampled from colony #8 collected in October 2014, and primary reproductives were sampled from incipient colonies founded by alates that emerged from colonies #5 and #7. For *in situ* hybridization, each caste (alates, female neotenics, workers, and soldiers) was sampled from mature colonies collected in Himi, Toyama Prefecture, Japan (colonies #9 and #10) in April–May 2019. Queens were sampled from incipient colonies newly founded by alates that emerged from colonies #9 and #10. Sexes of individuals were identified by means of the morphological characteristics of the 7th and 8th abdominal sternites (workers and soldiers)[155,156] or abdominal tergites (reproductives)[157].

#### Genome sequencing and assembly

We used female secondary reproductives (nymphoids I and II) (see above for details; colony #1 in Supplementary Table 1) for genome sequencing. We excluded the gut and ovaries of nymphoids to avoid contamination by DNAs from the king or other microorganisms. Remaining body parts were frozen in liquid nitrogen and stored at -80°C until DNA extraction. Genomic DNA was isolated from each individual using a QIAGEN Genomic-tip 20/G (Qiagen, Venlo, Netherlands). We used 5 microsatellite loci (Rf6-1, Rf21-1, Rf24-2, Rs02, and Rs03) to confirm whether they were homozygous at these loci and shared the same genotype. Primer sequences for the amplification of Rf and Rs loci are described in Vargo and Hayashi et al., respectively [158,159]. The quantity and quality of extracted DNA were analyzed using a NanoVue spectrophotometer (Cytiva, Marlborough, MA) and Qubit 2.0 fluorometer (Thermo Fisher Scientific, Waltham, MA). The integrity of genomic DNA was analyzed using pulsed-field electrophoresis in a 0.75% agarose gel (80 V, 16 hours).

Genomic DNA (derived from nymphoid I) purified as described above was fragmented with a Covaris S2 sonicator (Covaris, Woburn, MA), size-selected with BluePippin (Sage Science, Beverly, MA), and then used to create two pair-end libraries using a TruSeq DNA Sample Preparation Kit (Illumina, San Diego, CA) with insert sizes of ~250 and ~800 bp [Supplementary Table 3]. The enrichment PCR was done using six cycles. These libraries were sequenced using an Illumina HiSeq 1500 with a 2 × 151 bp paired-end sequencing protocol in Rapid mode at the NIBB Functional Genomics Facility (Okazaki, Japan). Four Mate-pair libraries with peaks at ~3 kb, ~5 kb, ~8 kb and ~10 kb, respectively, were created from the genomic DNA (derived from nymphoid II with the same genotype as described above) using a Nextera Mate Pair Sample Preparation Kit (Illumina) [Supplementary Table 3], and sequenced on a HiSeq system using a 2 × 151 bp paired-end sequencing protocol at the National Institute of Genetics (Mishima, Japan). Reads of the pair-end and mate-pair libraries were assembled using ALLPATHS-LG (build# 47878)[160], with default parameters. BUSCO v4.0.6[161] was used in quantitative measuring for the assessment of genome assembly, using insecta_odb10 as the lineage input. A genome browser was built using JBrowse[162] and is available at http://www.termites.nibb.info.

#### Gene prediction

A protein-coding gene reference set was generated using EvidenceModeler (EVM)[163] with two main sources of evidence, aligned *R. speratus* transcripts and aligned homologous proteins of other insects, and a set of *ab initio* gene predictions. RNA-seq reads were assembled *de novo* using Trinity[164], and ORFs were predicted using TransDecoder[165]. We used CD-HIT-EST[166] to reduce the redundancy of the predicted ORFs. The ORF sequences were mapped to the genome using Exonerate[167] in est2genome mode for splice-aware alignment. We processed homology evidence at the protein level using the reference proteomes of *Drosophila melanogaster* (FlyBase; Attrill et al. 2016), *Tribolium castaneum* (accession no. AAJJ00000000), *Apis mellifera* (accession no. AADG05000000), *Acyrthosiphon pisum* (accession no. ABLF01000000), *Daphnia pulex* (accession no. ACJG00000000), *Pediculus humanus* (accession no. AAZO00000000), and *Zootermopsis nevadensis* (accession no. AUST00000000). We also included Blattodea protein sequences predicted from *de novo* assembly of RNA-seq reads of *Periplaneta americana*[168] and *Nasutitermes takasagoensis*[169]. These reference proteins were split-mapped to the *R. speratus* genome in two steps: first with BLASTX to find approximate loci, and then with Exonerate in protein2genome mode to obtain more refined alignments. For an *ab initio* gene prediction, Augustus[170] was trained against a set of preliminary gene models of *R. speratus* (earlier version of EVM set) and then was used to predict the gene models in the *R. speratus* genome. These gene models derived from multiple evidence were merged using the EVM program to obtain the reference annotation for the genome, which yielded 15584 predicted genes. Lastly, genes of interest were manually inspected and corrected. In particular, tandemly duplicated genes discussed in the main text such as GGPPS and lysozyme genes were liable to be incorrect gene models with erroneous exon–exon connections between paralogous genes in the tandemly repeated cluster. In total, 74 gene models were manually updated. The final gene set composed of 15591 genes was designated as 'Rspe OGS1.0" [Supplementary Data 2 (DOI: 10.6084/m9.figshare.14267381)].

**Functional annotation of gene models**

Functional annotation of Rspe OGS1.0 was carried out by homology searches and motif searches. We scanned protein sequences of the Emsembl release-33 of *D. pulex*, *P. humanus*, *A. mellifera*, *D. melanogaster*, and *T. castaneum*, and the gene models of *Z. nevadensis* (Znev OGS v2.229), and *Macrotermes natalensis* (Mnat OGS3). We also scanned the protein sequences of Rspe OGS1.0 using InterProScan v5.17-56.0[171], to annotate domains and motifs of predicted *R. speratus* coding genes. The Kyoto Encyclopedia of Genes and Genomes (KEGG) annotation for Rspe OGS1.0 was performed on the web server (https://www.kegg.jp/blastkoala/) with the BlastKOALA algorithm[172,173]. Gene Ontology terms were assigned to Rspe OGS1.0 gene models by analyzing the results of the BLASTP searches against the NCBI nr database (version on March 3rd, 2016) and InterProScan searches with the Blast2GO pipeline (B2G4Pipe version 2.5.0)[174].

The quality of the Rspe OGS1.0 gene set was evaluated by assessing two types of evidence, homology evidence and expression evidence. Among 15591 genes, 12996 (83.3%) showed any hits in the NCBI nr database, 10440 (70.0%) included known protein motifs defined in the Pfam database, and 14302 (91.7%) showed evidence of expression with a threshold of RPKM = 1.0 in any sample of caste-specific RNA-seq data. In sum, 15577 (99.9%) had any evidence for the presence of homologs and/or expression.

**Orthology inference and gene duplication analysis**

Orthology determination among three termites: Orthologous genes among the proteomes of three termite species, namely, *R. speratus*, *Z. nevadensis*, and *M. natalensis* (gene models Rspe OGS1.0, Znev OGS v2.229, and Mnat OGS3, respectively), were determined by pairwise comparisons with InParanoid v4.1 followed by three-species comparison with MultiParanoid[175,176]. Note that the *M. natalensis* gene set, Mnat OGS3, was built in this study using a similar pipeline as used for *R. speratus* gene prediction. The BUSCO analysis indicated that Mnat OGS3 recovered 95.0% of insecta benchmarking universal single-copy orthologs (BUSCOs) showing significant improvement from original gene models, Mnat_gene_v1.2[49], which captured 83.1% of insecta BUSCOs.

Ortholog analysis with arthropod proteomes: Orthology relationships of *R. speratus* genes (OGS1.0) with other arthropod genes were analyzed by referring to the OrthoDB gene orthology database. We downloaded the arthropod ortholog table and all protein sequences provided by the OrthoDB ver.8 database (87 arthropod species)[177]. We grouped *R. speratus* genes with the OrthoDB ortholog group using a two-step clustering procedure implemented in custom Ruby scripts. For each *R. speratus* protein, BLASTP was used to find similar proteins among the arthropod proteins, and the ortholog group of the top hit was provisionally assigned to the query *R. speratus* gene. Then, the ortholog grouping was evaluated by comparing the similarity level (BLAST bit score was used as a proxy) among members within the focal ortholog group. We keep the grouping if the BLAST bit score between the query *R. speratus* gene and top arthropod gene was higher than the minimal score within the original cluster members. Among 15591 *R. speratus* OGS1.0 genes, 12434 genes were clustered into 9033 OrthoDB Arthropod ortholog groups. Gene duplication was assessed based on this clustering. If two or more members of one species were found in a single ortholog group, they were regarded as a multigene family.

**Repeat sequence annotation**

To annotate repeat sequences, first, we generated repeat sequence models using RepeatModeler (http://www.repeatmasker.org) from the genome assemblies of *R. speratus* (this work), *Z. nevadensis*[27], and *M. natalensis*[49]. Then, we pooled the repeat models of the three species. Using CD-HIT-EST[166], we clustered those sequences with >90% of sequence similarity and used only the longest sequences in the clusters as repeat models. We identified repeat sequences using RepeatMasker with the retained repeat models in each of the three species genome assemblies.

**RNA-seq**

W4–5 workers (old workers) and soldiers were collected from each colony according to the body size and antennal segments[178]. To collect primary reproductives, dealated adults, were chosen randomly from each colony in accordance with the method of the previous literature[108], and female–male pairs were mated (colonies #5 and #6: 10 pairs, #5 and #7: 10 pairs, #6 and #7: 10 pairs; Supplementary Table 1). Each pair was placed in a 20-mL glass vial with c. 8 g of mixed sawdust food (Mitani, Ibaraki, Japan) and kept at 25°C in constant darkness. Colonies were then sampled after 4 months. We observed plural larvae and several workers in each colony, and kings and queens were sampled. Each individual was divided into head and body parts (including thorax and abdomen with the guts) on ice, immediately frozen in liquid nitrogen and stored at -80°C until use.

We prepared RNA-seq libraries for 12 categories based on castes (reproductives, workers and soldiers), sexes (males and females) and body parts (head, and thorax and abdomen). Ten individuals were combined for each head sample of each caste and each sex, and five individuals for the thorax and abdomen sample. Three biological replications of the 12 categories were made with three different field colonies totaling 36 RNA-seq libraries [Supplementary Table 2]. Total RNA was isolated from each category using an SV total RNA isolation system (Promega, Madison, WI, USA). DNA was digested with RNase-free DNase I for 20 min at 37°C. The quantity and quality of extracted RNA were checked using a NanoVue spectrophotometer (Cytiva), Qubit 2.0 fluorometer (Thermo Fisher Scientific), and Agilent 2100 bioanalyzer (Agilent Technologies, Palo Alto, CA). Illumina libraries for RNA-seq were prepared using a TruSeq Stranded mRNA Library Prep kit (Illumina) in accordance with the manufacturer's instructions. First- and second-strand cDNA synthesis, adaptor ligation, and amplification were performed. The generated libraries were evaluated using RT-qPCR with a KAPA qPCR SYBR green PCR kit (Geneworks, Thebarton, Australia) and electrophoresis in an Agilent 2100 bioanalyzer (Agilent Technologies). All libraries were subjected to a single-end sequencing of 101 bp fragments on HiSeq 2500 (Illumina).

The raw sequencing reads were filtered to remove adapter sequences and low-quality bases using Trimmomatic v0.32[179] with the following thresholds: leading and trailing bases with a Phred quality score (Q) lower than 20, and other sequences lower than Q20 for the average quality in a 4-bp sliding window, but with a minimum length of the sequence read of 50 bp. Subsequently, the filtered reads were mapped onto the genome assembly with TopHat v2.1.0[180] guided by the gene models. Transcript abundances were then estimated using the featureCounts program of the Subread package[181]. To compare gene expression levels among castes and between sexes, first, counts per million (CPM) were calculated from the estimated transcript abundances. We kept genes with at least CPM of 1 in at least three samples for subsequent analyses. CPM values were then normalized with the trimmed mean of M-values (TMM) algorism in edgeR[182]. Differentially expressed genes among castes and between sexes were detected in each body part (head / thorax and abdomen) using a generalized linear model with two factors, namely, caste and sex using edgeR with the conditions set as false discovery rate (FDR) < 0.01 and the log2 fold change of the expression level > 1. MDS plot was made using the plotMDS function implemented in edgeR. RPKM (Reads Per Kilobase Million) values were calculated by dividing the CPM values by the length of the genes in kilobases.

**Methylome analysis**

W4–5 workers (old workers) and soldiers were collected from the colony as described in the previous section. To collect primary reproductives, female–male pairs and incipient colonies were prepared as shown in the previous section. Colonies were then sampled after 6 months, and primary reproductives were sampled. The head of each individual was frozen in liquid nitrogen and stored at -80°C until use. For DNA extraction, we used 10 heads per category. We prepared 6 categories based on castes (reproductives, workers, soldiers) and sexes (males, females) [Supplementary Table 3]. Total DNA was isolated from each category using a QIAGEN Genomic-tip 20/G (Qiagen). The quantity and quality of extracted DNA were checked using a NanoVue spectrophotometer (Cytiva) and Qubit 2.0 fluorometer (Thermo Fisher Scientific).
Samples containing 200 ng of genomic DNA each for the 6 categories were used to construct Methylome libraries using a post-bisulfite adaptor tagging (PBAT) technique to perform whole-genome bisulfite sequencing (WGBS)[183]. The isolated *R. speratus* genomic DNA, together with 1% unmethylated lambda DNA as a control, were subjected to bisulfite conversion using an EZ DNA Methylation-Gold Kit (ZYMO Research, Irvine, CA) in accordance with manufacturer's instructions. The bisulfite-converted templates were then subjected to adaptor tagging as described previously[183]. The generated libraries were assessed on the Agilent Bioanalyzer 2100 platform and quantified with a standard curve-based qPCR assay (KAPA Biosystems, Wilmington, MA). The final quality-ensured libraries were pooled and sequenced using an Illumina HiSeq 2500 sequencer for 101 bp single-end sequencing.

De-muliplexed raw reads were trimmed of sequencing adapters and low-quality ends (<Q30) using Cutadapt[184]. The 909,207,083 processed reads (91.8 Gb) were mapped to the *R. speratus* genome using the Bismark program (v0.16.1)[185] with the options "--bowtie2 --pbat --unmapped". The mapping rate ranged from 52% to 56%. The mapping results in BAM format were subjected to methylation calling using the bismark_methylation_extractor command included in the Bismark package. Significant methylation sites were identified based on a binomial model using the binom.test function in R (R Core Team 2015), where we excluded sites with too high (>100) or too low (<10) coverage. Differentially methylated regions (DMRs) among castes were investigated using two pipelines. First, we used ANOVA of the ratio of methylation sites by exons. After multiple comparison correction (FDR) and filtering by difference (>30.0%), no DMRs were detected. Next, we used BSmooth software[186] to find DMRs. Bismark CpG report files were loaded into BSmooth, and CpG sites with enough read coverage (more than 2 in all samples) were used for smoothing and DMR analysis.

**Manual annotations and analyses of specific categories**

**Lipocalins**. Sequence alignments of lipocalin-related Pfam domains (PF00061, PF08212, PF02087, PF07137, and PF02098) of arthropods were generated using the Pfam website (https://pfam.xfam.org)[187] and downloaded. Profile hidden Markov models (HMMs) of the alignments were then obtained with hmmbuild of the HMMER package version 3.1 beta2 (http://hmmer.org/). We searched gene models of each of the arthropod species (*D. pulex*, *Z. nevadensis*, *R. speratus*, *M. natalensis*, *P. humanus*, *A. mellifera*, *D. melanogaster*, and *T. castaneum*) with the HMMs using hmmsearch in the HMMER package[188]. We annotated significantly similar sequence matches from the hmmsearch as lipocalins. In addition, we also searched the InterProScan results of the arthropod species mentioned above for proteins with the lipocalin-related PROSITE (PS00213) and Pfam domains (IDs mentioned above) signatures, and annotated them as lipocalins. For phylogenetic tree construction, protein sequences of the lipocalins annotated above, SOL1 of *Hodotermopsis sjostedti* (NCBI accession no. BAA87882) and its homologous sequences of *Coptotermes formosanus* (AGM32427) were aligned using the E-INS-i strategy in MAFFT v7.3.1[189]. The best-fit model of amino acid replacement for the alignment was determined with ProtTest v3.4[190] based on the Bayesian information criterion[191]. A maximum likelihood phylogenetic tree was generated from the alignment using RAxML[75] with 100 bootstrap replicates.

**Cellulases.** Amino acid sequences deduced from the *R. speratus*, *Z. nevadensis*, *M. natalensis*, *A. mellifera*, and *D. melanogaster* genomes were annotated for CAZy (Carbohydrate Active Enzymes) families based on the dbCAN database v3[192]. Hmmscan on dbCAN was performed with default settings; E-value < 1e–3 and < 1e–5 as the cutoff values for alignments shorter and longer than 80 amino acids, respectively. Because results with E-value < 1e–3 frequently contained false positives based on BLASTP, these were manually removed. For phylogenetic tree construction, 38 amino acid sequences of GH1 (β-glucosidase) genes were aligned using the Clustal method with MEGA v6.06[193], and gaps were excluded manually. The best-fit model of amino acid sequence evolution was determined using the model selection option implemented in MEGA. A maximum likelihood phylogenetic tree was constructed using MEGA with 1000 bootstrap replicates.

**Lysozymes.** To identify lysozyme genes in *R. speratus*, BLASTP searches were carried out against the *R. speratus* gene model Rspe OGS1.0 using lysozyme protein sequences of *D. melanogaster*, *Anopheles gambiae*, and *Aedes aegypti* retrieved from ImmunoDB[194] as the query. Moreover, BLASTP searches for i-type lysozyme protein sequences of various insects[195] were also performed against the *R. speratus* gene model. E-value cutoff values of these BLASTP searches were set at 1e–20. In addition, we performed PfamScan[196] for the protein domain PF00062 (c-type lysozyme/alpha-lactalbumin family).

We further searched proteome datasets from various arthropod species for lysozyme genes to perform a phylogenetic analysis of lysozyme genes. Proteome sequence data of the arthropods were downloaded from the following web databases: Ensembl[197] for *Bombyx mori*, *Nasonia vitripennis*, and *T. castaneum*, VectorBase[198] for *A. gambiae*, *Ixodes scapularis*, *P. humanus*, and *Rhodnius prolixus*, Hymenoptera Genome Database[199] for *Acromyrmex echinatior*, *A. mellifera*, and *Camponotus floridanus*, AphidBase[200] for *A. pisum*, wFleaBase[201] for *D. pulex*, FlyBase[202] for *D. melanogaster*, Znev OGS v2.229 and Mnat OGS3. When the proteome data sets included isoforms derived from the same genes, we retained only the longest one for the further analyses. BLASTP searches and PfamScan were performed in proteome data sets of those species using the same method as for *R. speratus*. The protein sequences of the lysozymes identified in the arthropods were aligned with the E-INS-i strategy of the MAFFT program[189]. Then, a codon-based alignment for the coding sequences of the lysozymes was generated using the PAL2NAL program[203] in accordance with the protein alignment. A phylogenetic tree was reconstructed from the codon-based alignment with a maximum likelihood approach based on GTR + gamma using RAxML[204]. The codon-based alignment was partitioned into three codon positions and a particular parameter set was estimated for each position. One thousand bootstrap replicates were made to assess the branch support.

**GGPP synthases.** We performed TBLASTX with the nucleotide sequences of *A. pisum* GGPP synthase (accession no. XP_008184262.1) as a query, using the *R. speratus* gene model (older version of OGS1.0) and the genome final assembly. We found that GGPP synthase homologs were tandemly duplicated on scaffold 31, and the identified gene model was manually curated according to the homologous DNA and deduced amino acid sequences. We identified the homologous synteny blocks in the *Z. nevadensis* and *M. natalensis* genomes (scafolds 797 and 103, respectively), and conservation of synteny between termite genomes was revealed using dot plots. For phylogenetic tree construction, 26 amino acid sequences of GGPP synthase genes, including 13 homologs of *R. speratus*, were aligned using the Muscle method with MEGA v7[205]. The best-fit model of amino acid sequence evolution was determined using the model selection option implemented in MEGA. A maximum likelihood phylogenetic tree was constructed using MEGA with 100 bootstrap replicates.

For molecular evolutionary analysis, PRANK v170427[206] was used to align amino acid sequences of 13 homologs of *R. speratus* and those of six other insects, *A. pisum, A. mellifera, T. castaneum, D. melanogaster, Z. nevadensis*, and *M. natalensis*, and then back-translate to coding sequences. Gblocks v0.91b[207] was used to eliminate poorly aligned positions. The aligned 289 aa (867 bp) sites were subjected to downstream analyses. A gene tree was constructed by using RAxML ver. 8.2.12.[204] with the following parameters: -f a -# 100 -m PROTGAMMAAUTO. To test for signatures of positive selection acting on the lineages leading to *R. speratus* GGPP synthase paralogs, we compared the likelihood scores of selection models implemented in CODEML in the PAML package v4.9[208], using likelihood ratio tests. We used the branch-site test of positive selection (branch-site model A), where branches of interests are treated as foreground allowing three classes of sites (0<w<1, w=1, w=>1), and others, as background with two classes of sites (w=0, w=1), by specifying the following parameters: fix_omega = 0, omega = 1 and NSsites = 2. The likelihood ratio test statistics were compared against the $\lambda^2_{df} = 1$ distribution to calculate *P*-values followed by multiple testing correction using the method of Hommel implemented in R.

**TY family.** Secretion signal sequences in TY family proteins of *R. speratus* were predicted using the SignalP v4.1 program[209]. The TY family homologs in *Z. nevadensi*s and *M. natalensis* were identified based on the ortholog analysis described above and synteny information. Multiple alignments of TY family homologs from termites were constructed using MUSCLE software[210]. Non-synonymous (*Ka*) and synonymous (*Ks*) substitution rates of paired-wise paralogues were calculated with KaKs_Calculator v2.0[211] for codon-aligned sequences generated by the tranalign program included in the EMBOSS suite[212].

**RNA *in situ* hybridization**

All castes examined except for female primary reproductives (queens) were collected from mature colonies (#9 and #10). Queens were sampled 4 months after incipient colony foundation, as described in the previous section. To prepare RNA probes, specific primers for three *lipocalins* (*RS008881*, *RS008882*, and *RS008823*), two *GH1s* (*β-glucosidases*; *RS004136* and *RS004624*) and one *GGPP synthase* (*RS100016*) were designed using Primer3Plus[213] (Supplementary Table 6). Total RNA for probe synthesis was extracted from the whole bodies of female neotenics (*RS008881*), queens (*RS004624*), workers (*RS008882* and *RS004136*), and soldiers (*RS008823* and *RS100016*) using Isogene II (Nippon Gene, Tokyo, Japan). After treatment with DNase I (Takara Bio, Shiga, Japan), the quality and quantity of total RNA were measured using a NanoVue spectrophotometer (Cytiva). cDNA was synthesized using a High-Capacity cDNA Reverse Transcription Kit (Thermo Fisher Scientific). The PCR products from specific primers (Supplementary Table 6) were purified using a QIAquick Gel Extraction Kit (Qiagen) and subcloned into a pGEM easy T-vector (Promega, Madison, WI). The inserted DNA was amplified, and PCR products were sequenced using a BigDye Terminator v. 3.1 Cycle Sequencing Kit and an automatic DNA Sequencer 3130 Genetic Analyzer (Thermo Fisher Scientific). Plasmids with the targeted fragments were extracted using a GenElute Plasmid Miniprep Kit (SIGMA-Aldrich, St. Louis, MO). The digoxygenin (DIG)-labeled sense or antisense RNA probes were produced using a DIG RNA Labeling Kit (SP6/T7) (SIGMA-Aldrich), and purified with Ethachinmate (Nippon Gene).

Prior to cryosectioning, the abdomens of queens [*RS008881* (n = 4) and *RS004624* (n = 3)], heads and thoraxes of workers [*RS008882* (n = 3) and *RS004136* (n = 3)], and heads of soldiers [*RS008823* (n = 3) and *RS100016* (n = 2)] were dissected from the bodies and fixed with 4% paraformaldehyde in phosphate-buffered saline. Samples were embedded in TissueTek O.C.T. Compound (Sakura Finetek USA Inc., Torrance, CA). Cryosections (10 µm) were collected on CREST-coated glass slides (Matsunami, Osaka, Japan) using a CM1510S cryostat (Leica Biosystems, Nussloch, Germany). The sections were hybridized with DIG-labeled sense or antisense RNA probes using *In situ* hybridization reagents (Nippon Gene) in accordance with the instructions provided by the manufacturer. Immunocytochemical detection of DIG-labeled RNA was performed using a DIG Nucleic Acid Detection Kit (Roche, Grenzacherstrasse, Basel, Switzerland) in accordance with the manufacturer's instructions. The images were captured using a Biozero microscope (Keyence, Tokyo, Japan).

**Data availability**

Data from whole-genome sequencing, transcriptome sequencing, and methylome sequencing have been deposited in the DDBJ database under BioProject accessions PRJDB2984, PRJDB5589 and PRJDB11323, respectively. The analyzed data including genome assembly, gene prediction, annotation, and gene expression are available through FigShare (doi: m9.figshare.14267342, doi: 10.6084/m9.figshare.14267381, doi: 10.6084/m9.figshare.14267498). The *R. speratus* genome browser is available at http://www.termite.nibb.info/retsp/.

**Code availability**

All software used in this study for data analyses are open source. Custom R, Ruby and Shell scripts were deposited into GitHub (https://github.com/termiteg/retsp_genome_paper).

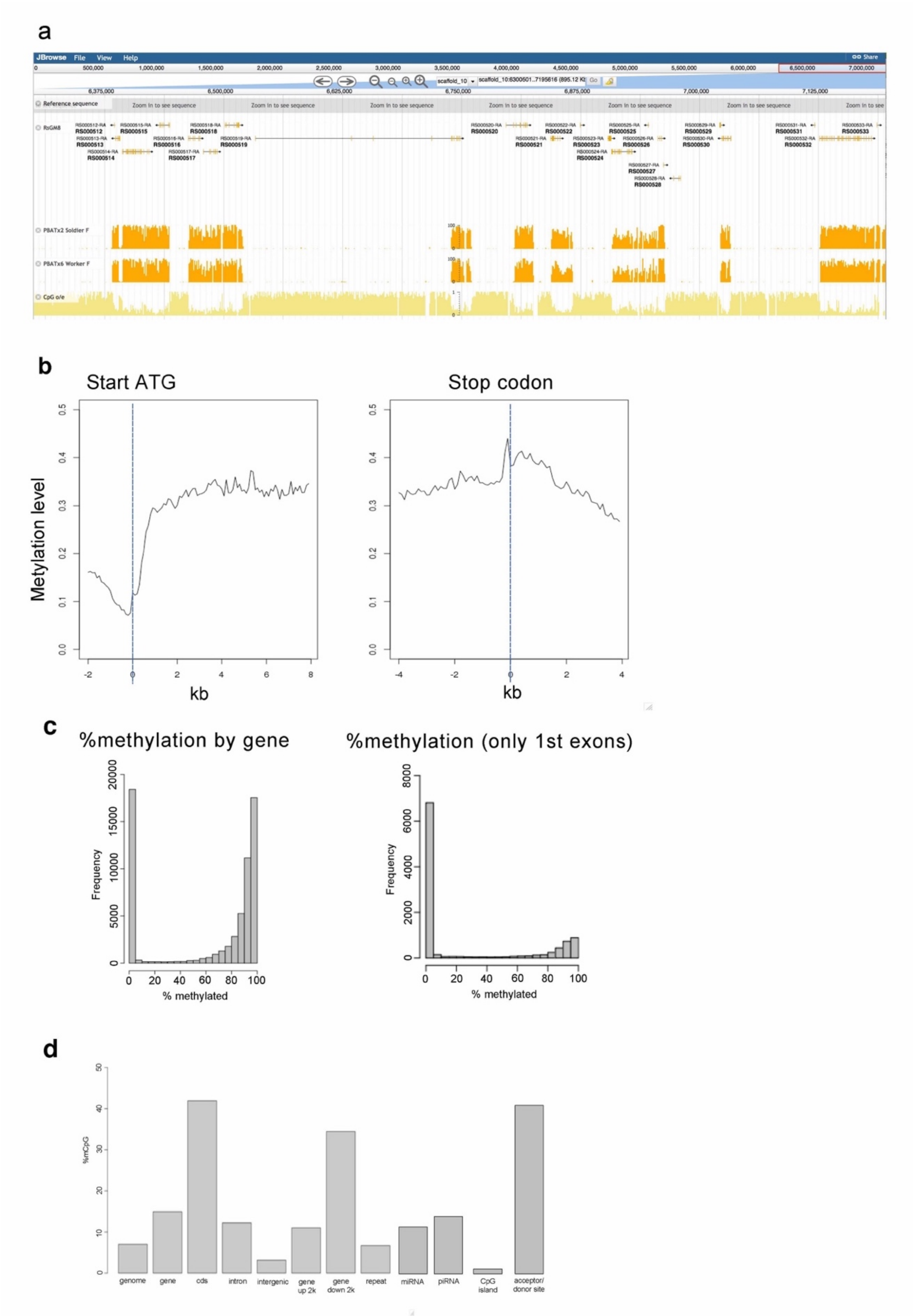
a
JBrowse File View Help
Share
Reference sequence
Zoom in to see sequence
RsGM8
PBATx2 Soldier F
PBATx6 Worker F
CpG o/e
b
Start ATG
Stop codon
Metylation level
kb
c
%methylation by gene
%methylation (only 1st exons)
Frequency
% methylated
d
%mCpG
genome
gene
cds
intron
intergenic
gene up 2k
gene down 2k
repeat
miRNA
piRNA
CpG island
acceptor/ donor site

**Fig. S1.** Global view of DNA methylation in the *Reticulitermes speratus* genome. (a) A snapshot of the genome browser showing the intensive gene body methylation. The top track indicates gene models with exon-intron structures. The orange barplots in the second (soldier female) and third (worker female) tracks indicate the methylation levels. The barplot in kahaki color in the bottom track shows the %CpG observed / expected value, which shows a notable negative correlation with the methylation levels. (b) Methylation levels around start codons (left) and stop codons (right) of all genes were averaged and plotted. (c) Methylation levels by gene. Methylation level was calculated by gene and summarized in the histogram showing a remarkable bimodal distribution (left). The same analysis conducted for only the first exons (right) indicated that the first exons are devoid of methylation in most genes. (d) Methylation level by the genomic context. CpG sites were partitioned by the genomic context, i.e., gene (exons and introns), protein-coding region (cds), intron, intergenic, 2k-upstream from start codon, 2k-downstream from stop codon, repeats, miRNA, piRNA, CpG island and acceptor-donor site), and calculated the %CpG for each category.

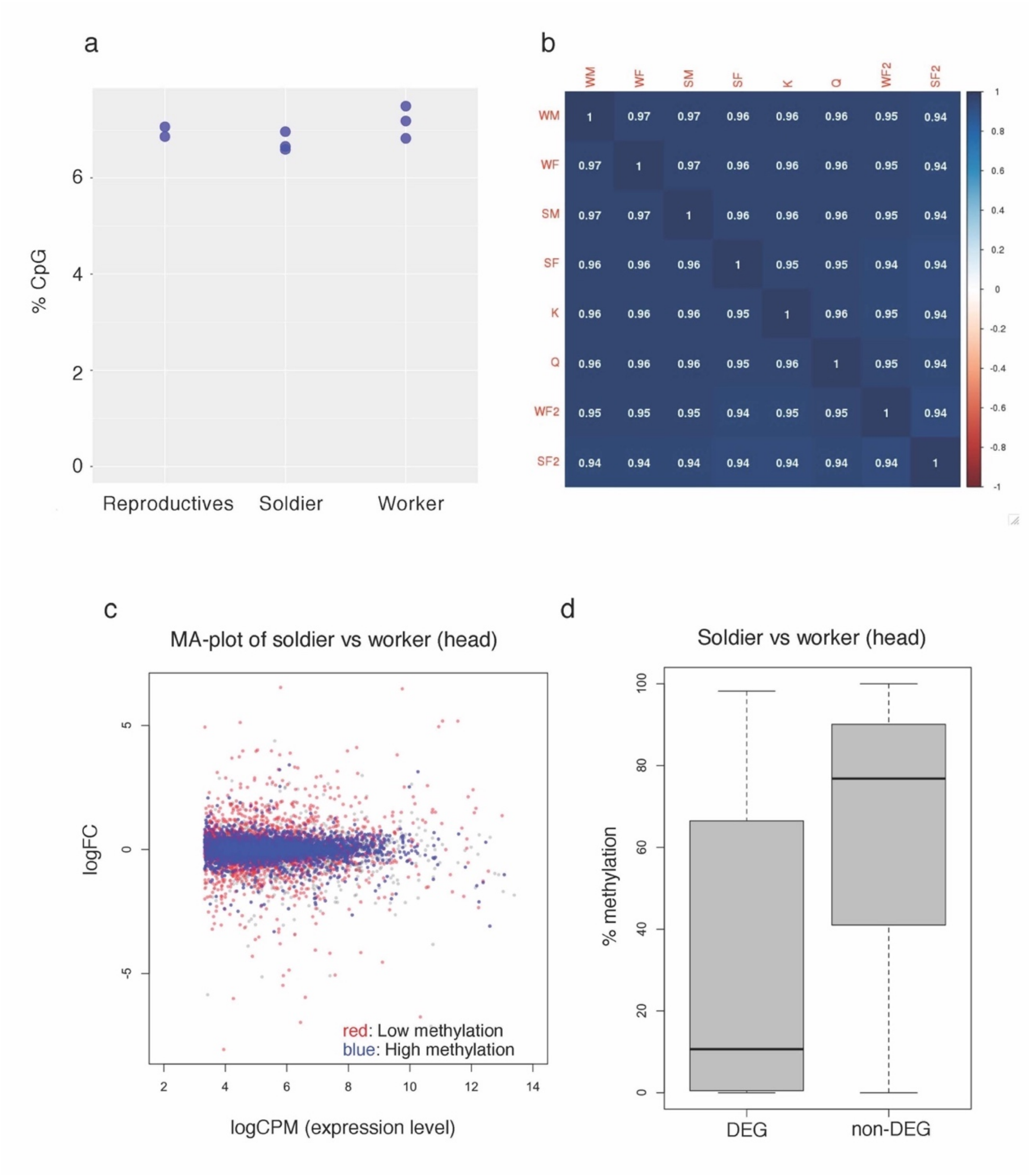


**Fig. S2.** Differential methylation pattern among castes. (a) Comparison of CpG levels among castes. No significant differences were detected. (b) Methylation pattern correlation among castes. Methylation levels were calculated by sliding the 200-bp window for each caste and the patterns were compared among castes. In any pair of comparisons, Pearson's correlation values were close to 1 suggesting the indistinguishable methylation pattern among castes. Male worker (WM), female worker (WF1-2), male soldier (SM), female soldier (SF1-2), king (K), queen (Q). (c, d) Caste-biased genes are unmethylated. Comparison of head transcriptomes between soldiers and workers is shown as a representative. The other comparisons of castes or body parts also showed similar patterns. In the MA plot comparing transcriptome of soldiers and workers (c), highly methylated genes (blue; >X%) showed a tendency to be plotted around logFC=0 (no changes between castes), while lowly methylated genes (red) are plotted away from the logFC=0 line. When genes are categorized into differentially expressed genes (DEG) and non-DE genes (non-DEG), the methylation levels are significantly different.

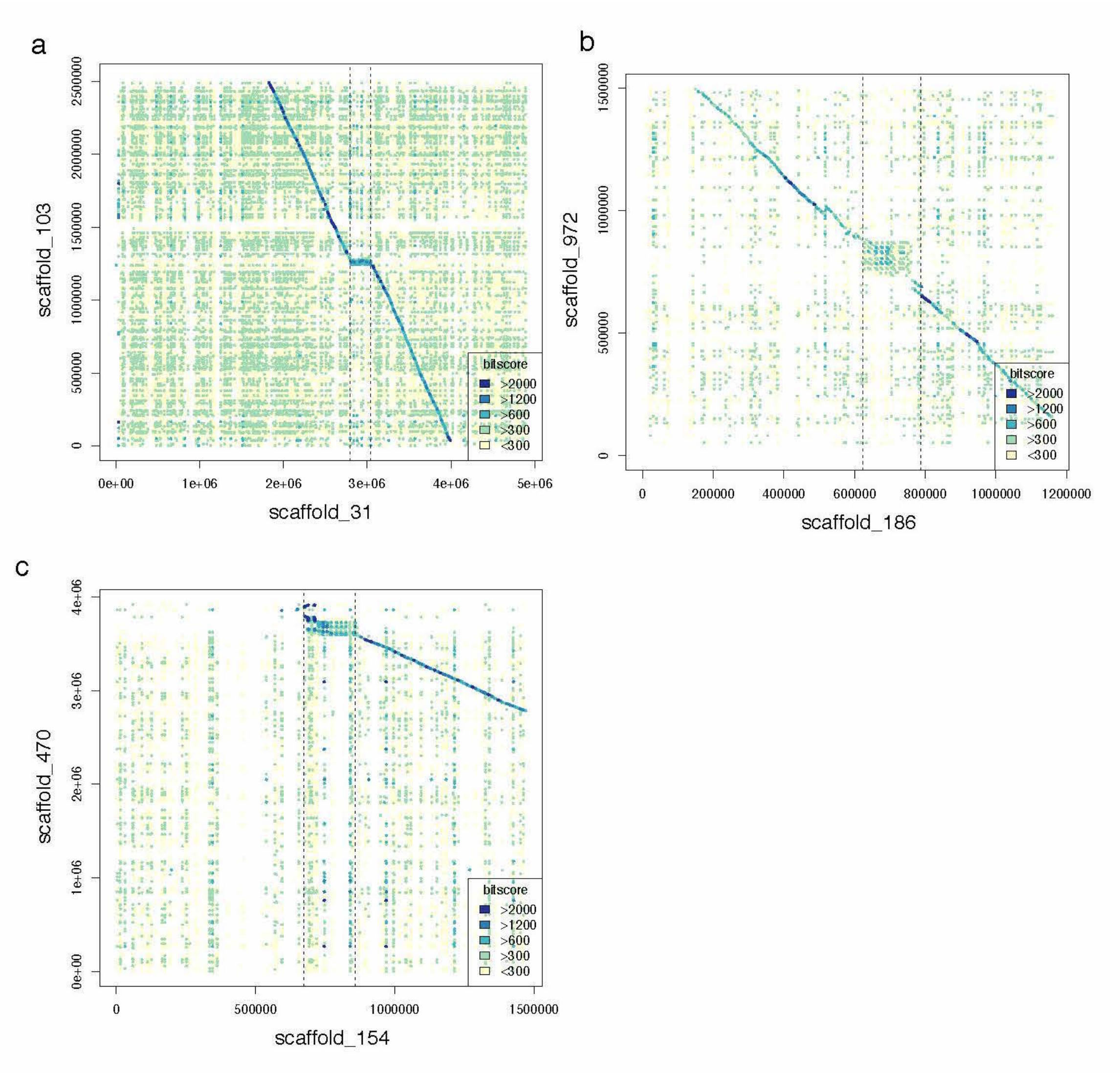


**Fig. S3.** Local syntenic break by tandem gene duplications. (a, b) Dot plots comparing the syntenic regions of the *Reticulitermes speratus* genome and the *Macrotermes natalensis* genome. The pair of syntenic sequences were compared with BLASTN and the aligned fragments were plotted with colors according to the bit scores. Scaffold_31 (a), scaffold_186 (b), and scaffold_154 (c) of the *R. speratus* assembly are shown.

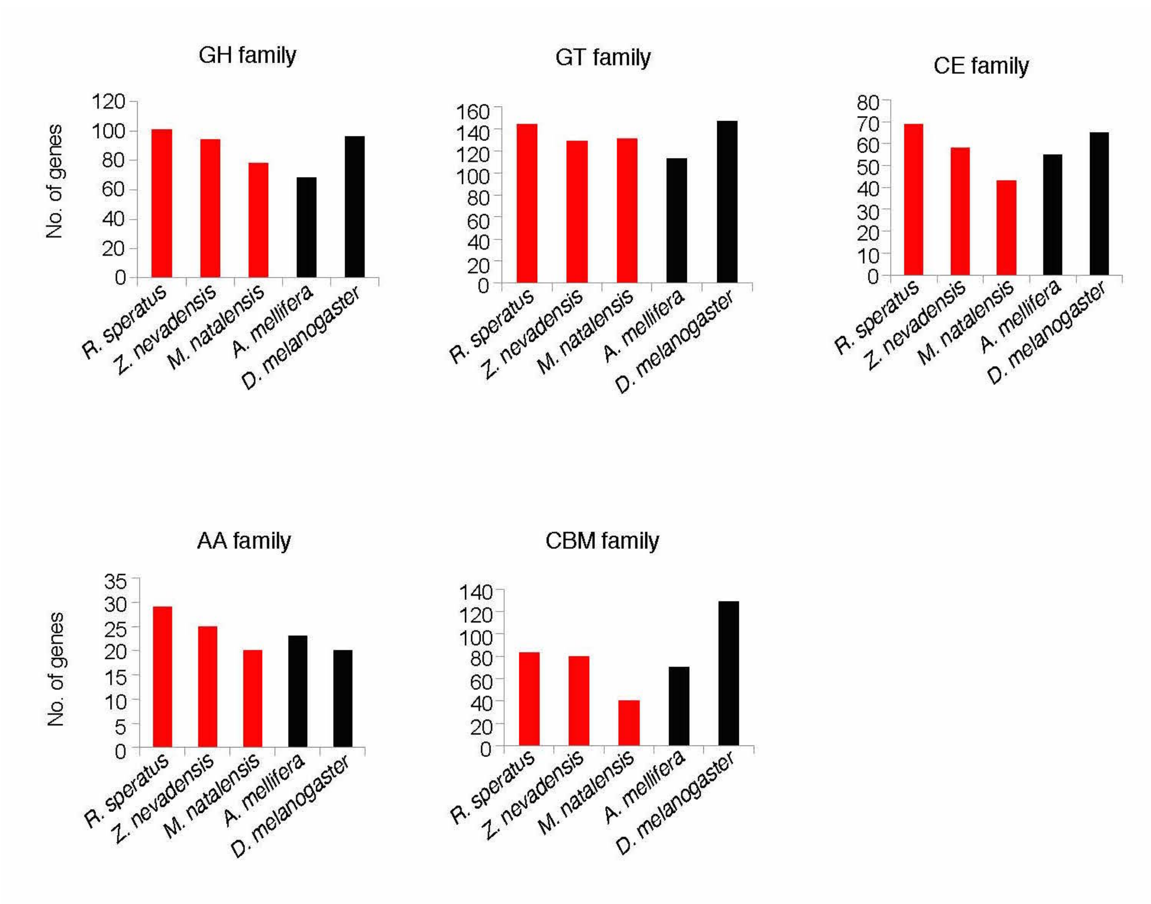


**Fig. S4.** Comparison of number of CAZyme-encoding genes among 5 insect species. The number of genes that belong to Glycoside Hydrolase (GH), Glycosyl Transferase (GT), Carbohydrate Esterase (CE), Auxiliary Activity (AA) and Carbohydrate-Binding Module (CBM) families are compared among *Reticulitermes speratus, Zootermopsis nevadensis*, *Macrotermes natalensis*, *Apis mellifera* and *Drosophila melanogaster*. Bar plots of termites are colored in red.

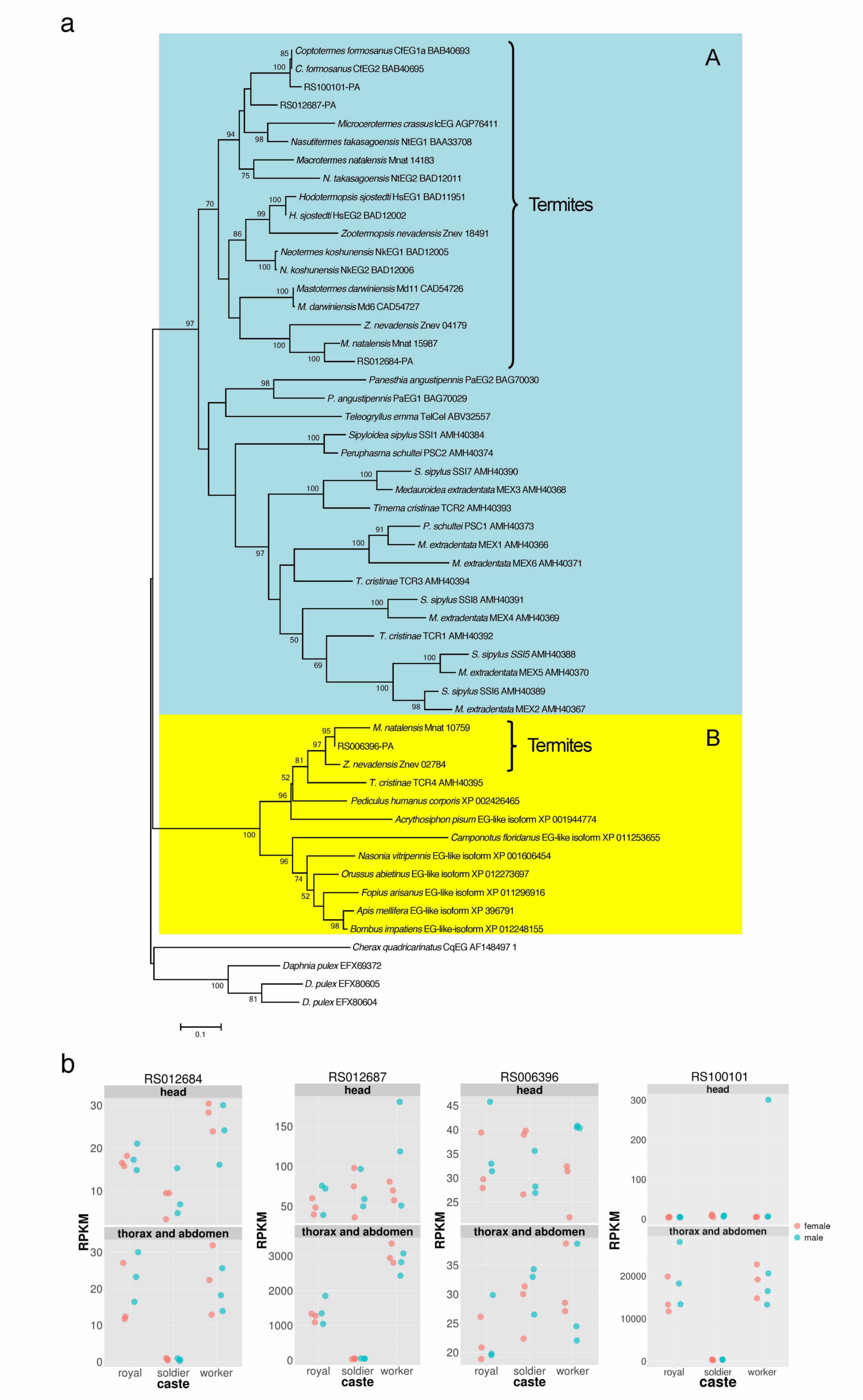

a
A
Coptotermes formosanus CfEG1a BAB40693
C. formosanus CfEG2 BAB40695
RS100101-PA
RS012687-PA
Microcerotermes crassus IcEG AGP76411
Nasutitermes takasagoensis NtEG1 BAA33708
Macrotermes natalensis Mnat 14183
N. takasagoensis NtEG2 BAD12011
Hodotermopsis sjostedti HsEG1 BAD11951
H. sjostedti HsEG2 BAD12002
Zootermopsis nevadensis Znev 18491
Neotermes koshunensis NkEG1 BAD12005
N. koshunensis NkEG2 BAD12006
Mastotermes darwiniensis Md11 CAD54726
M. darwiniensis Md6 CAD54727
Z. nevadensis Znev 04179
M. natalensis Mnat 15987
RS012684-PA
Termites
Panesthia angustipennis PaEG2 BAG70030
P. angustipennis PaEG1 BAG70029
Teleogryllus emma TelCel ABV32557
Sipyloidea sipylus SSI1 AMH40384
Peruphasma schultei PSC2 AMH40374
S. sipylus SSI7 AMH40390
Medauroidea extradentata MEX3 AMH40368
Timema cristinae TCR2 AMH40393
P. schultei PSC1 AMH40373
M. extradentata MEX1 AMH40366
M. extradentata MEX6 AMH40371
T. cristinae TCR3 AMH40394
S. sipylus SSI8 AMH40391
M. extradentata MEX4 AMH40369
T. cristinae TCR1 AMH40392
S. sipylus SSI5 AMH40388
M. extradentata MEX5 AMH40370
S. sipylus SSI6 AMH40389
M. extradentata MEX2 AMH40367
B
M. natalensis Mnat 10759
RS006396-PA
Z. nevadensis Znev 02784
Termites
T. cristinae TCR4 AMH40395
Pediculus humanus corporis XP 002426465
Acrythosiphon pisum EG-like isoform XP 001944774
Camponotus floridanus EG-like isoform XP 011253655
Nasonia vitripennis EG-like isoform XP 001606454
Orussus abietinus EG-like isoform XP 012273697
Fopius arisanus EG-like isoform XP 011296916
Apis mellifera EG-like isoform XP 396791
Bombus impatiens EG-like-isoform XP 012248155
Cherax quadricarinatus CqEG AF148497 1
Daphnia pulex EFX69372
D. pulex EFX80605
D. pulex EFX80604
0.1
b
RS012684
RS012687
RS006396
RS100101
head
thorax and abdomen
RPKM
royal
soldier
worker
caste
female
male

**Fig. S5.** Glycoside hydrolase family (GH) 9 in the *Reticulitermes speratus* genome. (a) Maximum likelihood (ML) tree of GH9 homologs based on the amino acid sequences obtained with a Le_Gascuel_2008 + Gamma model. The bootstrap percentages of 1000 ML trees in which the associated taxa clustered together are shown next to the nodes. The analysis involved 53 amino acid sequences. All positions containing gaps and missing data were eliminated. There were a total of 312 positions in the final dataset. Branches leading to clade A and clade B, which are 2 discrete GH9 groups of insects, are marked in blue and yellow, respectively. Two groups derived from termites are also marked. (b) Caste-biased expression patterns of GH9 family genes. Expression levels are indicated as RPKM calculated from RNA-sequencing analysis. Orange and blue points indicate females and males, respectively.

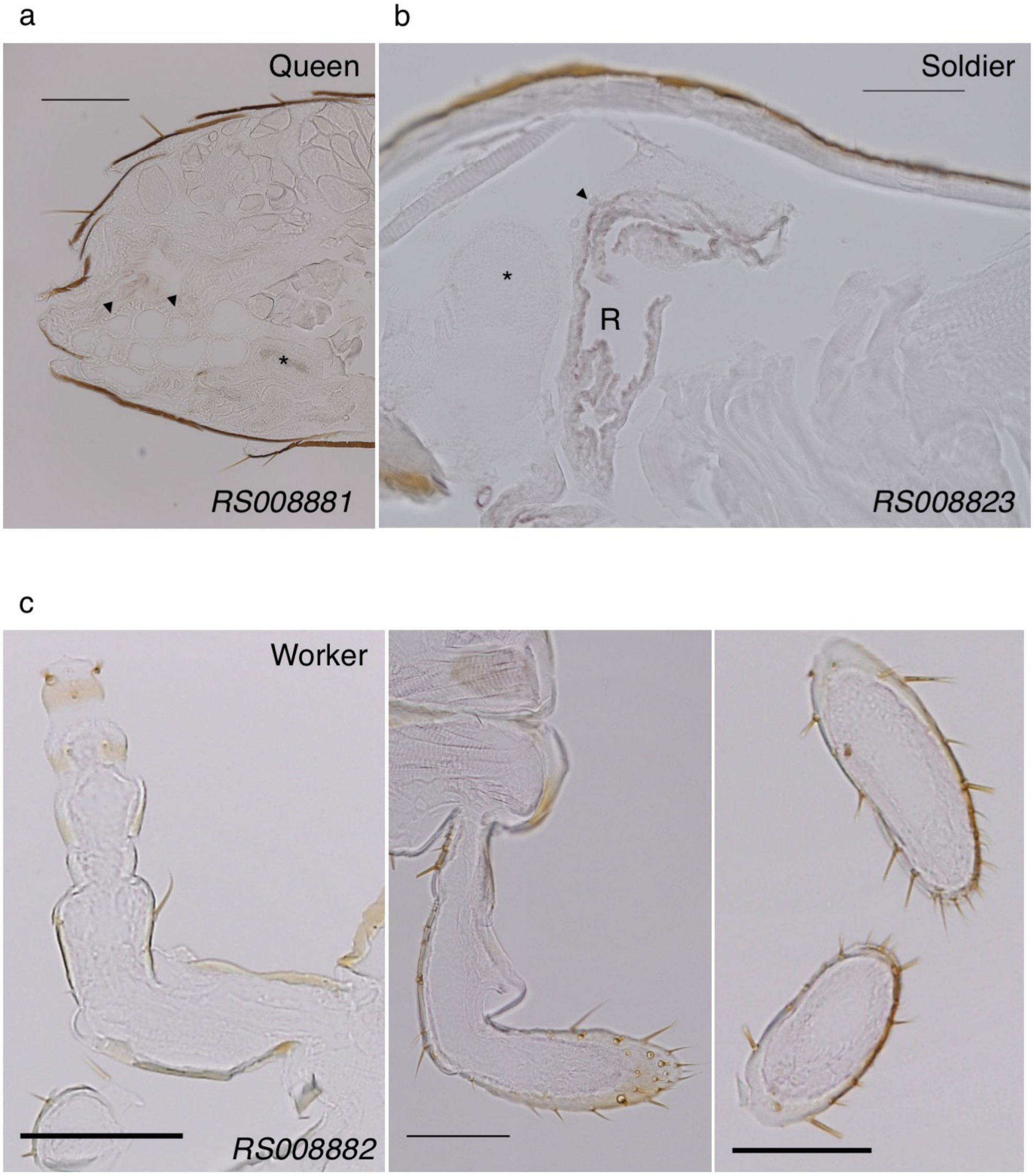


**Fig. S6.** *In situ* hybridization with *lipocalin* mRNA sense probe (negative control). (a) Vertical cryosection of the queen abdomen subjected to *in situ* hybridization with a sense DIG-labeled *RS008881* mRNA probe. Arrowheads indicate the accessory gland cell layer (stained dark with an antisense probe in Fig. 3d). Asterisk indicates the spermatica containing sperms. Bar = 0.2 mm. (b) Vertical cryosection of the soldier head subjected to *in situ* hybridization with a sense DIG-labeled *RS008823* mRNA probe. The front of the head is on the left side. Arrowhead indicates the gland cell layer surrounding the frontal gland reservoir (R) (stained dark with an antisense probe in Fig. 3e). Asterisk indicates the brain. Bar = 0.1 mm. (c) Left to right: vertical cryosection of the worker antenna, horizontal cryosection of the worker labial palp (right palp) and maxillary palp [the last segment of left (upper) and right (lower) palp] subjected to *in situ* hybridization with a sense DIG-labeled *RS008882* mRNA probe. Bar = 0.1 mm.

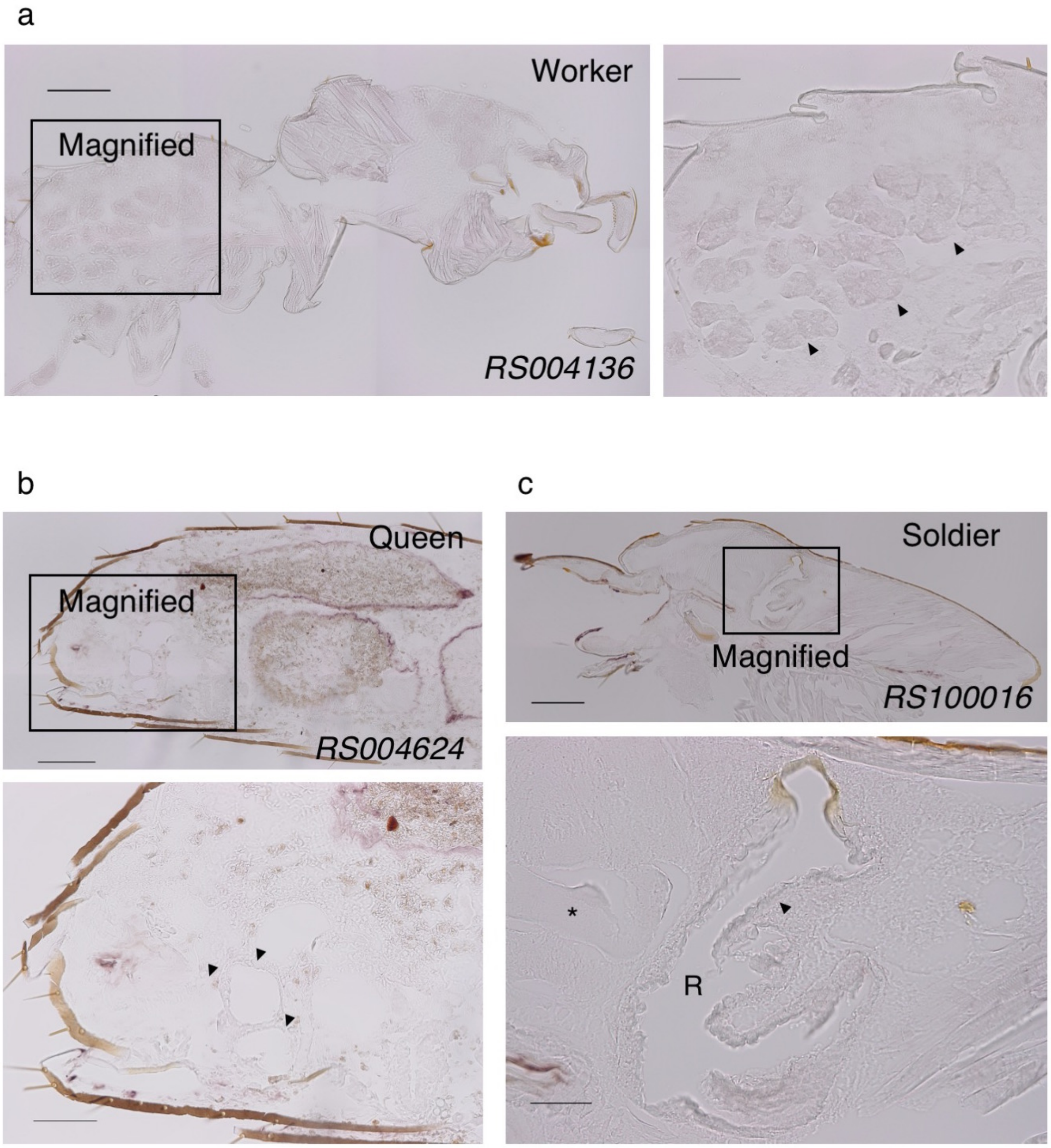


**Fig. S7.** *In situ* hybridization with *GH1* and *GGPPS* mRNA sense probe (negative control). (a) Vertical cryosection of the worker thorax subjected to *in situ* hybridization with a sense DIG-labeled *RS004136* mRNA probe. Magnified view is shown in the right panel. Arrowheads indicate the salivary gland cells (stained dark with an antisense probe in Fig. 4d). Bar = 0.2 (left) and 0.1 (right) mm. (b) Vertical cryosection of the queen abdomen subjected to *in situ* hybridization with a sense DIG-labeled *RS004624* mRNA probe. Magnified view is shown in the lower panel. Arrowheads indicate the accessory gland cell layers (stained dark with an antisense probe in Fig. 4f). Bar = 0.2 (upper) and 0.1 (lower) mm. (c) Vertical cryosection of the soldier head subjected to *in situ* hybridization with a sense DIG-labeled *RS100016* mRNA probe. Magnified view is shown in the lower panel. Arrowhead indicates the gland cell layer surrounding the frontal gland reservoir (R) (stained dark with an antisense probe in Fig. 6d). Asterisk indicates the brain. Bar = 0.2 (upper) and 0.05 (lower) mm.

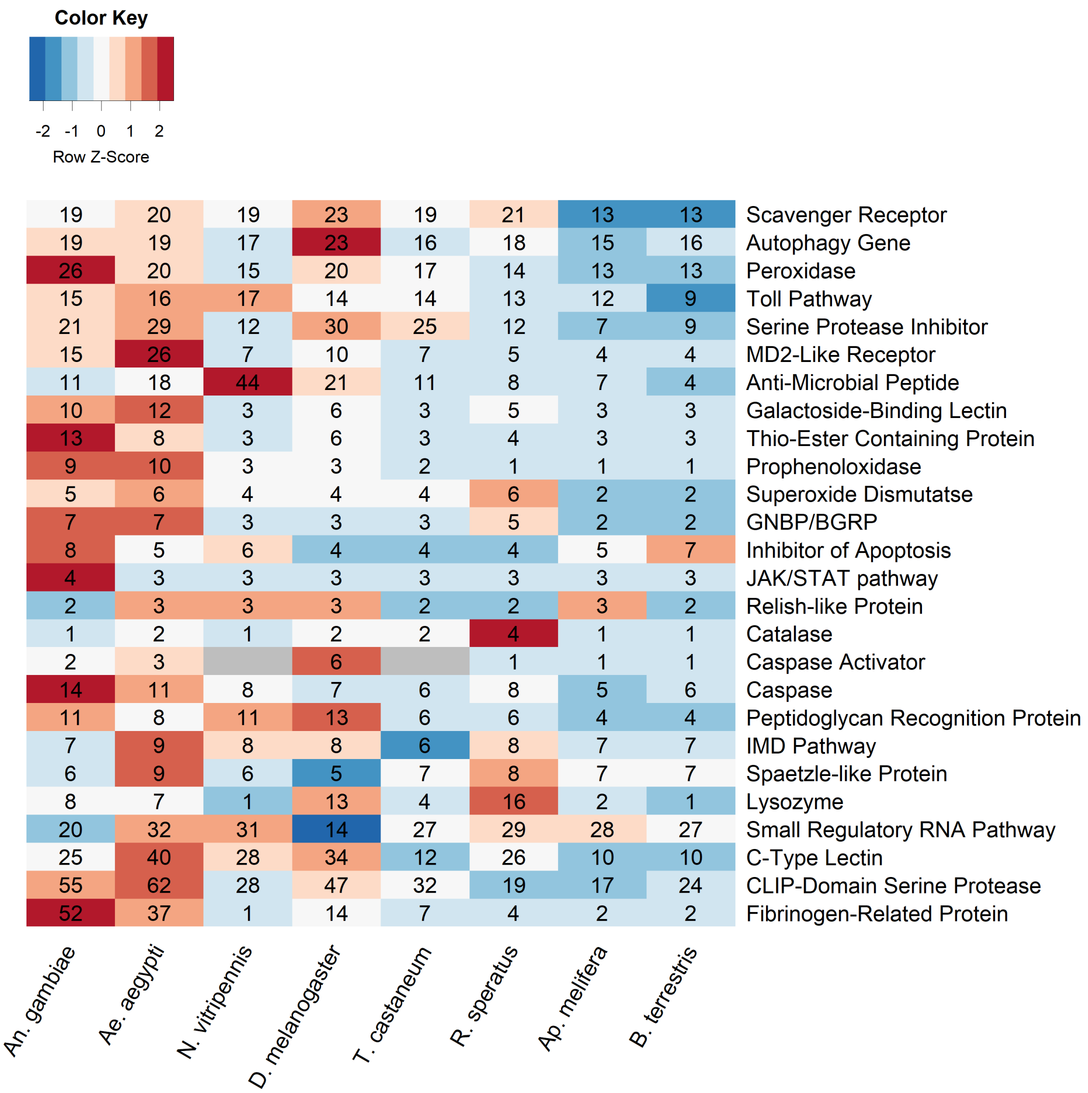


**Fig. S8.** Comparison of numbers of immune-related genes among 8 insect species. Species examined are *Anopheles gambiae*, *Aedes aegypti*, *Apis mellifera*, *Bombus terrestris*, *Drosophila melanogaster*, *Nasonia vitripennis*, *Reticulitermes speratus* and *Tribolium castaneum*. Gene numbers of *D. melanogaster*, *An. gambiae* and *Ae. aegypti* were obtained from ImmunoDB (Waterhouse et al. 2007), and those of *N. vitripennis*, *T. castaneum Ap. Mellifera* and *B. terrestris* were from Barribeau et al. (2015). Genes belonging to Caspase and Caspase Activator families were combined into Caspase, and Toll receptor and Toll pathway families into Toll pathway.

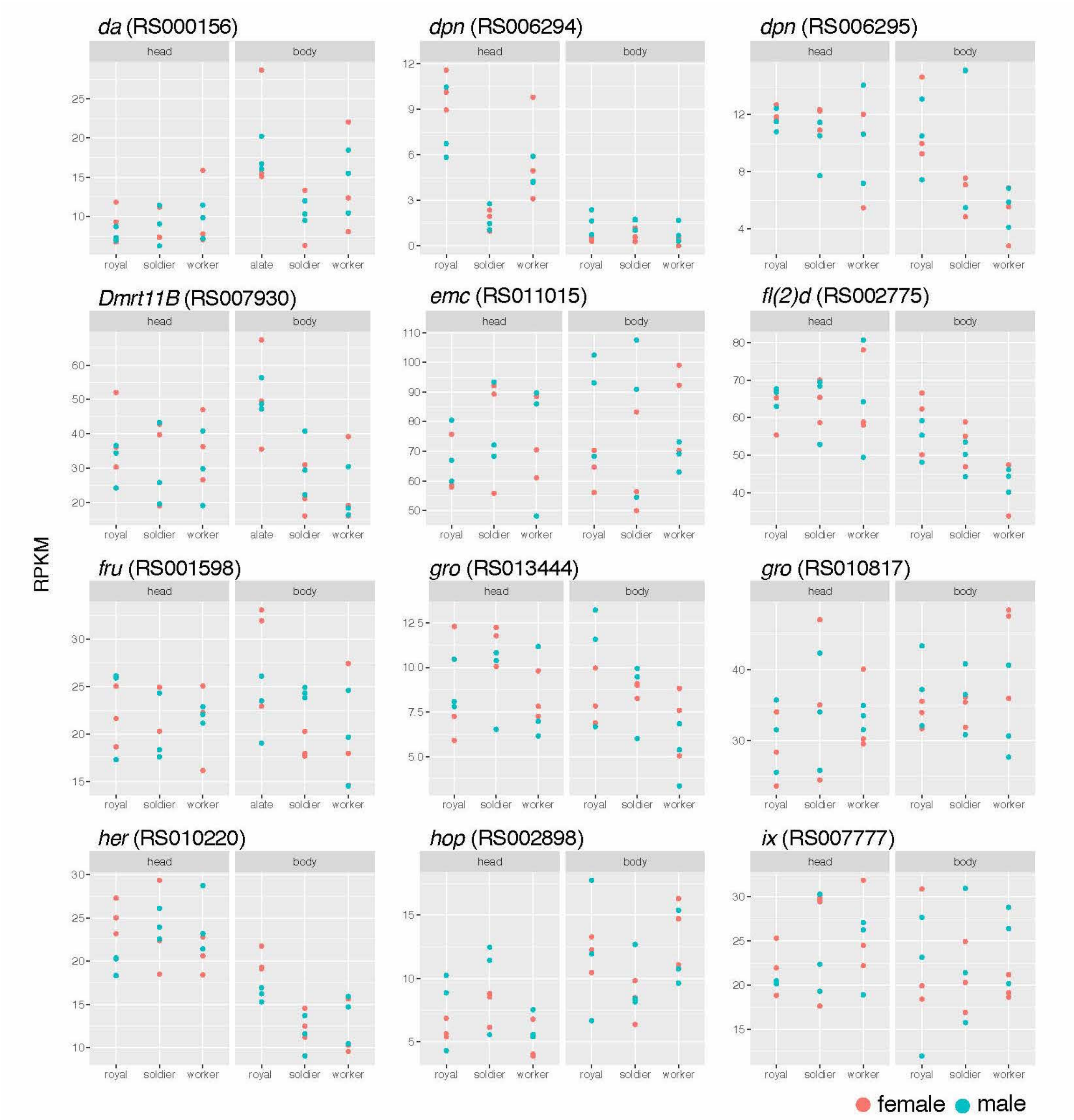


**Fig. S9.** Expression levels of sex determination genes among royals (reproductives), soldiers and workers in *Reticulitermes speratus*. Expression levels are indicated as RPKM calculated from RNA-sequencing analysis. Orange and blue points indicate females and males, respectively.

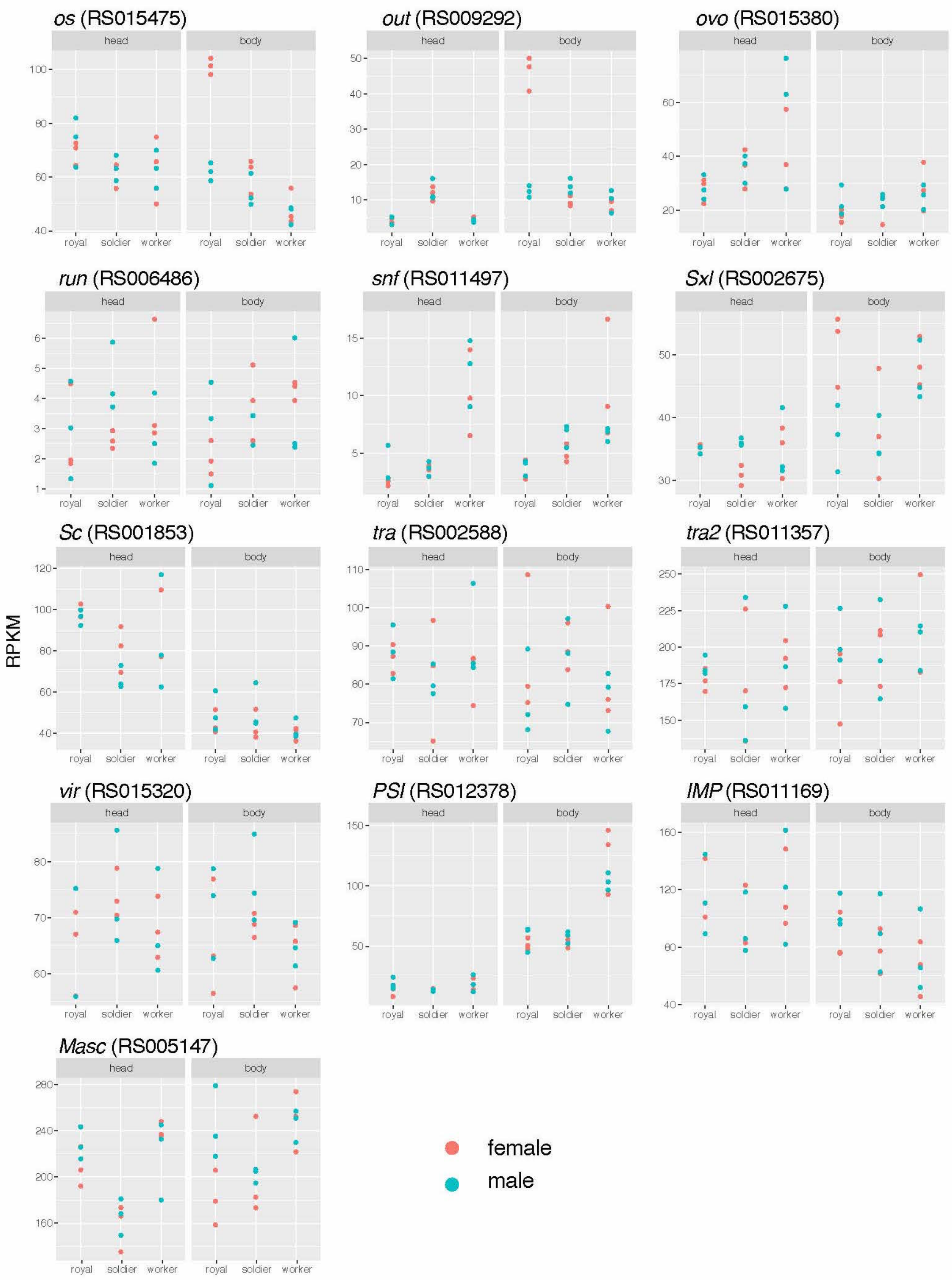


**Fig. S9.** *Continued*

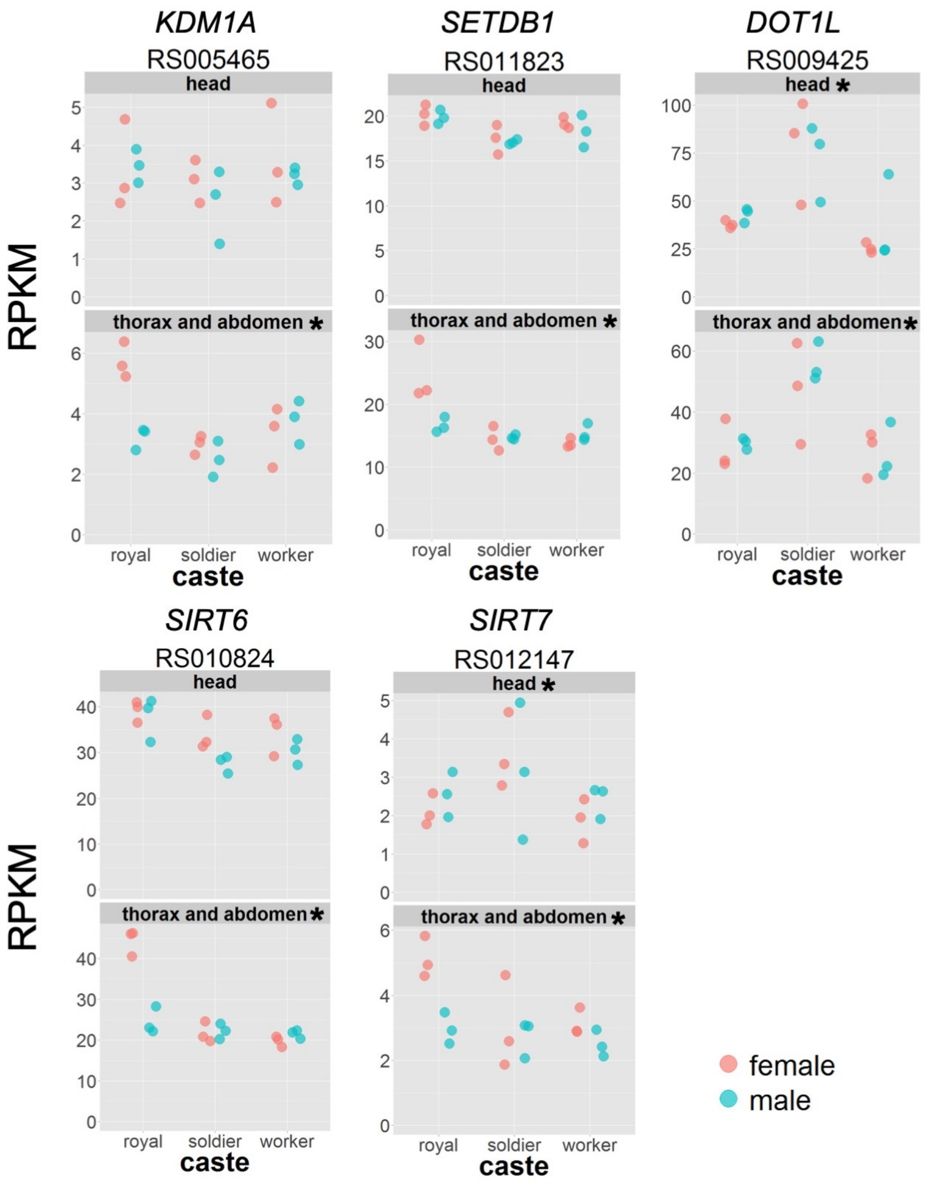


**Fig. S10.** Expression levels of histone modifying enzyme genes among royals (reproductives), soldiers and workers in *Reticulitermes speratus*. Expression levels are indicated as RPKM calculated from RNA-sequencing analysis. Orange and blue points indicate females and males, respectively. *KDM1A* (*RS005465*) encodes histone demethylase, *SETDB1* (*RS011823*) and *DOT1L* (*RS009425*) encode histone methyltransferases, and *SIRT6* (*RS010824*) and *SIRT7* (*RS012147*) encode histone deacetylases. These genes show the significant differences among castes in heads and/or thorax and abdomen samples (*FDR < 0.05).

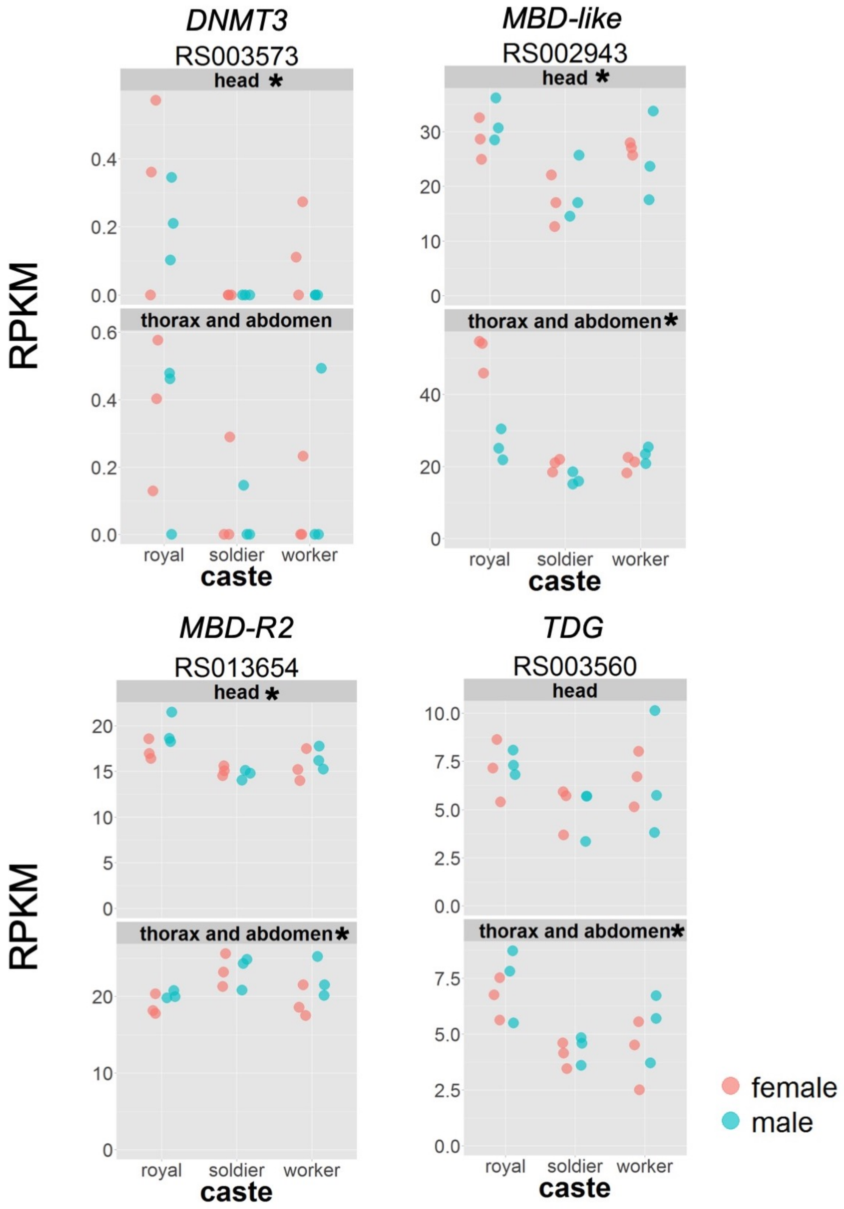


**Fig. S11.** Expression levels of DNA methylation-related genes among royals (reproductives), soldiers and workers in *Reticulitermes speratus*. Expression levels are indicated as RPKM calculated from RNA-sequencing analysis. Orange and blue points indicate females and males, respectively. All these 4 genes show the significant differences among castes in heads and/or thorax and abdomen samples (*FDR < 0.05).

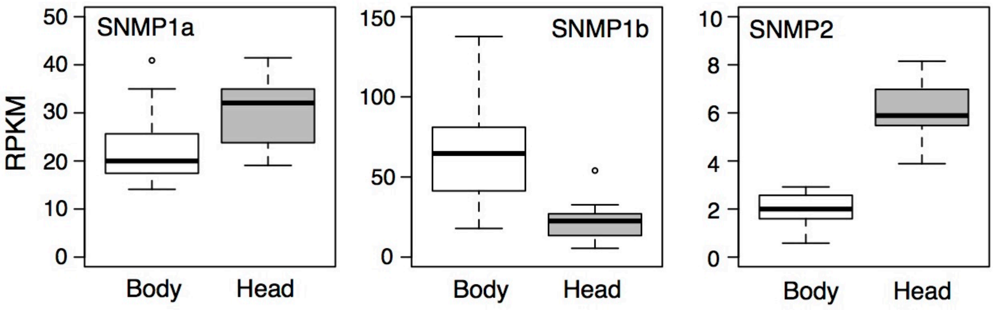


**Fig. S12.** Expression levels of sensory neuron membrane protein (SNMP) genes in the heads and bodies (throax + abdomen) of *Reticulitermes speratus*. Expression levels are indicated as RPKM calculated from RNA-sequencing analysis.

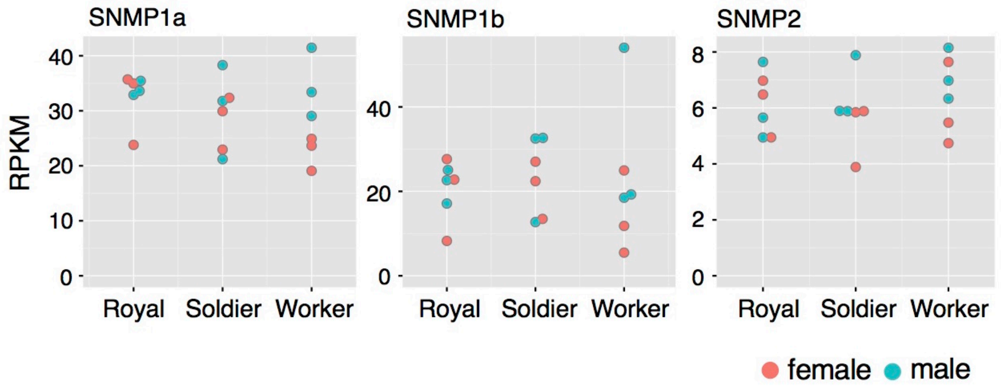


**Fig. S13.** Expression levels of sensory neuron membrane protein (SNMP) genes among the heads of royals (reproductives), soldiers and workers in *Reticulitermes speratus*. Expression levels are indicated as RPKM calculated from RNA-sequencing analysis. Orange and blue points indicate females and males, respectively.

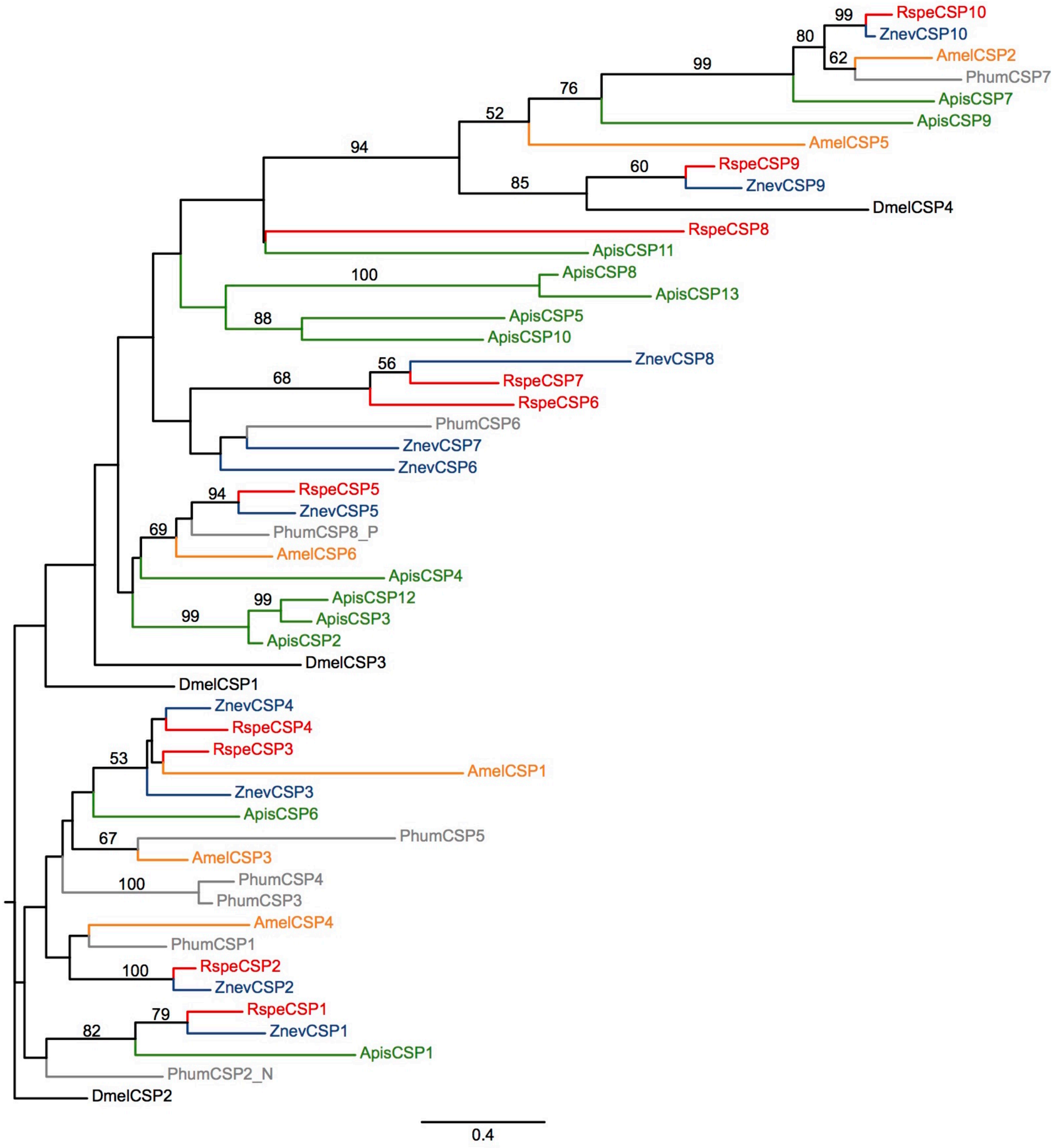


**Fig. S14.** Maximal likelihood (ML) tree of chemosensory protein (CSP) homologs based on the amino acid sequences obtained with PROTGAMMALG model. Sequences from *Drosophila melanogaster* (Dmel), *Apis mellifera* (Amel), *Acyrthosiphon pisum* (Apis), *Pediculus humanus* (Phum), *Zootermopsis nevadensis* (Znev) and *Reticultermes speratus* (Rspe) were included. Numbers above branches indicate the bootstrap probabilities more than 50% based on 100 ML trees.

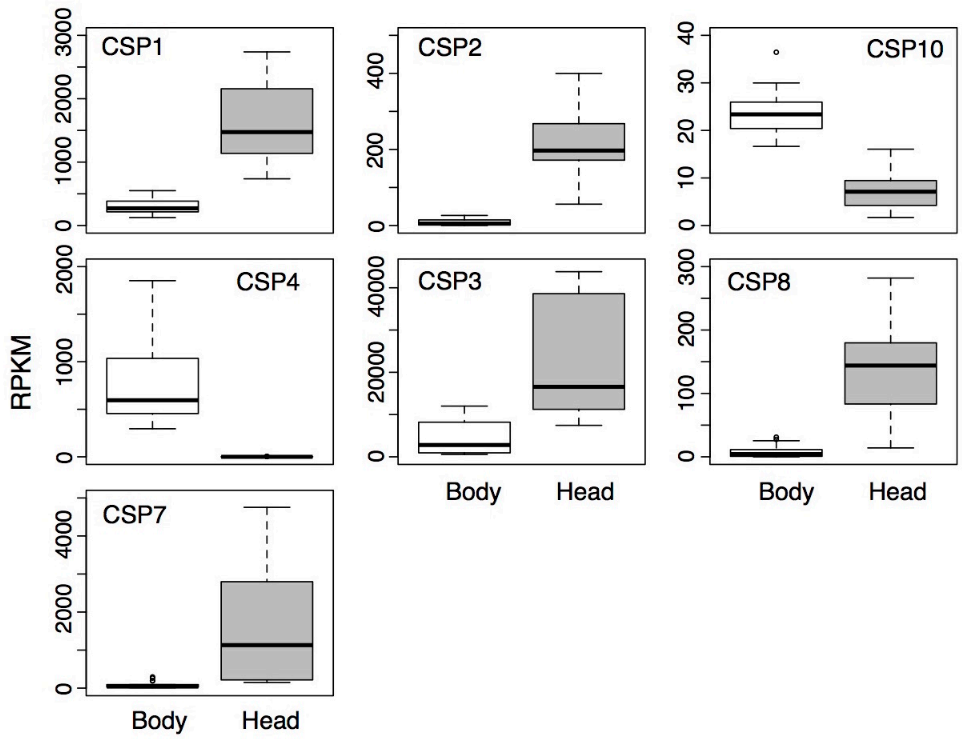


**Fig. S15.** Expression levels of chemosensory protein (CSP) genes in the heads and bodies (throax + abdomen) of *Reticulitermes speratus*. Expression levels are indicated as RPKM calculated from RNA-sequencing analysis.

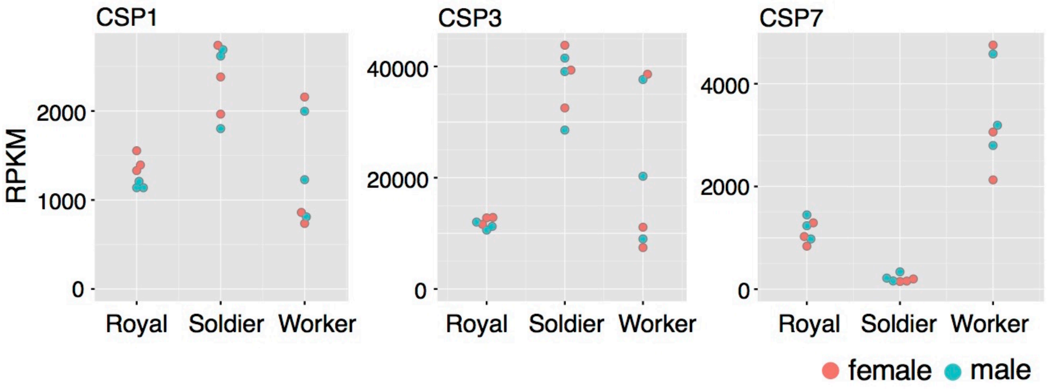


**Fig. S16.** Expression levels of chemosensory protein (CSP) genes among the heads of royals (reproductives), soldiers and workers in *Reticulitermes speratus*. Expression levels are indicated as RPKM calculated from RNA-sequencing analysis. Orange and blue points indicate females and males, respectively.

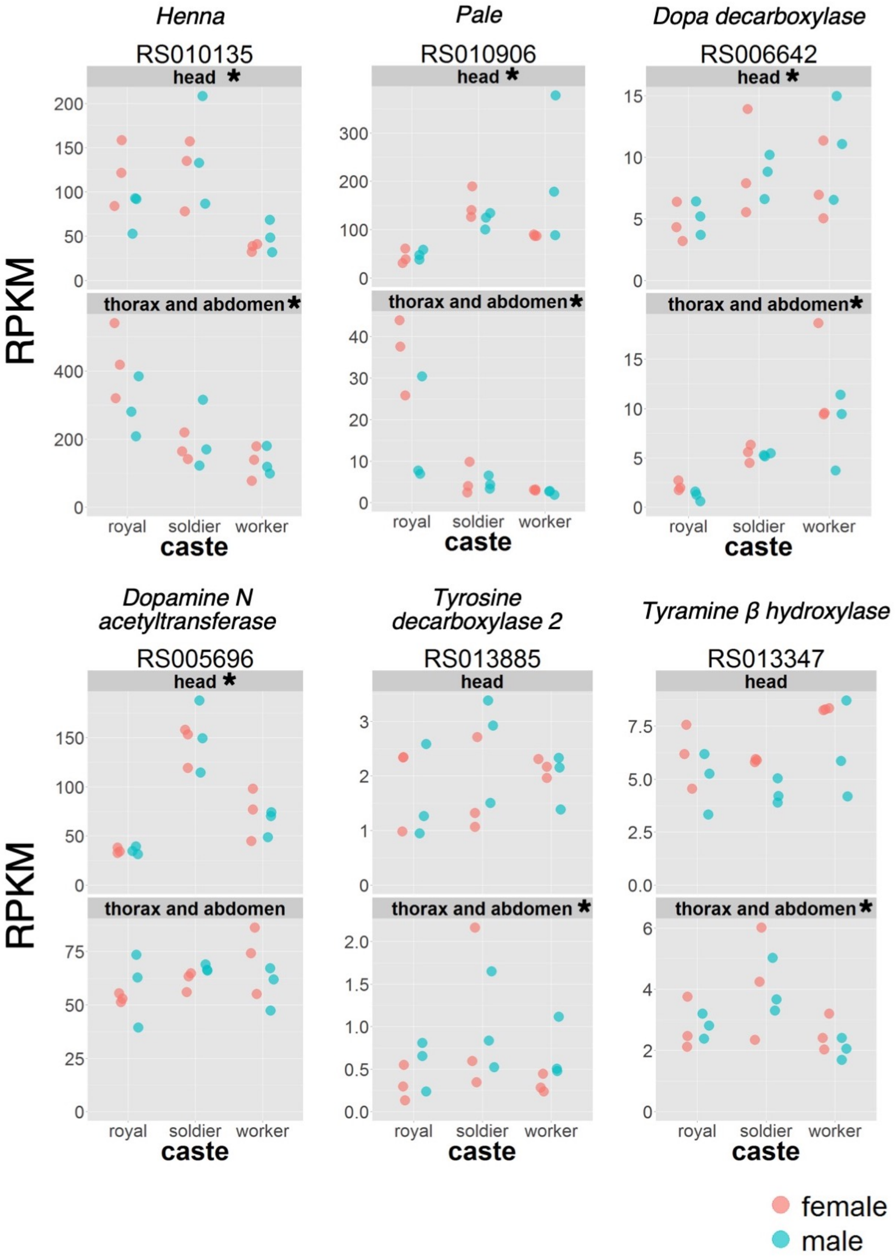


**Fig. S17.** Expression levels of biosynthetic genes of biogenic amines among royals (reproductives), soldiers and workers in *Reticulitermes speratus*. Expression levels are indicated as RPKM calculated from RNA-sequencing analysis. Orange and blue points indicate females and males, respectively. All these 6 genes show the significant differences among castes in heads and/or thorax and abdomen samples (*FDR < 0.05).

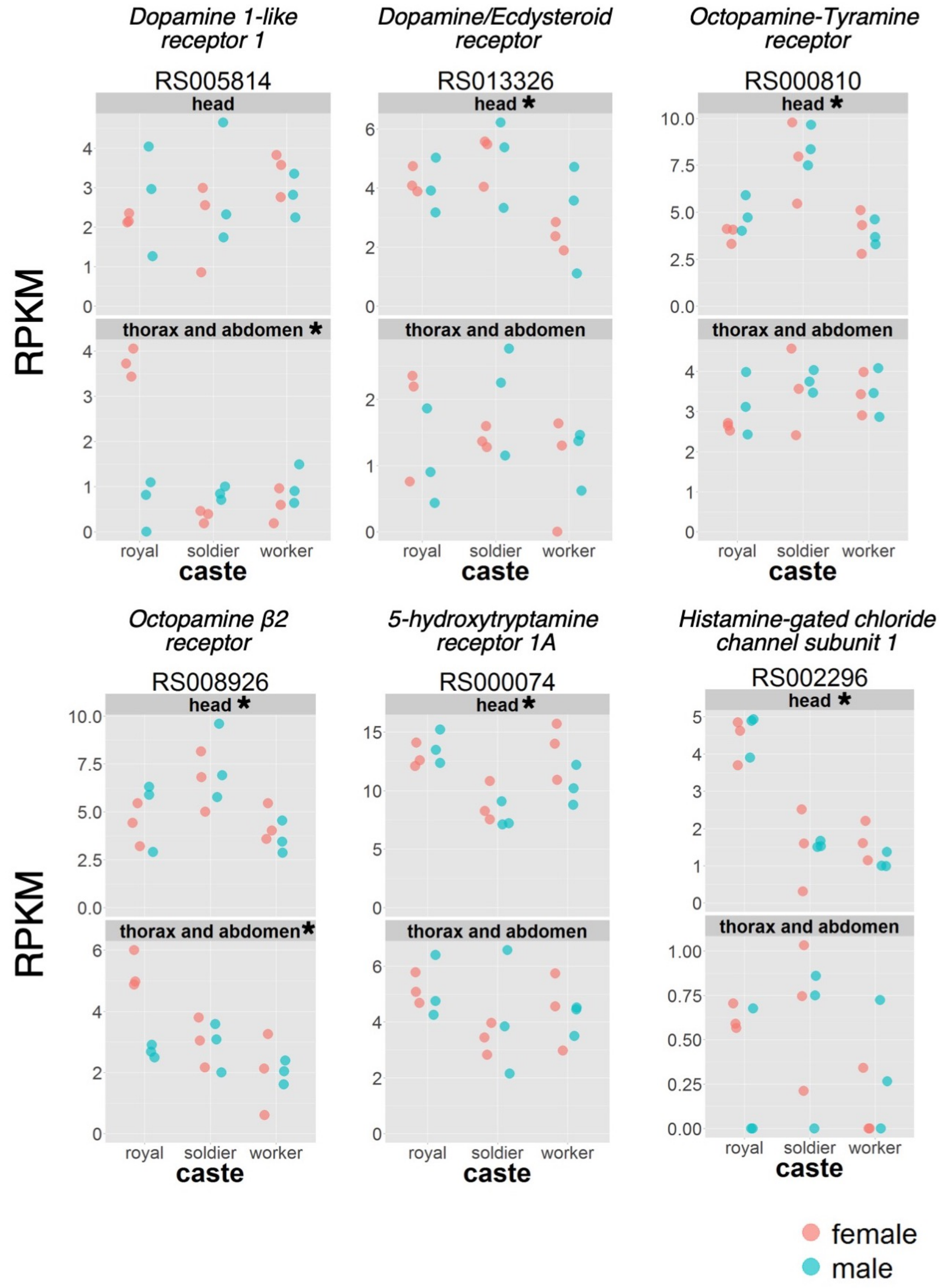


**Fig. S18.** Expression levels of receptor genes of biogenic amines among royals (reproductives), soldiers and workers in *Reticulitermes speratus*. Expression levels are indicated as RPKM calculated from RNA-sequencing analysis. Orange and blue points indicate females and males, respectively. All these 6 genes show the significant differences among castes in heads and/or thorax and abdomen samples (*FDR < 0.05).

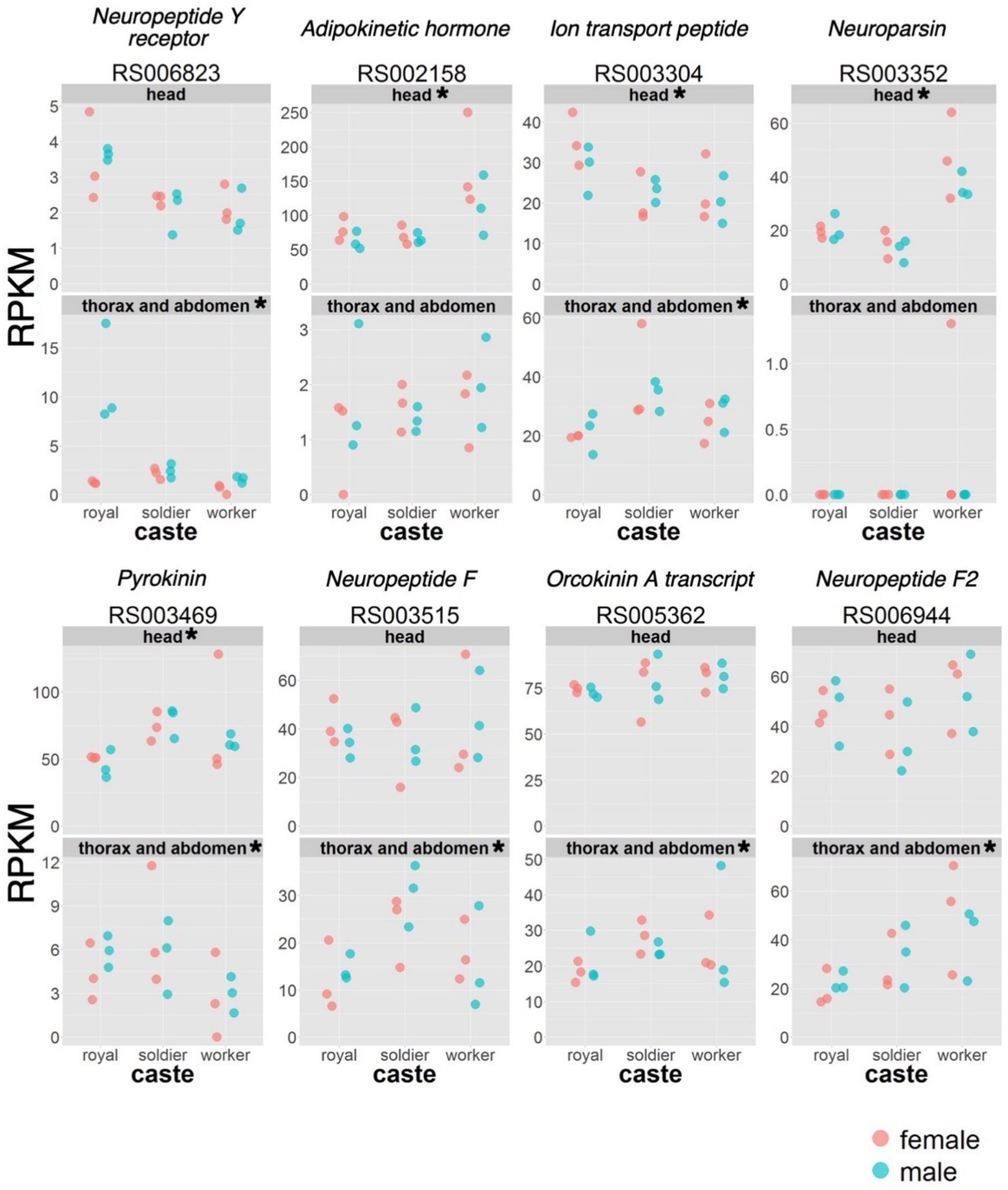


**Fig. S19.** Expression levels of neuropeptide genes among royals (reproductives), soldiers and workers in *Reticulitermes speratus*. Expression levels are indicated as RPKM calculated from RNA-sequencing analysis. Orange and blue points indicate females and males, respectively. All these 15 genes show the significant differences among castes in heads and/or thorax and abdomen samples (*FDR < 0.05).

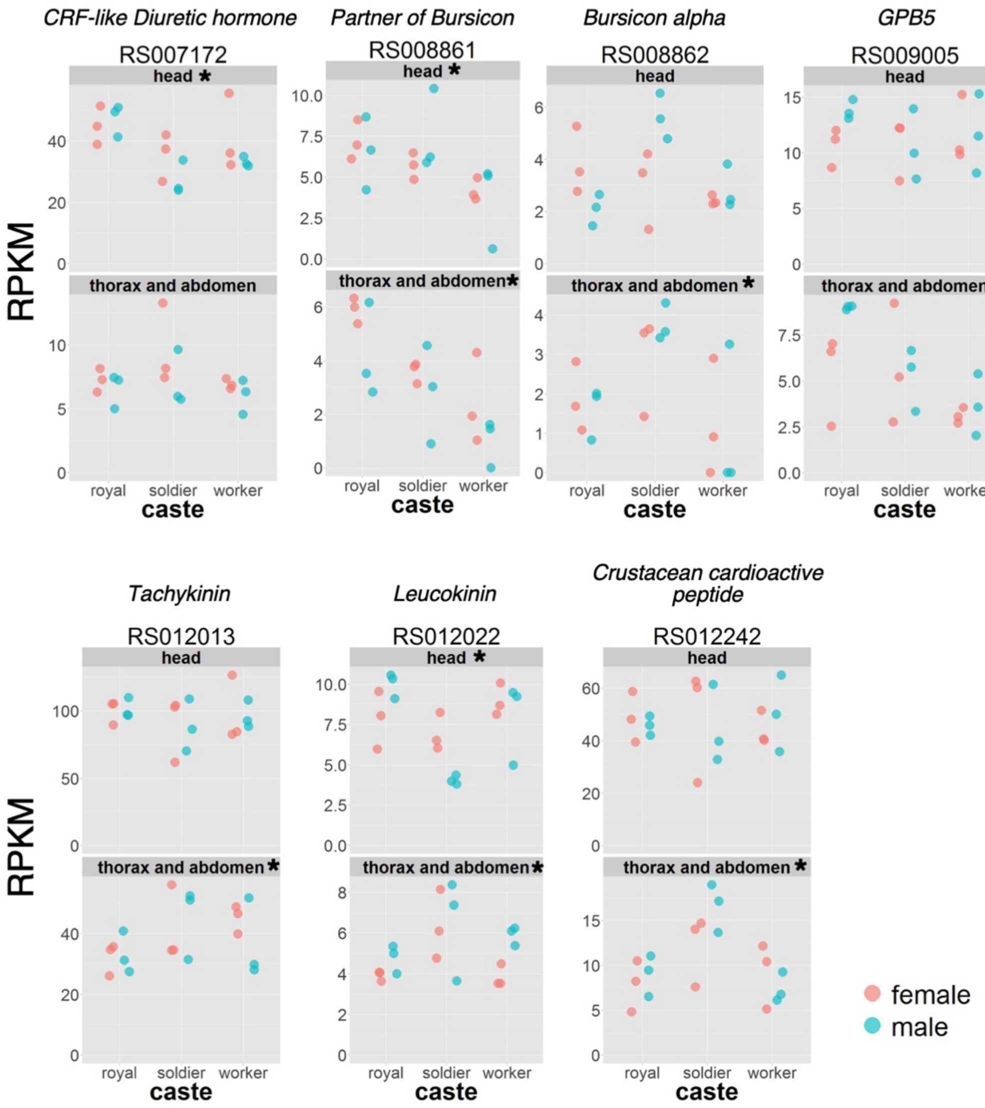


**Fig. S19.** *Continued*.

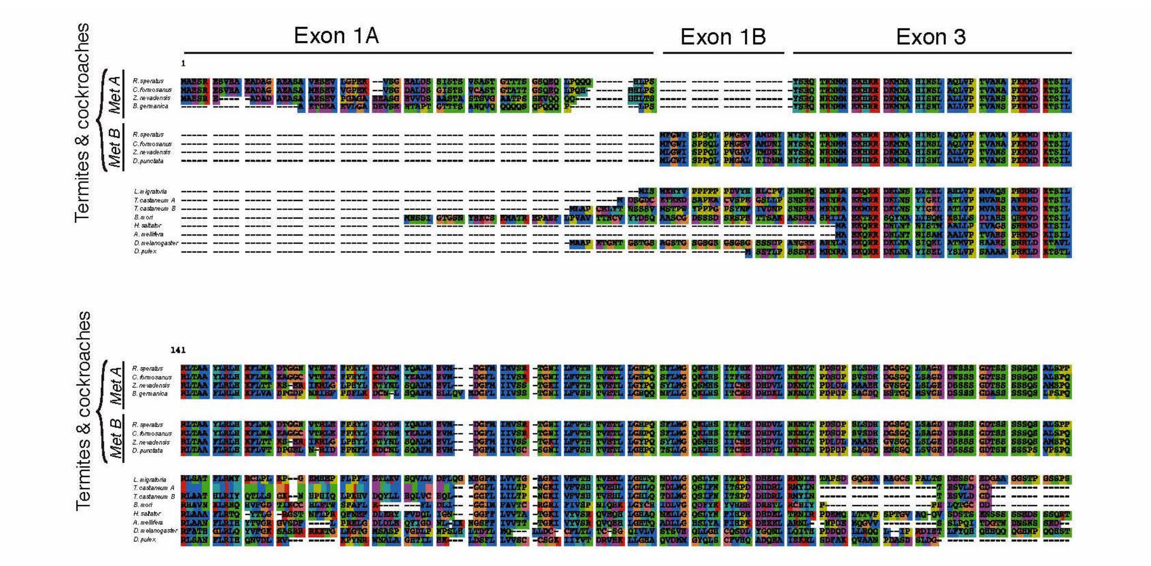


**Fig. S20.** Amino acid alignment of JH receptor gene (*Methoprene-torelant*; *Met*). Species (gene ID) examined are *Reticulitermes speratus* (RS010120), *Coptotermes formosanus* (A: GFG37549.1, B: GFG37551.1), *Zootermopsis nevadensis* (BAR92640.1), *Blattera germanica* (CDO33887.1), *Diploptera punctata* (AIM47235.1), *Locusta migratoria* (AHA42531.1), *Tribolium castaneum* (NP_001092812.1, XP_008191439.1), *Bombyx mori* (NP_001108458.1), *Harpegnathos saltator* (EFN85711.1), *Apis mellifera* (XP_395005.5), *Drosophila melanogaster* (NP_001285132.1) and *Daphnia pulex* (BAM83853.1).

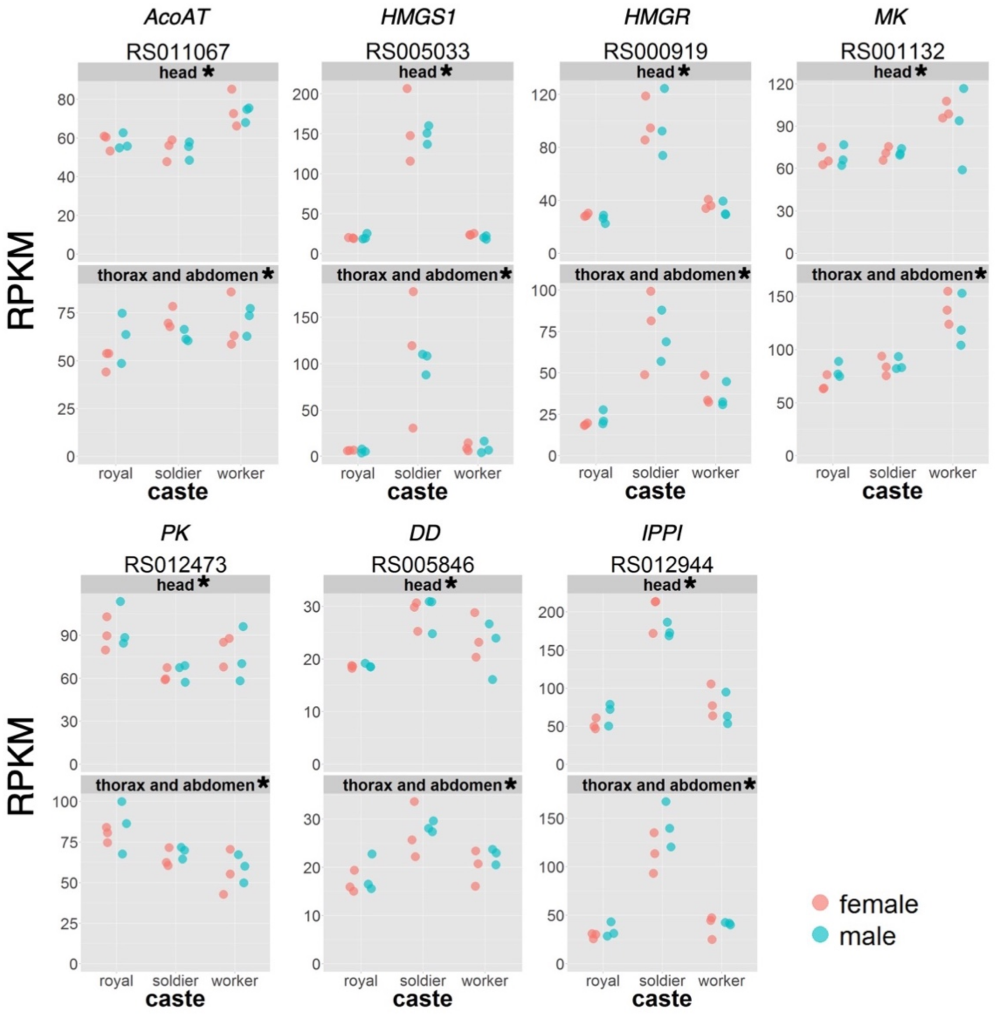


**Fig. S21.** Expression levels of JH biosynthetic genes (early steps) among royals (reproductives), soldiers and workers in *Reticulitermes speratus*. Expression levels are indicated as RPKM calculated from RNA-sequencing analysis. Orange and blue points indicate females and males, respectively. All these 7 genes show the significant differences among castes in heads and/or thorax and abdomen samples (*FDR < 0.05).

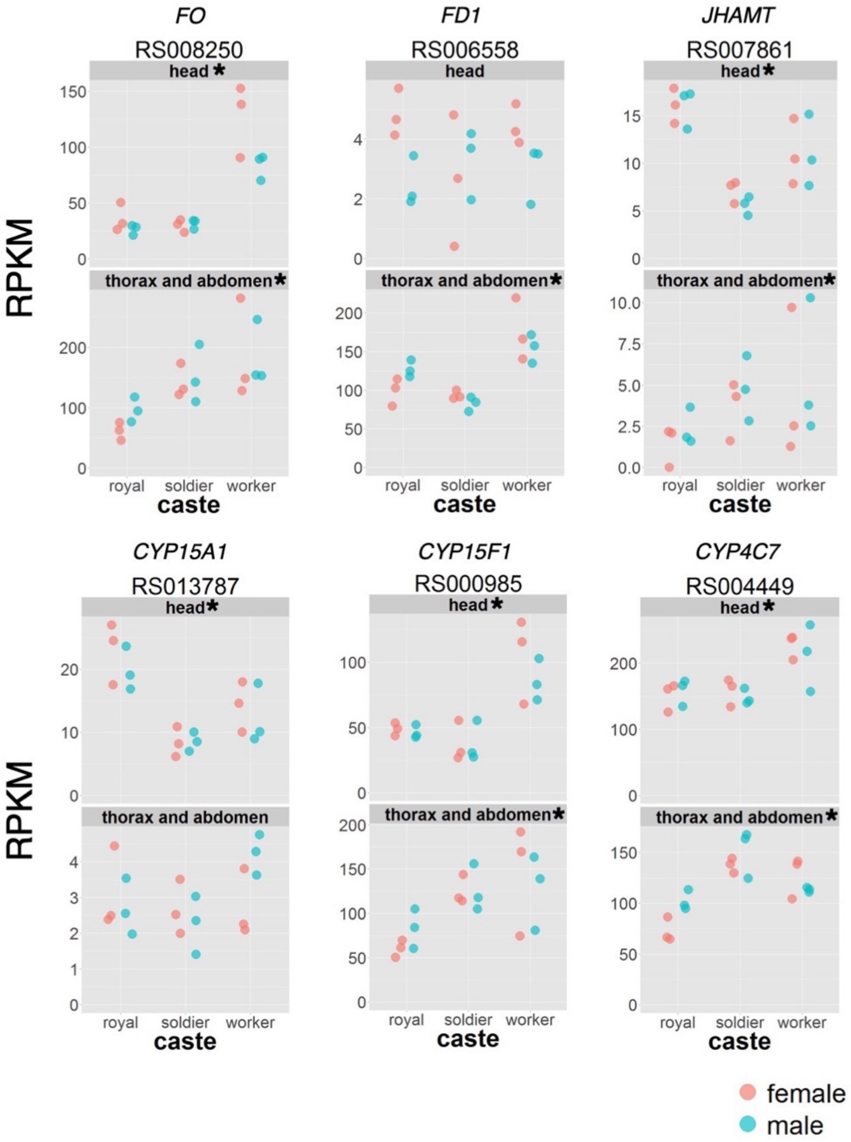


**Fig. S22.** Expression levels of JH biosynthetic genes (late steps) among royals (reproductives), soldiers and workers in *Reticulitermes speratus*. Expression levels are indicated as RPKM calculated from RNA-sequencing analysis. Orange and blue points indicate females and males, respectively. All these 6 genes show the significant differences among castes in heads and/or thorax and abdomen samples (*FDR < 0.05).

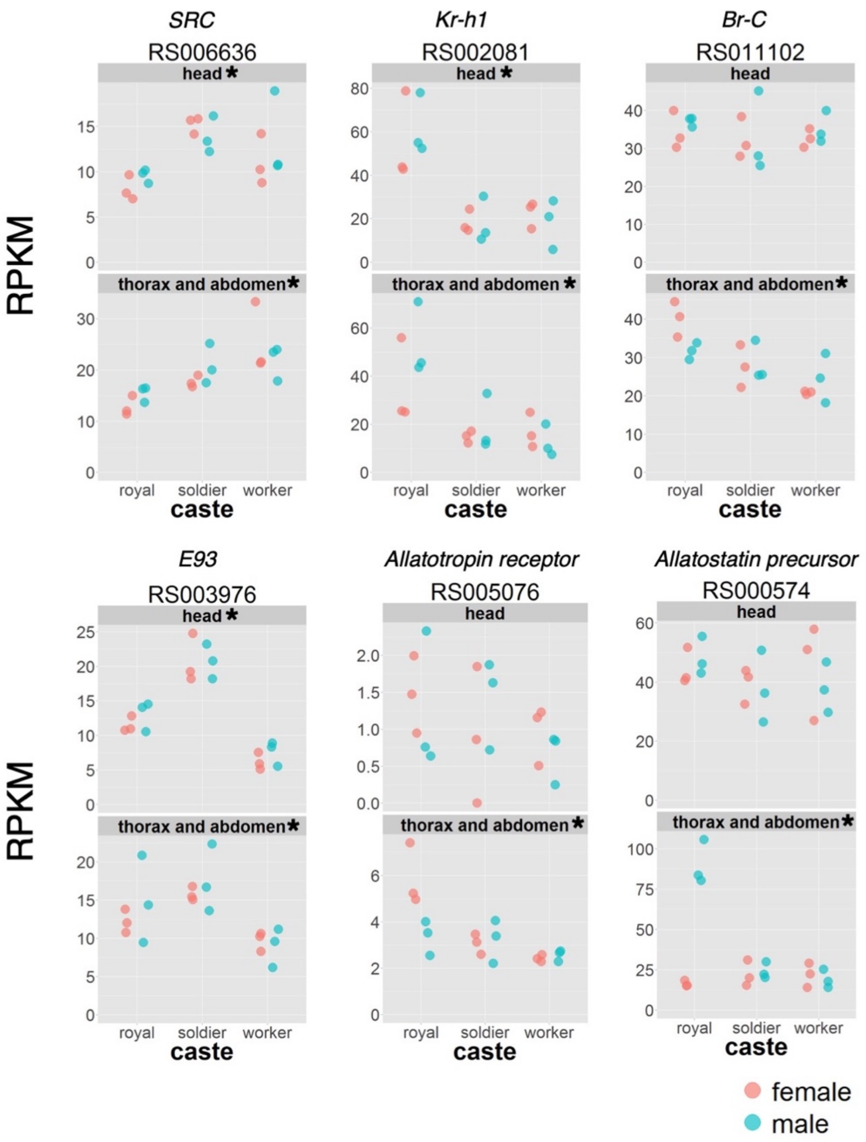


**Fig. S23.** Expression levels of JH signaling and neuropeptide related genes among royals (reproductives), soldiers and workers in *Reticulitermes speratus*. Expression levels are indicated as RPKM calculated from RNA-sequencing analysis. Orange and blue points indicate females and males, respectively. All these 6 genes show the significant differences among castes in heads and/or thorax and abdomen samples (*FDR < 0.05).

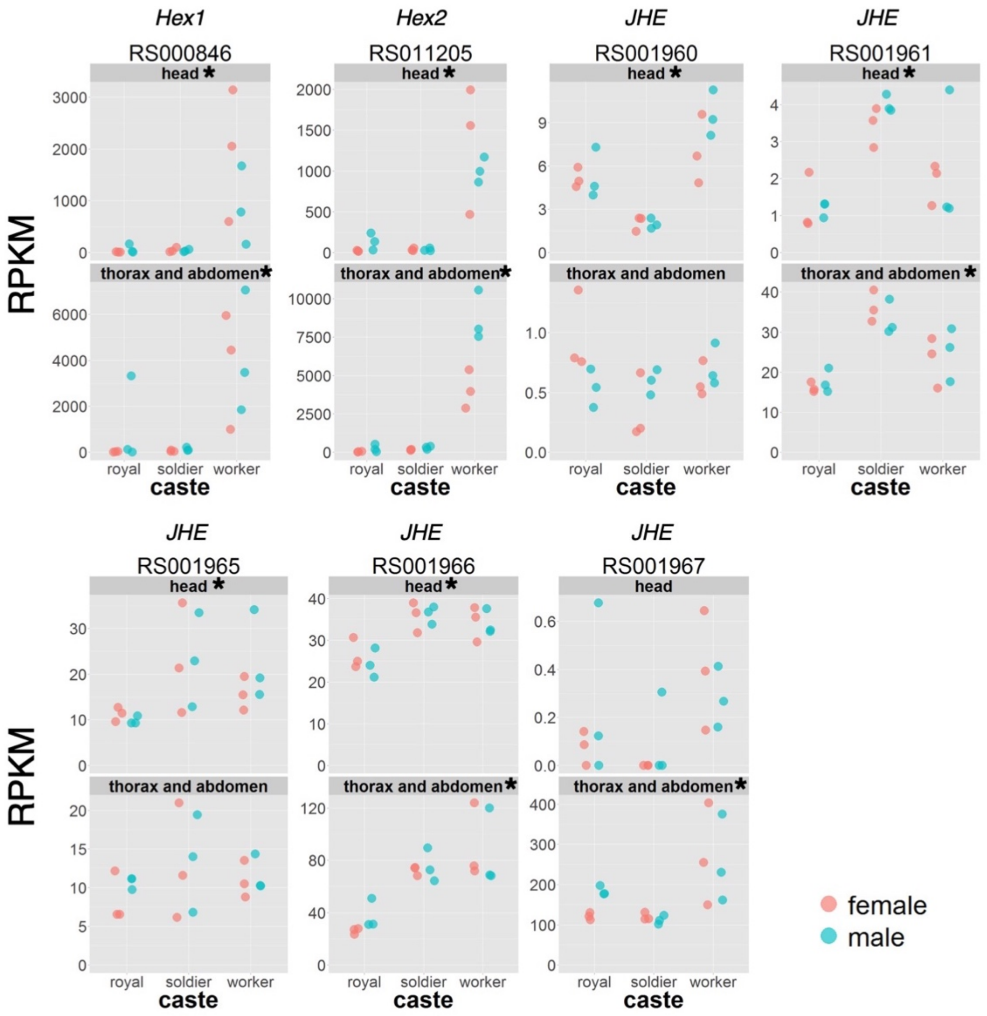


**Fig. S24.** Expression levels of JH binding and degradation genes among royals (reproductives), soldiers and workers in *Reticulitermes speratus*. Expression levels are indicated as RPKM calculated from RNA-sequencing analysis. Orange and blue points indicate females and males, respectively. All these 7 genes show the significant differences among castes in heads and/or thorax and abdomen samples (*FDR < 0.05).

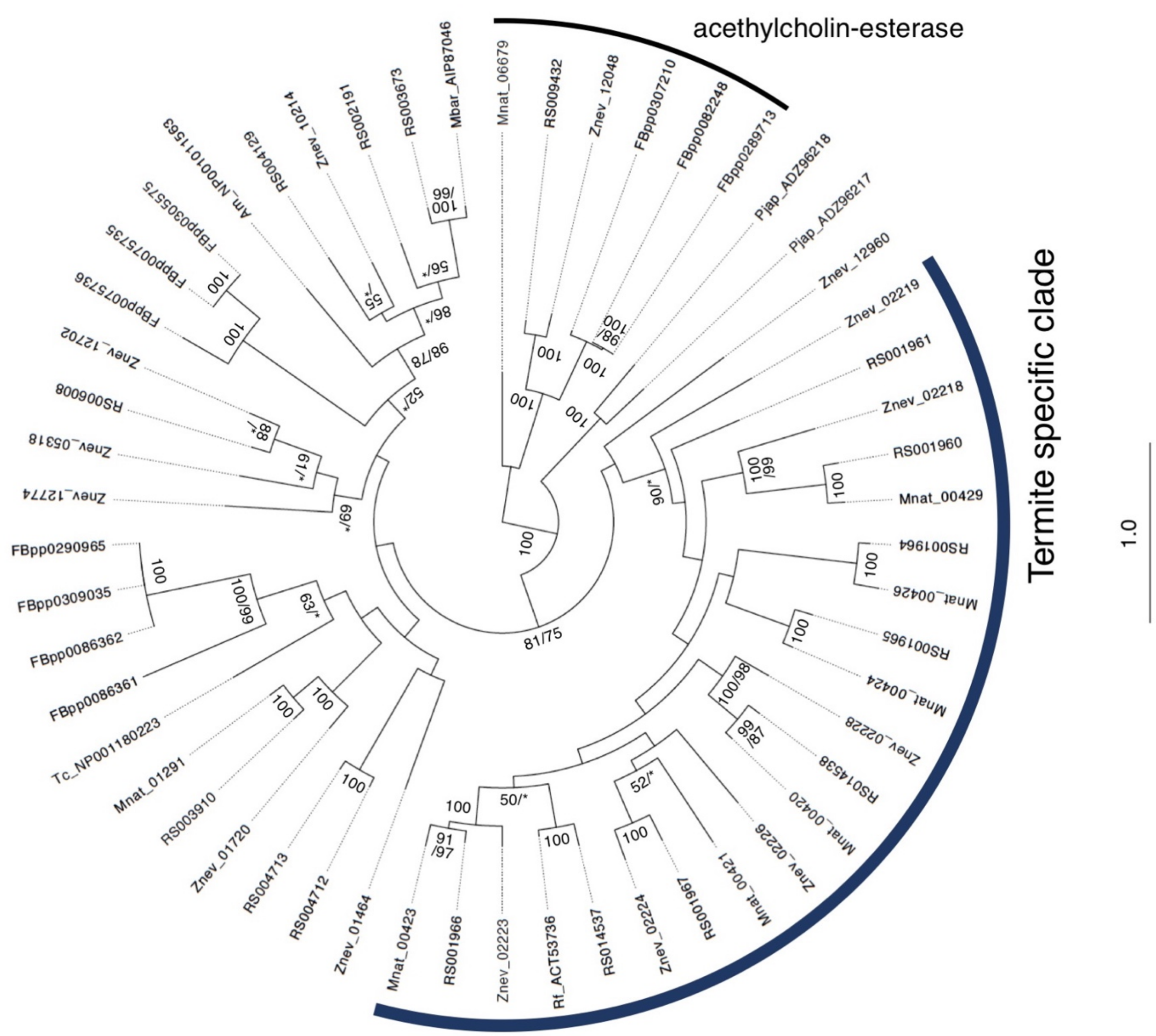


**Fig. S25.** Maximum likelihood (ML) tree of JH esterase (JHE) homologs based on the amino acid sequences with the highest log likelihood (-48105.8335) based on the Dayhoff model. Sequences from *Reticulitermes speratus* (RS), *R. flavipes* (Rf), *Zootermopsis nevadensis* (Znev), *Macrotermes natalensis* (Mnat), *M. barneyi* (Mbar), *Tribolium castaneum* (Tc), *Apis mellifera* (Am), *Pandalopsis japonica* (Pjap) and *Drosophila melanogaster* (FBpp) are included. Total 6 sequences of acethylcholin-esterases are used for outgroups. We also obtained estimations of tree topology under the Neighbor-joining (NJ) method based on the JTT+G model. Numbers above or below each branch indicate the bootstrap probabilities (BP) more than 50% (100 and 5,000 replicates in ML and NJ method, respectively). Only one number is given if BP was identical at that node. An asterisk indicates a node that was not supported by NJ method.

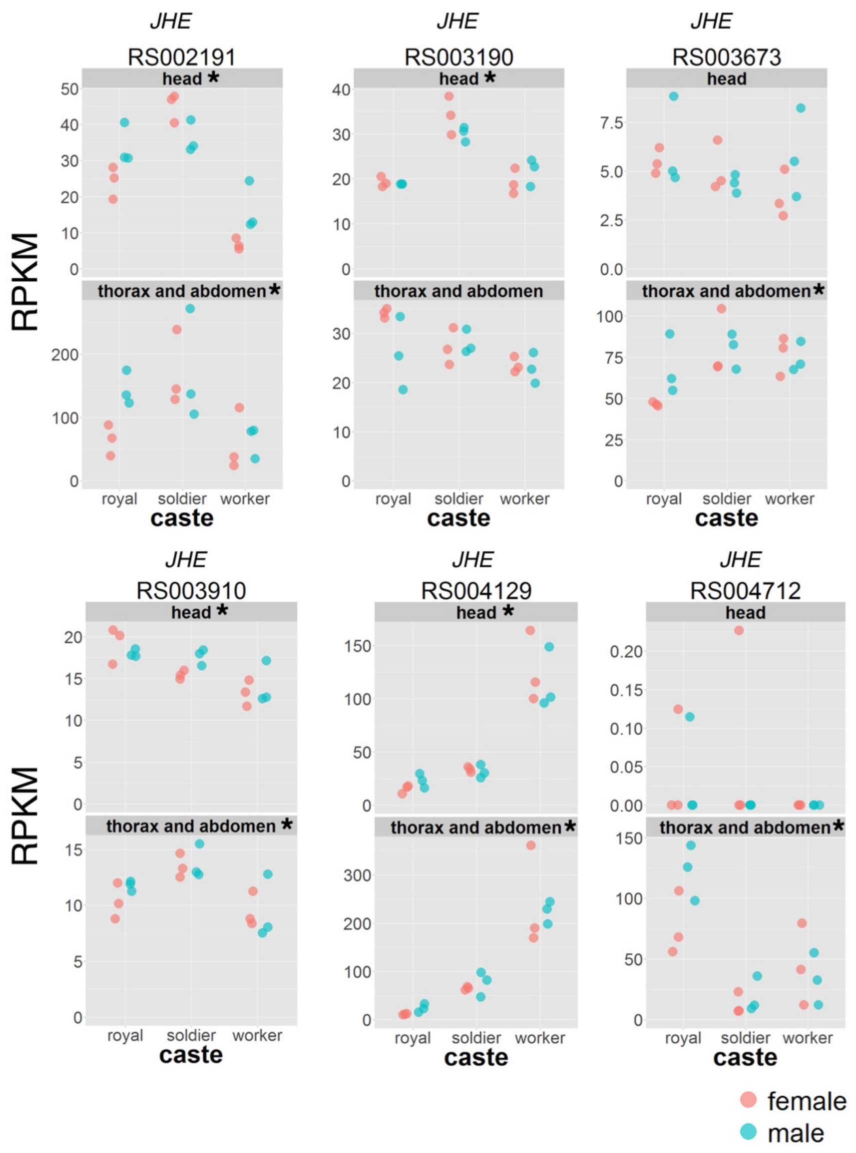


**Fig. S26.** Expression levels of JH degradation genes (*JHEs*) among royals (reproductives), soldiers and workers in *Reticulitermes speratus*. Expression levels are indicated as RPKM calculated from RNA-sequencing analysis. Orange and blue points indicate females and males, respectively. All these 12 genes show the significant differences among castes in heads and/or thorax and abdomen samples (*FDR < 0.05).

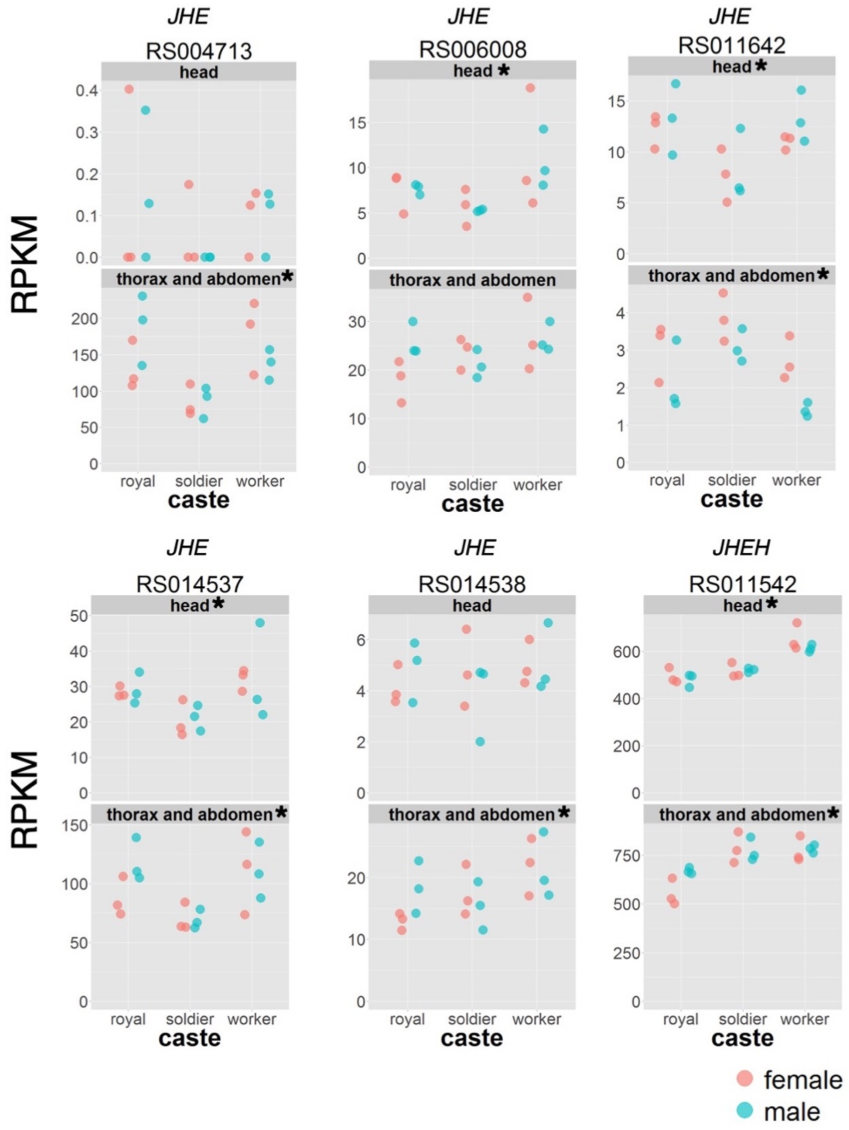


**Fig. S26.** *Continued*.

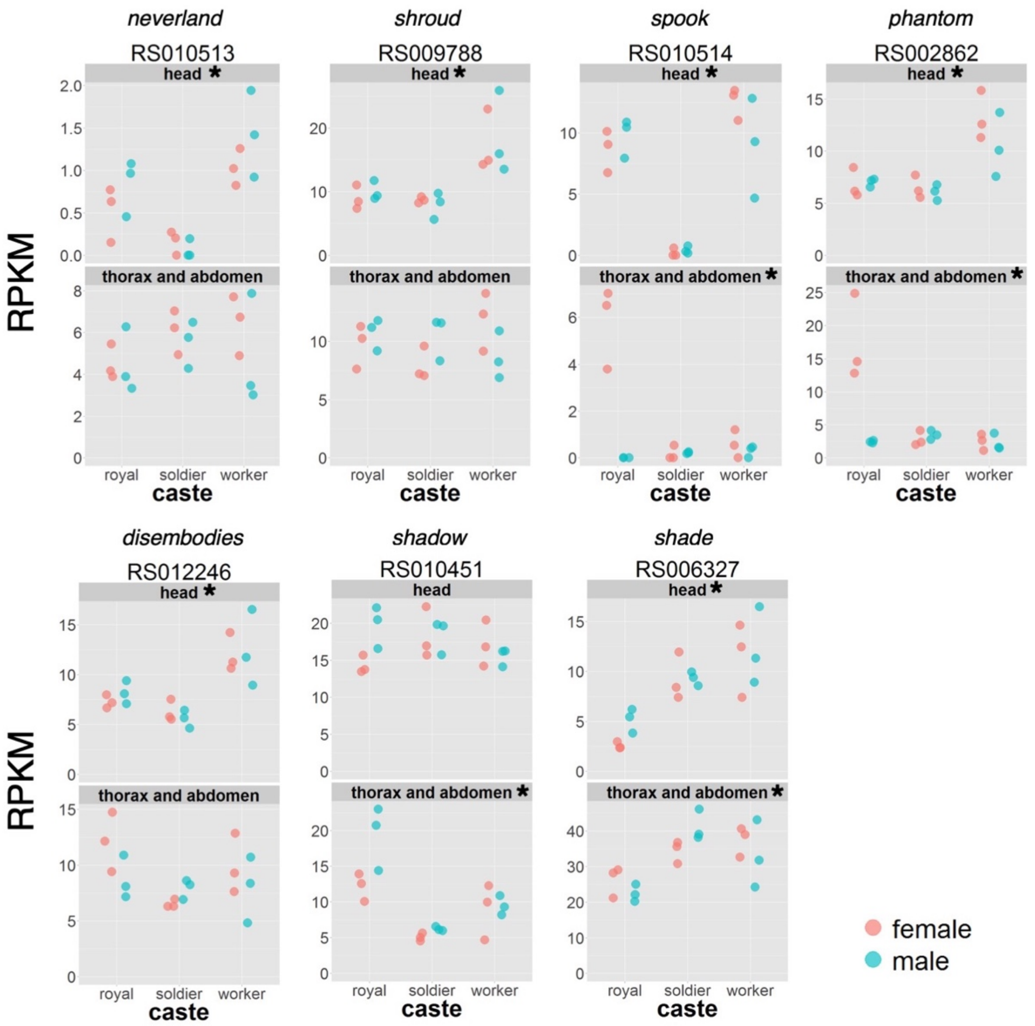


**Fig. S27.** Expression levels of ecdysone synthesis genes among royals (reproductives), soldiers and workers in *Reticulitermes speratus*. Expression levels are indicated as RPKM calculated from RNA-sequencing analysis. Orange and blue points indicate females and males, respectively. All these 7 genes show the significant differences among castes in heads and/or thorax and abdomen samples (*FDR < 0.05).

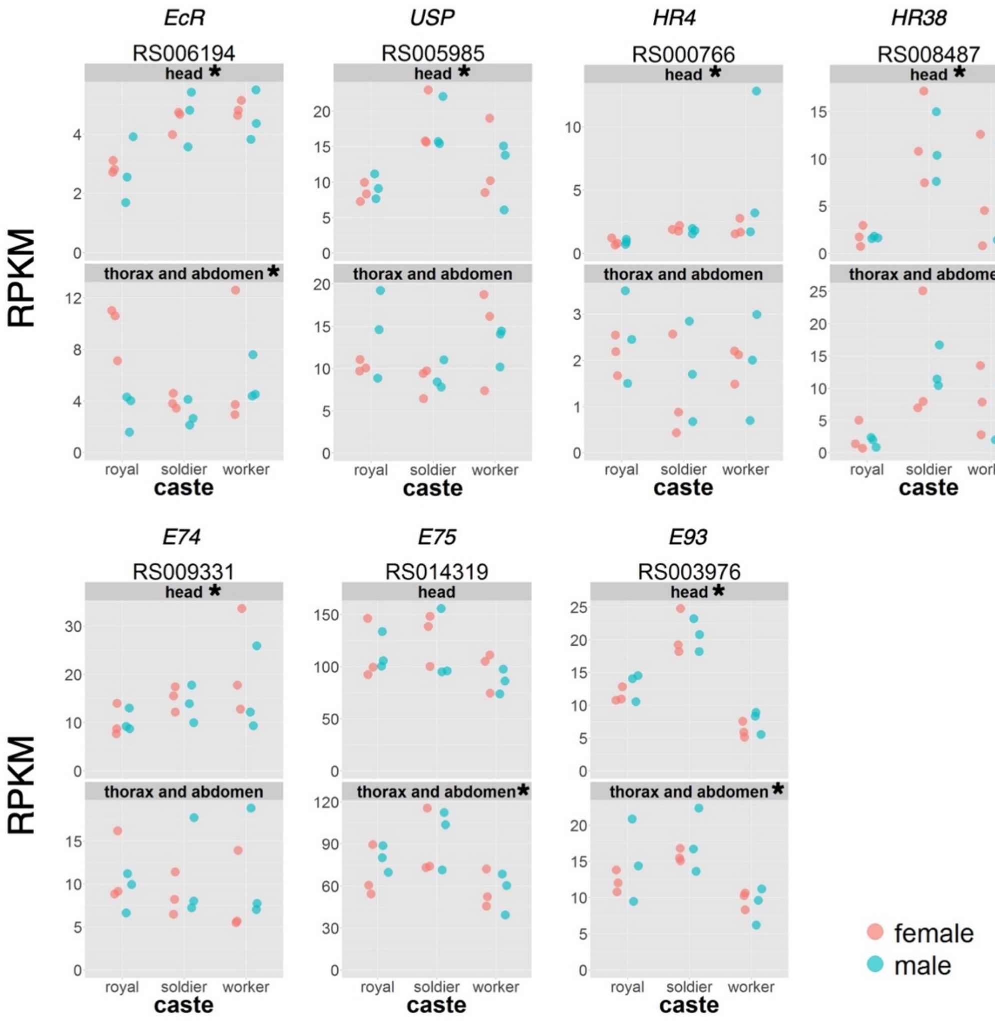


**Fig. S28.** Expression levels of ecdysone receptor and signaling genes among royals (reproductives), soldiers and workers in *Reticulitermes speratus*. Expression levels are indicated as RPKM calculated from RNA-sequencing analysis. Orange and blue points indicate females and males, respectively. All these 7 genes show the significant differences among castes in heads and/or thorax and abdomen samples (*FDR < 0.05).

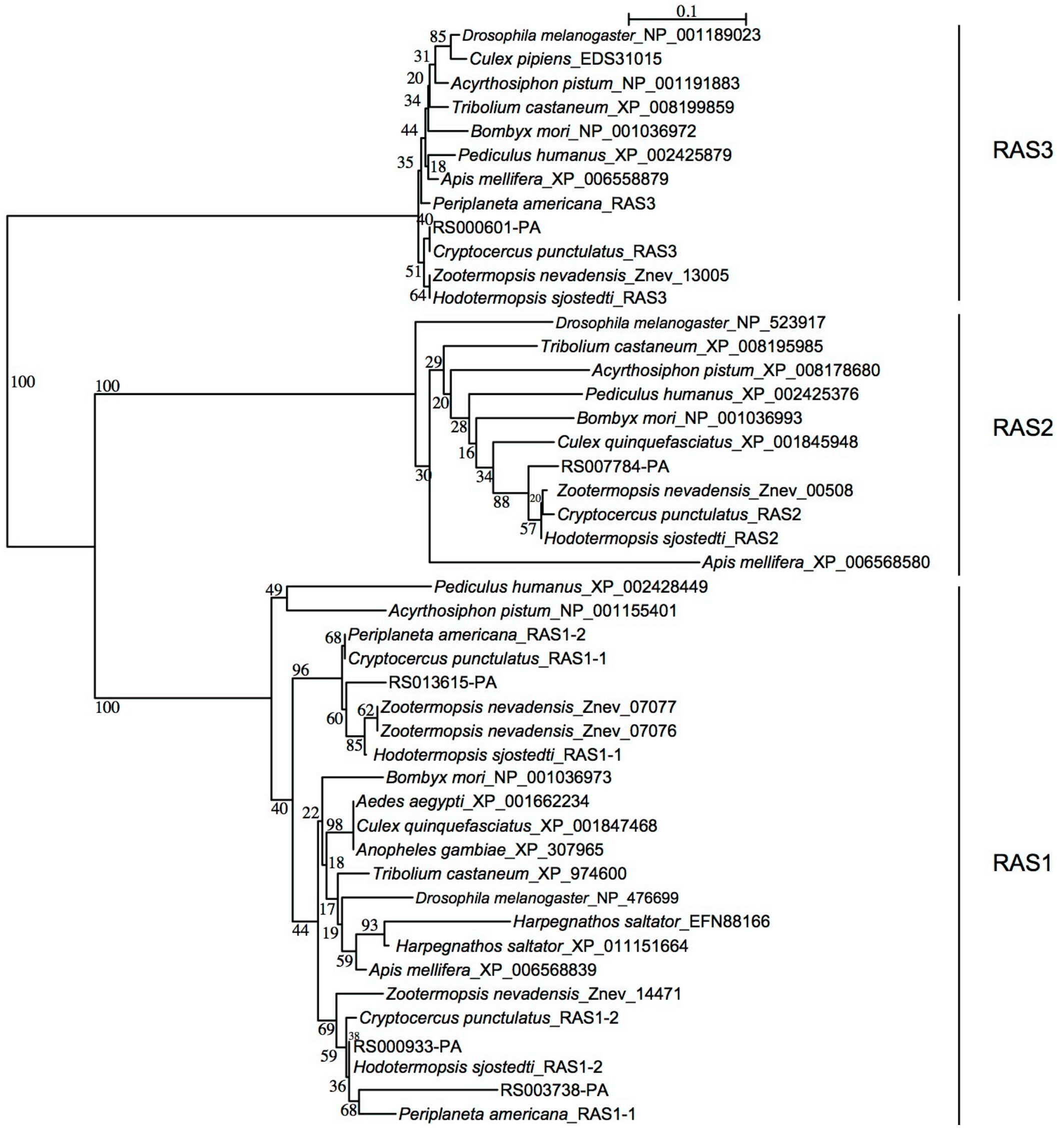


**Fig. S29.** Neighbor-joining (NJ) tree of Ras85D (Ras1) homologs based on the amino acid sequences obtained with Poisson model. The bootstrap percentages of 1000 NJ trees in which the associated taxa clustered together are shown next to the nodes. The Ras1 homologs of *R. speratus* are *RS013615*, *RS000933* and *RS003738*. The Ras64B (Ras2) and Rap1 (Ras3) homologs of *R. speratus* are *RS007784* and *RS000601*, respectively. Amino acid sequences of *Periplaneta americana* and *Cryptocercus punctulatus* are obtained from the assembled contig sequences using Trinity (Grabherr et al. 2011) (transcriptome data: DRA001254 and DRA004598, respectively).

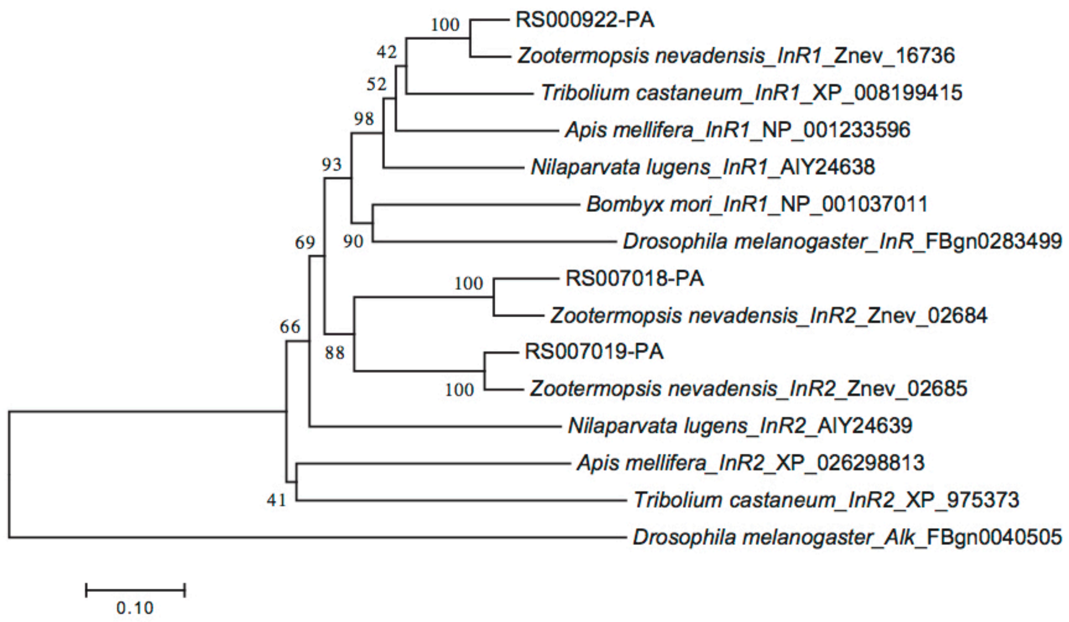


**Fig. S30.** Neighbor-joining (NJ) tree of insulin receptor (InR) homologs based on the amino acid sequences obtained with Poisson model. The bootstrap percentages of 1000 NJ trees in which the associated taxa clustered together are shown next to the nodes. Nucleotide sequences exept for *Reticulitermes speratus* were referred to Xu and Zhang (2015). The InR homologs of *R. speratus* are *RS000922*, *RS007018* and *RS007019*.

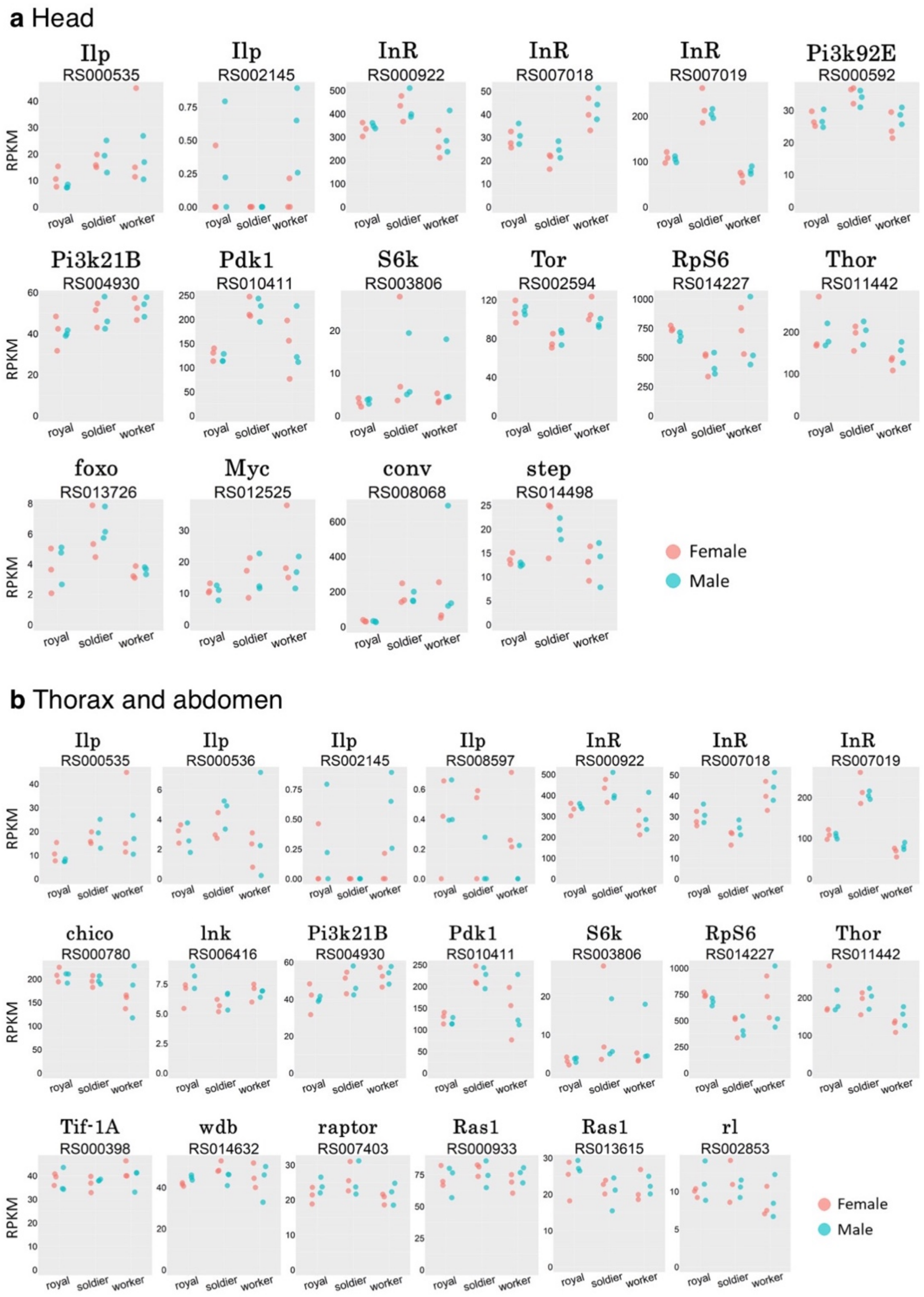


**Fig. S31.** Expression levels of Insulin/insulin-like signaling pathway genes among royals (reproductives), soldiers and workers in *Reticulitermes speratus*. Expression levels are indicated as RPKM calculated from RNA-sequencing analysis using the head (a) and the thorax and abdomen (b) samples. Orange and blue points indicate females and males, respectively.

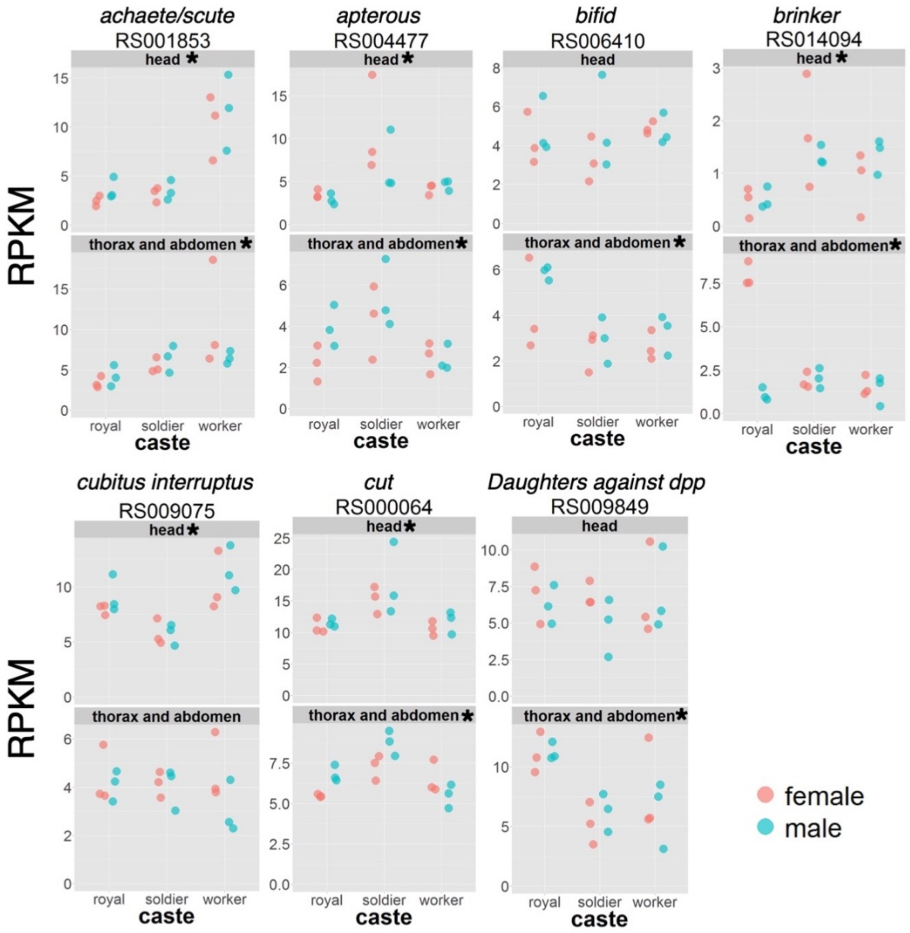


**Fig. S32.** Expression levels of toolkit genes among royals (reproductives), soldiers and workers in *Reticulitermes speratus*. Expression levels are indicated as RPKM calculated from RNA-sequencing analysis. Orange and blue points indicate females and males, respectively. All these 19 genes show the significant differences among castes in heads and/or thorax and abdomen samples (*FDR < 0.05).

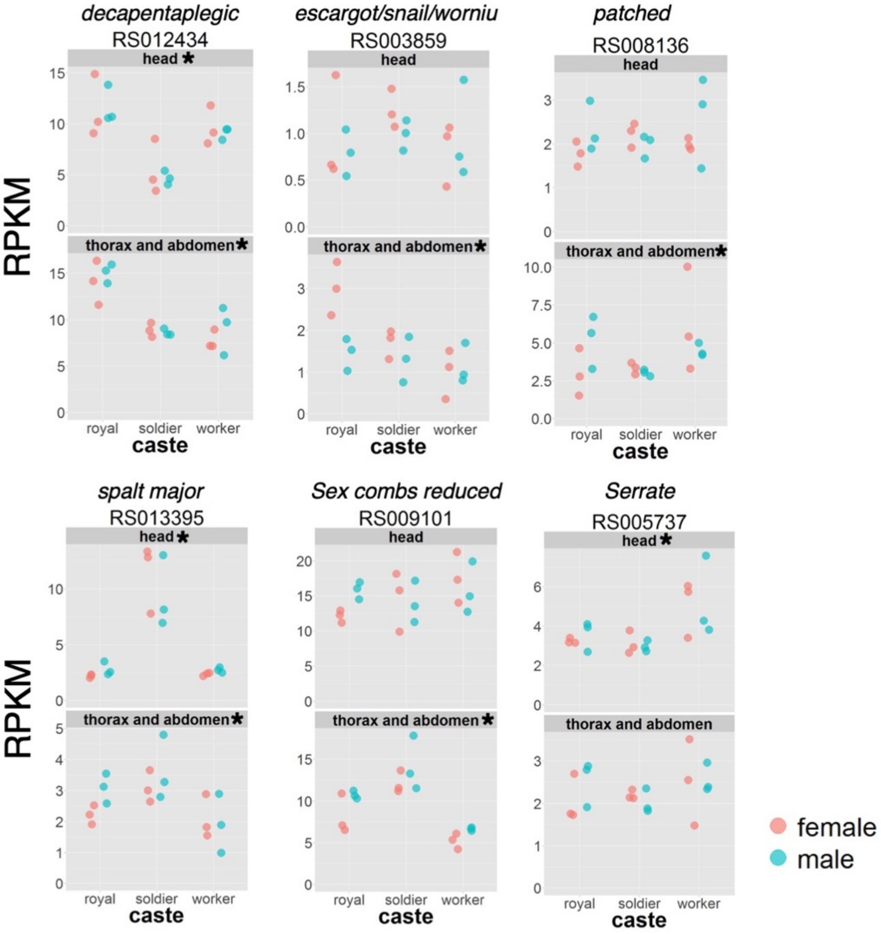


**Fig. S32.** *Continued.*

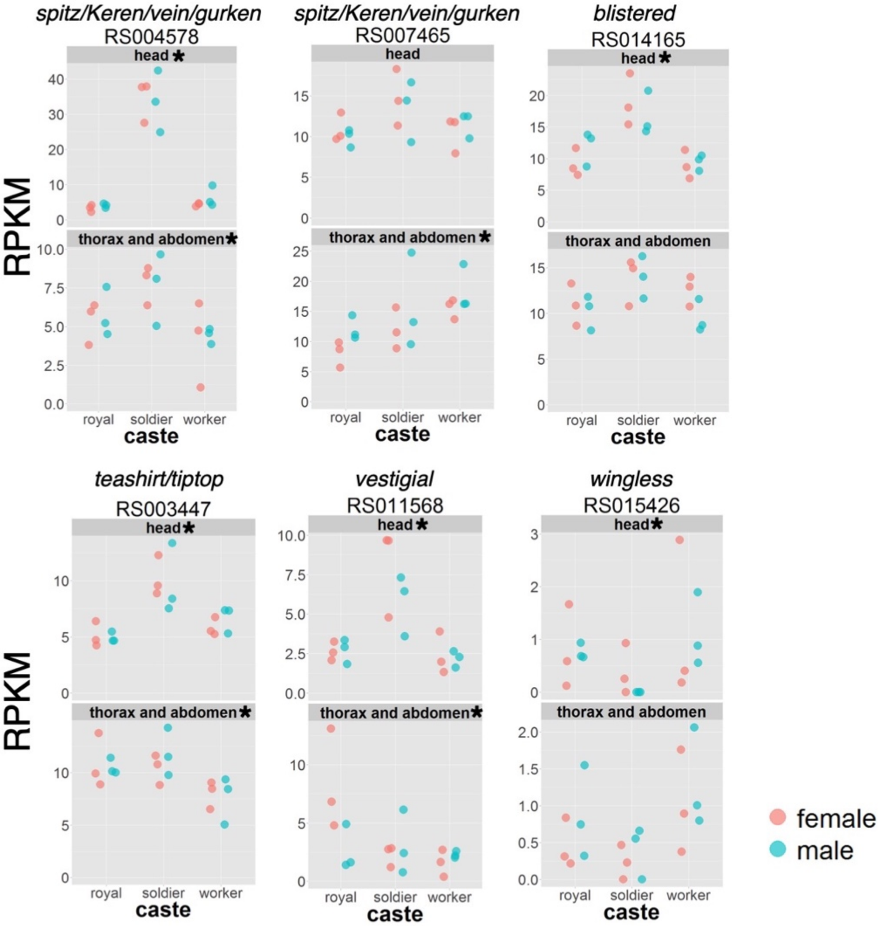


**Fig. S32.** *Continued*.

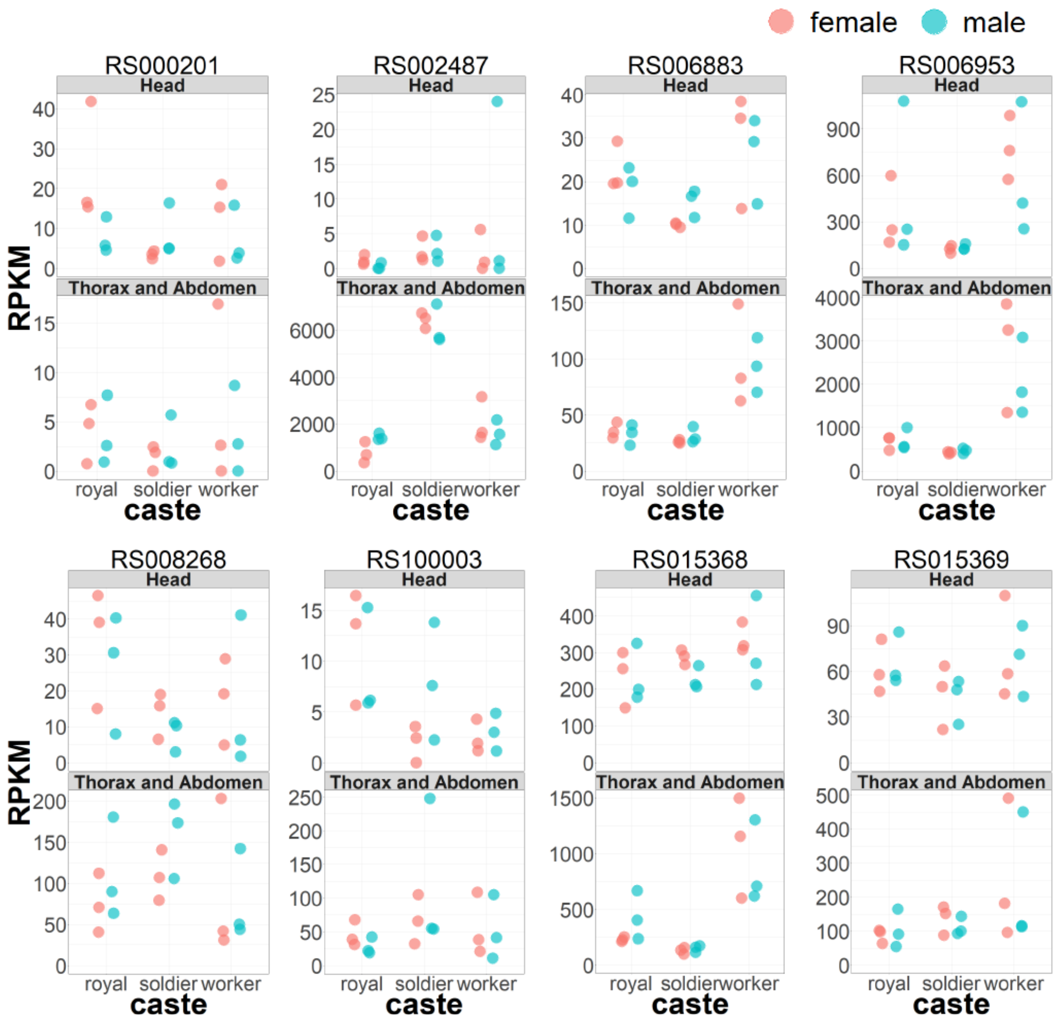


**Fig. S33.** Gene expression levels of antimicrobial peptides among royals (reproductives), soldiers and workers in *Reticulitermes speratus*. Expression levels are indicated as RPKM calculated from RNA-sequencing analysis. Orange and blue points indicate females and males, respectively.

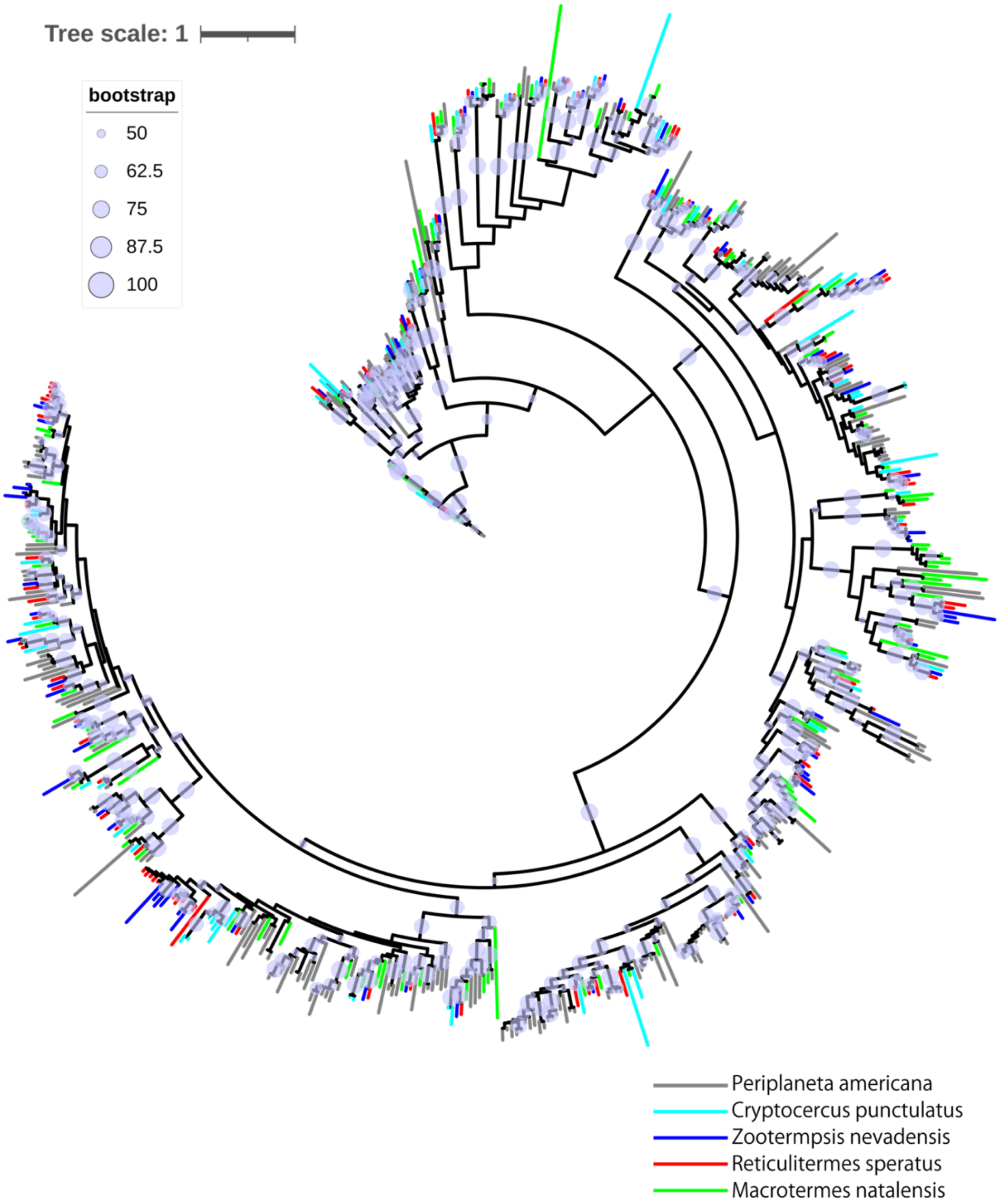


**Fig. S34.** Maximum likelihood (ML) tree of cytochrome P450 monooxygenases (CYP) homologs based on the amino acid sequences obtained with LG+I+G model. The bootstrap percentages of 100 ML trees in which the associated taxa clustered together are marked as circles on the nodes. Sequences from 5 species of cockroaches and termites are indicated as different color branches. Gene IDs of termites are shown in Supplementary Table 28. Amino acid sequences of *Periplaneta americana* and *Cryptocercus punctulatus* are obtained from the assembled contig sequences using Trinity (Grabherr et al. 2011) (transcriptome data: DRA001254 and DRA004598, respectively).

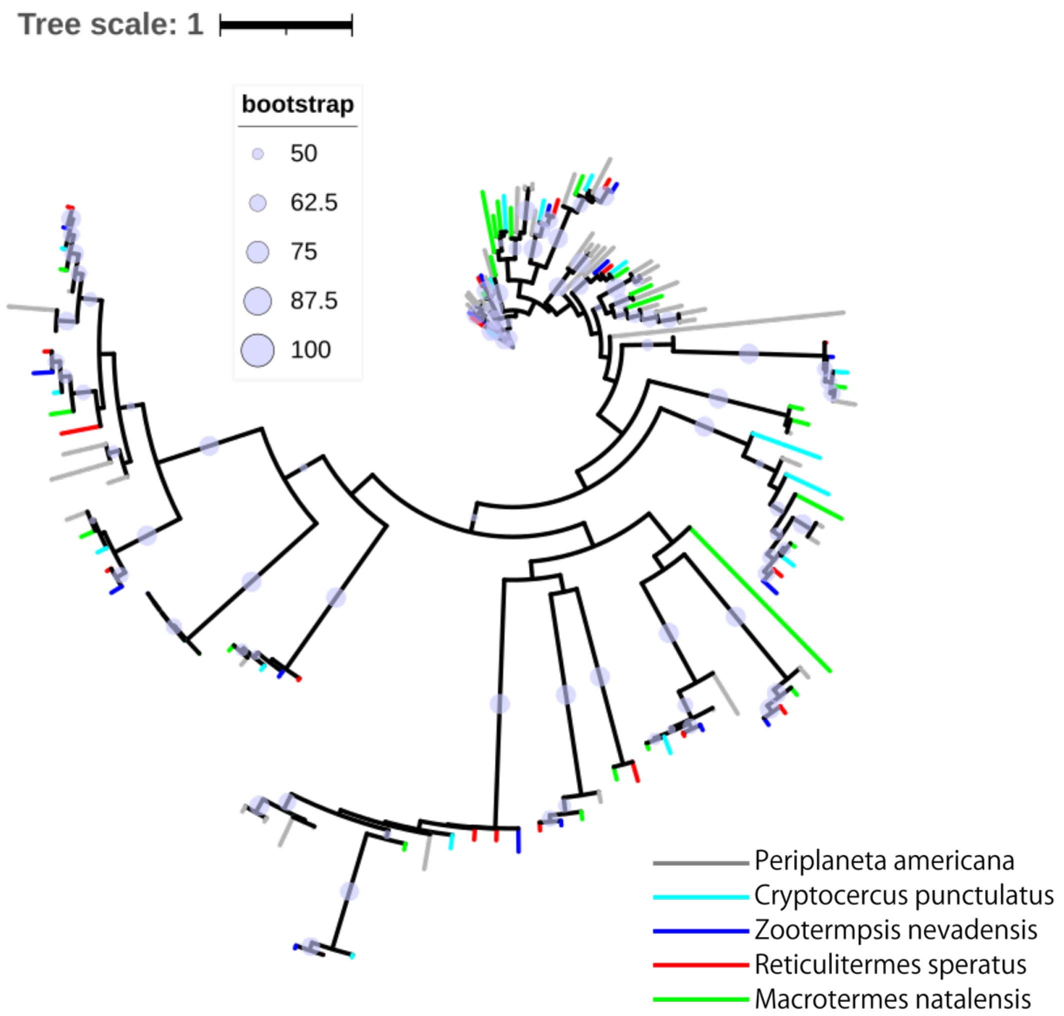


**Fig. S35.** Maximum likelihood (ML) tree of glutathione S-transferases (GST) homologs based on the amino acid sequences obtained with JTT+I+G model. The bootstrap percentages of 100 ML trees in which the associated taxa clustered together are marked as circles on the nodes. Sequences from 5 species of cockroaches and termites are indicated as different color branches. Gene IDs of termites are shown in Supplementary Table 28. Amino acid sequences of *Periplaneta americana* and *Cryptocercus punctulatus* are obtained from the assembled contig sequences using Trinity (Grabherr et al. 2011) (transcriptome data: DRA001254 and DRA004598, respectively).

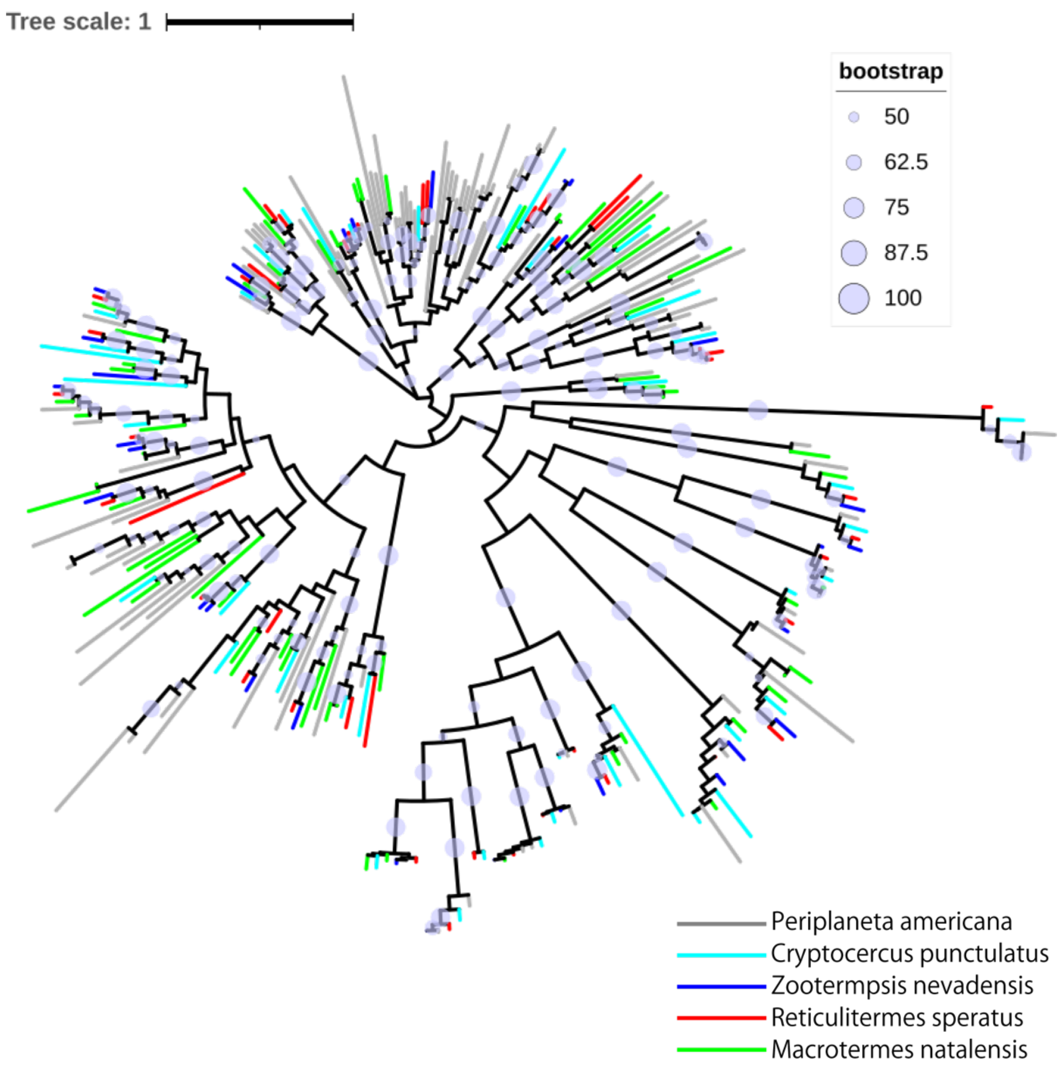


**Fig. S36.** Maximum likelihood (ML) tree of carboxylesterases (CCE) homologs based on the amino acid sequences obtained with JTT+I+G model. The bootstrap percentages of 100 ML trees in which the associated taxa clustered together are marked as circles on the nodes. Sequences from 5 species of cockroaches and termites are indicated as different color branches. Gene IDs of termites are shown in Supplementary Table 28. Amino acid sequences of *Periplaneta americana* and *Cryptocercus punctulatus* are obtained from the assembled contig sequences using Trinity (Grabherr et al. 2011) (transcriptome data: DRA001254 and DRA004598, respectively).

**Table S1.** Termite colonies used for next-generation sequencing.

| Analysis | Colony no | Location collected | Date collected | Castes |
|---|---|---|---|---|
| genome | #1 | Furudo, Toyama | November, 2013 | female secondary reprpductives (nymphoids) |
| RNAseq | #2 | Furudo, Toyama | September, 2014 | worker and soldier |
| RNAseq | #3 | Furudo, Toyama | September, 2014 | worker and soldier |
| RNAseq | #4 | Furudo, Toyama | September, 2014 | worker and soldier |
| RNAseq | #5 | Furudo, Toyama | April-May, 2014 | reproductive |
| RNAseq | #6 | Furudo, Toyama | April-May, 2014 | reproductive |
| RNAseq | #7 | Furudo, Toyama | April-May, 2014 | reproductive |
| PBAT | #8 | Furudo, Toyama | October, 2014 | worker and soldier |
| PBAT | #5, #7 | Furudo, Toyama | May, 2014 | reproductive |

**Table S2.** *Reticulitermes speratus* RNA sequencing libraries.

| Library name | SRA ID | Sample description | Caste | Sex | Body part | Colony* |
|---|---|---|---|---|---|---|
| Rspe_RMH1 | DRR090830 | *Reticulitermes speratus*, royal, male, head, rep1 | reproductive | male | head | #5 |
| Rspe_SFTA3 | DRR090831 | *Reticulitermes speratus*, soldier, female, thorax and abdomen, rep3 | soldier | female | thorax and abdomen | #4 |
| Rspe_RMTA1 | DRR090832 | *Reticulitermes speratus*, royal, male, thorax and abdomen, rep1 | reproductive | male | thorax and abdomen | #5 |
| Rspe_RMH3 | DRR090833 | *Reticulitermes speratus*, royal, male, head, rep3 | reproductive | male | head | #6 |
| Rspe_RFH1 | DRR090834 | *Reticulitermes speratus*, royal, female, head, rep1 | reproductive | female | head | #6 |
| Rspe_RMTA3 | DRR090835 | *Reticulitermes speratus*, royal, male, thorax and abdomen, rep3 | reproductive | male | thorax and abdomen | #6 |
| Rspe_RFTA1 | DRR090836 | *Reticulitermes speratus*, royal, female, thorax and abdomen, rep1 | reproductive | female | thorax and abdomen | #6 |
| Rspe_RFH3 | DRR090837 | *Reticulitermes speratus*, royal, female, head, rep3 | reproductive | female | head | #7 |
| Rspe_WMH2 | DRR090838 | *Reticulitermes speratus*, worker, male, head, rep2 | worker | male | head | #3 |
| Rspe_RFTA3 | DRR090839 | *Reticulitermes speratus*, royal, female, thorax and abdomen, rep3 | reproductive | female | thorax and abdomen | #7 |
| Rspe_WMTA2 | DRR090840 | *Reticulitermes speratus*, worker, male, thorax and abdomen, rep2 | worker | male | thorax and abdomen | #3 |
| Rspe_WFH2 | DRR090841 | *Reticulitermes speratus*, worker, female, head, rep2 | worker | female | head | #3 |
| Rspe_WFTA2 | DRR090842 | *Reticulitermes speratus*, worker, female, thorax and abdomen, rep2 | worker | female | thorax and abdomen | #3 |
| Rspe_SMH2 | DRR090843 | *Reticulitermes speratus*, soldier, male, head, rep2 | soldier | male | head | #3 |
| Rspe_WMH1 | DRR090844 | *Reticulitermes speratus*, worker, male, head, rep1 | worker | male | head | #2 |
| Rspe_RFTA2 | DRR090845 | *Reticulitermes speratus*, royal, female, thorax and abdomen, rep2 | reproductive | female | thorax and abdomen | #7 |
| Rspe_SMTA2 | DRR090846 | *Reticulitermes speratus*, soldier, male, thorax and abdomen, rep2 | soldier | male | thorax and abdomen | #3 |
| Rspe_SFH2 | DRR090847 | *Reticulitermes speratus*, soldier, female, head, rep2 | soldier | female | head | #3 |
| Rspe_SFTA2 | DRR090848 | *Reticulitermes speratus*, soldier, female, thorax and abdomen, rep2 | soldier | female | thorax and abdomen | #3 |
| Rspe_RMH2 | DRR090849 | *Reticulitermes speratus*, royal, male, head, rep2 | reproductive | male | head | #5 |
| Rspe_RMTA2 | DRR090850 | *Reticulitermes speratus*, royal, male, thorax and abdomen, rep2 | reproductive | male | thorax and abdomen | #5 |
| Rspe_RFH2 | DRR090851 | *Reticulitermes speratus*, royal, female, head, rep2 | reproductive | female | head | #7 |
| Rspe_WMTA1 | DRR090852 | *Reticulitermes speratus*, worker, male, thorax and abdomen, rep1 | worker | male | thorax and abdomen | #2 |
| Rspe_WMH3 | DRR090853 | *Reticulitermes speratus*, worker, male, head, rep3 | worker | male | head | #4 |
| Rspe_WFH1 | DRR090854 | *Reticulitermes speratus*, worker, female, head, rep1 | worker | female | head | #2 |
| Rspe_WMTA3 | DRR090855 | *Reticulitermes speratus*, worker, male, thorax and abdomen, rep3 | worker | male | thorax and abdomen | #4 |
| Rspe_WFTA1 | DRR090856 | *Reticulitermes speratus*, worker, female, thorax and abdomen, rep1 | worker | female | thorax and abdomen | #2 |

| | | | | | | |
|---|---|---|---|---|---|---|
| Rspe_WFH3 | DRR090857 | *Reticulitermes speratus*, worker, female, head, rep3 | worker | female | head | #4 |
| Rspe_SMH1 | DRR090858 | *Reticulitermes speratus*, soldier, male, head, rep1 | soldier | male | head | #2 |
| Rspe_WFTA3 | DRR090859 | *Reticulitermes speratus*, worker, female, thorax and abdomen, rep3 | worker | female | thorax and abdomen | #4 |
| Rspe_SMTA1 | DRR090860 | *Reticulitermes speratus*, soldier, male, thorax and abdomen, rep1 | soldier | male | thorax and abdomen | #2 |
| Rspe_SMH3 | DRR090861 | *Reticulitermes speratus*, soldier, male, head, rep3 | soldier | male | head | #4 |
| Rspe_SFH1 | DRR090862 | *Reticulitermes speratus*, soldier, female, head, rep1 | soldier | female | head | #2 |
| Rspe_SMTA3 | DRR090863 | *Reticulitermes speratus*, soldier, male, thorax and abdomen, rep3 | soldier | male | thorax and abdomen | #4 |
| Rspe_SFTA1 | DRR090864 | *Reticulitermes speratus*, soldier, female, thorax and abdomen, rep1 | soldier | female | thorax and abdomen | #2 |
| Rspe_SFH3 | DRR090865 | *Reticulitermes speratus*, soldier, female, head, rep3 | soldier | female | head | #4 |

*corresponds to the colony # in Supplementary Table 1.

**Table S3.** *Reticulitermes speratus* Illumina libraries for genome sequencing and whole-genome bisulfite sequencing.

| Library name | SRA ID | Sample description | Caste | Sex | Body part | Colony* |
|---|---|---|---|---|---|---|
| **Genome** | | | | | | |
| Rspe_GPE250 | DRR000000 | Paired-end 250 bp insert | female secondary reprpductives (nymphoids) | female | whole body; gut and ovaries are excluded | #1 |
| Rspe_GPE800 | DRR000000 | Paired-end 800 bp insert | female secondary reprpductives (nymphoids) | female | whole body; gut and ovaries are excluded | #1 |
| Rspe_GMP3k | DRR252502 | Mate-pair 3k bp insert | female secondary reprpductives (nymphoids) | female | whole body; gut and ovaries are excluded | #1 |
| Rspe_GMP5k | DRR252503 | Mate-pair 5k bp insert | female secondary reprpductives (nymphoids) | female | whole body; gut and ovaries are excluded | #1 |
| Rspe_GMP8k | DRR252504 | Mate-pair 8k bp insert | female secondary reprpductives (nymphoids) | female | whole body; gut and ovaries are excluded | #1 |
| Rspe_GMP10k | DRR252505 | Mate-pair 10k bp insert | female secondary reprpductives (nymphoids) | female | whole body; gut and ovaries are excluded | #1 |
| **Methylome** | | | | | | |
| Rspe_PBAT | DRR000000 | PBAT Primary reproductive, Male, Head | reproductive | male | head | #5 |
| Rspe_PBAT | DRR000000 | PBAT Primary reproductive, Female, Head | reproductive | female | head | #7 |
| Rspe_PBAT | DRR000000 | PBAT Soldier, Male, Head | soldier | male | head | #8 |
| Rspe_PBAT | DRR000000 | PBAT Soldier, Female, Head | soldier | female | head | #8 |
| Rspe_PBAT | DRR000000 | PBAT Worker, Male, Head | worker | male | head | #8 |
| Rspe_PBAT | DRR000000 | PBAT Worker Female, Head | worker | female | head | #8 |

* corresponds to the colony # in Supplementary Table 1.

**Table S4.** Over-represented Gene Ontology terms in caste-DEG.

| body part | category | Pfam domain | % in caste-biased genes | % in all | q-value | motif description |
|---|---|---|---|---|---|---|
| **Thorax + abdomen** | | | | | | |
| | BP | GO:0000270 | 0.34% | 0.08% | 0.02242181 | peptidoglycan metabolic process |
| | | GO:0005975 | 3.85% | 2.54% | 0.009769257 | carbohydrate metabolic process |
| | | GO:0006022 | 1.69% | 0.80% | 0.003451719 | aminoglycan metabolic process |
| | | GO:0006027 | 0.53% | 0.15% | 0.009769257 | glycosaminoglycan catabolic process |
| | | GO:0006030 | 1.16% | 0.56% | 0.034011861 | chitin metabolic process |
| | | GO:0006040 | 1.35% | 0.71% | 0.040302871 | amino sugar metabolic process |
| | | GO:0006629 | 4.43% | 2.84% | 0.002648555 | lipid metabolic process |
| | | GO:0006633 | 1.01% | 0.44% | 0.017959474 | fatty acid biosynthetic process |
| | | GO:0006720 | 1.20% | 0.44% | 0.00066074 | isoprenoid metabolic process |
| | | GO:0006721 | 0.87% | 0.22% | 4.46E-05 | terpenoid metabolic process |
| | | GO:0006726 | 0.39% | 0.08% | 0.004304289 | eye pigment biosynthetic process |
| | | GO:0008299 | 0.92% | 0.29% | 0.00066074 | isoprenoid biosynthetic process |
| | | GO:0008610 | 2.17% | 1.23% | 0.009769257 | lipid biosynthetic process |
| | | GO:0009253 | 0.34% | 0.06% | 0.004446678 | peptidoglycan catabolic process |
| | | GO:0016063 | 0.29% | 0.05% | 0.012532405 | rhodopsin biosynthetic process |
| | | GO:0016108 | 0.24% | 0.05% | 0.047939137 | tetraterpenoid metabolic process |
| | | GO:0016114 | 0.58% | 0.16% | 0.004446678 | terpenoid biosynthetic process |
| | | GO:0016116 | 0.24% | 0.05% | 0.047939137 | carotenoid metabolic process |
| | | GO:0042441 | 0.39% | 0.08% | 0.007327479 | eye pigment metabolic process |
| | | GO:0043052 | 0.34% | 0.05% | 0.001447822 | thermotaxis |
| | | GO:0043324 | 0.39% | 0.08% | 0.007327479 | pigment metabolic process involved in developmental pigmentation |
| | | GO:0043474 | 0.39% | 0.08% | 0.007327479 | pigment metabolic process involved in pigmentation |
| | | GO:0044255 | 3.56% | 2.40% | 0.026230783 | cellular lipid metabolic process |
| | | GO:0046154 | 0.29% | 0.06% | 0.034011861 | rhodopsin metabolic process |
| | | GO:0048069 | 0.39% | 0.08% | 0.007327479 | eye pigmentation |
| | | GO:1901136 | 1.11% | 0.49% | 0.010846089 | carbohydrate derivative catabolic process |
| | MF | GO:0003796 | 0.48% | 0.11% | 0.000303785 | lysozyme activity |
| | | GO:0004175 | 3.37% | 1.82% | 1.04E-05 | endopeptidase activity |
| | | GO:0004252 | 1.97% | 0.86% | 1.04E-05 | serine-type endopeptidase activity |
| | | GO:0004311 | 0.29% | 0.07% | 0.019075313 | farnesyltranstransferase activity |
| | | GO:0004312 | 0.39% | 0.09% | 0.003311523 | fatty acid synthase activity |

| | | | | | | |
|---|---|---|---|---|---|---|
| | | GO:0004497 | 1.64% | 0.84% | 0.002474851 | monooxygenase activity |
| | | GO:0004553 | 2.17% | 0.88% | 1.12E-06 | hydrolase activity, hydrolyzing O-glycosyl compounds |
| | | GO:0004659 | 0.48% | 0.18% | 0.041592279 | prenyltransferase activity |
| | | GO:0004806 | 0.19% | 0.04% | 0.04711786 | triglyceride lipase activity |
| | | GO:0004871 | 3.81% | 2.77% | 0.034606867 | signal transducer activity |
| | | GO:0004872 | 4.09% | 2.83% | 0.005844382 | receptor activity |
| | | GO:0004888 | 3.32% | 2.15% | 0.003311523 | transmembrane signaling receptor activity |
| | | GO:0004930 | 2.07% | 1.29% | 0.020926341 | G-protein coupled receptor activity |
| | | GO:0005214 | 0.39% | 0.13% | 0.044135237 | structural constituent of chitin-based cuticle |
| | | GO:0005506 | 2.17% | 1.15% | 0.000633723 | iron ion binding |
| | | GO:0008061 | 0.92% | 0.46% | 0.037508132 | chitin binding |
| | | GO:0008194 | 0.87% | 0.44% | 0.045884796 | UDP-glycosyltransferase activity |
| | | GO:0008233 | 4.91% | 3.20% | 0.000226142 | peptidase activity |
| | | GO:0008236 | 2.31% | 1.02% | 1.67E-06 | serine-type peptidase activity |
| | | GO:0008237 | 1.59% | 0.95% | 0.033595482 | metallopeptidase activity |
| | | GO:0008422 | 0.34% | 0.08% | 0.005252854 | beta-glucosidase activity |
| | | GO:0008745 | 0.34% | 0.06% | 0.000750055 | N-acetylmuramoyl-L-alanine amidase activity |
| | | GO:0015020 | 0.43% | 0.16% | 0.047143438 | glucuronosyltransferase activity |
| | | GO:0015926 | 0.39% | 0.13% | 0.044135237 | glucosidase activity |
| | | GO:0016160 | 0.19% | 0.04% | 0.04711786 | amylase activity |
| | | GO:0016297 | 0.29% | 0.06% | 0.007854275 | acyl-[acyl-carrier-protein] hydrolase activity |
| | | GO:0016614 | 1.35% | 0.66% | 0.003419662 | oxidoreductase activity, acting on CH-OH group of donors |
| | | GO:0016620 | 0.87% | 0.33% | 0.002073942 | oxidoreductase activity, acting on the aldehyde or oxo group of donors, NAD or NADP as acceptor |
| | | GO:0016705 | 2.31% | 1.17% | 0.000109621 | oxidoreductase activity, acting on paired donors, with incorporation or reduction of molecular oxygen |
| | | GO:0016717 | 0.39% | 0.09% | 0.003311523 | oxidoreductase activity, acting on paired donors, with oxidation of a pair of donors resulting in the reduction of molecular oxygen to two molecules of water |
| | | GO:0016798 | 2.36% | 1.03% | 1.67E-06 | hydrolase activity, acting on glycosyl bonds |
| | | GO:0016903 | 0.92% | 0.40% | 0.006585745 | oxidoreductase activity, acting on the aldehyde or oxo group of donors |
| | | GO:0017171 | 2.31% | 1.02% | 1.67E-06 | serine hydrolase activity |
| | | GO:0020037 | 2.12% | 1.14% | 0.000941207 | heme binding |
| | | GO:0038023 | 3.47% | 2.36% | 0.009272631 | signaling receptor activity |
| | | GO:0042302 | 1.30% | 0.44% | 3.78E-06 | structural constituent of cuticle |

| | | | | | | |
|---|---|---|---|---|---|---|
| | | GO:0046906 | 2.17% | 1.16% | 0.000669074 | tetrapyrrole binding |
| | | GO:0060089 | 4.09% | 2.83% | 0.005844382 | molecular transducer activity |
| | | GO:0070011 | 4.72% | 2.97% | 9.27E-05 | peptidase activity, acting on L-amino acid peptides |
| | | GO:0080019 | 0.72% | 0.23% | 0.000633723 | fatty-acyl-CoA reductase (alcohol-forming) activity |
| | | GO:0099600 | 3.61% | 2.38% | 0.003311523 | transmembrane receptor activity |
| | CC | GO:0005576 | 4.91% | 2.72% | 2.48E-07 | extracellular region |
| **Head** | | | | | | |
| | BP | GO:0006022 | 2.03% | 0.80% | 0.000141493 | aminoglycan metabolic process |
| | | GO:0006030 | 1.71% | 0.56% | 2.18E-05 | chitin metabolic process |
| | | GO:0006040 | 1.84% | 0.71% | 0.000189878 | amino sugar metabolic process |
| | | GO:0006629 | 5.19% | 2.84% | 1.55E-05 | lipid metabolic process |
| | | GO:0006694 | 0.82% | 0.27% | 0.021334218 | steroid biosynthetic process |
| | | GO:0006714 | 0.38% | 0.07% | 0.021334218 | sesquiterpenoid metabolic process |
| | | GO:0006716 | 0.38% | 0.07% | 0.021334218 | juvenile hormone metabolic process |
| | | GO:0006718 | 0.38% | 0.07% | 0.021334218 | juvenile hormone biosynthetic process |
| | | GO:0006720 | 1.77% | 0.44% | 1.03E-08 | isoprenoid metabolic process |
| | | GO:0006721 | 1.27% | 0.22% | 2.48E-09 | terpenoid metabolic process |
| | | GO:0006816 | 0.63% | 0.17% | 0.021334218 | calcium ion transport |
| | | GO:0008202 | 0.89% | 0.32% | 0.025400444 | steroid metabolic process |
| | | GO:0008299 | 1.39% | 0.29% | 1.10E-08 | isoprenoid biosynthetic process |
| | | GO:0008610 | 2.66% | 1.23% | 0.000189878 | lipid biosynthetic process |
| | | GO:0016106 | 0.38% | 0.07% | 0.021334218 | sesquiterpenoid biosynthetic process |
| | | GO:0016114 | 0.95% | 0.16% | 2.16E-07 | terpenoid biosynthetic process |
| | | GO:0034754 | 0.57% | 0.14% | 0.021334218 | cellular hormone metabolic process |
| | | GO:0040003 | 0.76% | 0.24% | 0.021334218 | chitin-based cuticle development |
| | | GO:0042335 | 0.89% | 0.30% | 0.021334218 | cuticle development |
| | | GO:0042445 | 0.70% | 0.19% | 0.012512124 | hormone metabolic process |
| | | GO:0044255 | 4.31% | 2.40% | 0.000209167 | cellular lipid metabolic process |
| | | GO:0070588 | 0.57% | 0.15% | 0.025400444 | calcium ion transmembrane transport |
| | | GO:1901071 | 1.71% | 0.59% | 6.45E-05 | glucosamine-containing compound metabolic process |
| | MF | GO:0004161 | 0.38% | 0.05% | 0.000793969 | dimethylallyltranstransferase activity |
| | | GO:0004311 | 0.38% | 0.07% | 0.005865833 | farnesyltranstransferase activity |
| | | GO:0004497 | 2.98% | 0.84% | 9.46E-14 | monooxygenase activity |
| | | GO:0004553 | 2.22% | 0.88% | 1.32E-05 | hydrolase activity, hydrolyzing O-glycosyl compounds |
| | | GO:0004659 | 0.70% | 0.18% | 0.001470475 | prenyltransferase activity |

| | | | | | | |
|---|---|---|---|---|---|---|
| | | GO:0005198 | 3.74% | 2.14% | 0.000580658 | structural molecule activity |
| | | GO:0005201 | 0.44% | 0.08% | 0.001216335 | extracellular matrix structural constituent |
| | | GO:0005214 | 0.63% | 0.13% | 0.000267327 | structural constituent of chitin-based cuticle |
| | | GO:0005216 | 2.28% | 1.41% | 0.047855518 | ion channel activity |
| | | GO:0005262 | 0.51% | 0.12% | 0.006835974 | calcium channel activity |
| | | GO:0005319 | 0.63% | 0.20% | 0.018197038 | lipid transporter activity |
| | | GO:0005506 | 3.86% | 1.15% | 2.29E-16 | iron ion binding |
| | | GO:0008010 | 0.38% | 0.08% | 0.019573356 | structural constituent of chitin-based larval cuticle |
| | | GO:0008061 | 1.33% | 0.46% | 0.000213188 | chitin binding |
| | | GO:0008083 | 0.57% | 0.19% | 0.038334846 | growth factor activity |
| | | GO:0008422 | 0.38% | 0.08% | 0.011539152 | beta-glucosidase activity |
| | | GO:0015085 | 0.63% | 0.16% | 0.002009273 | calcium ion transmembrane transporter activity |
| | | GO:0015171 | 0.63% | 0.23% | 0.047855518 | amino acid transmembrane transporter activity |
| | | GO:0015248 | 0.25% | 0.05% | 0.047855518 | sterol transporter activity |
| | | GO:0015926 | 0.57% | 0.13% | 0.001746016 | glucosidase activity |
| | | GO:0016614 | 1.84% | 0.66% | 1.38E-05 | oxidoreductase activity, acting on CH-OH group of donors |
| | | GO:0016620 | 1.14% | 0.33% | 7.89E-05 | oxidoreductase activity, acting on the aldehyde or oxo group of donors, NAD or NADP as acceptor |
| | | GO:0016705 | 3.61% | 1.17% | 1.19E-13 | oxidoreductase activity, acting on paired donors, with incorporation or reduction of molecular oxygen |
| | | GO:0016709 | 0.32% | 0.07% | 0.044419513 | oxidoreductase activity, acting on paired donors, with incorporation or reduction of molecular oxygen, NAD(P)H as one donor, and incorporation of one atom of oxygen |
| | | GO:0016765 | 0.95% | 0.37% | 0.011539152 | transferase activity, transferring alkyl or aryl (other than methyl) groups |
| | | GO:0016798 | 2.28% | 1.03% | 0.00017759 | hydrolase activity, acting on glycosyl bonds |
| | | GO:0016903 | 1.20% | 0.40% | 0.00026769 | oxidoreductase activity, acting on the aldehyde or oxo group of donors |
| | | GO:0020037 | 3.80% | 1.14% | 4.06E-16 | heme binding |
| | | GO:0022891 | 5.07% | 3.53% | 0.014299106 | substrate-specific transmembrane transporter activity |
| | | GO:0042302 | 2.41% | 0.44% | 5.57E-19 | structural constituent of cuticle |
| | | GO:0042813 | 0.25% | 0.05% | 0.047855518 | Wnt-activated receptor activity |
| | | GO:0046906 | 3.86% | 1.16% | 2.29E-16 | tetrapyrrole binding |
| | | GO:0046943 | 0.76% | 0.31% | 0.047855518 | carboxylic acid transmembrane transporter activity |
| | | GO:0048037 | 2.34% | 1.41% | 0.029166002 | cofactor binding |
| | | GO:0050660 | 1.27% | 0.57% | 0.012190946 | flavin adenine dinucleotide binding |

| | | | | | | |
|---|---|---|---|---|---|---|
| | | GO:0072509 | 0.63% | 0.23% | 0.040436179 | divalent inorganic cation transmembrane transporter activity |
| | | GO:0080019 | 0.89% | 0.23% | 0.000173742 | fatty-acyl-CoA reductase (alcohol-forming) activity |
| | CC | GO:0005576 | 5.51% | 2.72% | 2.39E-08 | extracellular region |
| | | GO:0005578 | 0.89% | 0.28% | 0.007096274 | proteinaceous extracellular matrix |
| | | GO:0031012 | 1.46% | 0.45% | 3.09E-05 | extracellular matrix |

**Table S5.** Over-represented Pfam domains in caste-DEG.

| body part | Pfam domain | % in caste-biased genes | % in all | q-value | motif description |
|---|---|---|---|---|---|
| **Thorax + abdomen** | | | | | |
| | PF00019 | 0.29% | 0.07% | 0.018855562 | Transforming growth factor beta like domain |
| | PF00049 | 0.19% | 0.04% | 0.047330812 | Insulin/IGF/Relaxin family |
| | PF00059 | 0.58% | 0.16% | 0.00082408 | Lectin C-type domain |
| | PF00061 | 0.29% | 0.06% | 0.010173654 | Lipocalin / cytosolic fatty-acid binding protein family |
| | PF00062 | 0.48% | 0.10% | 0.000162778 | C-type lysozyme/alpha-lactalbumin family |
| | PF00067 | 1.49% | 0.74% | 0.003042693 | Cytochrome P450 |
| | PF00084 | 0.39% | 0.11% | 0.018855562 | Sushi repeat (SCR repeat) |
| | PF00089 | 1.73% | 0.61% | 4.06E-07 | Trypsin |
| | PF00151 | 0.39% | 0.12% | 0.02933154 | Lipase |
| | PF00201 | 0.72% | 0.22% | 0.000469997 | UDP-glucoronosyl and UDP-glucosyl transferase |
| | PF00232 | 0.63% | 0.12% | 2.96E-06 | Glycosyl hydrolase family 1 |
| | PF00282 | 0.29% | 0.08% | 0.033068556 | Pyridoxal-dependent decarboxylase conserved domain |
| | PF00348 | 0.48% | 0.13% | 0.002788466 | Polyprenyl synthetase |
| | PF00379 | 1.20% | 0.38% | 2.96E-06 | Insect cuticle protein |
| | PF00394 | 0.24% | 0.05% | 0.02933154 | Multicopper oxidase |
| | PF00560 | 0.53% | 0.17% | 0.010173654 | Leucine Rich Repeat |
| | PF00650 | 0.77% | 0.25% | 0.000611959 | CRAL/TRIO domain |
| | PF00688 | 0.29% | 0.07% | 0.018855562 | TGF-beta propeptide |
| | PF01061 | 0.43% | 0.14% | 0.018249708 | ABC-2 type transporter |
| | PF01151 | 0.34% | 0.11% | 0.049368426 | GNS1/SUR4 family |
| | PF01400 | 0.29% | 0.06% | 0.010173654 | Astacin (Peptidase family M12A) |
| | PF01510 | 0.24% | 0.05% | 0.013999522 | N-acetylmuramoyl-L-alanine amidase |
| | PF01607 | 0.82% | 0.35% | 0.013820609 | Chitin binding Peritrophin-A domain |
| | PF01683 | 0.24% | 0.05% | 0.02933154 | EB module |
| | PF01757 | 0.34% | 0.09% | 0.01954313 | Acyltransferase family |
| | PF02244 | 0.19% | 0.04% | 0.047330812 | Carboxypeptidase activation peptide |
| | PF02958 | 0.53% | 0.13% | 0.000469997 | Ecdysteroid kinase |
| | PF03015 | 0.43% | 0.14% | 0.018249708 | Male sterility protein |
| | PF03145 | 0.87% | 0.29% | 0.000469997 | Seven in absentia protein family |
| | PF04083 | 0.34% | 0.06% | 0.00082408 | Partial alpha/beta-hydrolase lipase region |
| | PF06585 | 0.82% | 0.23% | 3.61E-05 | Haemolymph juvenile hormone binding protein (JHBP) |

| | | | | | |
|---|---|---|---|---|---|
| | PF07732 | 0.24% | 0.05% | 0.02933154 | Multicopper oxidase |
| | PF07993 | 0.58% | 0.18% | 0.003705549 | Male sterility protein |
| | PF12796 | 1.83% | 1.02% | 0.00667198 | Ankyrin repeats (3 copies) |
| | PF13637 | 0.92% | 0.41% | 0.013477688 | Ankyrin repeats (many copies) |
| | PF13855 | 2.12% | 0.97% | 2.41E-05 | Leucine rich repeat |
| **Head** | | | | | |
| | PF00019 | 0.38% | 0.07% | 0.00636893 | Transforming growth factor beta like domain |
| | PF00024 | 0.38% | 0.07% | 0.00636893 | PAN domain |
| | PF00067 | 2.98% | 0.74% | 4.79E-16 | Cytochrome P450 |
| | PF00094 | 0.32% | 0.07% | 0.048052598 | von Willebrand factor type D domain |
| | PF00100 | 0.57% | 0.11% | 0.000419475 | Zona pellucida-like domain |
| | PF00106 | 1.33% | 0.44% | 0.000146402 | short chain dehydrogenase |
| | PF00128 | 0.38% | 0.09% | 0.039392288 | Alpha amylase, catalytic domain |
| | PF00151 | 0.44% | 0.12% | 0.039392288 | Lipase |
| | PF00201 | 0.95% | 0.22% | 2.40E-05 | UDP-glucoronosyl and UDP-glucosyl transferase |
| | PF00232 | 0.57% | 0.12% | 0.00127824 | Glycosyl hydrolase family 1 |
| | PF00348 | 0.76% | 0.13% | 4.21E-06 | Polyprenyl synthetase |
| | PF00379 | 2.28% | 0.38% | 3.78E-20 | Insect cuticle protein |
| | PF00688 | 0.32% | 0.07% | 0.048052598 | TGF-beta propeptide |
| | PF00732 | 0.70% | 0.17% | 0.00081502 | GMC oxidoreductase |
| | PF01061 | 0.51% | 0.14% | 0.020057828 | ABC-2 type transporter |
| | PF01151 | 0.51% | 0.11% | 0.00295571 | GNS1/SUR4 family |
| | PF01347 | 0.32% | 0.06% | 0.030771274 | Lipoprotein amino terminal region |
| | PF01391 | 0.38% | 0.07% | 0.00636893 | Collagen triple helix repeat (20 copies) |
| | PF01562 | 0.32% | 0.07% | 0.048052598 | Reprolysin family propeptide |
| | PF01607 | 1.08% | 0.35% | 0.00081502 | Chitin binding Peritrophin-A domain |
| | PF01757 | 0.38% | 0.09% | 0.039392288 | Acyltransferase family |
| | PF03015 | 0.57% | 0.14% | 0.003726942 | Male sterility protein |
| | PF04083 | 0.32% | 0.06% | 0.030771274 | Partial alpha/beta-hydrolase lipase region |
| | PF05199 | 0.70% | 0.17% | 0.00081502 | GMC oxidoreductase |
| | PF06585 | 1.14% | 0.23% | 9.04E-08 | Haemolymph juvenile hormone binding protein (JHBP) |
| | PF07993 | 0.76% | 0.18% | 0.000419475 | Male sterility protein |

**Table S6.** Primer sequences used for RNA probe synthesis and *in situ* hybridization.

| **Gene** | **Gene ID** | **Forward (5-3')** | **Reverse (5'-3')** | **Length (bases)** |
|---|---|---|---|---|
| **lipocalin** | | | | |
| | RS008823 | TCGACGACAATCTCGACTGC | CGACCATCTGGCTGACATCA | 360 |
| | RS008881 | TGTCACAACCGAGACTGTGG | TAACAGGCGGAGTTGTCGAC | 403 |
| | RS008882 | ATTCTGCTTCGGACTGGTGT | ACAGTTCCTTGCACGCATGT | 407 |
| **GH1 (β-glucosidase)** | | | | |
| | RS004136 | GCTCATCCCATCTTCTCTGA | TGGGTCGATGAAATTCACTT | 567 |
| | RS004624 | TGCAAGAGCAAGAACACACC | GACGGCTCTTTTCAGCAATC | 1013 |
| **GGPP synthase** | | | | |
| | RS100016 | TGGAGGACATATTTGGCGTG | TGACGAGGTCAGCTTTGTTC | 378 |

**Table S7.** Lipocalin family genes in *Reticulitermes speratus*.

| Gene ID | Genomic position | Subclass | RNA-Seq, head (rpkm)* | | | | | | RNA-Seq, thorax + abdomen (rpkm)* | | | | | |
|---|---|---|---|---|---|---|---|---|---|---|---|---|---|---|
| | | | RF | RM | SF | SM | WF | WM | RF | RM | SF | SM | WF | WM |
| RS000420 | scaffold_1:11779874-11785972(+) | | 296.4 | 277.2 | 185.4 | 197.7 | 302.8 | 267.3 | 252.6 | 296.1 | 227.2 | 239.7 | 223.1 | 245.5 |
| RS004657 | scaffold_20:145341-153000(+) | Clade A (SOL1 family) | 80.2 | 142.4 | 385.9 | 320.6 | 1078.5 | 689.9 | 364.0 | 713.9 | 1738.9 | 2026.9 | 2685.3 | 2349.5 |
| RS005301 | scaffold_222:512009-520780(+) | | 46.0 | 42.3 | 35.2 | 42.5 | 43.5 | 32.1 | 111.6 | 116.7 | 112.2 | 86.2 | 107.1 | 108.8 |
| RS005409 | scaffold_228:179179-188294(+) | | 0.7 | 1.9 | 0.3 | 0.6 | 2.2 | 1.5 | 0.5 | 0.7 | 0.8 | 0.9 | 17.9 | 39.9 |
| RS008601 | scaffold_378:44567-47642(-) | Clade A (SOL1 family) | 11.2 | 9.4 | 8.2 | 9.3 | 12.0 | 13.5 | 8.1 | 8.2 | 5.4 | 7.5 | 6.5 | 6.8 |
| RS008823 | scaffold_387:404890-413918(-) | Clade A (SOL1 family) | 6.2 | 7.0 | 82.8 | 80.6 | 8.3 | 6.0 | 0.4 | 0.6 | 25.3 | 21.9 | 1.2 | 2.4 |
| RS008824 | scaffold_387:433200-439194(-) | Clade A (SOL1 family) | 0.2 | 0.2 | 8.7 | 8.4 | 2.6 | 2.7 | 0.0 | 0.0 | 4.6 | 3.5 | 1.2 | 0.7 |
| RS008881 | scaffold_39:1097117-1100793(-) | Clade B | 13.5 | 13.8 | 9.4 | 10.6 | 13.9 | 15.6 | 4005.2 | 11.0 | 4.1 | 5.2 | 5.0 | 5.3 |
| RS008882 | scaffold_39:1102331-1104831(-) | Clade B | 2228.8 | 2301.4 | 1432.2 | 1426.5 | 2993.8 | 3237.5 | 60.9 | 73.2 | 151.9 | 170.0 | 124.5 | 96.2 |
| RS008884 | scaffold_39:1122915-1126368(-) | Clade B | 3.8 | 2.4 | 2.6 | 2.9 | 2.7 | 3.4 | 755.1 | 1.7 | 3.0 | 1.7 | 1.0 | 1.8 |
| RS009761 | scaffold_43:3529390-3534162(+) | | 11.5 | 5.7 | 19.4 | 19.8 | 13.9 | 17.3 | 9.8 | 15.1 | 9.6 | 6.7 | 17.5 | 18.8 |
| RS010556 | scaffold_48:3522458-3532425(-) | | 0.0 | 0.0 | 0.0 | 0.0 | 0.0 | 0.0 | 0.0 | 0.0 | 0.0 | 0.0 | 0.0 | 0.0 |
| RS011706 | scaffold_55:2524382-2537041(-) | | 176.8 | 167.1 | 224.3 | 233.4 | 275.8 | 266.7 | 733.1 | 731.5 | 996.7 | 830.5 | 2103.0 | 1829.7 |
| RS012785 | scaffold_62:2793705-2806073(+) | | 3460.8 | 3284.3 | 2435.8 | 2137.1 | 797.0 | 798.1 | 1368.6 | 1480.2 | 349.6 | 313.0 | 185.7 | 306.4 |
| RS013912 | scaffold_757:39734-48788(-) | Clade A (SOL1 family) | 0.6 | 0.6 | 1.3 | 2.0 | 63.9 | 41.4 | 0.4 | 2.1 | 1.4 | 0.8 | 10.0 | 8.3 |
| RS013913 | scaffold_757:58631-120503(-) | Clade A (SOL1 family) | 20.8 | 11.2 | 1.4 | 0.8 | 16.8 | 12.5 | 7.3 | 1.6 | 0.5 | 0.0 | 7.8 | 10.8 |
| RS013914 | scaffold_757:88485-96225(-) | Clade A (SOL1 family) | 13.5 | 18.1 | 45.5 | 32.7 | 10.1 | 10.7 | 9.5 | 4.0 | 13.5 | 13.5 | 1.2 | 1.6 |
| RS014740 | scaffold_867:52755-60543(-) | | 172.9 | 161.9 | 199.5 | 226.0 | 130.8 | 196.1 | 107.5 | 116.0 | 267.4 | 267.9 | 151.0 | 142.7 |

*RF: female reproductives (queens), RM: male reproductives (kings), SF: female soldiers, SM: male soldiers, WF: female workers, WM: male workers.

**Table S8.** Cellulase genes in *Reticulitermes speratus*.

| Gene ID | Genomic position | Subclass | RNA-Seq, head (rpkm)* | | | | | | RNA-Seq, thorax+abdomen (rpkm)* | | | | | |
|---|---|---|---|---|---|---|---|---|---|---|---|---|---|---|
| | | | RF | RM | SF | SM | WF | WM | RF | RM | SF | SM | WF | WM |
| RS004136 | scaffold_186: 490588-509600(-) | GH1 | 5.0 | 3.2 | 15.6 | 9.8 | 1.1 | 13.0 | 1603.9 | 2018.8 | 68.3 | 55.1 | 2105.6 | 2036.8 |
| RS004137 | scaffold_186: 540509-564347(-) | GH1 | 60.7 | 65.7 | 22.3 | 22.6 | 80.2 | 78.0 | 7.9 | 11.8 | 10.6 | 11.1 | 17.5 | 17.2 |
| RS004143 | scaffold_186: 624299-638822(+) | GH1 | 8.3 | 10.0 | 4.1 | 3.5 | 1.3 | 1.7 | 38.9 | 52.9 | 39.9 | 47.2 | 8.6 | 9.4 |
| RS004144 | scaffold_186: 640651-659530(+) | GH1 | 1.7 | 2.0 | 1.1 | 1.4 | 1.9 | 3.0 | 67.8 | 56.6 | 61.6 | 55.3 | 58.5 | 57.6 |
| RS004146 | scaffold_186: 661301-665599(+) | GH1 | 16.6 | 13.4 | 7.2 | 5.3 | 25.1 | 24.1 | 25.6 | 30.8 | 33.7 | 33.2 | 76.9 | 73.4 |
| RS100005 | scaffold_186: 740796-759807(+) | GH1 | 3.2 | 3.7 | 4.1 | 4.5 | 13.1 | 13.1 | 31.0 | 47.1 | 44.6 | 49.5 | 137.6 | 124.7 |
| RS100006 | scaffold_186: 765348-787391(+) | GH1 | 103.9 | 100.9 | 97.9 | 96.0 | 76.5 | 106.9 | 127.1 | 195.3 | 418.6 | 411.0 | 198.1 | 211.7 |
| RS100007 | scaffold_2:3954538-3976450(+) | GH1 | 134.5 | 143.0 | 70.7 | 71.1 | 135.7 | 164.7 | 5.5 | 5.9 | 10.5 | 9.7 | 11.9 | 12.1 |
| RS004147 | scaffold_2:3989223-4009606(+) | GH1 | 24.1 | 23.7 | 10.8 | 8.0 | 22.0 | 20.5 | 17.8 | 22.5 | 31.3 | 33.6 | 49.6 | 47.5 |
| RS004149 | scaffold_6:2019334-2035300(+) | GH1 | 45.0 | 38.6 | 44.2 | 39.1 | 67.1 | 50.1 | 26.1 | 18.4 | 56.6 | 61.3 | 46.6 | 47.8 |
| RS004623 | scaffold_6:2047153-2060696(+) | GH1 | 37.6 | 34.2 | 71.5 | 80.0 | 17.8 | 24.5 | 36.5 | 57.7 | 81.9 | 93.0 | 41.0 | 42.0 |
| RS004624 | scaffold_6:2070232-2087920(+) | GH1 | 2.0 | 2.5 | 12.1 | 12.4 | 6.7 | 7.6 | 1748.4 | 0.8 | 4.1 | 6.8 | 2.1 | 2.9 |
| RS012436 | scaffold_6:2095616-2111294(+) | GH1 | 6.5 | 7.1 | 3.9 | 5.8 | 81.5 | 76.3 | 0.6 | 0.6 | 1.5 | 3.3 | 25.0 | 19.8 |
| RS012437 | scaffold_186: 676910-679770(+) | GH1 | 1.4 | 0.8 | 1.3 | 1.2 | 3.2 | 5.8 | 7.3 | 13.5 | 26.2 | 38.4 | 18.3 | 19.5 |
| RS012439 | scaffold_186: 692051-703746(+) | GH1 | 99.1 | 106.4 | 136.0 | 134.0 | 126.7 | 113.4 | 47.0 | 65.2 | 96.3 | 106.0 | 129.5 | 108.4 |
| RS012440 | scaffold_186: 706674-735672(+) | GH1 | 45.4 | 44.3 | 42.4 | 35.7 | 38.8 | 34.9 | 40.6 | 50.7 | 40.0 | 34.4 | 40.3 | 36.2 |
| RS006396 | scaffold_27:1888536-1904998(+) | GH9 | 32.3 | 36.7 | 35.1 | 30.2 | 28.5 | 40.5 | 21.9 | 23.0 | 27.9 | 31.2 | 31.4 | 28.4 |
| RS012684 | scaffold_611: 179946-189568(+) | GH9 | 16.8 | 17.7 | 7.4 | 9.0 | 27.5 | 23.4 | 17.0 | 23.2 | 0.7 | 0.6 | 22.4 | 19.2 |
| RS012687 | scaffold_611: 209423-219382(+) | GH9 | 49.3 | 62.2 | 69.6 | 68.5 | 69.3 | 116.7 | 1232.1 | 1411.5 | 36.8 | 43.4 | 3032.0 | 2772.4 |
| RS100101 ** | scaffold_611: 225890-228919(+) + scaffold_564: 265652-267717(-) | GH9 | 4.6 | 4.8 | 8.2 | 7.6 | 5.2 | 104.4 | 14962.2 | 19873.5 | 304.6 | 366.9 | 18894.0 | 16784.5 |

*RF: female reproductives (queens), RM: male reproductives (kings), SF: female soldiers, SM: male soldiers, WF: female workers, WM: male workers.

**RS100101 (GH9) lies between scaffold_611 and scaffold_564 in the current gene model (Rspe OGS1.0).

**Table S9.** Lisozyme genes in *Reticulitermes speratus*.

| Gene ID | Genomic position | Subclass | RNA-Seq, head (rpkm)* | | | | | | RNA-Seq, thorax + abdomen (rpkm)* | | | | | |
|---|---|---|---|---|---|---|---|---|---|---|---|---|---|---|
| | | | RF | RM | SF | SM | WF | WM | RF | RM | SF | SM | WF | WM |
| RS000427 | scaffold_10:4251-11117(-) | lysozyme c-type | 0.0 | 0.2 | 0.2 | 0.2 | 0.0 | 0.0 | 0.2 | 0.0 | 1.8 | 2.9 | 1.0 | 3.3 |
| RS002400 | scaffold_1370:6473-16438(-) | lysozyme c-type | 0.0 | 0.0 | 0.0 | 0.0 | 0.0 | 0.7 | 2.4 | 2.5 | 7.1 | 14.2 | 15.9 | 23.5 |
| RS003406 | scaffold_16:6014313-6016559(-) | lysozyme c-type | 0.0 | 0.2 | 5.1 | 3.6 | 0.0 | 1.0 | 22.5 | 33.0 | 153.6 | 285.8 | 64.6 | 64.3 |
| RS008613 | scaffold_378:288875-289873(-) | lysozyme c-type | 0.6 | 0.7 | 0.0 | 1.6 | 0.0 | 0.3 | 39.6 | 44.8 | 1676.8 | 1384.0 | 250.9 | 224.9 |
| RS014698 | scaffold_859:24243-30080(+) | lysozyme c-type | 0.5 | 0.0 | 0.9 | 1.9 | 0.2 | 5.9 | 9.1 | 203.6 | 2144.6 | 2615.8 | 1739.5 | 1161.5 |
| RS100001 | scaffold_1097:7906-14032(+) | lysozyme c-type | 0.0 | 0.0 | 0.0 | 0.2 | 0.0 | 2.0 | 4.9 | 23.3 | 5.3 | 1.9 | 269.3 | 219.4 |
| RS100002 | scaffold_1097:42864-48573(+) | lysozyme c-type | 97.4 | 78.3 | 266.1 | 276.1 | 127.1 | 178.9 | 131.3 | 137.7 | 211.0 | 191.1 | 148.7 | 156.0 |
| RS100004 | scaffold_16:6112988-6118555(-) | lysozyme c-type | 1.6 | 2.3 | 1.2 | 1.6 | 1.2 | 2.5 | 2.9 | 1.2 | 2.2 | 2.2 | 0.3 | 2.1 |
| RS100022 | scaffold_859:4935-6501(+) | lysozyme c-type | 1.2 | 0.5 | 2.8 | 2.3 | 1.8 | 2.8 | 0.9 | 1.1 | 2.6 | 1.8 | 4.3 | 6.1 |
| RS100023 | scaffold_859:48011-50947(+) | lysozyme c-type | 0.4 | 0.4 | 0.6 | 0.0 | 2.0 | 1.6 | 0.8 | 0.3 | 1.7 | 1.4 | 2.2 | 1.7 |
| RS100024 | scaffold_859:63516-69373(+) | lysozyme c-type | 0.5 | 0.8 | 0.7 | 0.7 | 0.2 | 1.0 | 35.4 | 92.5 | 951.1 | 728.6 | 1517.7 | 1464.3 |
| RS100025 | scaffold_859:85037-90894(+) | lysozyme c-type | 0.0 | 0.0 | 0.2 | 0.2 | 0.5 | 0.6 | 19.1 | 40.6 | 353.4 | 298.3 | 621.1 | 639.9 |
| RS100026 | scaffold_859:99866-105566(+) | lysozyme c-type | 24.4 | 18.8 | 17.4 | 14.3 | 8.9 | 9.1 | 19.6 | 30.4 | 14.6 | 13.1 | 13.8 | 13.7 |
| RS006054 | scaffold_257:1332-27487(-) | lysozyme i-type | 0.8 | 0.0 | 1.9 | 3.8 | 0.8 | 2.1 | 50.9 | 130.6 | 4359.1 | 4978.4 | 609.9 | 511.6 |
| RS008547 | scaffold_374:174359-193956(+) | lysozyme i-type | 0.4 | 0.6 | 1.5 | 4.7 | 0.0 | 0.2 | 67.3 | 180.6 | 3849.1 | 4184.8 | 694.7 | 545.6 |
| RS015579 | scaffold_997:67510-70385(+) | lysozyme i-type | 0.2 | 0.0 | 0.4 | 0.0 | 0.2 | 0.2 | 0.6 | 5.1 | 10.5 | 19.4 | 45.6 | 35.2 |

*RF: female reproductives (queens), RM: male reproductives (kings), SF: female soldiers, SM: male soldiers, WF: female workers, WM: male workers.

**Table S10.** GGPP synthase genes in *Reticulitermes speratus*.

| Gene ID | Genomic position | Subclass | RNA-Seq, head (rpkm)* | | | | | | RNA-Seq, thorax + abdomen (rpkm)* | | | | | |
|---|---|---|---|---|---|---|---|---|---|---|---|---|---|---|
| | | | RF | RM | SF | SM | WF | WM | RF | RM | SF | SM | WF | WM |
| RS100010 | scaffold_31:2794158-2806101(+) | derived | 44.1 | 37.0 | 59.8 | 59.9 | 11.3 | 5.6 | 3.5 | 3.8 | 53.4 | 51.9 | 2.2 | 2.2 |
| RS007480 | scaffold_31:2813416-2826556(+) | derived | 11.8 | 9.6 | 93.5 | 88.3 | 6.4 | 6.0 | 4.6 | 0.3 | 52.9 | 55.9 | 0.5 | 0.3 |
| RS007481 | scaffold_31:2831788-2846259(+) | derived | 47.9 | 41.2 | 2.2 | 3.5 | 21.7 | 20.2 | 15.3 | 2.0 | 2.7 | 1.8 | 0.9 | 1.5 |
| RS100011 | scaffold_31:2854401-2866719(+) | derived | 14.8 | 15.2 | 6.9 | 8.7 | 14.8 | 12.1 | 5.9 | 6.5 | 7.1 | 6.3 | 5.4 | 5.8 |
| RS100012 | scaffold_31:2881695-2891432(+) | derived | 3.8 | 5.4 | 115.8 | 117.2 | 2.6 | 4.2 | 6.3 | 8.6 | 85.4 | 82.3 | 6.7 | 6.5 |
| RS007482 | scaffold_31:2898315-2910945(+) | derived | 126.6 | 133.6 | 51.9 | 50.2 | 79.8 | 69.5 | 4.9 | 5.1 | 23.5 | 28.7 | 1.2 | 1.3 |
| RS100013 | scaffold_31:2918062-2928930(+) | derived | 76.8 | 71.9 | 3.0 | 5.0 | 34.0 | 22.6 | 0.3 | 0.2 | 2.9 | 0.6 | 0.1 | 0.2 |
| RS100014 | scaffold_31:2939721-2948893(+) | derived | 0.4 | 0.3 | 1.5 | 2.3 | 0.4 | 0.2 | 0.7 | 1.0 | 1.6 | 1.3 | 0.1 | 0.6 |
| RS100015 | scaffold_31:2964788-2973091(+) | derived | 11.1 | 10.0 | 619.1 | 582.8 | 20.4 | 15.7 | 0.8 | 0.0 | 419.8 | 426.1 | 1.7 | 1.0 |
| RS100016 | scaffold_31:2982793-2992579(+) | derived | 192.0 | 202.8 | 1482.2 | 1440.8 | 387.8 | 278.4 | 3.2 | 3.7 | 998.3 | 836.4 | 38.1 | 38.5 |
| RS100017 | scaffold_31:3003886-3010980(+) | derived | 3.0 | 9.1 | 43.4 | 37.9 | 44.5 | 33.6 | 0.2 | 0.2 | 20.9 | 25.4 | 0.0 | 0.1 |
| RS007483 | scaffold_31:3020421-3029674(+) | derived | 10.8 | 13.4 | 36.3 | 35.9 | 32.4 | 26.9 | 1.1 | 0.6 | 26.1 | 26.4 | 2.4 | 1.7 |
| RS007484 | scaffold_31:3031695-3042573(+) | possibly ancestral | 7.4 | 8.7 | 8.9 | 9.3 | 9.5 | 9.5 | 9.1 | 8.5 | 8.7 | 7.9 | 10.5 | 7.9 |

*RF: female reproductives (queens), RM: male reproductives (kings), SF: female soldiers, SM: male soldiers, WF: female workers, WM: male workers.

**Table S11.** TY family genes in *Reticulitermes speratus*.

| Gene ID | Genomic position | RNA-Seq, head (rpkm)* | | | | | | RNA-Seq, thorax + abdomen (rpkm)* | | | | | |
|---|---|---|---|---|---|---|---|---|---|---|---|---|---|
| | | RF | RM | SF | SM | WF | WM | RF | RM | SF | SM | WF | WM |
| RS001196 | scaffold_113:185192-185350(+) | 112 | 101 | 2,268 | 2,229 | 49,356 | 95,342 | 68 | 30 | 46 | 69 | 168 | 128 |
| RS001197 | scaffold_113:224999-225178(+) | 13,008 | 12,554 | 66,324 | 64,899 | 149,936 | 193,248 | 125 | 213 | 12,895 | 16,751 | 10,586 | 9,606 |
| RS001198 | scaffold_113:240134-240322(+) | 4,781 | 4,683 | 30,257 | 32,502 | 53,571 | 136,254 | 88 | 203 | 3,755 | 4,045 | 5,145 | 4,993 |

*RF: female reproductives (queens), RM: male reproductives (kings), SF: female soldiers, SM: male soldiers, WF: female workers, WM: male workers.

1324 **Table S12.** Sex determination genes in *Reticulitermes speratus*.

| Gene name | Symbol | OrthoDB7_ID | *Drosophila* homolog accesion No | Gene ID | Expression Differences between sexes (FDR) | | Expression Differences among castes (FDR) | |
|---|---|---|---|---|---|---|---|---|
| | | | | | Head | Thorax + abdomen | Head | Thorax + abdomen |
| daughterless | *da* | EOG7BGW18 | NM_001273411 | RS000156 | 1 | 0.99 | 0.75 | 0.0037 |
| Hairy/deadpan | *dpn* | EOG7X6ZFD | NM_057575 | RS006294 | 1 | 0.63 | 0.7 | 0.00073 |
| Hairy/deadpan | *dpn* | EOG7X6ZFD | NM_057575 | RS006295 | 1 | 0.92 | 0.86 | 1.12E-05 |
| degringolade | *dgrn* | EOG7KX50F | NM_141339 | - | - | - | - | - |
| dissatisfaction | *dsf* | EOG7VJ5CF | NM_001273180 | RS013719 | N/A | N/A | N/A | N/A |
| doublesex | *dsx* | EOG77DWNS | NM_169202 | - | - | - | - | - |
| Doublesex-Mab related 11B | *Dmrt11B* | EOG7TXKGH | NM_078591 | RS007930 | 1 | 0.084 | 7.85E-11 | 0.34 |
| Doublesex-Mab related 93B | *Dmrt93B* | EOG7B8S48 | NM_079704 | RS006912 | N/A | N/A | N/A | N/A |
| Doublesex-Mab related 99B | *Dmrt99B* | EOG718KC7 | NM_079825 | RS002870 | N/A | N/A | N/A | N/A |
| extramacrochaetae | *emc* | EOG7QCMBK | NM_079152 | RS011015 | 1 | 0.71 | 0.54 | 0.9 |
| female lethal d | *fl(2)d* | EOG7KT8QP | NM_166010 | RS002775 | 1 | 0.89 | 0.99 | 0.0064 |
| fruitless | *fru* | EOG7S84VN | NM_079673 | RS001598 | 1 | 0.99 | 0.85 | 0.065 |
| groucho | *gro* | EOG7BD0RV | NM_001260380 | RS013444 | 1 | 1 | 0.44 | 0.07 |
| groucho | *gro* | EOG7BD0RV | NM_001260380 | RS010817 | 1 | 0.91 | 0.44 | 0.78 |
| hermaphrodite | *her* | EOG7JXFJ7 | NM_001273577 | RS010220 | 1 | 0.87 | 0.89 | 0.0016 |
| hopscotch | *hop* | EOG74JNNN | NM_078564 | RS002898 | 1 | 0.99 | 0.06 | 0.08 |
| intersex | *ix* | EOG7162KM | NM_136833 | RS007777 | 1 | 0.86 | 0.44 | 0.98 |
| outstreched | *os* | EOG7TJFZ9 | NM_001103545 | RS015475 | 1 | 0.08 | 0.32 | 1.87E-08 |
| ovarian tumor | *out* | EOG7N9B8P | NM_001272403 | RS009292 | 1 | 0.51 | 3.79E-07 | 8.00E-07 |
| ovo/shavenbaby | *ovo* | EOG7C0766 | NM_001169202 | RS015380 | 1 | 0.83 | 0.003 | 0.24 |
| runt | *run* | EOG73RPRJ | NM_078700 | RS006486 | 1 | 0.99 | 0.71 | 0.23 |
| sansfille | *snf* | EOG78WZ6X | NM_078490 | RS011497 | 1 | 0.41 | 0.91 | 0.035 |
| Sex-lethal | *Sxl* | EOG72ZQVX | NM_001031891 | RS002675 | 1 | 0.69 | 0.024 | 0.32 |
| scute/sisterlessB | *sc* | EOG7HJ870 | NM_057455 | RS001853 | 1 | 0.94 | 2.13E-14 | 0.00021 |
| scute/sisterlessB | *sc* | EOG7HJ870 | NM_057455 | RS001854 | N/A | N/A | N/A | N/A |
| scute/sisterlessB | *sc* | EOG7HJ870 | NM_057455 | RS008845 | N/A | N/A | N/A | N/A |
| sisterless-A | *sisA* | EOG7FZBDN | NM_078561 | - | - | - | - | - |
| standstill | *stil* | EOG7RVNX6 | NM_057404 | - | - | - | - | - |
| transformer | *tra* | EOG7HN4HG | NM_079390 | RS002588 | 1 | 0.70 | 0.73 | 0.52 |

| | | | | | | | | |
|---|---|---|---|---|---|---|---|---|
| transformer2 | *tra2* | EOG7Z3SMD | NM_057416 | RS011357 | 1 | 0.87 | 0.82 | 0.76 |
| virillizer | *vir* | EOG7DRWGS | NM_080161 | RS015320 | 1 | 0.75 | 0.51 | 0.41 |
| P-element somatic inhibitor | *PSI* | EOG7K19SX | NM_001110343 * | RS012378 | 0.68 | 1 | 0.08 | 5.77E-04 |
| IGF-II mRNA BP | *IMP* | EOG738BJT | XM_004929848 * | RS011169 | 0.98 | 1 | 9.54E-13 | 0.14 |
| Masculinizer | *Masc* | EOG7QS39K | AB840788 * | RS005147 | 0.62 | 1 | 0.11 | 0.52 |
| Feminizer | *Fem* | | AB840787 * | - | - | - | - | - |

N/A: data is not available.
*Accession numbers of *Bombyx mori* orthologs, which are absent in the *Drosophila* genome.

**Table S13.** Histone modifying enzyme genes in *Reticulitermes speratus*, and their orthologs in *Zootermopsis nevadensis* and *Drosophila melanogaster*. Gene names were described based on those in human. Expression levels were compared between castes and between sexes in each of body parts (i.e., head and body). Asterisks indicate genes with significantly differential expressions (GLM analysis, FDR < 0.05).

| Gene name | Gene ID of orthologs | | | Expression Differences among castes (FDR) | | Expression Differences between sexes (FDR) | |
|---|---|---|---|---|---|---|---|
| | *D. melanogaster* | *Z. nevadensis* | *R. speratus* | Thorax + abdomen | Head | Thorax + abdomen | Head |
| **Histone acetyltransferases** | | | | | | | |
| HAT1 | FBgn0037376 | Znev_12488 | RS014176 | 0.41 | 0.409 | 0.985 | 1 |
| KAT2A | FBgn0020388 | Znev_04968 | RS007246 | 0.148 | 0.172 | 0.570 | 1 |
| EP300 | FBgn0261617 | Znev_08401 | RS009769 | 0.383 | 0.775 | 0.379 | 1 |
| TAF1 | FBgn0010355 | Znev_01981 | RS006173 | 8.02E-05* | 0.655 | 0.428 | 1 |
| KAT5 | FBgn0026080 | Znev_00128 | RS009782 | 0.003* | 0.360 | 0.794 | 1 |
| KAT6A | FBgn0034975 | Znev_04899, | RS008273 | 0.282 | 0.538 | 0.606 | 1 |
| | | Znev_04900, | | | | | |
| | | Znev_06347 | | | | | |
| KAT7 | FBgn28387 | Znev_14581 | RS001974 | 4.01E-04* | 0.192 | 0.476 | 1 |
| KAT8 | - | Znev_09388 | RS014043 | 0.010* | 0.731 | 0.985 | 1 |
| KAT8 | FBgn0014340 | Znev_09335 | RS014973 | 0.972 | 0.003* | 0.906 | 1 |
| ELP3 | FBgn0031604 | Znev_03596, | RS007305 | 0.002* | 0.568 | 0.428 | 1 |
| | | Znev_16248 | | | | | |
| GTF3C4 | - | Znev_04938 | RS006156 | 0.020* | 0.437 | 0.805 | 1 |
| NCOA2 | - | Znev_05082, | RS006636 | 1.56E-04* | 2.83E-04* | 0.840 | 1 |
| | | Znev_05083 | | | | | |
| CLOCK | FBgn0023076 | - | RS010134 | 0.197 | 0.855 | 0.965 | 1 |
| CSRP2BP | FBgn0032691 | Znev_02989 | RS007957 | 0.290 | 0.122 | 0.913 | 1 |
| ATF2 | FBgn0265193 | Znev_01083 | RS004334 | 0.233 | 0.471 | 0.795 | 1 |
| MGEA5 | FBgn0038870 | Znev_10779 | RS014601 | 8.72E-09* | 0.777 | 0.810 | 1 |
| NAA60 | FBgn0036039 | Znev_04119 | RS011057 | 0.019* | 0.941 | 0.725 | 1 |
| **Histone deacetylases** | | | | | | | |
| HDAC1 | FBgn0015805 | Znev_03795 | RS012536 | 0.020* | 0.005* | 0.683 | 1 |
| HDAC3 | FBgn0025825 | Znev_05602, | RS008767 | 0.008* | 0.067 | 0.891 | 1 |
| | | Znev_18002 | | | | | |
| HDAC4 | FBgn0041210 | Znev_00349 | RS004692 | 0.006* | 0.604 | 0.841 | 1 |
| HDAC6 | FBgn0026428 | Znev_02211 | RS001937 | 0.002* | 0.022* | 0.922 | 1 |
| HDAC8 | - | Znev_12928 | RS010779 | 0.246 | 0.706 | 0.458 | 1 |
| HDAC11 | FBgn0051119 | Znev_10901 | RS007375 | 0.606 | 1.34E-05* | 0.881 | 1 |
| SIRT1 | FBgn0024291 | Znev_11203 | RS007459 | 0.157 | 4.99E-04* | 0.992 | 1 |
| SIRT2 | FBgn0038788 | Znev_11971 | RS013047 | 0.291 | 0.723 | 0.598 | 1 |
| SIRT3 | - | Znev_01239 | RS012446 | 0.415 | 0.007* | 0.901 | 1 |
| SIRT4 | FBgn0029783 | Znev_10250 | RS011722 | 0.035* | 0.896 | 0.849 | 1 |

| | | | | | | | |
|---|---|---|---|---|---|---|---|
| SIRT5 | - | Znev_14842 | RS008195 | 0.131 | 0.315 | 0.598 | 1 |
| SIRT6 | FBgn0037802 | Znev_09433 | RS010824 | 3.54E-07* | 0.054 | 0.322 | 1 |
| SIRT7 | FBgn0039631 | Znev_03848 | RS012147 | 0.047* | 0.032* | 0.598 | 1 |
| **Histone methyltransferases** | | | | | | | |
| PRMT1 | FBgn0037834 | Znev_11976 | RS013040 | 0.342 | 7.78E-04* | 0.821 | 1 |
| CARM1 | FBgn0037770 | Znev_11468 | RS009701 | 0.669 | 0.879 | 0.978 | 1 |
| PRMT5 | FBgn0015925 | Znev_08220 | RS011600 | 4.00E-04* | 0.612 | 0.991 | 1 |
| PRMT7 | FBgn0034817 | Znev_07771 | RS014915 | 0.549 | 0.668 | 0.745 | 1 |
| SUV39H2 | FBgn0263755 | Znev_00097 | RS015134 | 0.759 | 0.633 | 0.662 | 1 |
| EHMT1 | FBgn0040372 | Znev_05631 | RS003423 | 0.010* | 0.181 | 0.929 | 1 |
| EHMT1 | - | - | RS001807 | 0.138 | 0.163 | 0.936 | 1 |
| SETDB1 | FBgn0086908 | Znev_15214 | RS011823 | 5.88E-06* | 0.224 | 0.634 | 1 |
| KMT2B | FBgn0003862 | Znev_03032 | RS005841 | 0.003* | 0.002* | 0.828 | 1 |
| KMT2C | FBgn0023518 | Znev_09224 | RS010172 | 0.263 | 0.765 | 0.502 | 1 |
| KMT2C | FBgn0263667 | Znev_09226 | RS010173 | 0.161 | 0.706 | 0.448 | 1 |
| KMT2E | FBgn0036398 | Znev_06854 | RS008681 | 0.083 | 0.972 | 0.298 | 1 |
| SETD1A | FBgn0040022 | Znev_03918 | RS010178 | 4.36E-04* | 0.286 | 0.810 | 1 |
| ASH1L | FBgn0005386 | Znev_16755 | RS005572 | 0.216 | 0.641 | 0.639 | 1 |
| SETD2 | FBgn0030486 | Znev_02205 | RS014526 | 0.081 | 0.723 | 0.906 | 1 |
| WHSC1L1 | FBgn0039559 | Znev_13059, | RS006777 | 4.42E-06* | 0.052 | 0.403 | 1 |
| | | Znev_14928 | | | | | |
| SMYD3 | FBgn0011566 | Znev_00597 | RS009274 | 0.020* | 0.008* | 0.443 | 1 |
| DOT1L | FBgn0264495 | Znev_02995 | RS009425 | 9.14E-05* | 1.16E-07* | 0.903 | 1 |
| SETD8 | FBgn0011474 | Znev_05607 | RS008772 | 0.472 | 1.13E-04* | 0.559 | 1 |
| SUV420H1 | FBgn0025639 | Znev_05656 | RS005423 | 0.024* | 0.003* | 0.961 | 1 |
| EXH2 | FBgn0000629 | Znev_02258 | RS003652 | 0.010* | 0.084 | 0.977 | 1 |
| SETMAR | FBgn0037841 | Znev_04699 | RS008713 | 6.48E-04* | 0.497 | 0.909 | 1 |
| SMYD4 | FBgn0033427 | Znev_01984 | RS006171 | 3.35E-04* | 1.69E-04* | 0.511 | 1 |
| SMYD5 | FBgn0038869 | Znev_01118 | RS004372 | 0.013* | 0.240 | 0.232 | 1 |
| SETD3 | FBgn0052732 | Znev_12254 | RS012270 | 0.545 | 0.005* | 0.874 | 1 |
| SETD4 | FBgn0053230 | Znev_05436 | RS004593 | 0.015* | 0.029* | 0.866 | 1 |
| **Histone demethylases** | | | | | | | |
| KDM1A | FBgn0260397 | Znev_07217 | RS006951 | 0.867 | 0.022* | 0.885 | 1 |
| KDM1A | - | Znev_00885 | RS005465 | 0.012* | 0.249 | 0.624 | 1 |
| KDM1A | - | Znev_00889 | RS005467 | 0.042* | 0.082 | 0.630 | 1 |
| KDM2A | FBgn0037659 | Znev_06646 | RS001102 | 0.294 | 0.297 | 0.915 | 1 |
| KDM3A | | Znev_17182, | RS010951 | 0.017* | 1.000 | 0.800 | 1 |
| | | Znev_09543 | | | | | |
| KDM4C | | Znev_07618 | RS014991 | 0.413 | 0.873 | 0.943 | 1 |
| KDM5A | KDM5 | Znev_17776 | RS002306 | 0.217 | 0.043* | 0.637 | 1 |
| KDM6A | Utx | Znev_00365 | RS002457 | 0.049* | 0.014* | 0.975 | 1 |

| | | | | | | | |
|---|---|---|---|---|---|---|---|
| KDM7A | -- | Znev_15990 | RS015526 | 0.061 | 0.294 | 0.760 | 1 |
| KDM8 | FBgn0035166 | Znev_07996 | RS011528 | 0.055 | 0.516 | 0.991 | 1 |
| JARID2 | FBgn0036004 | Znev_08621 | RS010046 | 0.035* | 0.165 | 0.560 | 1 |
| JMJD6 | FBgn0038948 | Znew_10481 | RS002889 | 0.008* | 0.153 | 0.876 | 1 |
| C14orf169 | FBgn0266570 | Znew_02698 | RS011375 | 1.51E-04* | 0.177 | 0.157 | 1 |

*FDR < 0.05

**Table S14.** DNA methylation-related genes in *Reticulitermes speratus*, and their orthologs in *Zootermopsis nevadensis* and *Drosophila melanogaster*. Gene names were described based on those in human. Expression levels were compared between castes and between sexes in each of body parts (i.e., head and body). Asterisks indicate genes with significantly differential expressions (GLM analysis, FDR < 0.05).

| **Gene name** | **Gene ID of orthologs** | | | **Expression Differences among castes (FDR)** | | **Expression Differences between sexes (FDR)** | |
|---|---|---|---|---|---|---|---|
| | ***D. melanogaster*** | ***Z. nevadensis*** | ***R. speratus*** | **Thorax + abdomen** | **Head** | **Thorax + abdomen** | **Head** |
| DNMT1 | - | Znev_18516 | RS003121 | 0.339 | 0.243 | 0.849 | 1 |
| DNMT3 | - | Znev_11906, Znev_06587 | RS003573 RS003574 | 0.147 | 0.006* | 0.993 | 1 |
| | | | | - | - | - | - |
| AGT | FBgn0024912 | Znev_07784 | RS014911 | 0.572 | 0.276 | 0.973 | 1 |
| CG9154 | FBgn0031777 | Znev_08521 | RS005202 | 0.023* | 0.512 | 0.219 | 1 |
| DMAP1 | FBgn0034537 | Znev_09468 | RS014390 | 0.069 | 0.269 | 0.842 | 1 |
| MBD-like | FBgn0027950 | Znev_00583 | RS002943 | 2.90E-06* | 0.002* | 0.202 | 1 |
| MBD-R2 | FBgn0038016 | Znev_06566, | RS013654, | 0.006* | 0.023* | 0.625 | 1 |
| | | Znev_01879 | RS012154 | 0.692 | 0.812 | 0.633 | 1 |
| TET | FBgn0263392 | Znev_11370 | RS012619 | 0.868 | 0.593 | 0.961 | 1 |
| TDG | FBgn0026869 | Znev_03074 | RS003560 | 0.012* | 0.083 | 0.711 | 1 |

*FDR < 0.05

**Table S15.** Odorant receptor (OR) genes in *Reticulitermes speratus.*

| Gene | Gene ID | Scaffold | Strand | Sequence* | Exons |
|---|---|---|---|---|---|
| RsOrco | RS006385 | 27 | - | complete | 7 |
| RsOr1 | RS010432 | 47 | - | fragment | 2 |
| RsOr2 | RS010431 | 47 | - | fragment | 1 |
| RsOr9 | RS007507 | 31 | + | NTE | 3 |
| RsOr10 | RS013588 | 703 | - | NTE | 3 |
| RsOr11 | RS005817 | 248 | - | fragment | 1 |
| RsOr12 | RS001899 | 1287 | + | fragment | 1 |
| RsOr13 | RS004141 | 186 | + | fragment | 2 |
| RsOr14 | RS004139 | 186 | + | complete | 3 |
| RsOr15 | RS004142 | 186 | + | fragment | 1 |
| RsOr16 | RS004140 | 186 | + | fragment | 1 |
| RsOr17 | RS004138 | 186 | + | fragment | 1 |
| RsOr26 | RS007955 | 337 | + | fragment | 1 |
| RsOr28 | RS013645 | 72 | + | fragment | 1 |
| RsOr30 | RS010974 | 50 | - | fragment | 1 |
| RsOr35 | RS000450 | 10 | - | complete | 6 |
| RsOr40 | RS013053 | 65 | - | fragment | 1 |
| RsOr41 | RS014428 | 807 | + | fragment | 1 |
| RsOr42 | RS001393 | 118 | - | NTE | 2 |
| RsOr43 | RS001394 | 118 | - | NTE | 5 |
| RsOr44 | RS001395 | 118 | - | fragment | 3 |
| RsOr45 | RS001396 | 118 | - | fragment | 4 |
| RsOr46 | RS011968 | 570 | + | fragment | 1 |
| RsOr47 | RS005073 | 212 | - | fragment | 4 |
| RsOr48 | RS008327 | 365 | - | fragment | 1 |
| RsOr56 | RS006316 | 269 | - | complete | 7 |
| RsOr66 | RS013051 | 65 | - | fragment | 1 |
| RsOr67 | RS013052 | 65 | - | fragment | 1 |
| RsOr68 | RS008460 | 37 | - | NTE | 3 |
| RsOr69 | RS008461 | 37 | - | fragment | 1 |
| RsOr70 | RS008462 | 37 | - | fragment | 1 |

*NTE: initial methionine (M) is missed in the transcript sequence. CTE: stop codon is missed in the transcript sequence.

**Table S16.** Gustatory receptor (GR) genes in *Reticulitermes speratus.*

| Genes | Gene ID | Scaffold | Strand | Sequence* | Exon |
|---|---|---|---|---|---|
| RsGr1 | RS001507 | 12 | - | complete | 8 |
| RsGr2 | RS001504 | 12 | - | NTE | 3 |
| RsGr3 | RS010223 | 458 | + | complete | 6 |
| RsGr4 | RS007965 | 338 | - | fragment | 3 |
| RsGr5a | RS013641 | 72 | + | fragment | 1 |
| RsGr5b | RS013642 | 72 | + | fragment | 1 |
| RsGr6 | RS013643 | 72 | - | complete | 7 |
| RsGr7 | RS001378 | 118 | + | complete | 6 |
| RsGr8 | RS011470 | 530 | + | CTE | 5 |
| RsGr9 | RS001377 | 118 | - | complete | 8 |
| RsGr10a | RS001379 | 118 | + | fragment | 1 |
| RsGr10b | RS001380 | 118 | + | fragment | 1 |
| RsGr11 | RS001506 | 12 | - | fragment | 1 |
| RsGr12 | RS001505 | 12 | - | fragment | 2 |
| RsGr13 | RS011406 | 53 | + | fragment | |
| RsGr14 | RS012780 | 62 | - | fragment | 1 |
| RsGr15 | RS005943 | 25 | + | NTE | 3 |
| RsGr16 | RS011323 | 524 | + | CTE | 1 |
| RsGr17 | RS009029 | 4 | - | complete | 1 |
| RsGr18 | RS011329 | 525 | + | complete | 1 |
| RsGr19 | RS015443 | 963 | - | NTE | 1 |
| RsGr20 | RS011328 | 525 | - | complete | 1 |
| RsGr21 | RS009385 | 410 | - | complete | 1 |
| RsGr22 | RS003203 | 155 | - | fragment | 1 |
| RsGr23 | RS003204 | 155 | + | fragment | 1 |

*NTE: initial methionine (M) is missed in the transcript sequence. CTE: stop codon is missed in the transcript sequence.

**Table S17.** Ionotropic receptor (IR) genes in *Reticulitermes speratus.*

| Gene | Gene ID | Scaffold | Strand | Sequence* | Exon |
|---|---|---|---|---|---|
| RsKAINATE1 | RS003449 | 1616 | - | CTE | 14 |
| RsKAINATE2 | RS001970 | 1291 | + | NTE | 16 |
| RsKAINATE3 | RS001366 | 1174 | + | complete | 16 |
| RsKAINATE4 | RS003544 | 166 | - | NTE | 16 |
| RsKAINATE5 | RS012852 | 63 | - | NTE | 16 |
| RsKAINATE6 | RS010902 | 5 | + | complete | 19 |
| RsNMDAR1 | RS012806 | 62 | - | complete | 17 |
| RsNMDAR2 | RS006920 | 293 | - | complete | 17 |
| RsNMDAR3 | RS004430 | 193 | + | complete | 13 |
| RsNMDAR4 | RS010428 | 47 | - | fragment | 1 |
| RsNMDAR4 | RS010430 | 47 | - | CTE | 1 |
| RsNMDAR4 | RS010426 | 47 | - | NTE | 8 |
| RsNMDAR4 | RS010427 | 47 | - | fragment | 4 |
| RsNMDAR4 | RS010429 | 47 | - | fragment | 1 |
| RsIR8a | RS015433 | 96 | - | complete | 16 |
| RsIR25a | RS008934 | 390 | + | complete | 17 |
| RsIR21a | RS004709 | 20 | - | NTE | 2 |
| RsIR21a | RS004710 | 20 | - | fragment | 4 |
| RsIR21b | RS011973 | 571 | + | complete | 9 |
| RsIR41a1 | RS004249 | 189 | - | complete | 4 |
| RsIR41a2 | RS001436 | 119 | - | NTE | 2 |
| RsIR41a2 | RS001437 | 119 | - | fragment | 1 |
| RsIR41a3 | RS009772 | 430 | + | complete | 5 |
| RsIR41a4 | RS013668 | 72 | + | NTE | 3 |
| RsIR41a4 | RS013667 | 72 | + | fragment | 3 |
| RsIR41a5 | RS004734 | 201 | - | fragment | 1 |
| RsIR41a5 | RS004733 | 201 | - | fragment | 1 |
| RsIR68a | RS000299 | 1 | - | fragment | 1 |
| RsIR68a | RS000298 | 1 | - | complete | 5 |
| RsIR68a | RS000297 | 1 | - | complete | 1 |
| RsIR75a | RS006718 | 281 | + | fragment | 2 |
| RsIR75b | RS009087 | 4 | + | complete | 11 |
| RsIR75c | RS011341 | 527 | - | NTE | 4 |
| RsIR75c | RS011342 | 527 | - | CTE | 5 |
| RsIR75d | RS012791 | 62 | + | fragment | 1 |
| RsIR75e | RS006720 | 281 | + | CTE | 1 |
| RsIR75e | RS006723 | 281 | + | NTE | 1 |
| RsIR75e | RS006722 | 281 | + | fragment | 1 |
| RsIR75e | RS006721 | 281 | + | fragment | 1 |
| RsIR75e | RS006719 | 281 | + | fragment | 1 |
| RsIR75f | RS006724 | 281 | + | complete | 6 |
| RsIR75g | RS012790 | 62 | + | fragment | 1 |
| RsIR75h | RS008223 | 353 | - | complete | 10 |
| RsIR75j | RS011943 | 57 | - | complete | 9 |
| RsIR75k | RS003408 | 160 | - | complete | 9 |

| | | | | | |
|---|---|---|---|---|---|
| RsIR75k | RS003409 | 160 | - | fragment | 1 |
| RsIR75m | RS012788 | 62 | + | complete | 3 |
| RsIR75m | RS012789 | 62 | + | CTE | 4 |
| RsIR75n | RS012793 | 62 | + | fragment | 2 |
| RsIR75n | RS012792 | 62 | + | fragment | 1 |
| RsIR75o | RS012794 | 62 | + | fragment | 1 |
| RsIR75p | RS012795 | 62 | + | CTE | 1 |
| RsIR76b | RS015388 | 955 | + | complete | 4 |
| RsIR76b | RS015389 | 955 | + | CTE | 4 |
| RsIR93a | RS004370 | 19 | - | fragment | 1 |
| RsIR93a | RS004368 | 19 | - | fragment | 5 |
| RsIR93a | RS004369 | 19 | - | fragment | 1 |
| RsIR100 | RS006921 | 293 | - | CTE | 1 |
| RsIR101 | RS000952 | 109 | + | complete | 2 |
| RsIR103 | RS000953 | 109 | + | complete | 3 |
| RsIR104 | RS011634 | 542 | + | complete | 11 |
| RsIR105 | RS009960 | 444 | + | complete | 11 |
| RsIR106 | RS006879 | 29 | + | complete | 8 |
| RsIR107 | RS008138 | 35 | + | complete | 1 |
| RsIR108 | RS007756 | 321 | + | fragment | 3 |
| RsIR109 | RS001336 | 116 | + | complete | 9 |
| RsIR110 | RS008632 | 379 | + | fragment | 1 |
| RsIR111 | RS015446 | 963 | + | fragment | 1 |
| RsIR111 | RS015444 | 963 | + | fragment | 1 |
| RsIR111 | RS015445 | 963 | + | fragment | 1 |
| RsIR112 | RS001346 | 1161 | - | NTE | 2 |
| RsIR113 | RS008629 | 379 | + | NTE | 1 |
| RsIR114 | RS013234 | 663 | + | fragment | 1 |
| RsIR115 | RS009235 | 401 | + | fragment | 1 |
| RsIR116 | RS008630 | 379 | + | CTE | 5 |
| RsIR117 | RS009236 | 401 | + | fragment | 1 |
| RsIR118 | RS014458 | 812 | + | fragment | 1 |
| RsIR119 | RS013409 | 69 | + | fragment | 1 |
| RsIR119 | RS013410 | 69 | + | fragment | 2 |
| RsIR120 | RS002319 | 1356 | - | CTE | 1 |
| RsIR121 | RS013712 | 728 | + | fragment | 1 |
| RsIR122 | RS015442 | 963 | - | fragment | 1 |
| RsIR123 | RS013711 | 728 | + | CTE | 1 |
| RsIR124 | RS008631 | 379 | + | fragment | 1 |
| RsIR125 | RS013235 | 663 | + | fragment | 2 |
| RsIR126 | RS013709 | 728 | - | fragment | 1 |
| RsIR127 | RS005838 | 248 | + | complete | 9 |
| RsIR128 | RS013236 | 663 | + | fragment | 1 |
| RsIR129 | RS001385 | 118 | + | fragment | 1 |
| RsIR130 | RS001384 | 118 | + | fragment | 8 |
| RsIR131 | RS001386 | 118 | + | fragment | 2 |
| RsIR132 | RS001387 | 118 | + | fragment | 1 |

| | | | | | |
|---|---|---|---|---|---|
| RsIR133 | RS003275 | 158 | + | complete | 13 |
| RsIR134 | RS001388 | 118 | + | fragment | 1 |
| RsIR135 | RS006500 | 273 | + | fragment | 1 |
| RsIR136 | RS003792 | 175 | + | complete | 19 |
| RsIR137 | RS007301 | 301 | - | NTE | 1 |
| RsIR138 | RS003198 | 155 | - | fragment | 3 |
| RsIR139 | RS006166 | 26 | - | complete | 25 |
| RsIR140 | RS009364 | 41 | - | complete | 3 |
| RsIR141 | RS015441 | 963 | - | fragment | 2 |
| RsIR142 | RS005465 | 23 | + | complete | 12 |
| RsIR143 | RS012712 | 615 | + | complete | 1 |
| RsIR144 | RS009304 | 41 | + | complete | 9 |
| RsIR145 | RS007402 | 308 | - | CTE | 1 |
| RsIR148 | RS002873 | 147 | - | complete | 2 |
| RsIR148 | RS002874 | 147 | - | fragment | 1 |
| RsIR149 | RS002123 | 130 | + | complete | 4 |
| RsIR149 | RS002121 | 130 | + | CTE | 1 |
| RsIR149 | RS002122 | 130 | + | fragment | 2 |
| RsIR150 | RS006006 | 250 | - | fragment | 2 |
| RsIR159 | RS011158 | 51 | - | complete | 1 |
| RsIR161 | RS009527 | 42 | - | NTE | 1 |
| RsIR162 | RS004299 | 19 | - | NTE | 1 |
| RsIR162 | RS004298 | 19 | - | complete | 1 |
| RsIR164 | RS006947 | 296 | - | NTE | 1 |
| RsIR165 | RS003995 | 18 | - | complete | 1 |
| RsIR169 | RS014584 | 84 | - | complete | 9 |
| RsIR172 | RS004637 | 2 | + | NTE | 1 |
| RsIR182 | RS006698 | 280 | + | complete | 1 |
| RsIR187 | RS003025 | 15 | - | CTE | 1 |
| RsIR192 | RS001765 | 1244 | - | complete | 1 |
| RsIR195 | RS003107 | 1510 | - | CTE | 1 |
| RsIR202 | RS010498 | 48 | - | complete | 1 |
| RsIR203 | RS007486 | 31 | - | complete | 1 |
| RsIR210 | RS013710 | 728 | - | complete | 1 |
| RsIR211 | RS014731 | 864 | - | complete | 2 |
| RsIR215 | RS008190 | 35 | - | complete | 1 |
| RsIR217 | RS008540 | 373 | + | complete | 1 |
| RsIR218 | RS008539 | 373 | - | complete | 1 |

*NTE: initial methionine (M) is missed in the transcript sequence. CTE: stop codon is missed in the transcript sequence.

**Table S18.** Sensory neuron membrane protein (SNMP) genes in *Reticulitermes speratus.*

| Gene | Gene ID | Scaffold | Strand | Sequence | Exons |
|---|---|---|---|---|---|
| RspeSNMP1a | RS007398 | 307 | - | complete | 9 |
| RspeSNMP1b | RS007395 | 307 | - | complete | 9 |
| RspeSNMP1c | RS007396 | 307 | - | fragment | 1 |
| RspeSNMP1c | RS007397 | 307 | - | fragment | 1 |
| RspeSNMP1d | RS007393 | 307 | - | fragment | 2 |
| RspeSNMP1d | RS007394 | 307 | - | fragment | 5 |
| RspeSNMP2 | RS007977 | 339 | - | complete | 9 |

**Table S19.** Chemosensory protein (CSP) genes in *Reticulitermes speratus.*

| Gene | Gene ID | Scaffold | Strand | Sequence* | Exons |
|---|---|---|---|---|---|
| RspeCSP1 | RS000584 | 1000 | - | complete | 2 |
| RspeCSP2 | RS000585 | 1000 | - | complete | 2 |
| RspeCSP3 | RS003292 | 1596 | + | complete | 2 |
| RspeCSP4 | RS003144 | 1536 | - | CTE | 1 |
| RspeCSP5 | RS001446 | 1196 | - | complete | 2 |
| RspeCSP6 | RS010441 | 471 | + | NTE | 1 |
| RspeCSP7 | RS010442 | 471 | + | complete | 2 |
| RspeCSP8 | RS009753 | 43 | + | NTE | 2 |
| RspeCSP9 | RS001447 | 1196 | - | complete | 2 |
| RspeCSP10 | RS002912 | 148 | - | complete | 3 |

*NTE: initial methionine (M) is missed in the transcript sequence. CTE: stop codon is missed in the transcript sequence.

**Table S20.** Biogenic amine- and neuropeptide-related genes in *Reticulitermes speratus*.

| **Gene Name** | **Gene ID** |
|---|---|
| **Biogenic amines (biosynthesis)** | |
| Henna (Phenylalanine hydroxylase) | RS010135 |
| Pale (Tyrosine hydroxylase) | RS010906 |
| Dopa decarboxylase | RS006642 |
| Dopamine N acetyltransferase | RS005696 |
| Tyrosine decarboxylase 2 | RS013885 |
| Tyramine β hydroxylase | RS013347 |
| Tryptophan hydroxylase | RS010115 |
| **Biogenic amines (receptor)** | |
| Dopamine 1-like receptor 1 (Dop1) | RS005814 |
| Dopamine 1-like receptor 2 (Dop2) | RS005701 |
| Dopamine 2-like receptor (Dop3) | RS012926 |
| Dopamine/Ecdysteroid receptor | RS013326 |
| Octopamine-Tyramine receptor | RS000810 |
| Octopamine receptor | RS003890 |
| Octopamine receptor in mushroom bodies | RS006582 |
| Octopamine β2 receptor | RS008926 |
| 5-hydroxytryptamine (serotonin) receptor 1A | RS000074 |
| 5-hydroxytryptamine (serotonin) receptor 1 | RS001623 |
| 5-hydroxytryptamine (serotonin) receptor 2 | RS007168 |
| 5-hydroxytryptamine (serotonin) receptor 2B | RS007166 |
| muscarinic Acetylcholine Receptor, A-type | RS008036 |
| muscarinic Acetylcholine Receptor, B-type | RS008383 |
| Adenosine receptor | RS012570 |
| Histamine-gated chloride channel subunit 1 | RS002296 |
| **Neuropeptides** | |
| Neuropeptide Y receptor | RS006823 |
| CCHamide-1 receptor | RS002615 |
| Adipokinetic hormone | RS002158 |
| Pigment dispersing factor | RS003293 |
| Ion transport peptide | RS003304 |
| Neuroparsin | RS003352 |
| Pyrokinin | RS003469 |
| Neuropeptide F | RS003515 |
| Orcokinin A transcript | RS005362 |
| FMRFamide | RS005650 |
| Vasotocin-neurophysin | RS006779 |
| Neuropeptide F 2 | RS006944 |
| CRF-like Diuretic hormone | RS007172 |

| | |
|---|---|
| RYamide | RS007800 |
| Sulfakinin | RS008491 |
| Partner of Bursicon | RS008861 |
| Bursicon alpha | RS008862 |
| Glycoprotein hormone beta5 (GPB5) | RS009005 |
| Glycoprotein hormone alpha2 (GPA2) | RS009006 |
| Trissin | RS009815 |
| short Neuropeptide F | RS009857 |
| Myosuppressin | RS010591 |
| Tachykinin | RS012013 |
| Leucokinin | RS012022 |
| Crustacean cardioactive peptide | RS012242 |
| Neuropeptide-like precursor 1 | RS012714 |
| Calcitonin-like Diuretic hormone, Diuretic hormone 31 | RS012750 |
| Corazonin | RS013838 |
| Natalisin | RS013901 |
| CCHamide 2 | RS014742 |

**Table S21.** Juvenile hormone (JH)-related genes in *Reticulitermes speratus*.

| Gene Name | Symbol | Gene ID |
|---|---|---|
| **JH biosynthesis genes** | | |
| Acetoacetyl-CoA thiolase | AcoAT | RS011067 |
| 3-Hydroxy-3-Methylglutaryl-CoA synthase 1 | HMGS1 | RS005033 |
| 3-Hydroxy-3-Methylglutaryl-CoA synthase 2 | HMGS2 | RS013349 |
| 3-Hydroxy-3-Methylglutaryl-CoA reductase | HMGR | RS000919 |
| Mevalonate kinase | MK | RS001132 |
| Phosphomevalonate kinase | PK | RS012473 |
| Diphosphomevalonate decarboxylase | DD | RS005846 |
| Isopentenyl-diphosphate σ-isomerase | IPPI | RS012944 |
| Farnesol oxidase | FO | RS008250 |
| Farnesal dehydrogenase 1 | FD1 | RS006558 |
| Farnesal dehydrogenase 2 | FD2 | RS014352 |
| JH acid methyltransferase | JHAMT | RS007861 |
| JH epoxidase | CYP15A1 | RS013787 |
| JH epoxidase homolog | CYP15F1 | RS000985 |
| JH epoxidase homolog | CYP4C7 | RS004449 |
| **JH signaling genes** | | |
| Methoprene-torelant | Met | RS010120 |
| Steroid receptor coactivator | SRC | RS006636 |
| Krüppel homolog-1 | Kr-h1 | RS002081 |
| Broad-Complex | Br-C | RS011102 |
| Ecdysone-induced protein 93 | E93 | RS003976 |
| **Neuropeptide related genes** | | |
| Allatotropin precursor | | RS006223 |
| Allatotropin receptor | | RS005076 |
| Allatostatin precursor | | RS000574 |
| Allatostatin receptor | | RS001538 |
| **JH binding protein genes** | | |
| Hexamerin 1 | Hex1 | RS000846 |
| Hexamerin 2 | Hex2 | RS011205 |
| **JH degradation genes** | | |
| JH esterase | JHE | RS001960 |
| JH esterase | JHE | RS001961 |
| JH esterase | JHE | RS001964 |
| JH esterase | JHE | RS001965 |
| JH esterase | JHE | RS001966 |
| JH esterase | JHE | RS001967 |
| JH esterase | JHE | RS002191 |
| JH esterase | JHE | RS003190 |

| | | |
|---|---|---|
| JH esterase | JHE | RS003673 |
| JH esterase | JHE | RS003910 |
| JH esterase | JHE | RS004129 |
| JH esterase | JHE | RS004712 |
| JH esterase | JHE | RS004713 |
| JH esterase | JHE | RS006008 |
| JH esterase | JHE | RS011642 |
| JH esterase | JHE | RS014537 |
| JH esterase | JHE | RS014538 |
| JH epoxide hydrolase | JHEH | RS011542 |

**Table S22.** Ecdysone-related genes in *Reticulitermes speratus*.

| Gene Name | Gene ID |
|---|---|
| **20E synthesis** | |
| neverland | RS010513 |
| shroud | RS009788 |
| CYP 307a1 (spook) | RS010514 |
| CYP 306a1 (phantom) | RS002862 |
| CYP 302a1 (disembodies) | RS012246 |
| CYP 315a1 (shadow) | RS010451 |
| CYP 314a1 (shade) | RS006327 |
| **20E receptor** | |
| Ecdysone receptor (EcR) | RS006194 |
| ultraspiracle (USP) | RS005985 |
| **20E signaling** | |
| Hormone receptor 3 (HR3) | RS006489 |
| Hormone receptor 4 (HR4) | RS000766 |
| Hormone receptor-like in 38 (HR38) | RS008487 |
| Hormone receptor-like in 39 (HR39) | RS004674 |
| Hormone-receptor-like in 78 (HR78) | RS003557 |
| fushi tarazu transcription factor 1 (FTZ-F1) | RS013785 |
| Ecdysone-induced protein 63 (E63) | RS002747 |
| Ecdysone-induced protein 74 (E74) | RS009331 |
| Ecdysone-induced protein 75 (E75) | RS014319 |
| Ecdysone-induced protein 93 (E93) | RS003976 |
| Ecdysone-induced protein 78 (E78) | RS011677 |

**Table S23.** Insulin signaling genes in *Drosophila melanogaster*, *Zootermepsis nevadensis* and *Reticulitermes speratus*.

| Gene name | Symbol | Function | *D. melanogaster* | *Z. nevadensis* | *R. speratus* |
|---|---|---|---|---|---|
| Insulin-like peptide 1 | Ilp1 | Ligand | FBgn0044051 | Znev_05166, Znev_05167, Znev_07008, Znev_07935, Znev_07936 | RS000535, RS000536, RS002145, RS008597, RS008598 |
| Insulin-like peptide 2 | Ilp2 | Ligand | FBgn0036046 | | |
| Insulin-like peptide 3 | Ilp3 | Ligand | FBgn0044050 | | |
| Insulin-like peptide 4 | Ilp4 | Ligand | FBgn0044049 | | |
| Insulin-like peptide 5 | Ilp5 | Ligand | FBgn0044048 | | |
| Insulin-like peptide 6 | Ilp6 | Ligand | FBgn0044047 | | |
| Insulin-like peptide 7 | Ilp7 | Ligand | FBgn0044046 | | |
| Insulin-like peptide 8 | Ilp8 | Ligand | FBgn0036690 | | |
| Insulin-like receptor | InR | Receptor | FBgn0283499 | Znev_02684, Znev_02685, Znev_16736 | RS000922, RS007018, RS007019 |
| chico | chico | Receptor | FBgn0024248 | Znev_13974 | RS000780 |
| Lnk | Lnk | Receptor | FBgn0028717 | Znev_12363 | RS006416 |
| Phosphatidylinositol 3-kinase 92E | Pi3K92E | Downstream of insulin receptor | FBgn0015279 | Znev_01400 | RS000592 |
| Pi3K21B | Pi3K21B | Downstream of insulin receptor | FBgn0020622 | Znev_04386, Znev_11082 | RS004930, RS006442 |
| Phosphoinositide-dependent kinase 1 | Pdk1 | Downstream of insulin receptor | FBgn0020386 | Znev_04009 | RS010411 |
| Akt1 | Akt1 | Downstream of insulin receptor | FBgn0010379 | Znev_04339 | RS000520 |
| Phosphatase and tensin homolog | Pten | Downstream of insulin receptor | FBgn0026379 | Znev_16636 | RS011450 |
| Ribosomal protein S6 kinase | S6k | Downstream of insulin receptor | FBgn0283472 | Znev_12092 | RS003806 |
| Target of rapamycin | Tor | Downstream of insulin receptor | FBgn0021796 | Znev_11128 | RS002594 |
| Tsc1 | Tsc1 | Downstream of insulin receptor | FBgn0026317 | Znev_11179 | RS004848 |
| gigas | gig | Downstream of insulin receptor | FBgn0005198 | Znev_09179 | RS006709 |
| Ribosomal protein S6 | RpS6 | Downstream of insulin receptor | FBgn0261592 | Znev_07260 | RS014227 |
| Thor | Thor | Downstream of insulin receptor | FBgn0261560 | Znev_16639 | RS011442 |
| forkhead box, sub-group O | foxo | Downstream of insulin receptor | FBgn0038197 | Znev_14322 | RS013726 |
| Tif-IA | Tif-IA | Downstream of insulin receptor | FBgn0032988 | Znev_17679 | RS000398 |
| shaggy | sgg | Downstream of insulin receptor | FBgn0003371 | - | RS007030 |
| Ras homolog enriched in brain | Rheb | Downstream of insulin receptor | FBgn0041191 | Znev_02068 | RS001480 |

| | | | | | |
|---|---|---|---|---|---|
| widerborst | wdb | Downstream of insulin receptor | FBgn0027492 | Znev_18447 | RS014632 |
| raptor | raptor | Downstream of insulin receptor | FBgn0029840 | Znev_09185 | RS007403 |
| Myc | Myc | Downstream of insulin receptor | FBgn0262656 | Znev_03784 | RS012525 |
| Ras oncogene at 85D | Ras85D (Ras1) | Ras signal | FBgn0003205 | Znev_07076, Znev_07077, Znev_14471 | RS000933, RS003738, RS013615 |
| happyhour | hppy | Ras signal | FBgn0263395 | Znev_03645 | RS015403 |
| rolled | rl | Ras signal | FBgn0003256 | Znev_00943 | RS002853 |
| Raf oncogene | Raf | Ras signal | FBgn0003079 | - | RS009493 |
| Downstream of raf1 | Dsor1 | Ras signal | FBgn0010269 | Znev_04859 | RS012464 |
| Ecdysone-inducible gene L2 | ImpL2 | Binding to ILP | FBgn0001257 | - | - |
| convoluted | conv | Binding to ILP | FBgn0261269 | Znev_01328 | RS008068 |
| steppke | step | other | FBgn0086779 | Znev_11787 | RS014498 |
| melted | melt | other | FBgn0023001 | Znev_04452 | RS000118 |

**Table S24.** Toolkit genes involved in wing formation in *Drosophila melanogaster* and *Reticulitemes speratus*.

| **Gene name** | **Gene symbol** | ***D. melanogaster*** | ***R. speratus*** | **Expression Differences among castes (FDR)** | |
|---|---|---|---|---|---|
| | | | | **Head** | **Thorax + abdomen** |
| achaete | ac | FBgn0000022 | RS001853 | 6.38E-13* | 0.0002* |
| scute | sc | FBgn0004170 | | | |
| asense | ase | FBgn0000137 | RS001854 | 0.719 | 0.261 |
| abdominal A | abd-A | FBgn0000014 | RS009109 | 0.536 | 0.423 |
| Antennapedia | Antp | FBgn0260642 | RS009104 | 0.150 | 0.052 |
| apterous | ap | FBgn0267978 | RS004477 | 1.25E-06* | 0.0099* |
| bifid | bi | FBgn0000179 | RS006410 | 0.731 | 0.008* |
| brinker | brk | FBgn0024250 | RS014094 | 0.011* | 0.023* |
| cubitus interruptus | ci | FBgn0004859 | RS009075 | 5.24E-06* | 0.909 |
| cut | cut | FBgn0004198 | RS000064 | 1.04E-05* | 0.005* |
| Daughters against dpp | dad | FBgn0020493 | RS009849 | 0.766 | 0.007* |
| Distal-less | dll | FBgn0000157 | RS010450 | 0.857 | 0.375 |
| decapentaplegic | dpp | FBgn0000490 | RS012434 | 1.40E-08* | 0.0001* |
| engrailed | en | FBgn0000577 | RS005170 | 0.517 | 0.862 |
| escargot | esg | FBgn0001981 | RS003859 | 0.669 | 0.039* |
| snail | sna | FBgn0003448 | | | |
| worniu | wor | FBgn0001983 | | | |
| extradenticle | exd | FBgn0000611 | RS014099 | 0.358 | 0.326 |
| hedgehog | hh | FBgn0004644 | RS014625 | 0.101 | 0.525 |
| homothorax | hth | FBgn0001235 | RS001474 | 0.068 | 0.916 |
| Notch | N | FBgn0004647 | RS013346 | 0.083 | 0.513 |
| nubbin | nub | FBgn0085424 | RS011952 | 0.594 | 0.968 |
| patched | ptc | FBgn0003892 | RS008136 | 0.907 | 0.028* |
| spalt major | sal | FBgn0261648 | RS013395 | 1.07E-33* | 0.009* |
| Sex combs reduced | scr | FBgn0003339 | RS009101 | 0.397 | 3.41E-07* |
| scalloped | sd | FBgn0003345 | RS008357 | 0.727 | 0.836 |
| Serrate | ser | FBgn0004197 | RS005737 | 0.001* | 0.500 |
| spitz | spi | FBgn0005672 | RS004578,<br>RS007465 | 1.29E-62*<br>0.145 | 0.009*<br>0.001* |
| Keren | Krn | FBgn0052179 | | | |
| vein | vn | FBgn0003984 | | | |
| gurken | grk | FBgn0001137 | | | |

| | | | | | |
|---|---|---|---|---|---|
| blistered | bs | FBgn0004101 | RS014165 | 4.05E-07* | 0.071 |
| teashirt | tsh | FBgn0003866 | RS003447 | 7.02E-09* | 0.018* |
| tiptop | tio | FBgn0028979 | | | |
| Ultrabithorax | Ubx | FBgn0003944 | RS009105 | 0.447 | 0.092 |
| vestigial | vg | FBgn0003975 | RS011568 | 0.0001* | 0.014* |
| ventral veins lacking | vvl | FBgn0086680 | RS011806 | 0.163 | 0.865 |
| wingless | wg | FBgn0284084 | RS015426 | 0.023* | 0.087 |

*FDR < 0.05

**Table S25.** Immune-related genes in *Reticulitermes speratus*.

| Family | Subfamily | Gene name | Gene ID |
|---|---|---|---|
| Antimicrobial peptide | crustin-like protein | | RS006883 |
| Antimicrobial peptide | defensin | | RS002487 |
| Antimicrobial peptide | defensin | | RS100003 |
| Antimicrobial peptide | locustin-like protein | | RS008268 |
| Antimicrobial peptide | prolixicin | | RS000201 |
| Antimicrobial peptide | termicin | | RS006953 |
| Antimicrobial peptide | thaumatin | | RS015368 |
| Antimicrobial peptide | thaumatin | | RS015369 |
| Autophagy | Autophgy-related | | RS000130 |
| Autophagy | Autophgy-related | | RS001459 |
| Autophagy | Autophgy-related | | RS002992 |
| Autophagy | Autophgy-related | | RS004579 |
| Autophagy | Autophgy-related | | RS006202 |
| Autophagy | Autophgy-related | | RS006853 |
| Autophagy | Autophgy-related | | RS006855 |
| Autophagy | Autophgy-related | | RS007186 |
| Autophagy | Autophgy-related | | RS007411 |
| Autophagy | Autophgy-related | | RS008423 |
| Autophagy | Autophgy-related | | RS008527 |
| Autophagy | Autophgy-related | | RS011626 |
| Autophagy | Autophgy-related | | RS011915 |
| Autophagy | Autophgy-related | | RS012458 |
| Autophagy | Autophgy-related | | RS014090 |
| Autophagy | Autophgy-related | | RS014951 |
| Autophagy | | Buffy | RS006694 |
| Autophagy | | Target of rapamycin | RS002594 |
| 1,3-beta-D glucan binding protein | | | RS002847 |
| 1,3-beta-D glucan binding protein | | | RS002848 |
| 1,3-beta-D glucan binding protein | | | RS004742 |
| 1,3-beta-D glucan binding protein | | | RS100018 |
| 1,3-beta-D glucan binding protein | | | RS100020 |
| Caspase | | | RS003133 |
| Caspase | | | RS003489 |

| | | | |
|---|---|---|---|
| Caspase | | | RS005313 |
| Caspase | | | RS008854 |
| Caspase | | | RS012279 |
| Caspase | | | RS014033 |
| Caspase | | | RS014035 |
| Caspase | | | RS014259 |
| Caspase activator | | | RS012758 |
| Catalase | | | RS001412 |
| Catalase | | | RS001654 |
| Catalase | | | RS001671 |
| Catalase | | | RS001672 |
| CLIP-domain serine protease | | | RS004304 |
| CLIP-domain serine protease | | | RS004305 |
| CLIP-domain serine protease | | | RS004306 |
| CLIP-domain serine protease | | | RS004307 |
| CLIP-domain serine protease | | | RS007082 |
| CLIP-domain serine protease | | | RS010297 |
| CLIP-domain serine protease | | | RS010298 |
| CLIP-domain serine protease | | | RS010333 |
| CLIP-domain serine protease | | | RS010976 |
| CLIP-domain serine protease | | | RS010995 |
| CLIP-domain serine protease | | | RS011937 |
| CLIP-domain serine protease | | | RS012021 |
| CLIP-domain serine protease | | | RS012936 |
| CLIP-domain serine protease | | | RS013065 |
| CLIP-domain serine protease | | | RS013066 |
| CLIP-domain serine protease | | | RS013067 |
| CLIP-domain serine protease | | | RS013068 |
| CLIP-domain serine protease | | | RS013070 |
| CLIP-domain serine protease | | | RS013766 |
| C-type lectine | | | RS000351 |
| C-type lectine | | | RS001294 |
| C-type lectine | | | RS001855 |
| C-type lectine | | | RS003812 |
| C-type lectine | | | RS005053 |

| | | | |
|---|---|---|---|
| C-type lectine | | | RS006091 |
| C-type lectine | | | RS006677 |
| C-type lectine | | | RS007270 |
| C-type lectine | | | RS007745 |
| C-type lectine | | | RS007746 |
| C-type lectine | | | RS008198 |
| C-type lectine | | | RS008199 |
| C-type lectine | | | RS008200 |
| C-type lectine | | | RS008201 |
| C-type lectine | | | RS008204 |
| C-type lectine | | | RS010439 |
| C-type lectine | | | RS010966 |
| C-type lectine | | | RS011333 |
| C-type lectine | | | RS011334 |
| C-type lectine | | | RS011520 |
| C-type lectine | | | RS011567 |
| C-type lectine | | | RS011914 |
| C-type lectine | | | RS013652 |
| C-type lectine | | | RS013842 |
| C-type lectine | | | RS014902 |
| C-type lectine | | | RS015474 |
| Fibrinogen-related protein | | | RS000785 |
| Fibrinogen-related protein | | | RS006501 |
| Fibrinogen-related protein | | | RS008150 |
| Fibrinogen-related protein | | | RS011288 |
| Galactoside-binding lectin | | | RS005311 |
| Galactoside-binding lectin | | | RS006436 |
| Galactoside-binding lectin | | | RS013017 |
| Galactoside-binding lectin | | | RS013896 |
| Galactoside-binding lectin | | | RS013897 |
| Inhibitor of apoptosis | | BIR repeat containing ubiquitin-conjugating enzyme | RS006599 |
| Inhibitor of apoptosis | | Death-associated inhibitor of apoptosis 1 | RS011068 |
| Inhibitor of apoptosis | | Death-associated inhibitor of apoptosis 2 | RS011069 |
| Inhibitor of apoptosis | | Deterin | RS011623 |

| | | | |
|---|---|---|---|
| IMD pathway member | | caspar | RS003316 |
| IMD pathway member | | Fas-associated death domain ortholog | RS010104 |
| IMD pathway member | | immune deficiency | RS008532 |
| IMD pathway member | | immune response deficient 5 | RS010217 |
| IMD pathway member | | kenny | RS007928 |
| IMD pathway member | | poor Imd response upon knock-in | RS006682 |
| IMD pathway member | | TAK1-associated binding protein 2 | RS007289 |
| IMD pathway member | | TGF-β activated kinase 1 | RS002792 |
| JAK/STAT pathway member | | domeless | RS014980 |
| JAK/STAT pathway member | | hopscotch | RS002898 |
| JAK/STAT pathway member | | Signal-transducer and activator of transcription protein at 92E | RS003724 |
| Lysozyme | c-type lysozyme | | RS000427 |
| Lysozyme | c-type lysozyme | | RS002400 |
| Lysozyme | c-type lysozyme | | RS003406 |
| Lysozyme | c-type lysozyme | | RS008613 |
| Lysozyme | c-type lysozyme | | RS014698 |
| Lysozyme | c-type lysozyme | | RS100001 |
| Lysozyme | c-type lysozyme | | RS100002 |
| Lysozyme | c-type lysozyme | | RS100004 |
| Lysozyme | c-type lysozyme | | RS100021 |
| Lysozyme | c-type lysozyme | | RS100022 |
| Lysozyme | c-type lysozyme | | RS100023 |
| Lysozyme | c-type lysozyme | | RS100024 |
| Lysozyme | c-type lysozyme | | RS100025 |
| Lysozyme | c-type lysozyme | | RS100026 |
| Lysozyme | i-type lysozyme | | RS006054 |
| Lysozyme | i-type lysozyme | | RS008547 |
| Lysozyme | i-type lysozyme | | RS015579 |
| MD2-like receptor | | | RS013029 |
| MD2-like receptor | | | RS013030 |
| MD2-like receptor | | | RS013031 |
| MD2-like receptor | | | RS013032 |
| MD2-like receptor | | | RS015155 |
| Peptidoglycan recognition protein | | | RS005405 |

| | | | |
|---|---|---|---|
| Peptidoglycan recognition protein | | | RS006292 |
| Peptidoglycan recognition protein | | | RS010040 |
| Peptidoglycan recognition protein | | | RS013012 |
| Peptidoglycan recognition protein | | | RS013013 |
| Peptidoglycan recognition protein | | | RS100019 |
| Prophenoloxidase | | | RS008762 |
| Peroxidase | | | RS002951 |
| Peroxidase | | | RS002966 |
| Peroxidase | | | RS004201 |
| Peroxidase | | | RS004202 |
| Peroxidase | | | RS004203 |
| Peroxidase | | | RS004289 |
| Peroxidase | | | RS004290 |
| Peroxidase | | | RS006166 |
| Peroxidase | | | RS009222 |
| Peroxidase | | | RS009427 |
| Peroxidase | | | RS012128 |
| Peroxidase | | | RS012299 |
| Peroxidase | | | RS013770 |
| Peroxidase | | | RS014619 |
| Relish-like protein | | dorsal | RS002016 |
| Relish-like protein | | Relish | RS007470 |
| Scavenger receptor | | | RS001883 |
| Scavenger receptor | | | RS001884 |
| Scavenger receptor | | | RS001885 |
| Scavenger receptor | | | RS001886 |
| Scavenger receptor | | | RS001887 |
| Scavenger receptor | | | RS002546 |
| Scavenger receptor | | | RS002548 |
| Scavenger receptor | | | RS002549 |
| Scavenger receptor | | | RS002550 |
| Scavenger receptor | | | RS003004 |
| Scavenger receptor | | | RS003767 |
| Scavenger receptor | | | RS005025 |
| Scavenger receptor | | | RS007394 |

| | | | |
|---|---|---|---|
| Scavenger receptor | | | RS007395 |
| Scavenger receptor | | | RS007398 |
| Scavenger receptor | | | RS007853 |
| Scavenger receptor | | | RS007977 |
| Scavenger receptor | | | RS012012 |
| Scavenger receptor | | | RS012281 |
| Scavenger receptor | | | RS013965 |
| Scavenger receptor | | | RS014587 |
| Superoxide dismutatse | | | RS001583 |
| Superoxide dismutatse | | | RS001584 |
| Superoxide dismutatse | | | RS002757 |
| Superoxide dismutatse | | | RS004133 |
| Superoxide dismutatse | | | RS009320 |
| Superoxide dismutatse | | | RS013005 |
| Spaetzle-like protein | | | RS000600 |
| Spaetzle-like protein | | | RS000979 |
| Spaetzle-like protein | | | RS003253 |
| Spaetzle-like protein | | | RS004223 |
| Spaetzle-like protein | | | RS011245 |
| Spaetzle-like protein | | | RS011246 |
| Spaetzle-like protein | | | RS011463 |
| Spaetzle-like protein | | | RS011594 |
| Serine protease inhibitor | | | RS000470 |
| Serine protease inhibitor | | | RS001220 |
| Serine protease inhibitor | | | RS004075 |
| Serine protease inhibitor | | | RS005022 |
| Serine protease inhibitor | | | RS005471 |
| Serine protease inhibitor | | | RS006339 |
| Serine protease inhibitor | | | RS008737 |
| Serine protease inhibitor | | | RS011058 |
| Serine protease inhibitor | | | RS013650 |
| Serine protease inhibitor | | | RS014220 |
| Serine protease inhibitor | | | RS014221 |
| Serine protease inhibitor | | | RS014444 |
| Small regulatory RNA pathway | | Argonaute 1 | RS005621 |

| | | | |
|---|---|---|---|
| Small regulatory RNA pathway | | Argonaute 2a | RS010079 |
| Small regulatory RNA pathway | | Argonaute 2b | RS003821 |
| Small regulatory RNA pathway | | Argonaute 3 | RS003744 |
| Small regulatory RNA pathway | | armitage | RS006369 |
| Small regulatory RNA pathway | | aubergine | RS004296 |
| Small regulatory RNA pathway | | Dicer 1 | RS010489 |
| Small regulatory RNA pathway | | Dicer 2 | RS013700 |
| Small regulatory RNA pathway | | drosha | RS007949 |
| Small regulatory RNA pathway | | loquacious | RS001759 |
| Small regulatory RNA pathway | | partner of drosha | RS011750 |
| Small regulatory RNA pathway | | piwi 1 | RS002624 |
| Small regulatory RNA pathway | | piwi 2 | RS011008 |
| Small regulatory RNA pathway | | R2D2 | RS004300 |
| Small regulatory RNA pathway | Rm62 | | RS001900 |
| Small regulatory RNA pathway | Rm62 | | RS001901 |
| Small regulatory RNA pathway | Rm62 | | RS001902 |
| Small regulatory RNA pathway | Rm62 | | RS001905 |
| Small regulatory RNA pathway | Rm62 | | RS001906 |
| Small regulatory RNA pathway | Rm62 | | RS002802 |
| Small regulatory RNA pathway | Rm62 | | RS002903 |
| Small regulatory RNA pathway | Rm62 | | RS005517 |
| Small regulatory RNA pathway | Rm62 | | RS006077 |
| Small regulatory RNA pathway | Rm62 | | RS006940 |
| Small regulatory RNA pathway | Rm62 | | RS012401 |
| Small regulatory RNA pathway | Rm62 | | RS013373 |
| Small regulatory RNA pathway | | spindle E | RS010943 |
| Small regulatory RNA pathway | | Tudor staphylococcal nuclease | RS004901 |
| Small regulatory RNA pathway | | vasa intronic gene | RS011955 |
| Thio-ester containing protein | | | RS000697 |
| Thio-ester containing protein | | | RS010197 |
| Thio-ester containing protein | | | RS013791 |
| Thio-ester containing protein | | | RS014232 |
| Toll pathway | | cactus | RS009203 |
| Toll pathway | | Myd88 | RS011504 |
| Toll pathway | | pelle | RS014381 |

| | | | |
|---|---|---|---|
| Toll pathway | | TNF-receptor-associated factor-like | RS011260 |
| Toll pathway | | tube | RS013249 |
| Toll pathway | Toll receptor | | RS002528 |
| Toll pathway | Toll receptor | | RS005466 |
| Toll pathway | Toll receptor | | RS005470 |
| Toll pathway | Toll receptor | | RS005472 |
| Toll pathway | Toll receptor | | RS005474 |
| Toll pathway | Toll receptor | | RS007296 |
| Toll pathway | Toll receptor | | RS007975 |
| Toll pathway | Toll receptor | | RS008286 |

**Table S26.** Insecticide-target genes in *Reticulitermes speratus* and their homologs in *Drosophila melanogaster*.

| Gene name | *D. melanogaster* | *R. speratus* |
|---|---|---|
| **Ion channels** | | |
| Acetylcholine esterase | FBgn0000024 | RS013479, RS009432 |
| nicotinic Acetylcholine Receptor α1 | FBgn0000036 | RS002929 |
| nicotinic Acetylcholine Receptor α2 | FBgn0000039 | RS002930 |
| nicotinic Acetylcholine Receptor α3 | FBgn0015519 | RS001773 |
| nicotinic Acetylcholine Receptor α4 | FBgn0266347 | RS001774 |
| nicotinic Acetylcholine Receptor α5 | FBgn0028875 | RS003729 |
| nicotinic Acetylcholine Receptor α6 | FBgn0032151 | RS014310 |
| nicotinic Acetylcholine Receptor α7 | FBgn0086778 | RS003145 |
| nicotinic Acetylcholine Receptor α8 | - | RS015063 |
| nicotinic Acetylcholine Receptor α9 | - | RS015059 |
| nicotinic Acetylcholine Receptor α10 | - | RS015059 |
| nicotinic Acetylcholine Receptor β1 | FBgn0000038 | RS003147 |
| nicotinic Acetylcholine Receptor β2 | FBgn0004118 | RS002931 |
| nicotinic Acetylcholine Receptor β3 | FBgn0031261 | RS012432 |
| GABA type A receptor | FBgn0004244 | RS010905 |
| GABA type A receptor | FBgn0030707 | RS013195 |
| Ligand-gated chloride channel | FBgn0010240 | RS013193 |
| Glycine receptor | FBgn0001134 | RS007641 |
| Histamine-gated chloride channel subunit | FBgn0037950 | RS002296 |
| Histamine-gated chloride channel subunit | FBgn0003011 | - |
| Voltage-gated chloride channel subunit | FBgn0051116 | RS008419 |
| Voltage-gated chloride channel subunit | FBgn0033755 | RS003825 |
| Voltage-gated chloride channel subunit | FBgn0036566 | RS008614 |

| | | |
|---|---|---|
| Voltage-gated chloride channel subunit | FBgn0038721 | RS011813 |
| Voltage-gated sodium channel subunit | FBgn0264255 | RS008214 |
| Voltage-gated sodium channel subunit | FBgn0085434 | RS015165 |
| **Chitin synthesis** | | |
| Glycogen phosphorylase | FBgn0004507 | RS013552, RS006448, RS006452 |
| Trehalase | FBgn0003748 | RS004154, RS005093, RS013303 |
| Hexokinase A | FBgn0001186 | RS007194 |
| Hexokinase C | FBgn0001187 | no homolog |
| Phosphoglucose isomerase | FBgn0003074 | RS012390 |
| Glutamine:fructose-6-phosphate aminotransferase | FBgn0027341 | RS005235 |
| Glutamine:fructose-7-phosphate aminotransferase | FBgn0039580 | - |
| Glucosamine-6-phosphate N-acetyltransferase | FBgn0039690 | RS004411 |
| phosphoglucose mutase | FBgn0003076 | RS009458 |
| mummy | FBgn0259749 | RS010490 |
| Chitin synthase | FBgn0001311 | RS006626 |
| Chitin synthase | FBgn0029091 | RS006625 |
| **Muscle-related** | | |
| Ryanodine receptor | FBgn0011286 | RS000001, RS015395 |
| sarcoplasmic/endoplasmic reticulum Ca2+ ATPase | FBgn0263006 | RS000365 |
| myosin light chain kinase | FBgn0265045 | RS008592, RS012598, RS012599, RS014874, RS014875 |
| Calcium/Calmodulin-dependent protein kinase | FBgn0005666 | RS014950 |
| Calmodulin | FBgn0000253 | RS014134, RS013623 |
| Calcium/calmodulin-dependent protein kinase I | FBgn0016126 | RS010237 |
| Calcium/calmodulin-dependent protein kinase II | FBgn0264607 | RS005236 |
| Calmodulin-binding transcription activator | FBgn0259234 | RS000076 |
| Ras-related protein interacting with calmodulin | FBgn0265605 | RS000452 |

| | | |
|---|---|---|
| Trimeric intracellular cation chanel | FBgn0030745 | RS011850 |
| Calmodulin-binding protein related to a Rab3 GDP/GTP exchange protein | FBgn0025864 | RS014554 |
| **Aquaporin family** | | |
| aquaporin | FBgn0015872 | RS000720 |
| aquaporin | FBgn0000180 | RS005584, RS005585, RS005586 |
| aquaporin | FBgn0033807 | RS011988 |
| aquaporin | FBgn0034883 | RS013221 |
| aquaporin | FBgn0034884 | - |
| aquaporin | FBgn0034885 | - |
| aquaporin | FBgn0034882 | - |
| aquaporin | FBgn0033635 | RS000719 |

**Table S27.** Cytochrome P450s (CYP), Glutathione S-transferases (GST) and Carboxylesterases (CCE) genes in *Reticulitermes speratus*, *Zootermopsis nevadensis* and *Macrotermes natalensis*.

| Gene name | *R. speratus* | *Z. nevadensis* | *M. natalensis* |
|---|---|---|---|
| CYP | RS000202, RS000210, RS000211, RS000212, RS000444, RS000445, RS000465, RS000466, RS000467, RS000812, RS000815, RS000859, RS000980, RS000981, RS000982, RS000983, RS000984, RS000985, RS000986, RS000987, RS001122, RS001799, RS001847, RS002862, RS002863, RS002871, RS002872, RS002910, RS003017, RS003149, RS003150, RS003151, RS003152, RS003153, RS003154, RS003155, RS003156, RS003157, RS003158, RS003160, RS003206, RS003207, RS003672, RS003688, RS003709, RS003788, RS004073, RS004422, RS004428, RS004445, RS004446, RS004448, RS004449, RS004450, RS005356, RS005658, RS005672, RS006327, RS007757, RS008449, RS008850, RS008851, RS008852, RS009217, RS010162, RS010163, RS010164, RS010165, RS010451, RS010514, RS010600, RS010601, RS010602, RS010608, RS010609, RS010610, RS010611, RS010975, RS011306, RS011875, RS011957, RS012246, RS012921, RS013159, RS013250, RS013635, RS013647, RS013663, RS013664, RS013665, RS013784, RS013787, RS013788, RS013835, RS014290, RS014291, RS014292, RS014413, RS014414, RS014484, RS014485, RS014624, RS014933, RS015215, RS015219, RS015221 | Znev_00012, Znev_00957, Znev_00958, Znev_01139, Znev_01838, Znev_01867, Znev_01868, Znev_02456, Znev_02808, Znev_03004, Znev_03222, Znev_04232, Znev_04417, Znev_04827, Znev_04985, Znev_05339, Znev_05340, Znev_05390, Znev_05391, Znev_05398, Znev_06057, Znev_06128, Znev_06541, Znev_06629, Znev_07037, Znev_08570, Znev_08701, Znev_08930, Znev_09012, Znev_09132, Znev_09277, Znev_09478, Znev_09480, Znev_09481, Znev_11665, Znev_12901, Znev_12912, Znev_13255, Znev_13889, Znev_13890, Znev_13891, Znev_13892, Znev_13893, Znev_14063, Znev_14143, Znev_14286, Znev_14287, Znev_14299, Znev_14300, Znev_14301, Znev_14302, Znev_14590, Znev_14632, Znev_14659, Znev_14677, Znev_14802, Znev_14833, Znev_15638, Znev_15869, Znev_15870, Znev_16120, Znev_16125, Znev_16153, Znev_16218, Znev_16223, Znev_16398, Znev_16438, Znev_16439, Znev_16771, Znev_17183, Znev_18486, Znev_18620, Znev_18647 | MN000227, MN000228, MN000347, MN001136, MN001137, MN001432, MN002329, MN002525, MN002815, MN002965, MN003123, MN003207, MN003446, MN003447, MN003965, MN004138, MN004139, MN004141, MN004142, MN004409, MN004466, MN004803, MN004999, MN005205, MN005352, MN005353, MN005542, MN005559, MN005560, MN005562, MN005695, MN005696, MN005697, MN005770, MN005961, MN006039, MN006125, MN006126, MN006243, MN006414, MN006416, MN006418, MN006598, MN006609, MN006690, MN006691, MN006758, MN007379, MN007380, MN007381, MN007382, MN007383, MN007384, MN007385, MN007386, MN007388, MN007624, MN007639, MN007913, MN007915, MN007918, MN008303, MN008754, MN008793, MN008794, MN008892, MN008893, MN009093, MN009118, MN009119, MN009543, MN009788, MN009789, MN010562, MN010563, MN010564, MN011715, MN011716, MN011746, MN011747, MN011748, MN012022, MN012023 |
| GST | RS000232, RS001168, RS001200, RS003134, RS003137, RS005177, RS005799, RS007488, RS007489, RS007490, RS009342, RS009343, RS009344, RS009865, RS010039, RS011657, RS011756, RS013048, RS014031, RS015157 | Znev_00124, Znev_00817, Znev_00818, Znev_04286, Znev_04467, Znev_04468, Znev_04470, Znev_08437, Znev_11969, Znev_12236, Znev_13081, Znev_13082, Znev_14234, Znev_14305, Znev_15569, Znev_15570, Znev_15795, Znev_15978 | MN000171, MN000172, MN000173, MN000543, MN000546, MN001107, MN001778, MN002944, MN004973, MN005428, MN005538, MN006379, MN006654, MN009351, MN010121, MN010281, MN011310 |
| CCE | RS001422, RS001423, RS001960, RS001961, RS001964, RS001965, RS001966, RS001967, RS002191, RS003673, RS003894, RS003895, RS003910, RS004129, RS004711, RS004712, RS004713, RS006008, RS006009, RS006010, RS006537, RS006938, RS007152, RS007153, RS007154, RS007155, RS007631, RS007632, RS007668, RS007829, RS009273, RS009432, RS010553, RS011141, RS011142, RS011143, RS011477, RS011642, RS012006, RS013406, RS013479, RS014537, RS014538 | Znev_00356, Znev_00985, Znev_01464, Znev_01490, Znev_01491, Znev_01695, Znev_01720, Znev_01734, Znev_01735, Znev_01778, Znev_01779, Znev_01844, Znev_02218, Znev_02219, Znev_02223, Znev_02224, Znev_02226, Znev_02227, Znev_02228, Znev_05318, Znev_07005, Znev_07007, Znev_07097, Znev_10214, Znev_12048, Znev_12702, Znev_12703, Znev_12704, Znev_12774, Znev_12960, Znev_13209, Znev_14317, Znev_14604, Znev_14951, Znev_15823, Znev_15907, Znev_17070, Znev_17401, Znev_17972, Znev_18481, Znev_18523 | MN000642, MN001220, MN001683, MN002124, MN002125, MN003389, MN003390, MN004198, MN005717, MN005718, MN006035, MN006244, MN006980, MN007226, MN008137, MN008387, MN010399, MN010706, MN011531, MN011532, MN011533, MN011534, MN011536, MN011537, MN011538, MN011539, MN011540, MN011541, MN011670, MN011672, MN011673, MN011731 |

**Table S28.** Gene numbers of Cytochrome P450s (CYP), Glutathione S-transferases (GST) and Carboxylesterases (CCE) in 6 insect species.

| | *Drosophila melanogaster** | *Periplaneta americana* | *Cryptocercus punctulatus* | *Zootermopsis nevadensis* | *Reticulitermes speratus* | *Macrotermes natalensis* |
|---|---|---|---|---|---|---|
| CYP | 89 | 264 | 124 | 75 | 106 | 94 |
| GST | 39 | 55 | 26 | 18 | 20 | 17 |
| CCE | 35 | 108 | 66 | 41 | 43 | 32 |

*Gene numbers are referred to Ranson et al. (2002).

**Dataset S1 (separate file).** Caste-biased genes [Excel] (SI_Data_1_caste-biased_genes.xlsx).
