## Supplementary Datasets S1 for "Genomic and transcriptomic analyses of the subterranean termite *Reticulitermes speratus:* gene duplication facilitates social evolution"

SI_Dataset_1
Caste-biased genes: Reproductive (head)

| Gene ID | Head Reproductive v.s. Soldier | | | Head Reproductive v.s. Worker | | | Head Soldier v.s. Worker | | | Gene annotation |
|---|---|---|---|---|---|---|---|---|---|---|
| | logFC | logCPM | FDR | logFC | logCPM | FDR | logFC | logCPM | FDR | |
| RS000444 | -2.8 | 2.6 | 2.5E-08 | -2.4 | 2.6 | 3.7E-07 | 0.4 | 2.6 | 7.2E-01 | cytochrome p450 |
| RS000445 | -1.9 | 5.1 | 6.1E-20 | -2.5 | 5.1 | 2.3E-32 | -0.6 | 5.1 | 2.6E-02 | cytochrome p450 |
| RS000610 | -7.8 | 6.5 | 9.3E-25 | -8.9 | 6.5 | 1.5E-27 | -1.0 | 6.5 | 4.2E-01 | vitellogenin |
| RS000616 | -8.1 | 5.0 | 1.4E-25 | -6.8 | 5.0 | 2.7E-20 | 1.3 | 5.0 | 3.7E-01 | vitellogenin |
| RS001422 | -3.0 | 5.8 | 7.9E-18 | -4.0 | 5.8 | 1.5E-27 | -1.0 | 5.8 | 1.5E-02 | |
| RS001423 | -3.1 | 3.0 | 7.3E-11 | -4.5 | 3.0 | 1.8E-18 | -1.4 | 3.0 | 2.7E-02 | venom carboxylesterase-6-like |
| RS001606 | -3.3 | -0.3 | 3.1E-04 | -3.5 | -0.3 | 1.5E-04 | -0.2 | -0.3 | 9.3E-01 | 101 kda malaria antigen-like |
| RS001654 | -1.3 | 5.5 | 5.1E-04 | -1.8 | 5.5 | 3.0E-06 | -0.5 | 5.5 | 3.5E-01 | catalase |
| RS001804 | -3.7 | 5.0 | 1.5E-65 | -1.8 | 5.0 | 3.0E-21 | 1.9 | 5.0 | 2.7E-16 | udp-glucuronosyltransferase |
| RS001884 | -1.6 | 6.0 | 8.7E-23 | -1.5 | 6.0 | 7.8E-21 | 0.1 | 6.0 | 8.4E-01 | protein croquemort-like |
| RS002081 | -1.7 | 5.7 | 1.9E-06 | -1.5 | 5.7 | 4.0E-05 | 0.2 | 5.7 | 8.1E-01 | krueppel homolog 1-like |
| RS002296 | -1.5 | 1.4 | 3.3E-08 | -1.7 | 1.4 | 1.7E-10 | -0.2 | 1.4 | 7.1E-01 | glycine receptor subunit alpha-4 |
| RS002324 | -1.3 | 4.7 | 2.0E-09 | -1.1 | 4.7 | 6.9E-07 | 0.2 | 4.7 | 5.8E-01 | dynein heavy chain axonemal |
| RS002398 | -3.6 | 4.6 | 5.1E-06 | -3.4 | 4.6 | 7.1E-06 | 0.2 | 4.6 | 9.2E-01 | endonuclease-reverse transcriptase |
| RS002524 | -1.0 | 7.6 | 1.2E-03 | -1.5 | 7.6 | 1.0E-06 | -0.5 | 7.6 | 1.9E-01 | fatty acyl- reductase cg5065 |
| RS002945 | -1.0 | 2.1 | 9.1E-04 | -1.1 | 2.1 | 5.5E-04 | -0.1 | 2.1 | 9.3E-01 | dna helicase mcm9 |
| RS003059 | -1.5 | 4.4 | 3.9E-06 | -1.8 | 4.4 | 1.0E-07 | -0.3 | 4.4 | 6.0E-01 | |
| RS003335 | -1.0 | 8.2 | 2.0E-04 | -1.7 | 8.2 | 1.3E-09 | -0.6 | 8.2 | 4.7E-02 | apolipophorins-like |
| RS003351 | -3.0 | 6.3 | 6.7E-38 | -1.1 | 6.3 | 9.5E-06 | 1.9 | 6.3 | 8.6E-17 | erad-associated e3 ubiquitin-protein ligase hrd1b-like |
| RS003387 | -2.7 | 2.3 | 3.5E-12 | -2.7 | 2.3 | 1.3E-13 | -0.1 | 2.3 | 9.5E-01 | |
| RS003472 | -1.4 | 2.9 | 5.6E-04 | -6.0 | 2.9 | 4.3E-23 | -4.6 | 2.9 | 4.6E-11 | nose resistant to fluoxetine protein 6-like |
| RS003544 | -1.7 | 2.8 | 8.2E-15 | -1.1 | 2.8 | 3.6E-07 | 0.6 | 2.8 | 3.5E-02 | glutamate receptor kainate 2 |
| RS003911 | -1.8 | 1.2 | 3.4E-05 | -1.7 | 1.2 | 7.5E-05 | 0.1 | 1.2 | 9.6E-01 | cgg triplet repeat-binding protein 1 |
| RS003912 | -2.1 | 1.4 | 1.4E-07 | -2.4 | 1.4 | 1.2E-08 | -0.2 | 1.4 | 8.1E-01 | |
| RS003954 | -2.7 | 1.4 | 9.4E-16 | -2.7 | 1.4 | 1.2E-16 | 0.0 | 1.4 | 1.0E+00 | guanine nucleotide-binding protein subunit beta-2 |
| RS004143 | -1.2 | 4.8 | 9.8E-05 | -2.6 | 4.8 | 2.5E-15 | -1.4 | 4.8 | 3.5E-04 | beta-glucosidase |
| RS005167 | -5.0 | 5.9 | 9.2E-14 | -4.9 | 5.9 | 1.6E-13 | 0.1 | 5.9 | 9.4E-01 | serine threonine-protein phosphatase 6 regulatory ankyrin repeat subunit b-like |
| RS005176 | -2.8 | 5.2 | 5.2E-13 | -1.2 | 5.2 | 2.2E-03 | 1.6 | 5.2 | 3.5E-04 | alkaline nuclease |
| RS005439 | -2.6 | 0.3 | 4.9E-03 | -2.8 | 0.3 | 3.6E-03 | -0.2 | 0.3 | 9.4E-01 | agap008309-pa-like protein |
| RS005611 | -1.5 | 5.6 | 8.2E-11 | -1.7 | 5.6 | 3.9E-13 | -0.2 | 5.6 | 6.2E-01 | ras-related protein rab-32 |
| RS005652 | -1.9 | 2.3 | 2.7E-06 | -1.1 | 2.3 | 8.4E-03 | 0.8 | 2.3 | 1.5E-01 | ras-related protein rab-7l1 |
| RS006200 | -1.4 | 2.9 | 3.9E-07 | -1.2 | 2.9 | 9.3E-06 | 0.2 | 2.9 | 7.6E-01 | dynein regulatory complex subunit 7 |
| RS006467 | -2.1 | 4.3 | 8.8E-07 | -2.4 | 4.3 | 6.7E-08 | -0.2 | 4.3 | 8.3E-01 | fibroblast growth factor receptor-like 1 |
| RS006505 | -1.5 | 1.1 | 7.5E-04 | -1.6 | 1.1 | 4.6E-04 | -0.1 | 1.1 | 9.2E-01 | |

| | | | | | | | | | | |
|---|---|---|---|---|---|---|---|---|---|---|
| RS006838 | -1.1 | 7.6 | 8.5E-07 | -1.1 | 7.6 | 1.7E-06 | 0.0 | 7.6 | 9.9E-01 | |
| RS007003 | -3.1 | 4.8 | 7.9E-29 | -2.3 | 4.8 | 2.7E-18 | 0.8 | 4.8 | 3.0E-02 | 2-hydroxyacylsphingosine 1-beta-galactosyltransferase-like |
| RS007115 | -3.0 | 0.2 | 1.9E-05 | -3.5 | 0.2 | 1.0E-06 | -0.5 | 0.2 | 7.4E-01 | chaoptin |
| RS007116 | -3.6 | 0.7 | 2.7E-10 | -3.5 | 0.7 | 2.2E-10 | 0.1 | 0.7 | 9.6E-01 | chaoptin |
| RS007118 | -2.5 | 2.7 | 1.7E-18 | -3.4 | 2.7 | 1.5E-29 | -0.9 | 2.7 | 2.6E-02 | chaoptin |
| RS007134 | -2.1 | 4.9 | 1.1E-07 | -1.1 | 4.9 | 4.7E-03 | 1.1 | 4.9 | 6.3E-02 | arylsulfatase b-like |
| RS007218 | -3.6 | -0.4 | 3.1E-04 | -3.8 | -0.4 | 1.6E-04 | -0.2 | -0.4 | 9.5E-01 | |
| RS007497 | -2.4 | 8.0 | 4.9E-07 | -3.4 | 8.0 | 5.9E-12 | -1.1 | 8.0 | 1.6E-01 | |
| RS007613 | -1.5 | 1.9 | 2.2E-05 | -2.1 | 1.9 | 1.0E-08 | -0.6 | 1.9 | 3.2E-01 | carotenoid isomerooxygenase |
| RS007636 | -3.1 | 6.8 | 1.5E-33 | -1.3 | 6.8 | 3.7E-07 | 1.8 | 6.8 | 5.4E-12 | leucine-rich repeat protein soc-2 |
| RS007637 | -3.0 | 5.0 | 3.6E-12 | -1.5 | 5.0 | 1.7E-03 | 1.5 | 5.0 | 1.3E-03 | |
| RS008372 | -1.1 | 1.3 | 3.4E-03 | -1.3 | 1.3 | 8.9E-04 | -0.2 | 1.3 | 8.1E-01 | s68306pol truncated - red flour beetle retrotransposon woot |
| RS008469 | -3.2 | 2.5 | 5.7E-14 | -3.2 | 2.5 | 4.2E-15 | 0.0 | 2.5 | 1.0E+00 | transcription factor |
| RS008849 | -2.0 | 4.3 | 1.7E-09 | -1.8 | 4.3 | 1.8E-07 | 0.2 | 4.3 | 6.9E-01 | |
| RS008850 | -1.6 | 6.3 | 2.2E-04 | -2.4 | 6.3 | 2.7E-08 | -0.8 | 6.3 | 1.1E-01 | cytochrome p450 6k1-like |
| RS009035 | -1.7 | 1.8 | 7.5E-06 | -1.3 | 1.8 | 7.2E-04 | 0.4 | 1.8 | 5.3E-01 | |
| RS009427 | -2.3 | 3.6 | 3.7E-27 | -2.6 | 3.6 | 6.4E-33 | -0.3 | 3.6 | 4.1E-01 | chorion peroxidase |
| RS009745 | -2.5 | 9.0 | 2.6E-13 | -3.3 | 9.0 | 2.1E-20 | -0.8 | 9.0 | 8.9E-02 | sphingomyelin phosphodiesterase-like |
| RS010590 | -5.1 | 7.3 | 3.6E-79 | -2.5 | 7.3 | 6.5E-27 | 2.7 | 7.3 | 1.7E-21 | acyl- synthetase short-chain family member mitochondrial |
| RS010661 | -3.4 | 1.8 | 8.9E-17 | -2.3 | 1.8 | 4.5E-10 | 1.1 | 1.8 | 6.6E-02 | long-wavelength opsin |
| RS011038 | -1.1 | 4.0 | 2.5E-03 | -1.1 | 4.0 | 5.1E-03 | 0.0 | 4.0 | 9.9E-01 | protein giant-lens-like |
| RS011042 | -1.8 | 1.8 | 1.7E-06 | -1.3 | 1.8 | 5.5E-04 | 0.5 | 1.8 | 3.9E-01 | eri1 exoribonuclease 2 |
| RS011327 | -2.3 | 4.6 | 5.2E-40 | -5.5 | 4.6 | 6.6E-132 | -3.2 | 4.6 | 9.3E-38 | fatty acyl- reductase cg5065 |
| RS011729 | -1.6 | 6.1 | 5.2E-06 | -2.8 | 6.1 | 2.4E-14 | -1.1 | 6.1 | 3.3E-03 | |
| RS011857 | -2.0 | 5.5 | 4.6E-33 | -1.8 | 5.5 | 9.3E-27 | 0.2 | 5.5 | 4.0E-01 | 2-amino-3-ketobutyrate coenzyme a mitochondrial |
| RS012257 | -2.0 | 1.4 | 1.2E-07 | -3.0 | 1.4 | 1.3E-13 | -1.0 | 1.4 | 1.2E-01 | pr domain zinc finger protein 1 |
| RS012330 | -2.0 | 3.3 | 1.2E-03 | -2.0 | 3.3 | 2.1E-03 | 0.0 | 3.3 | 9.8E-01 | ankyrin repeat protein |
| RS012610 | -3.0 | 2.2 | 8.2E-16 | -2.1 | 2.2 | 1.1E-08 | 1.0 | 2.2 | 4.9E-02 | arrestin homolog |
| RS012980 | -1.3 | 2.6 | 1.7E-03 | -1.4 | 2.6 | 1.2E-03 | -0.1 | 2.6 | 9.1E-01 | lim homeobox transcription factor 1-beta |
| RS013083 | -1.7 | 4.5 | 2.6E-03 | -4.3 | 4.5 | 6.9E-12 | -2.7 | 4.5 | 1.7E-04 | maltase |
| RS013482 | -2.8 | 3.6 | 7.2E-16 | -1.5 | 3.6 | 2.0E-06 | 1.3 | 3.6 | 3.9E-03 | |
| RS013977 | -5.3 | 1.5 | 1.3E-06 | -5.5 | 1.5 | 2.6E-07 | -0.1 | 1.5 | 1.0E+00 | vitellogenin |
| RS014586 | -1.0 | 2.3 | 3.8E-04 | -1.1 | 2.3 | 2.8E-04 | -0.1 | 2.3 | 9.3E-01 | bride of sevenless |
| RS014971 | -2.5 | 2.8 | 5.1E-12 | -2.4 | 2.8 | 8.9E-12 | 0.1 | 2.8 | 9.2E-01 | growth differentiation factor 8 |
| RS014972 | -2.4 | 1.9 | 5.7E-06 | -3.0 | 1.9 | 2.5E-08 | -0.7 | 1.9 | 5.0E-01 | growth differentiation factor 8 |
| RS015143 | -1.8 | 3.1 | 3.9E-07 | -1.7 | 3.1 | 1.8E-06 | 0.1 | 3.1 | 8.8E-01 | |
| RS015418 | -2.8 | 1.1 | 1.7E-09 | -2.0 | 1.1 | 4.6E-06 | 0.8 | 1.1 | 2.9E-01 | |
| RS015422 | -2.1 | 0.9 | 2.3E-08 | -2.1 | 0.9 | 1.2E-08 | 0.0 | 0.9 | 9.9E-01 | |
| RS015423 | -2.8 | 3.5 | 1.7E-21 | -2.4 | 3.5 | 5.1E-17 | 0.4 | 3.5 | 4.0E-01 | neither inactivation nor afterpotential protein c |
| RS100008 | -2.7 | -0.4 | 5.1E-03 | -2.9 | -0.4 | 3.7E-03 | -0.2 | -0.4 | 9.3E-01 | glycine receptor subunit alpha-3 |
| RS100013 | -4.2 | 4.2 | 2.5E-35 | -1.4 | 4.2 | 1.2E-05 | 2.8 | 4.2 | 2.6E-16 | geranylgeranyl pyrophosphate synthase |

SI_Dataset_1
Caste-biased genes: Reproductive (thorax + abdomen)

| Gene ID | Thorax + abdomen Reproductive v.s. Soldier | | | Thorax + abdomen Reproductive v.s. Worker | | | Thorax + abdomen Soldier v.s. Worker | | | Gene annotation |
|---|---|---|---|---|---|---|---|---|---|---|
| | logFC | logCPM | FDR | logFC | logCPM | FDR | logFC | logCPM | FDR | |
| RS000203 | -1.6 | 2.9 | 7.4E-08 | -1.8 | 2.9 | 5.7E-09 | -0.2 | 2.9 | 7.3E-01 | |
| RS000251 | -1.2 | 4.2 | 2.7E-06 | -1.1 | 4.2 | 6.4E-06 | 0.0 | 4.2 | 9.4E-01 | |
| RS000340 | -3.7 | 2.3 | 1.0E-07 | -3.2 | 2.3 | 1.0E-06 | 0.5 | 2.3 | 6.9E-01 | |
| RS000346 | -1.2 | 3.3 | 1.3E-03 | -1.2 | 3.3 | 3.0E-03 | 0.1 | 3.3 | 8.8E-01 | hemicentin-2 |
| RS000358 | -3.1 | 0.5 | 1.4E-08 | -1.3 | 0.5 | 2.9E-03 | 1.8 | 0.5 | 3.1E-02 | dynein intermediate chain ciliary |
| RS000417 | -2.1 | 2.1 | 1.6E-07 | -2.4 | 2.1 | 2.4E-08 | -0.3 | 2.1 | 7.6E-01 | krueppel homolog 1 |
| RS000445 | -1.7 | 5.1 | 7.2E-20 | -2.4 | 5.1 | 1.2E-36 | -0.7 | 5.1 | 9.3E-04 | cytochrome p450 |
| RS000509 | -1.0 | 3.7 | 4.5E-04 | -1.2 | 3.7 | 1.4E-05 | -0.2 | 3.7 | 5.9E-01 | |
| RS000529 | -1.9 | 1.6 | 3.5E-04 | -2.3 | 1.6 | 6.6E-05 | -0.4 | 1.6 | 6.9E-01 | |
| RS000534 | -8.4 | 1.4 | 3.1E-21 | -4.9 | 1.4 | 1.3E-13 | 3.5 | 1.4 | 2.9E-02 | insulin-like peptide 1 |
| RS000536 | -2.7 | 2.0 | 6.1E-05 | -2.9 | 2.0 | 1.2E-04 | -0.1 | 2.0 | 9.2E-01 | lirp-like |
| RS000610 | -9.1 | 6.5 | 6.5E-30 | -9.3 | 6.5 | 1.9E-30 | -0.3 | 6.5 | 8.7E-01 | vitellogenin |
| RS000616 | -9.2 | 5.0 | 1.3E-29 | -8.0 | 5.0 | 1.4E-25 | 1.2 | 5.0 | 4.9E-01 | vitellogenin |
| RS000618 | -2.2 | 2.3 | 3.6E-05 | -1.4 | 2.3 | 8.8E-03 | 0.8 | 2.3 | 2.4E-01 | structural maintenance of chromosomes protein 1a |
| RS000643 | -1.0 | 2.2 | 3.4E-03 | -1.5 | 2.2 | 6.9E-05 | -0.4 | 2.2 | 4.5E-01 | beat protein |
| RS000762 | -5.0 | 2.0 | 1.3E-13 | -3.0 | 2.0 | 1.4E-07 | 2.0 | 2.0 | 1.0E-01 | |
| RS000785 | -1.5 | 2.6 | 3.1E-03 | -1.3 | 2.6 | 6.5E-03 | 0.1 | 2.6 | 8.5E-01 | |
| RS000786 | -9.1 | 1.8 | 3.4E-13 | -4.7 | 1.8 | 2.8E-07 | 4.5 | 1.8 | 3.5E-03 | serine protease snake-like |
| RS000840 | -1.2 | 4.2 | 2.8E-03 | -1.1 | 4.2 | 4.7E-03 | 0.1 | 4.2 | 9.3E-01 | |
| RS000932 | -2.1 | 0.8 | 6.6E-03 | -3.0 | 0.8 | 1.2E-04 | -0.9 | 0.8 | 4.5E-01 | single-minded homolog 1-like |
| RS001024 | -1.7 | 3.0 | 1.5E-06 | -1.3 | 3.0 | 4.5E-04 | 0.4 | 3.0 | 3.8E-01 | |
| RS001113 | -1.4 | 2.3 | 2.2E-04 | -2.3 | 2.3 | 3.9E-08 | -0.9 | 2.3 | 1.4E-01 | bmp and activin membrane-bound inhibitor homolog |
| RS001261 | -1.4 | 5.8 | 1.9E-08 | -2.2 | 5.8 | 6.8E-17 | -0.9 | 5.8 | 1.7E-02 | elongation of very long chain fatty acids protein 7 |
| RS001362 | -3.4 | 2.4 | 9.2E-19 | -1.2 | 2.4 | 7.9E-04 | 2.2 | 2.4 | 2.4E-07 | serine proteases 1 2-like |
| RS001418 | -4.3 | 3.2 | 2.7E-23 | -2.2 | 3.2 | 3.2E-08 | 2.1 | 3.2 | 7.2E-06 | acyl- synthetase family member mitochondrial |
| RS001422 | -3.6 | 5.8 | 2.9E-23 | -5.7 | 5.8 | 4.3E-42 | -2.0 | 5.8 | 4.7E-06 | |
| RS001423 | -3.2 | 3.0 | 1.6E-10 | -5.6 | 3.0 | 1.3E-19 | -2.5 | 3.0 | 2.7E-03 | venom carboxylesterase-6-like |
| RS001472 | -3.0 | 1.5 | 3.0E-04 | -2.5 | 1.5 | 3.8E-03 | 0.5 | 1.5 | 7.2E-01 | |
| RS001520 | -4.2 | -0.2 | 4.4E-03 | -5.4 | -0.2 | 9.7E-04 | -1.2 | -0.2 | 5.0E-01 | zinc finger protein 530 |
| RS001535 | -2.9 | 6.3 | 4.2E-07 | -3.0 | 6.3 | 1.4E-07 | -0.1 | 6.3 | 8.8E-01 | vitellogenin receptor |
| RS001539 | -1.4 | 3.3 | 4.2E-04 | -1.6 | 3.3 | 5.2E-05 | -0.2 | 3.3 | 7.7E-01 | transient receptor potential cation channel protein painless |
| RS001654 | -1.6 | 5.5 | 1.8E-05 | -1.8 | 5.5 | 1.2E-06 | -0.2 | 5.5 | 7.4E-01 | catalase |
| RS001672 | -1.0 | 6.1 | 7.0E-03 | -1.8 | 6.1 | 2.1E-06 | -0.7 | 6.1 | 1.1E-01 | catalase |
| RS001732 | -6.6 | 1.6 | 8.8E-20 | -8.5 | 1.6 | 2.2E-20 | -1.9 | 1.6 | 3.0E-01 | |
| RS001733 | -6.8 | 1.2 | 9.2E-42 | -8.0 | 1.2 | 2.4E-37 | -1.2 | 1.2 | 4.8E-01 | |
| RS001746 | -1.2 | 1.1 | 5.2E-03 | -1.3 | 1.1 | 2.9E-03 | -0.1 | 1.1 | 8.9E-01 | forkhead box protein j3 |
| RS001772 | -1.7 | 2.6 | 8.6E-04 | -2.5 | 2.6 | 2.0E-06 | -0.8 | 2.6 | 2.6E-01 | udp-glucuronosyltransferase 2b13-like |
| RS001804 | -3.4 | 5.0 | 1.7E-54 | -2.4 | 5.0 | 7.3E-32 | 1.0 | 5.0 | 4.6E-04 | udp-glucuronosyltransferase |
| RS001883 | -1.1 | 7.1 | 2.0E-10 | -1.8 | 7.1 | 7.6E-29 | -0.8 | 7.1 | 1.3E-05 | protein croquemort-like |
| RS001884 | -1.4 | 6.0 | 3.7E-16 | -1.5 | 6.0 | 1.4E-19 | -0.2 | 6.0 | 5.2E-01 | protein croquemort-like |
| RS001962 | -1.9 | 0.8 | 6.0E-04 | -1.4 | 0.8 | 8.9E-03 | 0.5 | 0.8 | 6.2E-01 | gastrin-releasing peptide receptor-like |
| RS002039 | -1.1 | 4.0 | 3.4E-04 | -1.2 | 4.0 | 1.8E-04 | 0.0 | 4.0 | 9.6E-01 | low quality protein: fibrillin-1 |

| | | | | | | | | | | |
|---|---|---|---|---|---|---|---|---|---|---|
| RS002081 | -1.3 | 5.7 | 3.8E-04 | -1.5 | 5.7 | 4.6E-05 | -0.2 | 5.7 | 7.5E-01 | krueppel homolog 1-like |
| RS002145 | -8.5 | 1.5 | 2.4E-41 | -5.7 | 1.5 | 7.5E-31 | 2.8 | 1.5 | 1.0E-01 | lirp-like |
| RS002205 | -4.4 | -0.6 | 6.2E-03 | -4.3 | -0.6 | 9.8E-03 | 0.2 | -0.6 | 1.0E+00 | |
| RS002305 | -1.4 | 2.3 | 4.0E-06 | -1.3 | 2.3 | 1.8E-05 | 0.1 | 2.3 | 8.9E-01 | |
| RS002324 | -1.1 | 4.7 | 5.2E-06 | -1.1 | 4.7 | 6.9E-06 | 0.0 | 4.7 | 9.8E-01 | dynein heavy chain axonemal |
| RS002355 | -5.4 | -0.3 | 8.9E-04 | -5.2 | -0.3 | 1.6E-03 | 0.2 | -0.3 | 1.0E+00 | |
| RS002427 | -2.3 | 2.1 | 5.6E-05 | -1.8 | 2.1 | 1.5E-03 | 0.5 | 2.1 | 5.2E-01 | |
| RS002469 | -5.1 | 0.2 | 7.7E-07 | -4.8 | 0.2 | 5.3E-06 | 0.3 | 0.2 | 9.3E-01 | |
| RS002470 | -6.2 | 1.7 | 1.0E-34 | -6.4 | 1.7 | 1.8E-31 | -0.2 | 1.7 | 9.2E-01 | sialin |
| RS002493 | -1.0 | 2.7 | 9.6E-03 | -1.1 | 2.7 | 6.1E-03 | -0.1 | 2.7 | 9.3E-01 | hypothetical protein |
| RS002557 | -1.5 | 6.4 | 8.7E-07 | -1.4 | 6.4 | 3.4E-06 | 0.1 | 6.4 | 8.8E-01 | ankyrin-1-like |
| RS002624 | -2.6 | 2.2 | 1.5E-10 | -1.8 | 2.2 | 7.0E-06 | 0.8 | 2.2 | 1.7E-01 | protein aubergine-like |
| RS002625 | -2.1 | 0.0 | 2.7E-03 | -3.0 | 0.0 | 1.4E-04 | -0.9 | 0.0 | 5.2E-01 | protein aubergine-like |
| RS002649 | -1.2 | 4.0 | 6.3E-03 | -2.3 | 4.0 | 3.8E-08 | -1.1 | 4.0 | 1.8E-02 | suppressor protein srp40-like |
| RS002740 | -1.6 | 3.7 | 1.7E-12 | -1.7 | 3.7 | 4.9E-14 | -0.1 | 3.7 | 7.1E-01 | hypothetical protein |
| RS002754 | -1.2 | 3.2 | 3.6E-04 | -1.1 | 3.2 | 2.0E-03 | 0.1 | 3.2 | 8.0E-01 | cell division cycle-associated protein 7-like |
| RS002862 | -1.3 | 3.4 | 1.7E-03 | -1.7 | 3.4 | 1.6E-05 | -0.4 | 3.4 | 4.3E-01 | cytochrome p450 306a1 |
| RS002878 | -7.1 | 1.5 | 2.8E-28 | -6.9 | 1.5 | 4.6E-25 | 0.3 | 1.5 | 9.4E-01 | e3 ubiquitin-protein ligase trim71 |
| RS003000 | -1.2 | 1.3 | 9.2E-03 | -1.4 | 1.3 | 3.6E-03 | -0.2 | 1.3 | 8.4E-01 | gamma-1-syntrophin |
| RS003059 | -1.1 | 4.4 | 7.6E-04 | -1.0 | 4.4 | 2.7E-03 | 0.1 | 4.4 | 8.4E-01 | |
| RS003073 | -1.3 | 3.7 | 8.1E-07 | -1.3 | 3.7 | 4.7E-07 | -0.1 | 3.7 | 9.2E-01 | |
| RS003134 | -1.2 | 7.0 | 7.8E-05 | -1.5 | 7.0 | 2.3E-06 | -0.2 | 7.0 | 6.3E-01 | glutathione s-transferase |
| RS003147 | -1.3 | 3.5 | 3.6E-04 | -2.0 | 3.5 | 1.8E-08 | -0.8 | 3.5 | 1.1E-01 | acetylcholine receptor subunit beta-like 1 |
| RS003315 | -1.2 | 2.9 | 5.4E-05 | -1.1 | 2.9 | 5.3E-04 | 0.2 | 2.9 | 7.4E-01 | e3 ubiquitin-protein ligase traip |
| RS003332 | -1.8 | 3.1 | 6.6E-12 | -1.9 | 3.1 | 2.0E-12 | -0.1 | 3.1 | 8.3E-01 | zinc finger protein 423 homolog |
| RS003385 | -2.6 | 1.6 | 1.4E-04 | -3.2 | 1.6 | 3.7E-06 | -0.6 | 1.6 | 6.1E-01 | kinesin-like protein kif23 |
| RS003488 | -2.4 | 4.0 | 1.9E-04 | -2.6 | 4.0 | 1.5E-05 | -0.2 | 4.0 | 8.6E-01 | heparan-alpha-glucosaminide n-acetyltransferase |
| RS003601 | -1.6 | 3.7 | 9.5E-06 | -1.3 | 3.7 | 2.1E-04 | 0.2 | 3.7 | 6.6E-01 | |
| RS003686 | -1.4 | 2.7 | 2.9E-06 | -1.4 | 2.7 | 2.8E-06 | 0.0 | 2.7 | 9.7E-01 | dynein heavy chain axonemal |
| RS003729 | -3.1 | 2.1 | 1.8E-04 | -5.0 | 2.1 | 5.7E-09 | -1.8 | 2.1 | 1.4E-01 | neuronal acetylcholine receptor subunit alpha-7 |
| RS003783 | -2.1 | 4.2 | 2.3E-13 | -1.7 | 4.2 | 3.2E-09 | 0.4 | 4.2 | 3.5E-01 | sprt-like domain-containing protein spartan |
| RS003816 | -4.2 | 9.1 | 5.4E-24 | -1.9 | 9.1 | 3.9E-06 | 2.4 | 9.1 | 7.1E-09 | serine proteases 1 2-like |
| RS003863 | -1.0 | 7.2 | 3.5E-07 | -1.2 | 7.2 | 1.3E-09 | -0.2 | 7.2 | 5.2E-01 | hemocyte protein-glutamine gamma-glutamyltransferase-like |
| RS003882 | -2.8 | 2.7 | 1.1E-06 | -2.7 | 2.7 | 3.2E-06 | 0.1 | 2.7 | 9.3E-01 | trithorax group protein osa-like |
| RS003932 | -1.8 | 7.9 | 1.6E-13 | -1.8 | 7.9 | 1.2E-12 | 0.1 | 7.9 | 8.8E-01 | membrane metallo-endopeptidase-like 1 |
| RS004029 | -1.5 | 1.6 | 5.0E-03 | -2.9 | 1.6 | 2.7E-07 | -1.5 | 1.6 | 5.2E-02 | lim homeobox protein awh |
| RS004064 | -6.1 | 0.1 | 6.8E-13 | -3.8 | 0.1 | 2.0E-08 | 2.3 | 0.1 | 2.0E-01 | pancreatic triacylglycerol lipase-like |
| RS004089 | -2.1 | -0.4 | 6.2E-03 | -3.4 | -0.4 | 2.4E-04 | -1.4 | -0.4 | 4.1E-01 | limulus clotting factor c-like |
| RS004194 | -1.2 | 4.5 | 2.7E-07 | -1.0 | 4.5 | 3.6E-05 | 0.2 | 4.5 | 5.3E-01 | |
| RS004223 | -1.5 | 4.5 | 1.1E-05 | -1.4 | 4.5 | 5.8E-05 | 0.2 | 4.5 | 7.7E-01 | spatzle-like protein |
| RS004361 | -3.2 | -0.2 | 6.1E-04 | -3.6 | -0.2 | 5.2E-04 | -0.4 | -0.2 | 8.4E-01 | homeobox protein not2 |
| RS004425 | -7.9 | 4.3 | 1.6E-51 | -6.6 | 4.3 | 5.7E-43 | 1.4 | 4.3 | 1.4E-01 | |
| RS004426 | -5.1 | 4.4 | 9.5E-06 | -4.0 | 4.4 | 1.3E-04 | 1.1 | 4.4 | 4.5E-01 | |
| RS004430 | -1.6 | 3.3 | 7.2E-07 | -2.4 | 3.3 | 7.3E-13 | -0.8 | 3.3 | 6.1E-02 | glutamate receptor nmda 3a-like |

| | | | | | | | | | | |
|---|---|---|---|---|---|---|---|---|---|---|
| RS004450 | -1.4 | 6.0 | 1.0E-13 | -1.0 | 6.0 | 1.8E-07 | 0.4 | 6.0 | 9.6E-02 | cytochrome p450 4c3 |
| RS004525 | -1.5 | 3.5 | 2.7E-06 | -2.3 | 3.5 | 1.8E-11 | -0.8 | 3.5 | 1.0E-01 | cyclin-dependent kinase-like 1 |
| RS004579 | -1.5 | 2.9 | 2.5E-05 | -1.3 | 2.9 | 1.6E-04 | 0.2 | 2.9 | 7.9E-01 | ubiquitin-like-conjugating enzyme atg10 |
| RS004624 | -6.5 | 7.9 | 4.7E-05 | -7.6 | 7.9 | 1.8E-06 | -1.2 | 7.9 | 3.4E-01 | myrosinase 1-like |
| RS004639 | -1.3 | 3.3 | 2.3E-06 | -1.4 | 3.3 | 2.4E-06 | 0.0 | 3.3 | 9.6E-01 | neural cell adhesion molecule 1 |
| RS004718 | -3.7 | 0.3 | 6.7E-05 | -2.4 | 0.3 | 4.7E-03 | 1.2 | 0.3 | 4.0E-01 | facilitated trehalose transporter tret1 |
| RS004822 | -1.6 | 5.5 | 1.4E-20 | -2.4 | 5.5 | 1.1E-41 | -0.8 | 5.5 | 5.2E-05 | mitochondrial amidoxime-reducing component 1-like |
| RS004825 | -1.6 | 4.0 | 7.1E-07 | -1.1 | 4.0 | 1.1E-03 | 0.5 | 4.0 | 2.2E-01 | hyaluronan mediated motility receptor-like |
| RS004832 | -1.4 | 1.9 | 2.0E-05 | -1.2 | 1.9 | 4.4E-04 | 0.3 | 1.9 | 6.5E-01 | |
| RS005041 | -1.2 | 3.8 | 1.9E-05 | -1.1 | 3.8 | 1.2E-04 | 0.1 | 3.8 | 8.2E-01 | zinc finger protein 91-like |
| RS005117 | -4.9 | 0.3 | 8.9E-06 | -3.2 | 0.3 | 2.7E-03 | 1.7 | 0.3 | 3.7E-01 | ankyrin repeat protein |
| RS005233 | -1.1 | 5.3 | 1.3E-04 | -1.4 | 5.3 | 2.5E-07 | -0.4 | 5.3 | 3.2E-01 | u3 small nucleolar rna-interacting protein 2 |
| RS005246 | -1.3 | 4.3 | 2.5E-08 | -1.4 | 4.3 | 5.1E-10 | -0.2 | 4.3 | 6.6E-01 | high-affinity choline transporter 1-like |
| RS005262 | -1.0 | 5.8 | 2.9E-03 | -1.7 | 5.8 | 1.2E-07 | -0.7 | 5.8 | 5.3E-02 | rho guanine nucleotide exchange factor at |
| RS005385 | -2.6 | -0.1 | 8.9E-04 | -3.5 | -0.1 | 1.5E-04 | -0.9 | -0.1 | 6.1E-01 | |
| RS005428 | -2.0 | 0.3 | 4.4E-04 | -1.5 | 0.3 | 6.0E-03 | 0.4 | 0.3 | 6.7E-01 | e3 ubiquitin-protein ligase siah1-like |
| RS005521 | -1.8 | 1.3 | 3.1E-04 | -1.5 | 1.3 | 3.8E-03 | 0.3 | 1.3 | 6.8E-01 | cyclin-dependent kinases regulatory subunit |
| RS005524 | -1.1 | 4.1 | 2.7E-06 | -1.0 | 4.1 | 2.7E-05 | 0.1 | 4.1 | 7.8E-01 | gonadotropin-releasing hormone ii receptor |
| RS005611 | -1.3 | 5.6 | 6.7E-08 | -1.5 | 5.6 | 6.0E-11 | -0.3 | 5.6 | 4.2E-01 | ras-related protein rab-32 |
| RS005612 | -3.7 | 3.1 | 3.2E-08 | -4.1 | 3.1 | 5.3E-09 | -0.3 | 3.1 | 7.8E-01 | |
| RS005652 | -1.8 | 2.3 | 2.2E-08 | -2.2 | 2.3 | 5.2E-11 | -0.4 | 2.3 | 4.5E-01 | ras-related protein rab-7l1 |
| RS005690 | -1.0 | 1.7 | 5.6E-03 | -1.0 | 1.7 | 6.8E-03 | 0.0 | 1.7 | 9.8E-01 | protein |
| RS005702 | -1.4 | 4.2 | 3.6E-10 | -1.1 | 4.2 | 6.0E-06 | 0.4 | 4.2 | 2.3E-01 | |
| RS005732 | -2.0 | 1.0 | 2.4E-03 | -1.9 | 1.0 | 6.3E-03 | 0.2 | 1.0 | 8.8E-01 | protein wnt-5b-like |
| RS005806 | -2.1 | 2.1 | 3.4E-08 | -2.3 | 2.1 | 1.9E-08 | -0.2 | 2.1 | 8.5E-01 | hypothetical protein |
| RS005812 | -1.0 | 1.4 | 6.7E-03 | -1.0 | 1.4 | 1.0E-02 | 0.0 | 1.4 | 9.8E-01 | zinc finger protein 239-like |
| RS005983 | -1.2 | 1.9 | 3.6E-04 | -1.1 | 1.9 | 1.5E-03 | 0.1 | 1.9 | 8.6E-01 | |
| RS005988 | -1.6 | 1.4 | 6.7E-04 | -1.2 | 1.4 | 9.5E-03 | 0.4 | 1.4 | 6.6E-01 | whirlin |
| RS006040 | -1.3 | 3.0 | 4.7E-04 | -1.9 | 3.0 | 6.5E-07 | -0.6 | 3.0 | 2.6E-01 | transcription factor hes-1 |
| RS006196 | -6.4 | 0.1 | 2.0E-14 | -6.2 | 0.1 | 2.8E-12 | 0.2 | 0.1 | 1.0E+00 | |
| RS006199 | -1.4 | 3.3 | 4.1E-06 | -1.1 | 3.3 | 2.7E-04 | 0.3 | 3.3 | 5.5E-01 | |
| RS006247 | -2.6 | 2.2 | 4.4E-10 | -1.3 | 2.2 | 1.4E-03 | 1.3 | 2.2 | 1.1E-02 | calpain-d-like |
| RS006280 | -2.3 | 8.0 | 2.2E-35 | -1.6 | 8.0 | 3.7E-18 | 0.7 | 8.0 | 7.7E-04 | aminopeptidase n |
| RS006331 | -1.8 | 3.1 | 8.3E-07 | -2.1 | 3.1 | 6.9E-08 | -0.3 | 3.1 | 6.9E-01 | |
| RS006412 | -1.4 | 4.1 | 2.8E-08 | -1.2 | 4.1 | 1.5E-06 | 0.2 | 4.1 | 6.6E-01 | fanconi anemia group j protein |
| RS006499 | -1.9 | 2.7 | 4.2E-06 | -2.6 | 2.7 | 1.9E-08 | -0.7 | 2.7 | 3.9E-01 | transient receptor potential cation channel subfamily v member 5 |
| RS006506 | -1.4 | 2.6 | 2.9E-03 | -2.0 | 2.6 | 1.5E-05 | -0.7 | 2.6 | 3.1E-01 | perphorin-1 |
| RS006632 | -1.5 | 2.4 | 3.9E-06 | -1.5 | 2.4 | 5.0E-06 | 0.0 | 2.4 | 9.7E-01 | synaptotagmin |
| RS006639 | -3.7 | 5.4 | 6.3E-15 | -2.3 | 5.4 | 9.5E-07 | 1.4 | 5.4 | 1.8E-02 | aromatic-l-amino-acid decarboxylase-like |
| RS006802 | -2.0 | 3.9 | 7.4E-09 | -1.1 | 3.9 | 1.6E-03 | 0.9 | 3.9 | 3.3E-02 | |
| RS006867 | -2.9 | -0.2 | 2.5E-04 | -1.9 | -0.2 | 8.1E-03 | 1.0 | -0.2 | 4.9E-01 | soluble guanylate cyclase 89db-like |

| | | | | | | | | | | |
|---|---|---|---|---|---|---|---|---|---|---|
| RS006881 | -4.1 | 3.3 | 1.7E-05 | -4.0 | 3.3 | 3.6E-05 | 0.1 | 3.3 | 9.7E-01 | serine protease nudel |
| RS006921 | -2.7 | 0.4 | 2.2E-05 | -3.9 | 0.4 | 3.3E-07 | -1.2 | 0.4 | 4.2E-01 | glutamate receptor nmda 2b |
| RS007010 | -1.4 | 2.5 | 2.5E-03 | -1.3 | 2.5 | 4.2E-03 | 0.1 | 2.5 | 9.1E-01 | |
| RS007048 | -1.4 | 2.1 | 7.9E-06 | -1.6 | 2.1 | 3.3E-06 | -0.1 | 2.1 | 8.5E-01 | dual 3 -cyclic-amp and -gmp phosphodiesterase 11a |
| RS007063 | -1.1 | 4.5 | 1.9E-07 | -1.0 | 4.5 | 1.3E-06 | 0.1 | 4.5 | 8.3E-01 | |
| RS007090 | -1.4 | 3.2 | 1.4E-04 | -1.2 | 3.2 | 8.0E-04 | 0.1 | 3.2 | 8.4E-01 | |
| RS007098 | -1.7 | 0.9 | 1.1E-04 | -2.8 | 0.9 | 7.7E-08 | -1.1 | 0.9 | 2.0E-01 | |
| RS007124 | -1.2 | 5.3 | 1.2E-08 | -2.0 | 5.3 | 4.1E-21 | -0.8 | 5.3 | 9.1E-04 | ets-domain protein |
| RS007144 | -1.0 | 3.1 | 4.6E-06 | -1.1 | 3.1 | 1.1E-06 | -0.1 | 3.1 | 8.4E-01 | |
| RS007153 | -1.4 | 3.5 | 1.2E-06 | -1.5 | 3.5 | 4.8E-07 | -0.1 | 3.5 | 8.9E-01 | neuroligin- y-linked-like |
| RS007222 | -1.1 | 2.5 | 3.5E-03 | -1.4 | 2.5 | 3.9E-04 | -0.3 | 2.5 | 6.8E-01 | muscle segmentation homeobox |
| RS007244 | -1.6 | 7.3 | 1.7E-05 | -2.1 | 7.3 | 4.1E-09 | -0.5 | 7.3 | 2.3E-01 | mucin |
| RS007331 | -1.4 | 3.0 | 2.2E-05 | -1.2 | 3.0 | 3.5E-04 | 0.2 | 3.0 | 6.9E-01 | ankyrin repeat and lem domain-containing protein 2 |
| RS007466 | -1.3 | 1.4 | 4.1E-03 | -1.8 | 1.4 | 1.1E-04 | -0.5 | 1.4 | 4.8E-01 | ras-related and estrogen-regulated growth inhibitor-like |
| RS007481 | -1.6 | 3.8 | 4.8E-03 | -2.3 | 3.8 | 1.0E-04 | -0.7 | 3.8 | 3.8E-01 | geranylgeranyl pyrophosphate synthase |
| RS007493 | -1.6 | 4.1 | 1.2E-05 | -1.1 | 4.1 | 5.2E-03 | 0.5 | 4.1 | 2.5E-01 | targeting protein for xklp2 |
| RS007597 | -1.5 | 3.4 | 6.7E-06 | -1.4 | 3.4 | 8.9E-06 | 0.0 | 3.4 | 9.8E-01 | endonuclease iii-like protein 1 |
| RS007636 | -2.1 | 6.8 | 9.7E-17 | -1.6 | 6.8 | 4.1E-10 | 0.5 | 6.8 | 1.0E-01 | leucine-rich repeat protein soc-2 |
| RS007637 | -2.1 | 5.0 | 1.9E-06 | -1.8 | 5.0 | 5.1E-05 | 0.3 | 5.0 | 6.3E-01 | |
| RS007699 | -5.0 | 1.5 | 1.1E-11 | -2.8 | 1.5 | 1.6E-05 | 2.1 | 1.5 | 4.9E-02 | hypothetical protein |
| RS007731 | -1.8 | 3.0 | 7.2E-04 | -1.4 | 3.0 | 7.5E-03 | 0.3 | 3.0 | 6.6E-01 | metabotropic glutamate receptor 3-like |
| RS007753 | -1.5 | 2.9 | 1.2E-03 | -1.3 | 2.9 | 4.6E-03 | 0.2 | 2.9 | 8.0E-01 | |
| RS007761 | -1.0 | 4.7 | 2.4E-06 | -1.1 | 4.7 | 8.8E-08 | -0.1 | 4.7 | 6.7E-01 | zinc finger protein 782 |
| RS007825 | -1.2 | 1.4 | 1.7E-03 | -1.1 | 1.4 | 7.9E-03 | 0.2 | 1.4 | 8.1E-01 | diphthine methyltransferase |
| RS007918 | -6.3 | -0.1 | 3.1E-10 | -6.5 | -0.1 | 2.3E-10 | -0.2 | -0.1 | 1.0E+00 | |
| RS007951 | -1.1 | 8.3 | 4.0E-07 | -1.7 | 8.3 | 5.0E-17 | -0.7 | 8.3 | 4.7E-03 | lipophorin receptor |
| RS007971 | -1.2 | 5.4 | 1.3E-08 | -1.1 | 5.4 | 2.9E-07 | 0.1 | 5.4 | 7.6E-01 | arylsulfatase b |
| RS008171 | -4.4 | 1.2 | 1.4E-11 | -3.4 | 1.2 | 1.5E-07 | 1.0 | 1.2 | 4.7E-01 | alpha-1a adrenergic receptor |
| RS008189 | -2.6 | 1.7 | 1.7E-09 | -2.2 | 1.7 | 4.9E-07 | 0.4 | 1.7 | 6.0E-01 | voltage-dependent calcium channel gamma-7 subunit |
| RS008245 | -1.6 | 3.5 | 3.2E-05 | -1.5 | 3.5 | 1.0E-04 | 0.1 | 3.5 | 8.6E-01 | fatty-acid amide hydrolase 2 |
| RS008256 | -3.5 | 0.1 | 1.8E-05 | -2.2 | 0.1 | 2.1E-03 | 1.3 | 0.1 | 3.4E-01 | protein gooseberry-like |
| RS008330 | -1.4 | 3.6 | 3.2E-07 | -1.2 | 3.6 | 1.1E-05 | 0.2 | 3.6 | 6.9E-01 | dimethyladenosine transferase mitochondrial |
| RS008332 | -1.3 | 4.4 | 4.6E-07 | -1.3 | 4.4 | 5.0E-07 | 0.0 | 4.4 | 9.9E-01 | thap domain-containing protein 4-like |
| RS008335 | -1.6 | 5.1 | 3.6E-05 | -2.1 | 5.1 | 6.2E-08 | -0.5 | 5.1 | 3.5E-01 | |
| RS008348 | -1.5 | 2.6 | 3.1E-06 | -1.2 | 2.6 | 2.2E-04 | 0.3 | 2.6 | 5.4E-01 | |
| RS008360 | -1.7 | 2.8 | 1.7E-09 | -1.2 | 2.8 | 3.1E-05 | 0.5 | 2.8 | 1.8E-01 | |
| RS008412 | -3.5 | 0.4 | 1.1E-06 | -2.7 | 0.4 | 1.2E-04 | 0.9 | 0.4 | 5.6E-01 | dynein assembly factor |
| RS008413 | -2.7 | 1.9 | 1.5E-08 | -2.0 | 1.9 | 2.1E-05 | 0.7 | 1.9 | 3.5E-01 | |
| RS008469 | -4.6 | 2.5 | 4.1E-36 | -4.5 | 2.5 | 1.9E-33 | 0.1 | 2.5 | 9.3E-01 | transcription factor |
| RS008621 | -1.6 | 4.8 | 9.8E-05 | -1.9 | 4.8 | 1.6E-06 | -0.3 | 4.8 | 5.5E-01 | katanin p60 atpase-containing subunit a1 |
| RS008687 | -1.2 | 3.5 | 1.4E-05 | -1.2 | 3.5 | 6.5E-05 | 0.1 | 3.5 | 8.6E-01 | origin recognition complex subunit 1 |
| RS008723 | -1.9 | 3.3 | 4.3E-06 | -3.1 | 3.3 | 3.1E-13 | -1.2 | 3.3 | 1.7E-02 | excitatory amino acid transporter |

| | | | | | | | | | | |
|---|---|---|---|---|---|---|---|---|---|---|
| RS008871 | -2.8 | 1.8 | 1.6E-12 | -2.8 | 1.8 | 2.1E-11 | 0.1 | 1.8 | 9.7E-01 | ras-specific guanine nucleotide-releasing factor 2-like |
| RS008881 | -7.6 | 7.6 | 4.3E-10 | -7.5 | 7.6 | 5.2E-10 | 0.1 | 7.6 | 9.4E-01 | apolipoprotein d |
| RS008884 | -6.1 | 5.3 | 3.2E-07 | -6.4 | 5.3 | 3.4E-07 | -0.3 | 5.3 | 8.3E-01 | apolipoprotein d |
| RS008921 | -4.6 | 2.7 | 1.9E-09 | -3.1 | 2.7 | 4.6E-05 | 1.5 | 2.7 | 1.2E-01 | hypothetical protein |
| RS008943 | -1.6 | 3.4 | 7.4E-08 | -1.2 | 3.4 | 1.6E-04 | 0.5 | 3.4 | 2.3E-01 | inosine-uridine preferring nucleoside hydrolase-like |
| RS008944 | -1.7 | 2.4 | 1.4E-08 | -1.2 | 2.4 | 1.2E-04 | 0.6 | 2.4 | 1.8E-01 | |
| RS008975 | -6.7 | 1.3 | 1.6E-13 | -4.1 | 1.3 | 1.1E-07 | 2.6 | 1.3 | 9.3E-02 | histone h4 |
| RS009049 | -1.6 | 2.9 | 3.9E-06 | -2.0 | 2.9 | 3.9E-09 | -0.5 | 2.9 | 3.7E-01 | |
| RS009082 | -1.7 | 2.0 | 2.7E-05 | -2.0 | 2.0 | 1.6E-06 | -0.3 | 2.0 | 7.0E-01 | glycine receptor subunit alpha-2-like |
| RS009186 | -4.1 | 8.8 | 2.0E-46 | -1.8 | 8.8 | 2.1E-11 | 2.3 | 8.8 | 2.2E-16 | pancreatic triacylglycerol lipase |
| RS009200 | -1.6 | 3.0 | 9.3E-05 | -1.3 | 3.0 | 2.7E-03 | 0.3 | 3.0 | 6.0E-01 | |
| RS009226 | -1.9 | 2.0 | 5.6E-07 | -1.4 | 2.0 | 4.1E-04 | 0.5 | 2.0 | 3.5E-01 | |
| RS009272 | -1.5 | 5.5 | 3.4E-22 | -1.3 | 5.5 | 1.9E-17 | 0.2 | 5.5 | 4.6E-01 | mitochondrial sodium hydrogen exchanger 9b2 |
| RS009281 | -1.4 | 4.1 | 2.0E-09 | -1.3 | 4.1 | 1.2E-07 | 0.2 | 4.1 | 6.9E-01 | dual specificity protein kinase ttk |
| RS009292 | -1.1 | 3.6 | 9.0E-04 | -1.4 | 3.6 | 9.5E-06 | -0.3 | 3.6 | 4.6E-01 | |
| RS009375 | -1.1 | 2.4 | 5.3E-05 | -1.3 | 2.4 | 4.6E-06 | -0.2 | 2.4 | 6.9E-01 | zinc finger protein ozf-like |
| RS009403 | -1.5 | 1.6 | 4.9E-03 | -1.4 | 1.6 | 7.8E-03 | 0.1 | 1.6 | 9.4E-01 | paired box |
| RS009441 | -6.7 | 0.3 | 5.9E-06 | -6.5 | 0.3 | 1.5E-05 | 0.2 | 0.3 | 1.0E+00 | |
| RS009444 | -4.7 | 0.0 | 1.3E-05 | -6.0 | 0.0 | 3.1E-06 | -1.2 | 0.0 | 4.9E-01 | e3 ubiquitin-protein ligase siah1b |
| RS009445 | -6.4 | 0.1 | 2.4E-11 | -6.2 | 0.1 | 7.3E-10 | 0.2 | 0.1 | 1.0E+00 | e3 ubiquitin-protein ligase sina |
| RS009606 | -3.0 | 0.7 | 4.8E-07 | -3.6 | 0.7 | 1.2E-07 | -0.6 | 0.7 | 6.8E-01 | t-box transcription factor tbx6-like |
| RS009727 | -3.8 | 3.3 | 3.8E-12 | -1.6 | 3.3 | 1.5E-03 | 2.2 | 3.3 | 3.5E-04 | |
| RS009738 | -2.0 | 1.3 | 5.1E-05 | -1.3 | 1.3 | 7.3E-03 | 0.7 | 1.3 | 3.9E-01 | aristaless-related homeobox protein |
| RS010016 | -1.2 | 1.4 | 1.4E-03 | -1.6 | 1.4 | 6.4E-05 | -0.4 | 1.4 | 5.7E-01 | oocyte zinc finger protein 6-like |
| RS010030 | -1.6 | 3.6 | 1.3E-07 | -1.5 | 3.6 | 7.5E-07 | 0.1 | 3.6 | 9.0E-01 | |
| RS010050 | -5.1 | 1.2 | 7.3E-25 | -5.7 | 1.2 | 1.9E-23 | -0.5 | 1.2 | 7.5E-01 | ankyrin repeat protein |
| RS010130 | -1.3 | 3.5 | 5.6E-04 | -1.4 | 3.5 | 2.6E-04 | -0.1 | 3.5 | 8.6E-01 | |
| RS010162 | -2.1 | 5.5 | 4.2E-26 | -1.4 | 5.5 | 3.4E-13 | 0.7 | 5.5 | 6.3E-03 | cytochrome p450 6k1-like |
| RS010208 | -2.0 | 0.7 | 3.2E-04 | -2.1 | 0.7 | 1.9E-04 | -0.1 | 0.7 | 9.4E-01 | acyl- delta desaturase-like |
| RS010258 | -2.1 | 1.6 | 6.9E-06 | -1.5 | 1.6 | 1.3E-03 | 0.6 | 1.6 | 4.0E-01 | |
| RS010325 | -1.7 | 1.9 | 4.4E-03 | -2.2 | 1.9 | 1.1E-04 | -0.5 | 1.9 | 5.8E-01 | dc-stamp domain-containing protein 2 |
| RS010339 | -1.1 | 5.3 | 1.2E-08 | -1.0 | 5.3 | 3.4E-07 | 0.1 | 5.3 | 7.2E-01 | |
| RS010397 | -1.3 | 1.9 | 8.2E-04 | -1.1 | 1.9 | 6.2E-03 | 0.2 | 1.9 | 7.5E-01 | 15-hydroxyprostaglandin dehydrogenase |
| RS010485 | -2.0 | 4.5 | 5.6E-11 | -1.0 | 4.5 | 6.3E-04 | 0.9 | 4.5 | 9.3E-03 | probable chitinase 3 |
| RS010514 | -3.4 | 2.8 | 1.1E-08 | -2.3 | 2.8 | 3.4E-05 | 1.1 | 2.8 | 3.0E-01 | cytochrome p450 307a1 |
| RS010524 | -2.1 | 2.2 | 1.1E-04 | -1.4 | 2.2 | 7.6E-03 | 0.7 | 2.2 | 3.7E-01 | |
| RS010768 | -1.3 | 3.4 | 1.0E-04 | -1.1 | 3.4 | 1.2E-03 | 0.2 | 3.4 | 7.1E-01 | serine threonine-protein kinase greatwall |
| RS010782 | -1.4 | 3.7 | 6.3E-10 | -1.7 | 3.7 | 2.9E-12 | -0.2 | 3.7 | 5.5E-01 | |
| RS010831 | -1.2 | 4.7 | 6.8E-06 | -1.2 | 4.7 | 1.7E-06 | -0.1 | 4.7 | 8.8E-01 | ccr4-not transcription complex subunit 4 |
| RS010879 | -2.4 | 1.9 | 3.5E-05 | -2.1 | 1.9 | 6.0E-04 | 0.3 | 1.9 | 7.5E-01 | serine threonine-protein kinase mos |
| RS010906 | -2.1 | 6.7 | 4.7E-08 | -3.0 | 6.7 | 7.4E-14 | -0.9 | 6.7 | 7.8E-02 | tyrosine 3-monooxygenase |
| RS010971 | -1.6 | 1.9 | 9.0E-05 | -1.6 | 1.9 | 1.3E-04 | 0.0 | 1.9 | 9.9E-01 | |
| RS010992 | -1.4 | 2.7 | 5.2E-05 | -1.1 | 2.7 | 1.6E-03 | 0.3 | 2.7 | 5.9E-01 | |

| | | | | | | | | | | |
|---|---|---|---|---|---|---|---|---|---|---|
| RS011006 | -1.9 | 3.7 | 2.0E-05 | -1.2 | 3.7 | 8.3E-03 | 0.7 | 3.7 | 2.1E-01 | |
| RS011061 | -1.4 | 3.3 | 3.5E-05 | -2.0 | 3.3 | 8.8E-09 | -0.6 | 3.3 | 2.0E-01 | probable basic-leucine zipper transcription factor c |
| RS011062 | -1.7 | 1.1 | 8.6E-03 | -1.8 | 1.1 | 3.8E-03 | -0.2 | 1.1 | 8.8E-01 | |
| RS011117 | -2.5 | 4.3 | 4.0E-07 | -2.7 | 4.3 | 2.8E-08 | -0.2 | 4.3 | 7.3E-01 | cytoplasmic polyadenylation element-binding protein 1 |
| RS011143 | -2.1 | 0.0 | 2.5E-03 | -2.0 | 0.0 | 4.9E-03 | 0.1 | 0.0 | 9.2E-01 | venom carboxylesterase-6 |
| RS011249 | -3.0 | 1.0 | 1.2E-07 | -1.8 | 1.0 | 9.5E-04 | 1.2 | 1.0 | 1.6E-01 | |
| RS011280 | -1.3 | 7.0 | 1.6E-24 | -1.2 | 7.0 | 1.1E-21 | 0.1 | 7.0 | 7.3E-01 | bone morphogenetic protein 1 |
| RS011300 | -4.8 | 0.8 | 2.0E-11 | -3.4 | 0.8 | 1.3E-07 | 1.4 | 0.8 | 3.0E-01 | zinc metalloproteinase nas-4-like |
| RS011327 | -3.1 | 4.6 | 3.5E-42 | -3.5 | 4.6 | 1.3E-46 | -0.4 | 4.6 | 3.7E-01 | fatty acyl- reductase cg5065 |
| RS011395 | -1.4 | 2.7 | 4.3E-04 | -1.3 | 2.7 | 1.0E-03 | 0.1 | 2.7 | 8.6E-01 | bone morphogenetic protein |
| RS011423 | -1.3 | 5.5 | 1.1E-13 | -2.0 | 5.5 | 3.1E-28 | -0.7 | 5.5 | 1.3E-03 | low density lipoprotein receptor adapter protein 1-like |
| RS011481 | -1.3 | 3.9 | 2.2E-05 | -1.6 | 3.9 | 5.6E-07 | -0.2 | 3.9 | 6.2E-01 | |
| RS011513 | -2.0 | 2.8 | 3.9E-04 | -1.7 | 2.8 | 3.2E-03 | 0.3 | 2.8 | 6.7E-01 | protein nanos |
| RS011575 | -2.2 | 1.5 | 8.4E-05 | -3.1 | 1.5 | 1.6E-07 | -0.9 | 1.5 | 2.9E-01 | neuromedin-k receptor |
| RS011580 | -3.6 | 0.2 | 2.6E-03 | -6.2 | 0.2 | 7.2E-06 | -2.6 | 0.2 | 2.2E-01 | |
| RS011831 | -5.2 | 2.0 | 7.1E-24 | -6.2 | 2.0 | 5.9E-25 | -1.0 | 2.0 | 4.7E-01 | p protein |
| RS011857 | -1.6 | 5.5 | 5.5E-20 | -1.5 | 5.5 | 1.5E-17 | 0.1 | 5.5 | 7.6E-01 | 2-amino-3-ketobutyrate coenzyme a mitochondrial |
| RS011931 | -2.7 | 0.0 | 8.3E-04 | -3.0 | 0.0 | 7.1E-04 | -0.3 | 0.0 | 9.0E-01 | hypothetical protein |
| RS012051 | -1.1 | 4.0 | 6.5E-05 | -1.1 | 4.0 | 1.1E-04 | 0.0 | 4.0 | 9.7E-01 | sodium-independent sulfate anion transporter |
| RS012257 | -3.4 | 1.4 | 3.2E-09 | -2.8 | 1.4 | 6.8E-07 | 0.6 | 1.4 | 6.2E-01 | pr domain zinc finger protein 1 |
| RS012258 | -2.3 | -0.4 | 7.5E-03 | -3.3 | -0.4 | 1.1E-03 | -1.0 | -0.4 | 6.0E-01 | protein lin-28 homolog |
| RS012549 | -2.0 | 2.9 | 4.9E-08 | -1.0 | 2.9 | 5.7E-03 | 0.9 | 2.9 | 3.3E-02 | |
| RS012635 | -1.3 | 4.0 | 3.2E-08 | -1.3 | 4.0 | 1.7E-08 | 0.0 | 4.0 | 9.3E-01 | hypothetical protein |
| RS012707 | -1.8 | 2.8 | 1.6E-05 | -1.5 | 2.8 | 2.7E-04 | 0.3 | 2.8 | 6.7E-01 | rho gtpase-activating protein 19 |
| RS012727 | -6.0 | 0.0 | 3.1E-06 | -4.3 | 0.0 | 3.1E-04 | 1.7 | 0.0 | 3.9E-01 | histone h1 |
| RS012732 | -5.2 | 1.4 | 3.3E-06 | -3.1 | 1.4 | 3.5E-03 | 2.1 | 1.4 | 1.2E-01 | histone h1 |
| RS012785 | -2.0 | 10.5 | 3.3E-09 | -2.5 | 10.5 | 4.1E-13 | -0.5 | 10.5 | 3.0E-01 | lysosomal aspartic |
| RS013159 | -2.4 | 3.1 | 1.4E-17 | -2.3 | 3.1 | 6.4E-16 | 0.1 | 3.1 | 8.6E-01 | probable cytochrome p450 4aa1 |
| RS013175 | -1.2 | 5.5 | 9.7E-08 | -1.3 | 5.5 | 3.1E-08 | -0.1 | 5.5 | 8.6E-01 | |
| RS013177 | -1.6 | 2.6 | 4.0E-05 | -1.4 | 2.6 | 1.9E-04 | 0.1 | 2.6 | 8.4E-01 | synaptotagmin-5 |
| RS013185 | -1.0 | 5.1 | 8.0E-11 | -1.4 | 5.1 | 1.4E-18 | -0.4 | 5.1 | 6.2E-02 | spondin-1-like |
| RS013195 | -3.4 | 0.1 | 1.0E-04 | -2.4 | 0.1 | 4.1E-03 | 1.1 | 0.1 | 5.2E-01 | gamma-aminobutyric acid receptor alpha-like |
| RS013247 | -4.3 | 5.7 | 2.6E-19 | -5.1 | 5.7 | 3.3E-21 | -0.8 | 5.7 | 4.9E-01 | |
| RS013378 | -1.4 | 5.1 | 7.9E-08 | -1.0 | 5.1 | 9.0E-05 | 0.4 | 5.1 | 2.9E-01 | |
| RS013406 | -1.7 | 2.3 | 8.1E-07 | -1.7 | 2.3 | 2.0E-06 | 0.0 | 2.3 | 9.8E-01 | neuroligin- x-linked-like |
| RS013520 | -1.3 | 2.1 | 3.6E-05 | -1.0 | 2.1 | 1.2E-03 | 0.3 | 2.1 | 6.0E-01 | upf0544 protein c5orf45 homolog |
| RS013643 | -2.2 | 4.7 | 3.9E-05 | -2.5 | 4.7 | 1.4E-06 | -0.3 | 4.7 | 6.6E-01 | gustatory receptor for sugar taste 64f-like |
| RS013661 | -1.3 | 1.1 | 6.8E-03 | -1.4 | 1.1 | 4.8E-03 | -0.1 | 1.1 | 9.3E-01 | |
| RS013699 | -3.9 | -0.3 | 3.3E-05 | -3.0 | -0.3 | 7.5E-04 | 0.9 | -0.3 | 6.6E-01 | protein giant |
| RS013710 | -5.6 | -0.1 | 5.1E-06 | -3.9 | -0.1 | 3.5E-04 | 1.7 | -0.1 | 4.0E-01 | hypothetical protein |
| RS013861 | -1.4 | 1.9 | 6.4E-03 | -2.0 | 1.9 | 3.5E-04 | -0.6 | 1.9 | 5.3E-01 | basic helix-loop-helix transcription factor amos |
| RS013880 | -2.4 | -0.2 | 3.3E-03 | -3.3 | -0.2 | 6.2E-04 | -0.9 | -0.2 | 6.1E-01 | versican core protein |
| RS013956 | -3.7 | 1.3 | 1.3E-09 | -2.0 | 1.3 | 3.2E-04 | 1.7 | 1.3 | 3.1E-02 | laccase-4-like |
| RS013971 | -1.1 | 3.1 | 6.0E-03 | -1.0 | 3.1 | 9.9E-03 | 0.1 | 3.1 | 9.2E-01 | larval cuticle protein a2b-like |
| RS013977 | -8.4 | 1.5 | 8.2E-20 | -6.7 | 1.5 | 1.3E-16 | 1.6 | 1.5 | 4.0E-01 | vitellogenin |

| | | | | | | | | | | |
|---|---|---|---|---|---|---|---|---|---|---|
| RS013986 | -2.0 | 0.4 | 2.0E-04 | -1.4 | 0.4 | 5.9E-03 | 0.6 | 0.4 | 5.3E-01 | zinc finger cw-type pwwp domain protein 1-like |
| RS014221 | -1.3 | 7.8 | 1.9E-06 | -1.4 | 7.8 | 1.9E-07 | -0.1 | 7.8 | 8.0E-01 | antichymotrypsin-2-like |
| RS014229 | -1.4 | 1.8 | 1.0E-03 | -1.3 | 1.8 | 1.4E-03 | 0.0 | 1.8 | 9.9E-01 | tyrosine-protein phosphatase lar-like |
| RS014279 | -2.3 | 5.3 | 2.9E-05 | -2.1 | 5.3 | 2.0E-04 | 0.3 | 5.3 | 7.2E-01 | |
| RS014454 | -6.5 | 2.2 | 2.1E-30 | -5.6 | 2.2 | 2.1E-24 | 0.9 | 2.2 | 5.2E-01 | |
| RS014469 | -3.0 | 0.2 | 3.6E-03 | -3.4 | 0.2 | 2.8E-03 | -0.4 | 0.2 | 8.5E-01 | |
| RS014533 | -2.4 | 1.6 | 5.4E-05 | -2.8 | 1.6 | 5.3E-06 | -0.4 | 1.6 | 7.2E-01 | tetratricopeptide repeat protein 25 |
| RS014639 | -1.3 | 3.9 | 3.3E-04 | -1.8 | 3.9 | 1.0E-06 | -0.5 | 3.9 | 3.5E-01 | down syndrome cell adhesion molecule-like protein dscam2 |
| RS014743 | -2.7 | 1.8 | 2.3E-08 | -2.3 | 1.8 | 1.3E-06 | 0.4 | 1.8 | 6.6E-01 | brain-specific angiogenesis inhibitor 1 |
| RS014871 | -1.9 | 3.2 | 4.6E-15 | -1.0 | 3.2 | 2.8E-05 | 0.9 | 3.2 | 2.8E-03 | zinc finger protein zic 1-like |
| RS014971 | -1.0 | 2.8 | 5.9E-04 | -1.5 | 2.8 | 3.3E-07 | -0.5 | 2.8 | 2.1E-01 | growth differentiation factor 8 |
| RS014972 | -1.4 | 1.9 | 1.3E-03 | -2.3 | 1.9 | 4.7E-07 | -0.9 | 1.9 | 1.3E-01 | growth differentiation factor 8 |
| RS015044 | -1.6 | 2.4 | 7.1E-05 | -1.8 | 2.4 | 6.1E-06 | -0.3 | 2.4 | 7.1E-01 | t-cell acute lymphocytic leukemia protein 1 |
| RS015090 | -1.0 | 1.9 | 4.8E-03 | -1.0 | 1.9 | 4.6E-03 | 0.0 | 1.9 | 9.9E-01 | |
| RS015149 | -1.6 | 1.9 | 3.8E-03 | -1.7 | 1.9 | 3.5E-03 | -0.1 | 1.9 | 9.6E-01 | chaoptin |
| RS015423 | -2.3 | 3.5 | 1.3E-07 | -1.6 | 3.5 | 5.0E-05 | 0.6 | 3.5 | 3.7E-01 | neither inactivation nor afterpotential protein c |
| RS015515 | -1.0 | 3.4 | 2.6E-04 | -1.4 | 3.4 | 4.6E-07 | -0.4 | 3.4 | 3.0E-01 | mam and ldl-receptor class a domain-containing protein 1-like |

SI_Dataset_1
Caste-biased genes: Soldier (head)

| Gene ID | Head Reproductive v.s. Soldier | | | Head Reproductive v.s. Worker | | | Head Soldier v.s. Worker | | | Gene annotation |
|---|---|---|---|---|---|---|---|---|---|---|
| | logFC | logCPM | FDR | logFC | logCPM | FDR | logFC | logCPM | FDR | |
| RS000001 | 1.7 | 9.4 | 6.8E-21 | -0.3 | 9.4 | 2.3E-01 | -2.0 | 9.4 | 4.0E-29 | ryanodine receptor 44f |
| RS000043 | 2.6 | 3.2 | 4.8E-09 | 1.4 | 3.2 | 1.6E-02 | -1.2 | 3.2 | 9.1E-03 | |
| RS000102 | 1.4 | 4.7 | 2.9E-09 | 0.3 | 4.7 | 5.8E-01 | -1.1 | 4.7 | 3.2E-06 | serine threonine-protein kinase 17b-like |
| RS000188 | 1.2 | 4.2 | 2.1E-14 | -0.2 | 4.2 | 6.1E-01 | -1.4 | 4.2 | 1.4E-17 | death-associated protein kinase 1-like |
| RS000193 | 1.2 | 6.6 | 2.5E-19 | -0.5 | 6.6 | 6.3E-04 | -1.7 | 6.6 | 2.2E-38 | b( +)-type amino acid transporter 1 |
| RS000202 | 5.2 | 2.3 | 4.1E-50 | 1.0 | 2.3 | 1.7E-01 | -4.2 | 2.3 | 2.5E-38 | probable cytochrome p450 49a1 |
| RS000208 | 2.8 | 4.6 | 1.0E-13 | 1.7 | 4.6 | 2.4E-05 | -1.1 | 4.6 | 9.2E-03 | |
| RS000216 | 1.2 | 2.1 | 8.2E-04 | -0.9 | 2.1 | 1.6E-01 | -2.1 | 2.1 | 1.4E-07 | innexin shaking-b |
| RS000285 | 1.5 | 5.1 | 7.9E-15 | -0.2 | 5.1 | 6.6E-01 | -1.7 | 5.1 | 6.2E-18 | facilitated trehalose transporter tret1-like |
| RS000312 | 1.9 | 7.5 | 2.7E-18 | 0.8 | 7.5 | 1.5E-03 | -1.1 | 7.5 | 1.1E-06 | |
| RS000328 | 1.1 | 5.2 | 3.0E-06 | -0.3 | 5.2 | 4.0E-01 | -1.4 | 5.2 | 1.1E-09 | farnesyl pyrophosphate synthase |
| RS000365 | 3.0 | 12.6 | 4.8E-33 | -0.1 | 12.6 | 9.1E-01 | -3.1 | 12.6 | 2.1E-34 | calcium-transporting atpase sarcoplasmic endoplasmic reticulum type |
| RS000381 | 2.1 | 5.4 | 2.1E-41 | 0.0 | 5.4 | 9.9E-01 | -2.1 | 5.4 | 4.6E-41 | |
| RS000382 | 1.8 | 3.2 | 4.5E-07 | -0.2 | 3.2 | 7.6E-01 | -2.1 | 3.2 | 1.2E-08 | |
| RS000609 | 1.1 | 7.7 | 1.5E-15 | -0.2 | 7.7 | 2.7E-01 | -1.3 | 7.7 | 1.4E-22 | muscle calcium channel subunit alpha-1 |
| RS000669 | 1.2 | 5.4 | 1.4E-09 | 0.2 | 5.4 | 7.0E-01 | -1.0 | 5.4 | 3.2E-07 | wd repeat-containing protein 26 |
| RS000700 | 1.1 | 7.9 | 2.4E-13 | -0.1 | 7.9 | 8.9E-01 | -1.2 | 7.9 | 1.8E-14 | synaptic vesicle membrane protein vat-1 homolog-like |
| RS000836 | 1.4 | 4.8 | 2.3E-18 | 0.4 | 4.8 | 9.3E-02 | -1.0 | 4.8 | 6.6E-10 | |
| RS000837 | 1.7 | 6.5 | 1.7E-21 | 0.2 | 6.5 | 4.6E-01 | -1.4 | 6.5 | 5.1E-16 | excitatory amino acid transporter 1-like |
| RS000842 | 2.0 | 2.9 | 2.5E-08 | -1.2 | 2.9 | 2.0E-02 | -3.2 | 2.9 | 2.0E-16 | cytochrome p450 4c1-like |
| RS000843 | 2.7 | 3.6 | 9.8E-09 | -0.5 | 3.6 | 5.8E-01 | -3.2 | 3.6 | 4.1E-11 | cytochrome p450-like protein |
| RS000909 | 2.0 | 3.1 | 8.1E-12 | 0.4 | 3.1 | 5.2E-01 | -1.7 | 3.1 | 3.6E-08 | mitochondrial uncoupling protein 2-like |
| RS000919 | 1.9 | 7.0 | 5.0E-28 | 0.3 | 7.0 | 1.9E-01 | -1.5 | 7.0 | 3.4E-19 | 3-hydroxy-3-methylglutaryl-coenzyme a reductase |
| RS000936 | 2.2 | 9.1 | 8.2E-26 | -2.4 | 9.1 | 1.1E-29 | -4.5 | 9.1 | 7.4E-92 | |
| RS000947 | 2.9 | 9.3 | 1.9E-44 | 0.5 | 9.3 | 7.1E-02 | -2.4 | 9.3 | 7.5E-32 | protein disulfide-isomerase |
| RS000948 | 2.8 | 7.1 | 2.9E-40 | 0.3 | 7.1 | 3.9E-01 | -2.5 | 7.1 | 9.1E-33 | protein disulfide-isomerase a5 |
| RS001012 | 1.5 | 5.7 | 3.0E-22 | 0.0 | 5.7 | 1.0E+00 | -1.5 | 5.7 | 8.0E-22 | sodium calcium exchanger 2 |
| RS001088 | 1.0 | 4.0 | 4.7E-10 | 0.0 | 4.0 | 9.6E-01 | -1.1 | 4.0 | 6.5E-10 | rims-binding protein 2-like |
| RS001106 | 1.7 | 6.2 | 1.4E-21 | 0.2 | 6.2 | 4.8E-01 | -1.5 | 6.2 | 3.1E-16 | phosphatidylinositol transfer protein alpha isoform |
| RS001172 | 1.3 | 1.5 | 6.7E-05 | 0.1 | 1.5 | 9.4E-01 | -1.3 | 1.5 | 3.8E-04 | cadherin-89d |
| RS001195 | 3.3 | 6.0 | 1.6E-28 | 2.2 | 6.0 | 1.3E-13 | -1.1 | 6.0 | 4.7E-04 | |
| RS001199 | 3.4 | 2.0 | 8.3E-12 | 1.9 | 2.0 | 1.4E-03 | -1.5 | 2.0 | 2.5E-03 | |
| RS001228 | 3.1 | 5.9 | 3.7E-66 | 0.0 | 5.9 | 9.9E-01 | -3.0 | 5.9 | 6.7E-65 | von willebrand factor type egf and pentraxin domain-containing protein |
| RS001241 | 3.2 | 12.2 | 9.8E-68 | 1.1 | 12.2 | 3.5E-09 | -2.1 | 12.2 | 7.8E-32 | arginine kinase |

| | | | | | | | | | | |
|---|---|---|---|---|---|---|---|---|---|---|
| RS001285 | 1.8 | 4.3 | 8.7E-13 | 0.5 | 4.3 | 1.8E-01 | -1.3 | 4.3 | 6.1E-07 | transcriptional-regulating factor 1 |
| RS001347 | 2.6 | 3.7 | 2.4E-09 | 0.1 | 3.7 | 9.5E-01 | -2.5 | 3.7 | 1.1E-08 | |
| RS001348 | 2.2 | 6.2 | 1.7E-23 | 0.0 | 6.2 | 9.8E-01 | -2.3 | 6.2 | 1.2E-23 | muscle m-line assembly protein unc-89 |
| RS001352 | 1.6 | 7.1 | 7.1E-37 | 0.2 | 7.1 | 3.8E-01 | -1.4 | 7.1 | 9.7E-29 | dual 3 -cyclic-amp and -gmp phosphodiesterase 11-like |
| RS001421 | 1.3 | 4.6 | 5.2E-03 | -0.5 | 4.6 | 5.1E-01 | -1.8 | 4.6 | 8.7E-05 | acyl- synthetase family member mitochondrial-like |
| RS001531 | 3.5 | 2.7 | 2.3E-17 | 0.7 | 2.7 | 3.3E-01 | -2.8 | 2.7 | 4.5E-12 | clavesin-2 |
| RS001581 | 1.0 | 5.0 | 7.1E-08 | -0.4 | 5.0 | 1.8E-01 | -1.4 | 5.0 | 1.5E-13 | |
| RS001613 | 1.8 | 5.5 | 5.8E-27 | -0.1 | 5.5 | 7.9E-01 | -1.9 | 5.5 | 1.3E-29 | |
| RS001614 | 1.6 | 7.3 | 2.4E-27 | -0.1 | 7.3 | 8.3E-01 | -1.7 | 7.3 | 8.8E-30 | tyrosine-protein phosphatase non-receptor type 13-like |
| RS001624 | 1.9 | 2.4 | 6.3E-10 | -3.6 | 2.4 | 6.9E-15 | -5.5 | 2.4 | 3.2E-40 | |
| RS001683 | 1.1 | 5.5 | 1.4E-12 | 0.0 | 5.5 | 9.9E-01 | -1.1 | 5.5 | 1.4E-12 | inactive phospholipase c-like |
| RS001705 | 1.9 | 5.8 | 6.2E-16 | 0.6 | 5.8 | 7.5E-02 | -1.3 | 5.8 | 3.6E-08 | tubulointerstitial nephritis antigen-like |
| RS001888 | 1.1 | 5.8 | 2.2E-11 | 0.0 | 5.8 | 9.7E-01 | -1.2 | 5.8 | 1.3E-11 | |
| RS001958 | 1.3 | 10.8 | 1.8E-10 | 0.2 | 10.8 | 6.6E-01 | -1.1 | 10.8 | 7.5E-08 | muscle m-line assembly protein unc-89 |
| RS002051 | 6.8 | 10.3 | 1.1E-148 | 0.0 | 10.3 | 9.9E-01 | -6.7 | 10.3 | 1.1E-147 | troponin |
| RS002055 | 1.2 | 1.9 | 5.2E-04 | 0.2 | 1.9 | 8.3E-01 | -1.0 | 1.9 | 5.4E-03 | |
| RS002057 | 1.6 | 5.6 | 1.4E-30 | -0.2 | 5.6 | 4.1E-01 | -1.8 | 5.6 | 1.3E-37 | zinc finger protein xfin-like |
| RS002058 | 1.2 | 7.4 | 4.6E-21 | 0.0 | 7.4 | 9.3E-01 | -1.1 | 7.4 | 7.1E-20 | dnaj homolog subfamily a member 2 |
| RS002180 | 1.7 | 3.9 | 1.4E-17 | -0.4 | 3.9 | 1.8E-01 | -2.2 | 3.9 | 2.7E-25 | serine threonine-protein phosphatase rdgc |
| RS002203 | 1.8 | 3.4 | 3.3E-10 | 0.3 | 3.4 | 5.8E-01 | -1.5 | 3.4 | 4.5E-07 | atp-binding cassette sub-family g member 4 |
| RS002211 | 2.7 | 3.3 | 2.8E-07 | 0.7 | 3.3 | 4.3E-01 | -2.0 | 3.3 | 1.8E-04 | |
| RS002240 | 3.5 | 6.3 | 7.0E-71 | 0.1 | 6.3 | 7.6E-01 | -3.4 | 6.3 | 1.9E-65 | probable pyruvate dehydrogenase e1 component subunit mitochondrial |
| RS002265 | 1.6 | 6.7 | 6.6E-05 | 0.0 | 6.7 | 9.8E-01 | -1.6 | 6.7 | 2.5E-04 | probable rna-binding protein 46 |
| RS002448 | 3.5 | 3.4 | 2.0E-11 | -2.3 | 3.4 | 1.5E-03 | -5.9 | 3.4 | 3.3E-22 | fatty acyl- reductase cg5065 |
| RS002601 | 1.8 | 6.7 | 1.3E-16 | 0.5 | 6.7 | 6.9E-02 | -1.2 | 6.7 | 1.4E-08 | |
| RS002602 | 1.8 | 5.5 | 4.7E-11 | 0.5 | 5.5 | 2.4E-01 | -1.3 | 5.5 | 3.4E-06 | |
| RS002685 | 1.1 | 5.3 | 1.1E-19 | 0.0 | 5.3 | 8.9E-01 | -1.1 | 5.3 | 1.0E-20 | calcium uptake protein mitochondrial |
| RS002693 | 3.9 | 1.4 | 1.0E-05 | 1.2 | 1.4 | 4.5E-01 | -2.7 | 1.4 | 2.3E-03 | |
| RS002694 | 3.4 | 1.6 | 5.6E-05 | 1.2 | 1.6 | 4.3E-01 | -2.3 | 1.6 | 9.1E-03 | protein takeout |
| RS002695 | 1.1 | 6.6 | 8.8E-09 | -0.5 | 6.6 | 9.3E-02 | -1.6 | 6.6 | 4.2E-16 | |
| RS002790 | 4.2 | 1.3 | 1.5E-20 | 0.4 | 1.3 | 8.3E-01 | -3.9 | 1.3 | 1.6E-17 | serine protease 33-like |
| RS002844 | 3.2 | 2.9 | 3.5E-14 | 1.7 | 2.9 | 4.0E-04 | -1.5 | 2.9 | 4.7E-04 | isoform i |
| RS002918 | 2.4 | 1.6 | 2.3E-03 | -0.1 | 1.6 | 9.9E-01 | -2.4 | 1.6 | 3.6E-03 | |
| RS003052 | 5.2 | 5.9 | 7.2E-09 | 0.1 | 5.9 | 9.6E-01 | -5.1 | 5.9 | 1.6E-08 | integrase core domain protein |
| RS003131 | 1.7 | 4.4 | 2.7E-34 | 0.0 | 4.4 | 9.8E-01 | -1.7 | 4.4 | 1.2E-32 | |
| RS003132 | 1.8 | 6.0 | 2.7E-43 | 0.1 | 6.0 | 7.2E-01 | -1.7 | 6.0 | 3.3E-38 | serine threonine-protein phosphatase 2b catalytic subunit 2-like |
| RS003189 | 2.0 | 0.5 | 1.4E-04 | -0.6 | 0.5 | 6.3E-01 | -2.6 | 0.5 | 5.9E-06 | zinc metalloproteinase nas-15 |
| RS003197 | 1.6 | 6.9 | 6.8E-15 | 0.4 | 6.9 | 2.9E-01 | -1.3 | 6.9 | 2.4E-09 | 17-beta-hydroxysteroid dehydrogenase type 6 |

| | | | | | | | | | | |
|---|---|---|---|---|---|---|---|---|---|---|
| RS003214 | 1.8 | 1.9 | 2.6E-12 | -0.1 | 1.9 | 9.3E-01 | -1.9 | 1.9 | 4.0E-12 | potassium voltage-gated channel subfamily kqt member 4-like |
| RS003216 | 2.1 | 0.5 | 1.4E-06 | 0.6 | 0.5 | 4.9E-01 | -1.5 | 0.5 | 9.2E-04 | potassium voltage-gated channel subfamily kqt member 4 |
| RS003217 | 3.5 | 3.3 | 3.3E-03 | 0.0 | 3.3 | 1.0E+00 | -3.5 | 3.3 | 8.0E-03 | |
| RS003255 | 2.7 | 5.0 | 2.4E-59 | -0.4 | 5.0 | 2.0E-01 | -3.0 | 5.0 | 8.0E-71 | rna pseudouridylate synthase domain-containing protein 2-like |
| RS003372 | 1.8 | 2.5 | 3.9E-08 | 0.6 | 2.5 | 2.0E-01 | -1.1 | 2.5 | 9.0E-04 | udp-glucuronosyltransferase 2c1-like |
| RS003433 | 3.7 | 0.7 | 3.8E-09 | 1.1 | 0.7 | 3.5E-01 | -2.6 | 0.7 | 1.9E-05 | retrovirus-related pol polyprotein from transposon |
| RS003527 | 1.4 | 6.3 | 8.4E-34 | 0.1 | 6.3 | 5.6E-01 | -1.3 | 6.3 | 1.4E-27 | protein lmbr1l |
| RS003552 | 1.4 | 5.2 | 6.5E-09 | 0.0 | 5.2 | 1.0E+00 | -1.4 | 5.2 | 1.0E-08 | xanthine dehydrogenase oxidase-like |
| RS003669 | 1.3 | 2.7 | 5.2E-06 | -0.6 | 2.7 | 2.2E-01 | -1.9 | 2.7 | 2.6E-10 | retinol dehydrogenase 11-like |
| RS003692 | 2.4 | 6.9 | 5.4E-90 | 0.2 | 6.9 | 2.5E-01 | -2.1 | 6.9 | 5.5E-75 | |
| RS003693 | 2.7 | 8.0 | 8.4E-78 | 0.3 | 8.0 | 1.9E-01 | -2.4 | 8.0 | 1.9E-63 | microtubule-associated protein futsch |
| RS003709 | 3.3 | 5.9 | 6.1E-23 | -2.2 | 5.9 | 2.0E-09 | -5.5 | 5.9 | 4.3E-49 | cytochrome p450 4v2 |
| RS003758 | 1.8 | 5.5 | 1.9E-14 | 0.8 | 5.5 | 7.1E-03 | -1.0 | 5.5 | 4.5E-05 | chitin deacetylase 3 |
| RS003832 | 4.2 | 3.6 | 4.1E-24 | 1.2 | 3.6 | 1.8E-01 | -3.0 | 3.6 | 1.3E-15 | |
| RS003868 | 1.7 | 3.9 | 1.6E-08 | 0.3 | 3.9 | 6.8E-01 | -1.4 | 3.9 | 3.1E-06 | |
| RS003986 | 1.3 | 2.8 | 8.5E-11 | 0.0 | 2.8 | 9.8E-01 | -1.4 | 2.8 | 2.1E-10 | chloride intracellular channel exl-1 |
| RS004040 | 1.1 | 7.7 | 5.5E-19 | -0.1 | 7.7 | 8.1E-01 | -1.1 | 7.7 | 2.6E-21 | poly -binding protein 3 |
| RS004044 | 1.1 | 3.1 | 6.4E-06 | 0.0 | 3.1 | 1.0E+00 | -1.1 | 3.1 | 1.3E-05 | glutamate-gated chloride channel |
| RS004058 | 2.4 | 5.2 | 4.6E-43 | 0.1 | 5.2 | 7.5E-01 | -2.3 | 5.2 | 1.9E-38 | sh3 and cysteine-rich domain-containing protein 2 |
| RS004086 | 1.6 | 5.1 | 1.1E-16 | 0.1 | 5.1 | 8.8E-01 | -1.5 | 5.1 | 5.0E-15 | protein phosphatase 1 regulatory subunit 3c-b |
| RS004177 | 1.3 | 7.4 | 5.2E-13 | 0.1 | 7.4 | 7.2E-01 | -1.2 | 7.4 | 2.4E-10 | |
| RS004195 | 1.9 | 2.8 | 1.9E-15 | 0.1 | 2.8 | 9.2E-01 | -1.8 | 2.8 | 7.2E-14 | zinc finger protein 346-like |
| RS004231 | 1.7 | 10.6 | 1.4E-17 | 0.2 | 10.6 | 5.8E-01 | -1.5 | 10.6 | 1.2E-13 | |
| RS004232 | 1.8 | 8.6 | 1.6E-32 | 0.6 | 8.6 | 1.6E-03 | -1.2 | 8.6 | 5.5E-16 | |
| RS004289 | 1.1 | 8.7 | 1.3E-08 | -0.1 | 8.7 | 8.9E-01 | -1.2 | 8.7 | 1.7E-09 | peroxidase-like |
| RS004341 | 2.8 | 4.1 | 6.9E-28 | 0.2 | 4.1 | 8.0E-01 | -2.6 | 4.1 | 5.2E-25 | fork head domain-containing protein crocodile |
| RS004477 | 1.5 | 2.9 | 1.1E-06 | 0.5 | 2.9 | 3.6E-01 | -1.0 | 2.9 | 1.8E-03 | protein apterous-like |
| RS004485 | 1.0 | 1.3 | 1.6E-03 | 0.0 | 1.3 | 9.9E-01 | -1.0 | 1.3 | 3.9E-03 | |
| RS004491 | 2.6 | 5.0 | 1.6E-26 | 1.1 | 5.0 | 7.3E-05 | -1.5 | 5.0 | 5.2E-10 | |
| RS004492 | 2.5 | 3.3 | 6.5E-21 | 1.1 | 3.3 | 1.1E-03 | -1.5 | 3.3 | 3.7E-08 | |
| RS004500 | 1.4 | 6.4 | 3.5E-16 | -0.6 | 6.4 | 2.6E-03 | -2.0 | 6.4 | 1.2E-31 | basement membrane-specific heparan sulfate proteoglycan core |
| RS004526 | 1.9 | 3.4 | 2.6E-31 | -0.1 | 3.4 | 9.0E-01 | -1.9 | 3.4 | 2.0E-31 | zinc finger cchc domain-containing protein 24-like |
| RS004527 | 1.3 | 4.5 | 1.8E-08 | 0.2 | 4.5 | 6.6E-01 | -1.1 | 4.5 | 4.9E-06 | |
| RS004547 | 7.0 | 3.9 | 4.3E-141 | -1.1 | 3.9 | 4.3E-01 | -8.1 | 3.9 | 3.2E-139 | structural constituent of |
| RS004548 | 3.4 | 3.3 | 9.5E-21 | 1.8 | 3.3 | 5.4E-06 | -1.6 | 3.3 | 1.2E-05 | cuticle protein 8 |
| RS004578 | 3.2 | 3.6 | 9.9E-46 | 0.5 | 3.6 | 1.9E-01 | -2.7 | 3.6 | 4.3E-34 | protein vein |
| RS004623 | 1.1 | 6.3 | 2.6E-06 | -0.8 | 6.3 | 3.4E-03 | -1.9 | 6.3 | 1.6E-16 | beta-glucosidase |
| RS004634 | 1.3 | 4.2 | 5.5E-10 | 0.2 | 4.2 | 6.1E-01 | -1.1 | 4.2 | 5.4E-07 | protein pellino |

| | | | | | | | | | | |
|---|---|---|---|---|---|---|---|---|---|---|
| RS004652 | 1.3 | 5.7 | 1.2E-18 | 0.1 | 5.7 | 6.7E-01 | -1.2 | 5.7 | 6.2E-15 | src kinase-associated phosphoprotein 2 |
| RS004890 | 1.2 | 5.6 | 2.0E-08 | -2.1 | 5.6 | 1.1E-20 | -3.3 | 5.6 | 5.1E-49 | |
| RS004891 | 1.6 | 3.3 | 4.6E-09 | -3.3 | 3.3 | 6.7E-19 | -5.0 | 3.3 | 2.9E-45 | |
| RS004931 | 1.3 | 6.3 | 1.4E-15 | 0.0 | 6.3 | 1.0E+00 | -1.3 | 6.3 | 3.9E-15 | inactive hydroxysteroid dehydrogenase-like protein 1 |
| RS005033 | 2.9 | 6.2 | 5.2E-26 | 0.1 | 6.2 | 8.7E-01 | -2.8 | 6.2 | 3.6E-24 | hydroxymethylglutaryl- synthase 1 |
| RS005034 | 2.9 | 6.1 | 8.3E-35 | 0.6 | 6.1 | 9.4E-02 | -2.4 | 6.1 | 4.3E-24 | |
| RS005088 | 1.2 | 2.0 | 8.1E-04 | 0.2 | 2.0 | 8.1E-01 | -1.0 | 2.0 | 8.4E-03 | |
| RS005096 | 2.3 | 4.4 | 1.6E-20 | -0.2 | 4.4 | 7.8E-01 | -2.5 | 4.4 | 1.1E-22 | |
| RS005133 | 2.2 | 4.0 | 1.4E-15 | 0.9 | 4.0 | 9.1E-03 | -1.3 | 4.0 | 5.0E-06 | activating signal cointegrator 1 complex subunit 2 homolog |
| RS005160 | 1.9 | 5.3 | 2.1E-30 | -0.3 | 5.3 | 3.1E-01 | -2.2 | 5.3 | 1.9E-38 | rna polymerase-associated protein rtf1 |
| RS005244 | 2.0 | 8.9 | 4.3E-24 | 0.3 | 8.9 | 2.8E-01 | -1.7 | 8.9 | 5.3E-17 | sparc |
| RS005458 | 2.0 | 9.4 | 2.1E-32 | -0.2 | 9.4 | 5.2E-01 | -2.2 | 9.4 | 1.4E-38 | trichohyalin |
| RS005488 | 2.2 | 8.3 | 4.0E-23 | 0.4 | 8.3 | 2.8E-01 | -1.8 | 8.3 | 3.0E-16 | basement membrane-specific heparan sulfate proteoglycan core protein |
| RS005696 | 2.1 | 5.8 | 1.1E-32 | 1.0 | 5.8 | 4.2E-07 | -1.1 | 5.8 | 2.3E-10 | dopamine n-acetyltransferase-like |
| RS005869 | 1.6 | 4.2 | 2.9E-23 | 0.6 | 4.2 | 5.6E-03 | -1.0 | 4.2 | 2.1E-10 | |
| RS005957 | 1.2 | 3.7 | 1.2E-08 | 0.1 | 3.7 | 9.1E-01 | -1.2 | 3.7 | 1.4E-07 | dipeptidyl aminopeptidase-like protein 6 |
| RS005962 | 1.5 | 8.4 | 2.0E-22 | 0.0 | 8.4 | 9.4E-01 | -1.5 | 8.4 | 1.7E-21 | |
| RS005987 | 1.7 | 5.1 | 3.4E-09 | 0.5 | 5.1 | 3.1E-01 | -1.2 | 5.1 | 4.1E-05 | ral guanine nucleotide dissociation stimulator |
| RS006140 | 1.9 | 2.4 | 9.6E-14 | -0.2 | 2.4 | 7.1E-01 | -2.1 | 2.4 | 9.1E-16 | two pore potassium channel protein sup-9 |
| RS006147 | 1.1 | 4.1 | 1.4E-08 | -0.2 | 4.1 | 6.3E-01 | -1.3 | 4.1 | 4.8E-11 | hypothetical protein |
| RS006270 | 1.2 | 3.2 | 5.2E-05 | -0.6 | 3.2 | 1.7E-01 | -1.8 | 3.2 | 1.8E-09 | organic cation transporter 1 |
| RS006354 | 2.0 | 9.9 | 4.6E-30 | -0.1 | 9.9 | 7.7E-01 | -2.1 | 9.9 | 2.8E-33 | |
| RS006382 | 1.4 | 5.7 | 1.6E-10 | 0.4 | 5.7 | 2.4E-01 | -1.0 | 5.7 | 1.2E-05 | chitooligosaccharidolytic beta-n-acetylglucosaminidase |
| RS006393 | 2.0 | 6.3 | 8.8E-38 | 0.3 | 6.3 | 3.2E-01 | -1.7 | 6.3 | 4.5E-29 | alpha-tocopherol transfer |
| RS006577 | 1.1 | 3.5 | 4.1E-07 | -0.2 | 3.5 | 7.4E-01 | -1.3 | 3.5 | 1.0E-08 | regulator of g-protein signaling 7 |
| RS006648 | 3.1 | 5.7 | 8.1E-38 | -0.1 | 5.7 | 9.0E-01 | -3.2 | 5.7 | 3.5E-39 | voltage-dependent l-type calcium channel subunit beta-2 |
| RS006650 | 2.0 | 3.7 | 1.5E-17 | -0.1 | 3.7 | 8.6E-01 | -2.1 | 3.7 | 7.6E-19 | |
| RS006676 | 3.7 | 5.7 | 1.7E-99 | 0.4 | 5.7 | 1.8E-01 | -3.3 | 5.7 | 2.1E-83 | adenylate kinase isoenzyme 1 |
| RS006696 | 1.6 | 6.1 | 4.1E-14 | 0.1 | 6.1 | 8.0E-01 | -1.5 | 6.1 | 5.0E-12 | uncharacterized f-box lrr-repeat protein |
| RS006743 | 1.3 | 1.6 | 2.6E-03 | -0.1 | 1.6 | 9.5E-01 | -1.3 | 1.6 | 2.2E-03 | |
| RS006884 | 1.1 | 2.2 | 2.8E-05 | 0.1 | 2.2 | 9.2E-01 | -1.0 | 2.2 | 2.3E-04 | |
| RS006977 | 1.2 | 5.2 | 1.2E-17 | 0.2 | 5.2 | 4.4E-01 | -1.0 | 5.2 | 3.0E-12 | synaptotagmin-1-like |
| RS007128 | 1.0 | 6.1 | 1.2E-16 | 0.0 | 6.1 | 1.0E+00 | -1.0 | 6.1 | 2.2E-16 | probable multidrug resistance-associated protein lethal 03659 |
| RS007137 | 2.0 | 0.1 | 3.2E-04 | -0.7 | 0.1 | 6.3E-01 | -2.7 | 0.1 | 1.6E-05 | follicle cell protein 3c-1 |
| RS007154 | 1.0 | 5.2 | 2.9E-13 | -0.4 | 5.2 | 2.7E-02 | -1.4 | 5.2 | 2.9E-24 | neuroligin- x-linked |
| RS007161 | 1.4 | 5.0 | 9.2E-21 | 0.2 | 5.0 | 5.2E-01 | -1.3 | 5.0 | 1.4E-15 | methionine-r-sulfoxide reductase b1 |
| RS007182 | 1.5 | 4.7 | 3.5E-26 | 0.1 | 4.7 | 8.1E-01 | -1.4 | 4.7 | 6.8E-23 | receptor-type tyrosine-protein phosphatase kappa |
| RS007183 | 1.3 | 4.9 | 7.7E-24 | 0.1 | 4.9 | 8.8E-01 | -1.2 | 4.9 | 1.7E-21 | |

| | | | | | | | | | | |
|---|---|---|---|---|---|---|---|---|---|---|
| RS007254 | 1.9 | 6.1 | 4.4E-07 | 0.0 | 6.1 | 9.6E-01 | -1.9 | 6.1 | 3.6E-07 | |
| RS007262 | 1.1 | 3.2 | 1.1E-04 | -0.2 | 3.2 | 7.7E-01 | -1.3 | 3.2 | 9.6E-06 | protein hos4-like |
| RS007263 | 1.2 | 1.4 | 3.5E-03 | -0.3 | 1.4 | 7.3E-01 | -1.5 | 1.4 | 4.4E-04 | |
| RS007344 | 1.4 | 12.4 | 9.6E-14 | 0.1 | 12.4 | 7.8E-01 | -1.3 | 12.4 | 1.5E-11 | tropomyosin-1 |
| RS007346 | 1.5 | 12.1 | 3.4E-15 | 0.0 | 12.1 | 9.6E-01 | -1.5 | 12.1 | 1.4E-15 | tropomyosin |
| RS007374 | 1.2 | 13.1 | 2.6E-09 | 0.2 | 13.1 | 6.1E-01 | -1.0 | 13.1 | 1.3E-06 | troponin skeletal muscle |
| RS007404 | 1.3 | 2.3 | 7.2E-03 | -0.2 | 2.3 | 8.5E-01 | -1.5 | 2.3 | 3.0E-03 | sodium-independent sulfate anion transporter |
| RS007435 | 2.4 | 5.2 | 1.0E-27 | 1.2 | 5.2 | 9.5E-07 | -1.2 | 5.2 | 2.9E-08 | a disintegrin and metalloproteinase with thrombospondin motifs 3-like |
| RS007451 | 3.4 | 10.7 | 1.1E-50 | -0.5 | 10.7 | 1.3E-01 | -3.8 | 10.7 | 4.9E-63 | |
| RS007452 | 3.8 | 7.4 | 2.1E-82 | -0.3 | 7.4 | 2.5E-01 | -4.2 | 7.4 | 3.7E-94 | |
| RS007480 | 3.1 | 4.7 | 2.6E-10 | -0.8 | 4.7 | 2.9E-01 | -3.9 | 4.7 | 1.3E-14 | geranylgeranyl pyrophosphate synthase |
| RS007516 | 2.0 | 15.4 | 1.3E-13 | 0.7 | 15.4 | 7.0E-02 | -1.4 | 15.4 | 1.2E-06 | muscle |
| RS007542 | 1.1 | 3.3 | 2.4E-07 | 0.0 | 3.3 | 9.4E-01 | -1.2 | 3.3 | 1.5E-07 | ski oncogene |
| RS007614 | 1.2 | 4.9 | 7.8E-10 | -0.3 | 4.9 | 4.7E-01 | -1.4 | 4.9 | 9.8E-14 | facilitated trehalose transporter tret1-like |
| RS007615 | 1.3 | 6.5 | 2.3E-10 | -0.1 | 6.5 | 8.2E-01 | -1.4 | 6.5 | 6.0E-12 | facilitated trehalose transporter tret1-like |
| RS007700 | 1.6 | 5.2 | 3.9E-28 | 0.1 | 5.2 | 8.8E-01 | -1.5 | 5.2 | 1.4E-25 | serine threonine-protein kinase plk1-like |
| RS007932 | 2.5 | 5.6 | 1.3E-05 | 1.0 | 5.6 | 3.8E-01 | -1.5 | 5.6 | 7.4E-03 | |
| RS007933 | 1.6 | 3.9 | 2.1E-05 | 0.3 | 3.9 | 7.1E-01 | -1.3 | 3.9 | 9.9E-04 | |
| RS007934 | 3.0 | 2.4 | 4.4E-09 | 0.0 | 2.4 | 9.8E-01 | -2.9 | 2.4 | 1.4E-08 | |
| RS008012 | 1.1 | 3.5 | 1.6E-06 | -0.1 | 3.5 | 8.0E-01 | -1.2 | 3.5 | 1.3E-07 | protein goliath |
| RS008064 | 1.3 | 5.7 | 6.1E-08 | -0.3 | 5.7 | 5.8E-01 | -1.6 | 5.7 | 1.1E-10 | aminopeptidase n |
| RS008106 | 5.4 | 4.3 | 2.2E-06 | -0.6 | 4.3 | 8.0E-01 | -6.0 | 4.3 | 4.2E-07 | nose resistant to fluoxetine protein 6-like |
| RS008107 | 3.6 | 5.8 | 5.9E-40 | -0.2 | 5.8 | 8.1E-01 | -3.8 | 5.8 | 2.3E-42 | nose resistant to fluoxetine protein 6 |
| RS008183 | 1.2 | 6.2 | 2.3E-22 | -0.7 | 6.2 | 4.3E-07 | -1.9 | 6.2 | 1.1E-51 | transmembrane protein 114 |
| RS008238 | 1.9 | 4.6 | 7.3E-39 | 0.5 | 4.6 | 1.4E-02 | -1.4 | 4.6 | 1.2E-22 | |
| RS008283 | 2.6 | 4.8 | 5.7E-86 | -0.2 | 4.8 | 3.9E-01 | -2.8 | 4.8 | 8.4E-95 | |
| RS008284 | 6.5 | 6.6 | 6.5E-221 | 0.5 | 6.6 | 1.1E-01 | -6.0 | 6.6 | 9.4E-198 | protein |
| RS008285 | 1.6 | 7.1 | 3.6E-33 | -0.4 | 7.1 | 1.8E-02 | -2.0 | 7.1 | 2.8E-50 | protein isoform a-like |
| RS008380 | 1.9 | 6.1 | 2.1E-10 | 0.9 | 6.1 | 1.2E-02 | -1.0 | 6.1 | 2.2E-03 | |
| RS008567 | 2.4 | 6.9 | 1.3E-65 | 0.5 | 6.9 | 1.1E-02 | -2.0 | 6.9 | 9.5E-45 | glycerol-3-phosphate mitochondrial |
| RS008591 | 1.9 | 7.6 | 1.1E-46 | 0.0 | 7.6 | 9.1E-01 | -1.9 | 7.6 | 2.7E-48 | myosin light chain smooth muscle-like |
| RS008592 | 1.8 | 4.8 | 1.2E-15 | -0.3 | 4.8 | 4.3E-01 | -2.1 | 4.8 | 1.1E-20 | myosin light chain smooth muscle-like |
| RS008617 | 2.0 | 4.9 | 3.9E-38 | 0.3 | 4.9 | 2.9E-01 | -1.7 | 4.9 | 9.7E-29 | proton-coupled amino acid transporter 1 |
| RS008620 | 1.6 | 2.7 | 1.2E-10 | 0.3 | 2.7 | 6.5E-01 | -1.4 | 2.7 | 1.4E-07 | protein jagunal |
| RS008630 | 3.0 | 4.6 | 1.1E-19 | 1.8 | 4.6 | 8.2E-08 | -1.1 | 4.6 | 9.2E-04 | |
| RS008741 | 1.6 | 13.0 | 8.0E-14 | -0.1 | 13.0 | 8.6E-01 | -1.7 | 13.0 | 3.4E-15 | titin |
| RS008743 | 1.1 | 10.3 | 9.1E-10 | -0.1 | 10.3 | 7.4E-01 | -1.2 | 10.3 | 5.4E-12 | titin |
| RS008795 | 1.1 | 5.8 | 2.4E-13 | -0.4 | 5.8 | 5.9E-02 | -1.5 | 5.8 | 4.4E-23 | cd151 antigen-like |
| RS008796 | 1.9 | 11.4 | 7.6E-16 | 0.2 | 11.4 | 6.7E-01 | -1.7 | 11.4 | 1.1E-12 | troponin i |
| RS008822 | 1.9 | 9.5 | 1.5E-43 | 0.1 | 9.5 | 7.8E-01 | -1.8 | 9.5 | 9.5E-40 | aspartyl asparaginyl beta-hydroxylase |
| RS008823 | 3.6 | 5.1 | 2.3E-76 | 0.1 | 5.1 | 8.6E-01 | -3.5 | 5.1 | 7.9E-72 | |

| | | | | | | | | | | |
|---|---|---|---|---|---|---|---|---|---|---|
| RS008824 | 4.8 | 1.0 | 1.8E-20 | 3.2 | 1.0 | 2.0E-06 | -1.7 | 1.0 | 3.3E-05 | |
| RS008874 | 1.5 | 13.4 | 1.5E-09 | -0.4 | 13.4 | 3.5E-01 | -1.8 | 13.4 | 2.8E-14 | myosin regulatory light chain 2 |
| RS009061 | 1.9 | 4.4 | 3.0E-13 | 0.5 | 4.4 | 2.0E-01 | -1.4 | 4.4 | 2.8E-07 | elongation of very long chain fatty acids protein aael008004 |
| RS009124 | 1.3 | 5.5 | 4.1E-07 | 0.3 | 5.5 | 5.9E-01 | -1.0 | 5.5 | 1.6E-04 | regucalcin-like |
| RS009206 | 1.5 | 5.9 | 1.1E-32 | -0.2 | 5.9 | 3.8E-01 | -1.7 | 5.9 | 2.2E-40 | calmodulin-lysine n-methyltransferase |
| RS009217 | 2.5 | 4.1 | 3.3E-18 | 0.8 | 4.1 | 2.7E-02 | -1.7 | 4.1 | 7.9E-09 | cytochrome p450 4c1 |
| RS009292 | 1.6 | 3.6 | 4.5E-07 | 0.0 | 3.6 | 9.6E-01 | -1.6 | 3.6 | 1.8E-06 | |
| RS009341 | 1.2 | 8.3 | 1.1E-13 | 0.0 | 8.3 | 9.6E-01 | -1.2 | 8.3 | 4.1E-13 | protein sda1 homolog |
| RS009529 | 2.8 | 1.2 | 9.4E-07 | 1.1 | 1.2 | 2.7E-01 | -1.8 | 1.2 | 2.3E-03 | transcription factor sum-1 |
| RS009599 | 1.1 | 9.5 | 4.6E-14 | 0.0 | 9.5 | 9.5E-01 | -1.0 | 9.5 | 2.2E-13 | fructose-bisphosphate aldolase |
| RS009650 | 1.0 | 4.3 | 6.1E-03 | -0.1 | 4.3 | 8.7E-01 | -1.2 | 4.3 | 2.5E-03 | f-box wd repeat-containing protein 9-like |
| RS009716 | 1.4 | 7.3 | 1.8E-22 | 0.0 | 7.3 | 9.6E-01 | -1.4 | 7.3 | 1.0E-21 | myocyte-specific enhancer factor 2 |
| RS009717 | 1.4 | 5.2 | 8.5E-25 | -0.1 | 5.2 | 8.2E-01 | -1.5 | 5.2 | 6.0E-27 | |
| RS009878 | 2.2 | 4.9 | 1.0E-19 | -0.5 | 4.9 | 2.0E-01 | -2.7 | 4.9 | 3.1E-27 | |
| RS009887 | 2.5 | 9.5 | 2.4E-19 | -0.1 | 9.5 | 8.4E-01 | -2.6 | 9.5 | 3.9E-21 | |
| RS009917 | 3.8 | 5.9 | 3.1E-37 | 2.0 | 5.9 | 3.5E-11 | -1.8 | 5.9 | 4.0E-10 | phosphoenolpyruvate carboxykinase |
| RS009918 | 4.2 | 3.0 | 3.8E-27 | 2.2 | 3.0 | 5.0E-06 | -1.9 | 3.0 | 4.1E-09 | phosphoenolpyruvate carboxykinase |
| RS009924 | 1.3 | 5.0 | 8.6E-22 | 0.0 | 5.0 | 9.4E-01 | -1.2 | 5.0 | 2.6E-20 | ankyrin repeat and ibr domain-containing protein 1-like |
| RS009933 | 1.8 | 9.6 | 6.9E-27 | 0.4 | 9.6 | 5.3E-02 | -1.4 | 9.6 | 6.0E-16 | |
| RS009934 | 2.5 | 7.6 | 2.8E-58 | 0.4 | 7.6 | 9.2E-02 | -2.1 | 7.6 | 1.7E-43 | |
| RS009935 | 2.6 | 8.4 | 8.3E-38 | 0.5 | 8.4 | 1.0E-01 | -2.1 | 8.4 | 1.3E-26 | pdz and lim domain protein 3 |
| RS009949 | 3.9 | 1.5 | 1.3E-11 | 0.9 | 1.5 | 5.6E-01 | -2.9 | 1.5 | 5.6E-08 | g-protein coupled receptor mth2 |
| RS010034 | 2.1 | 10.3 | 3.5E-18 | 0.1 | 10.3 | 8.3E-01 | -2.0 | 10.3 | 3.4E-16 | titin |
| RS010105 | 1.3 | 6.6 | 1.3E-04 | 0.1 | 6.6 | 9.4E-01 | -1.2 | 6.6 | 4.4E-04 | elongation of very long chain fatty acids protein 6 |
| RS010177 | 2.0 | 3.7 | 4.3E-15 | 0.1 | 3.7 | 9.1E-01 | -1.9 | 3.7 | 1.2E-13 | |
| RS010186 | 2.6 | 2.6 | 4.0E-16 | 0.2 | 2.6 | 8.5E-01 | -2.4 | 2.6 | 1.0E-13 | probable proline dehydrogenase 2 |
| RS010203 | 1.6 | 4.2 | 7.0E-12 | 0.6 | 4.2 | 6.4E-02 | -1.0 | 4.2 | 3.7E-05 | |
| RS010268 | 1.3 | 4.2 | 1.7E-11 | -0.2 | 4.2 | 5.5E-01 | -1.5 | 4.2 | 5.6E-15 | serine threonine-protein kinase sbk1 |
| RS010486 | 2.7 | 4.4 | 6.7E-22 | 1.5 | 4.4 | 3.9E-07 | -1.2 | 4.4 | 5.7E-05 | peritrophin-1-like |
| RS010508 | 1.3 | 7.0 | 2.3E-16 | -0.2 | 7.0 | 5.4E-01 | -1.5 | 7.0 | 7.6E-21 | protein tanc2 |
| RS010535 | 1.4 | 3.2 | 4.1E-12 | 0.2 | 3.2 | 7.1E-01 | -1.2 | 3.2 | 3.8E-09 | protein beta isoform |
| RS010561 | 2.1 | 10.8 | 3.2E-18 | 0.9 | 10.8 | 1.7E-03 | -1.2 | 10.8 | 1.1E-06 | tubulin alpha-1 chain |
| RS010570 | 2.3 | 9.4 | 3.0E-34 | 0.5 | 9.4 | 2.8E-02 | -1.8 | 9.4 | 3.4E-21 | sarcalumenin |
| RS010607 | 2.2 | 3.7 | 3.0E-19 | 0.8 | 3.7 | 1.1E-02 | -1.4 | 3.7 | 1.3E-08 | |
| RS010842 | 4.1 | 6.9 | 5.7E-16 | 0.1 | 6.9 | 9.5E-01 | -4.1 | 6.9 | 6.5E-15 | |
| RS010843 | 2.9 | 12.2 | 2.4E-22 | 0.6 | 12.2 | 1.9E-01 | -2.3 | 12.2 | 3.9E-15 | |
| RS010847 | 3.3 | 2.1 | 3.8E-22 | 1.0 | 2.1 | 5.1E-02 | -2.2 | 2.1 | 2.6E-12 | helix-loop-helix protein delilah |
| RS010848 | 1.5 | 3.3 | 1.3E-07 | 0.3 | 3.3 | 5.4E-01 | -1.2 | 3.3 | 8.7E-05 | helix-loop-helix protein delilah |
| RS010908 | 1.9 | 6.7 | 1.2E-61 | -0.4 | 6.7 | 2.2E-02 | -2.3 | 6.7 | 6.2E-83 | ras-related and estrogen-regulated growth inhibitor |
| RS010935 | 1.5 | 6.7 | 5.9E-23 | 0.5 | 6.7 | 1.1E-02 | -1.0 | 6.7 | 7.0E-11 | integrin-linked protein kinase |
| RS010967 | 2.2 | 4.2 | 2.5E-22 | -0.1 | 4.2 | 9.2E-01 | -2.2 | 4.2 | 5.0E-23 | protein fam135a |
| RS011024 | 2.1 | 0.4 | 3.0E-04 | -0.2 | 0.4 | 9.1E-01 | -2.2 | 0.4 | 2.6E-04 | heparan sulfate 2-o-sulfotransferase pipe |

| | | | | | | | | | | |
|---|---|---|---|---|---|---|---|---|---|---|
| RS011025 | 1.9 | 2.1 | 3.4E-08 | -0.1 | 2.1 | 9.2E-01 | -2.0 | 2.1 | 1.8E-08 | heparan sulfate 2-o-sulfotransferase pipe |
| RS011152 | 2.5 | 0.5 | 7.5E-08 | -0.5 | 0.5 | 7.2E-01 | -3.0 | 0.5 | 6.2E-09 | |
| RS011174 | 2.7 | 7.0 | 1.1E-71 | 0.3 | 7.0 | 1.4E-01 | -2.4 | 7.0 | 1.8E-56 | haloacid dehalogenase-like hydrolase domain-containing protein 2 |
| RS011193 | 1.6 | 5.9 | 4.9E-23 | -0.4 | 5.9 | 9.4E-02 | -2.0 | 5.9 | 4.7E-34 | organic cation transporter protein |
| RS011210 | 2.5 | 1.3 | 1.5E-12 | -0.2 | 1.3 | 8.9E-01 | -2.6 | 1.3 | 9.6E-13 | |
| RS011241 | 1.2 | 1.9 | 5.1E-03 | -0.1 | 1.9 | 8.9E-01 | -1.3 | 1.9 | 2.5E-03 | non-ltr retrotransposon cats |
| RS011282 | 2.3 | 0.1 | 2.3E-05 | 0.0 | 0.1 | 1.0E+00 | -2.3 | 0.1 | 8.0E-05 | |
| RS011283 | 1.8 | 9.6 | 6.6E-14 | 0.6 | 9.6 | 6.2E-02 | -1.2 | 9.6 | 1.2E-06 | |
| RS011284 | 1.9 | 9.5 | 1.9E-17 | 0.6 | 9.5 | 4.1E-02 | -1.3 | 9.5 | 9.6E-09 | pollen-specific leucine-rich repeat extensin-like protein 1 |
| RS011288 | 2.0 | 3.5 | 4.8E-13 | -0.1 | 3.5 | 8.4E-01 | -2.2 | 3.5 | 4.3E-14 | |
| RS011319 | 2.0 | 5.1 | 6.1E-08 | 0.4 | 5.1 | 6.5E-01 | -1.6 | 5.1 | 2.6E-05 | elongation of very long chain fatty acids protein aael008004 |
| RS011325 | 3.6 | 1.5 | 1.4E-11 | 0.8 | 1.5 | 5.4E-01 | -2.8 | 1.5 | 5.3E-08 | nose resistant to fluoxetine protein 6-like |
| RS011334 | 2.9 | 3.5 | 2.8E-15 | 1.0 | 3.5 | 3.9E-02 | -1.9 | 3.5 | 2.7E-07 | |
| RS011405 | 2.0 | 9.3 | 8.3E-09 | 0.9 | 9.3 | 3.5E-02 | -1.1 | 9.3 | 4.9E-03 | inositol-3-phosphate synthase 1-b |
| RS011410 | 2.8 | 6.9 | 2.9E-113 | 0.4 | 6.9 | 1.0E-02 | -2.4 | 6.9 | 6.4E-86 | protein |
| RS011411 | 4.0 | 6.3 | 5.5E-79 | 0.5 | 6.3 | 5.4E-02 | -3.4 | 6.3 | 4.7E-62 | protein isoform b-like |
| RS011412 | 3.9 | 5.0 | 7.2E-95 | 0.3 | 5.0 | 4.1E-01 | -3.6 | 5.0 | 8.8E-83 | endonuclease-reverse transcriptase |
| RS011454 | 3.7 | 0.3 | 4.5E-09 | -0.2 | 0.3 | 9.3E-01 | -3.9 | 0.3 | 4.8E-09 | |
| RS011494 | 1.6 | 7.5 | 8.1E-23 | 0.5 | 7.5 | 5.9E-03 | -1.1 | 7.5 | 2.4E-10 | laminin subunit beta-1 |
| RS011508 | 1.3 | 14.5 | 2.8E-09 | 0.0 | 14.5 | 9.6E-01 | -1.3 | 14.5 | 1.3E-09 | myosin heavy muscle |
| RS011519 | 2.7 | 5.9 | 6.3E-61 | 0.5 | 5.9 | 1.7E-02 | -2.2 | 5.9 | 5.1E-42 | cytosolic carboxypeptidase 1-like |
| RS011560 | 2.9 | 3.1 | 6.1E-16 | 1.7 | 3.1 | 1.2E-05 | -1.2 | 3.1 | 1.6E-03 | dehydrogenase reductase sdr family member 11-like |
| RS011568 | 1.4 | 2.2 | 2.2E-03 | -0.2 | 2.2 | 8.2E-01 | -1.6 | 2.2 | 5.0E-04 | protein vestigial |
| RS011646 | 3.0 | 4.7 | 2.7E-79 | -0.1 | 4.7 | 8.2E-01 | -3.1 | 4.7 | 7.8E-81 | serine threonine-protein phosphatase pp1-gamma catalytic subunit |
| RS011689 | 1.5 | 7.6 | 5.1E-05 | -3.6 | 7.6 | 2.3E-21 | -5.1 | 7.6 | 2.4E-38 | alpha-tocopherol transfer |
| RS011693 | 3.4 | 6.4 | 8.6E-25 | -3.5 | 6.4 | 4.7E-20 | -7.0 | 6.4 | 2.1E-66 | alpha-tocopherol transfer |
| RS011712 | 4.3 | 0.4 | 1.8E-09 | 2.5 | 0.4 | 1.2E-02 | -1.8 | 0.4 | 4.7E-03 | glucose dehydrogenase |
| RS011713 | 6.7 | 3.6 | 2.1E-29 | 5.3 | 3.6 | 3.7E-19 | -1.3 | 3.6 | 8.4E-03 | glucose dehydrogenase |
| RS011763 | 1.2 | 4.6 | 4.1E-07 | -0.4 | 4.6 | 3.5E-01 | -1.5 | 4.6 | 5.0E-11 | proton-coupled amino acid transporter 4 |
| RS011839 | 2.2 | 3.1 | 1.4E-14 | 1.0 | 3.1 | 3.9E-03 | -1.2 | 3.1 | 5.4E-05 | |
| RS011850 | 1.8 | 8.2 | 8.3E-40 | -0.1 | 8.2 | 8.1E-01 | -1.9 | 8.2 | 4.6E-43 | trimeric intracellular cation channel type b |
| RS011875 | 1.5 | 7.1 | 6.9E-08 | 0.3 | 7.1 | 5.9E-01 | -1.2 | 7.1 | 3.3E-05 | cytochrome p450 |
| RS011966 | 1.3 | 6.0 | 6.7E-17 | 0.0 | 6.0 | 9.5E-01 | -1.3 | 6.0 | 4.8E-16 | set and mynd domain-containing protein 4-like |
| RS011988 | 2.4 | 7.0 | 3.1E-34 | 0.5 | 7.0 | 4.2E-02 | -1.9 | 7.0 | 8.0E-22 | aquaporin-11 |
| RS012030 | 2.0 | 1.3 | 1.1E-07 | 0.6 | 1.3 | 3.7E-01 | -1.4 | 1.3 | 4.1E-04 | |
| RS012090 | 1.4 | 2.0 | 1.5E-04 | 0.3 | 2.0 | 6.6E-01 | -1.1 | 2.0 | 6.8E-03 | |
| RS012213 | 2.0 | 1.8 | 2.7E-08 | 0.8 | 1.8 | 1.8E-01 | -1.3 | 1.8 | 9.4E-04 | isoform b |
| RS012478 | 8.4 | 2.3 | 4.7E-57 | 5.3 | 2.3 | 1.2E-14 | -3.1 | 2.3 | 3.4E-21 | cell wall protein dan4 |
| RS012481 | 3.7 | 5.2 | 1.0E-28 | 0.9 | 5.2 | 2.0E-02 | -2.7 | 5.2 | 2.4E-17 | protein giant-lens |

| | | | | | | | | | | |
|---|---|---|---|---|---|---|---|---|---|---|
| RS012561 | 1.3 | 5.0 | 2.5E-10 | 0.0 | 5.0 | 9.7E-01 | -1.3 | 5.0 | 1.0E-09 | polycomb complex protein bmi-1 |
| RS012575 | 1.3 | 4.9 | 5.8E-18 | -0.2 | 4.9 | 5.9E-01 | -1.5 | 4.9 | 7.0E-22 | phosphatidylinositol 5-phosphate 4-kinase type-2 alpha |
| RS012585 | 2.6 | 5.2 | 6.7E-54 | 0.3 | 5.2 | 2.5E-01 | -2.3 | 5.2 | 3.0E-42 | gtp-binding protein 2 |
| RS012636 | 1.6 | 5.6 | 4.6E-09 | 0.3 | 5.6 | 5.7E-01 | -1.3 | 5.6 | 4.1E-06 | dopamine n-acetyltransferase-like |
| RS012638 | 2.2 | 3.3 | 2.8E-10 | 1.0 | 3.3 | 2.5E-02 | -1.2 | 3.3 | 1.3E-03 | |
| RS012649 | 5.5 | 10.7 | 1.2E-74 | -1.7 | 10.7 | 3.6E-09 | -7.1 | 10.7 | 5.7E-108 | troponin |
| RS012674 | 2.3 | 4.0 | 2.3E-33 | 0.4 | 4.0 | 2.5E-01 | -1.9 | 4.0 | 6.1E-24 | |
| RS012675 | 2.2 | 6.5 | 3.8E-32 | 0.6 | 6.5 | 6.5E-03 | -1.6 | 6.5 | 2.4E-17 | |
| RS012689 | 1.1 | 5.0 | 2.4E-13 | -0.2 | 5.0 | 3.3E-01 | -1.3 | 5.0 | 7.6E-19 | solute carrier family 41 member 1-like |
| RS012736 | 1.3 | 4.4 | 6.2E-06 | 0.1 | 4.4 | 9.1E-01 | -1.2 | 4.4 | 3.8E-05 | sine oculis-binding protein homolog |
| RS012935 | 1.1 | 4.9 | 9.2E-06 | 0.1 | 4.9 | 9.0E-01 | -1.1 | 4.9 | 7.0E-05 | leucine-rich repeat and transmembrane domain-containing protein |
| RS012944 | 1.7 | 5.9 | 2.8E-23 | 0.3 | 5.9 | 1.8E-01 | -1.3 | 5.9 | 5.0E-15 | isopentenyl-diphosphate delta-isomerase 1 |
| RS012952 | 1.2 | 3.8 | 1.6E-07 | -0.1 | 3.8 | 8.4E-01 | -1.3 | 3.8 | 1.6E-08 | rna-binding protein 24-like |
| RS013013 | 2.2 | 4.1 | 1.7E-11 | 0.7 | 4.1 | 1.3E-01 | -1.5 | 4.1 | 6.8E-06 | peptidoglycan-recognition protein lb |
| RS013075 | 1.7 | 5.2 | 3.5E-12 | 0.1 | 5.2 | 9.2E-01 | -1.7 | 5.2 | 3.6E-11 | maltase a8 |
| RS013142 | 2.9 | 2.2 | 3.4E-23 | 0.4 | 2.2 | 6.2E-01 | -2.5 | 2.2 | 5.1E-18 | spermatogenesis-associated protein 18-like protein |
| RS013151 | 1.9 | 5.1 | 3.3E-22 | 0.1 | 5.1 | 8.6E-01 | -1.8 | 5.1 | 6.8E-20 | camp-specific 3 -cyclic phosphodiesterase 4a-like |
| RS013178 | 1.3 | 9.0 | 5.4E-27 | -0.2 | 9.0 | 4.7E-01 | -1.4 | 9.0 | 1.2E-33 | pericentriolar material 1 protein |
| RS013259 | 2.4 | 1.6 | 9.2E-19 | 0.6 | 1.6 | 1.9E-01 | -1.7 | 1.6 | 4.8E-11 | t-box transcription factor tbx20-like |
| RS013260 | 2.4 | 2.4 | 5.3E-25 | 0.5 | 2.4 | 2.0E-01 | -1.9 | 2.4 | 3.5E-16 | t-box transcription factor tbx20-like |
| RS013343 | 1.8 | 2.1 | 5.8E-10 | 0.0 | 2.1 | 9.7E-01 | -1.8 | 2.1 | 1.1E-09 | |
| RS013344 | 1.6 | 2.1 | 2.1E-06 | 0.0 | 2.1 | 9.7E-01 | -1.6 | 2.1 | 7.6E-06 | |
| RS013361 | 1.9 | 3.8 | 2.1E-14 | 0.3 | 3.8 | 5.8E-01 | -1.6 | 3.8 | 1.3E-10 | |
| RS013376 | 1.9 | 4.4 | 4.5E-18 | 0.9 | 4.4 | 7.2E-04 | -1.0 | 4.4 | 5.9E-06 | |
| RS013395 | 2.1 | 4.3 | 3.1E-23 | 0.0 | 4.3 | 9.8E-01 | -2.0 | 4.3 | 2.2E-22 | homeotic protein spalt-major-like |
| RS013396 | 1.5 | 1.6 | 7.7E-05 | -0.3 | 1.6 | 6.9E-01 | -1.8 | 1.6 | 2.9E-06 | |
| RS013430 | 2.6 | 7.6 | 5.3E-20 | 0.5 | 7.6 | 2.7E-01 | -2.1 | 7.6 | 1.0E-13 | collagen alpha-1 chain |
| RS013524 | 1.5 | 7.1 | 2.1E-37 | 0.1 | 7.1 | 7.8E-01 | -1.4 | 7.1 | 1.3E-33 | uncharacterized protein |
| RS013540 | 3.2 | 2.1 | 5.1E-15 | 1.2 | 2.1 | 3.3E-02 | -2.0 | 2.1 | 3.9E-07 | probable serine threonine-protein kinase kinx |
| RS013541 | 2.0 | 5.1 | 6.9E-27 | 0.6 | 5.1 | 1.3E-02 | -1.4 | 5.1 | 5.0E-14 | hypothetical protein |
| RS013543 | 1.7 | 1.9 | 7.2E-07 | 0.5 | 1.9 | 3.9E-01 | -1.2 | 1.9 | 1.1E-03 | |
| RS013544 | 1.4 | 6.8 | 1.8E-29 | 0.2 | 6.8 | 4.3E-01 | -1.2 | 6.8 | 1.2E-22 | hemicentin-1 |
| RS013560 | 1.4 | 9.0 | 4.6E-24 | -0.1 | 9.0 | 5.9E-01 | -1.5 | 9.0 | 4.7E-29 | ring finger protein 31 |
| RS013610 | 1.2 | 6.3 | 6.5E-06 | 0.2 | 6.3 | 7.5E-01 | -1.0 | 6.3 | 2.8E-04 | glucose dehydrogenase |
| RS013626 | 2.8 | 3.9 | 5.5E-15 | 0.9 | 3.9 | 5.4E-02 | -1.9 | 3.9 | 1.7E-07 | |
| RS013635 | 2.2 | 4.9 | 7.4E-10 | -2.1 | 4.9 | 3.9E-08 | -4.3 | 4.9 | 1.7E-29 | cytochrome p450 4c1 |
| RS013636 | 1.2 | 3.0 | 4.4E-03 | -0.2 | 3.0 | 7.8E-01 | -1.4 | 3.0 | 6.8E-04 | |
| RS013637 | 1.5 | 5.4 | 7.0E-10 | -0.4 | 5.4 | 3.0E-01 | -1.9 | 5.4 | 5.6E-15 | hypothetical protein |
| RS013687 | 1.3 | 3.9 | 5.1E-05 | -0.2 | 3.9 | 7.5E-01 | -1.5 | 3.9 | 3.1E-06 | acyl- delta desaturase |

| | | | | | | | | | | |
|---|---|---|---|---|---|---|---|---|---|---|
| RS013740 | 3.2 | 1.8 | 2.1E-17 | -0.7 | 1.8 | 4.8E-01 | -3.8 | 1.8 | 7.7E-21 | pdz and lim domain protein 3 |
| RS013858 | 1.7 | 11.8 | 1.8E-18 | -0.2 | 11.8 | 7.3E-01 | -1.9 | 11.8 | 1.7E-21 | myosin light chain alkali |
| RS013865 | 1.2 | 6.3 | 1.6E-14 | -0.2 | 6.3 | 4.0E-01 | -1.5 | 6.3 | 9.4E-20 | |
| RS013915 | 3.3 | 8.1 | 5.8E-26 | 0.7 | 8.1 | 1.0E-01 | -2.6 | 8.1 | 4.4E-17 | collagen alpha-1 chain |
| RS013964 | 1.3 | 4.3 | 8.4E-13 | -0.4 | 4.3 | 2.0E-01 | -1.7 | 4.3 | 1.7E-19 | |
| RS013965 | 1.1 | 6.3 | 1.5E-08 | -0.5 | 6.3 | 3.3E-02 | -1.7 | 6.3 | 3.0E-17 | scavenger receptor class b member 1 |
| RS013966 | 4.5 | 1.7 | 4.4E-23 | 0.5 | 1.7 | 7.0E-01 | -4.0 | 1.7 | 4.0E-19 | larval cuticle protein a2b-like |
| RS013978 | 2.6 | 0.7 | 1.0E-07 | -0.6 | 0.7 | 6.5E-01 | -3.2 | 0.7 | 2.1E-09 | cytosolic carboxypeptidase 1-like |
| RS013979 | 2.6 | 2.5 | 1.4E-08 | 0.4 | 2.5 | 6.9E-01 | -2.3 | 2.5 | 1.7E-06 | |
| RS014237 | 1.9 | 6.7 | 4.8E-31 | 0.8 | 6.7 | 5.9E-06 | -1.1 | 6.7 | 8.5E-11 | calcium-binding mitochondrial carrier protein aralar1 |
| RS014264 | 1.0 | 6.6 | 5.9E-05 | -0.1 | 6.6 | 8.9E-01 | -1.1 | 6.6 | 1.6E-05 | |
| RS014332 | 1.1 | 7.0 | 8.9E-27 | -0.4 | 7.0 | 5.9E-04 | -1.5 | 7.0 | 9.4E-49 | kh domain-containing protein |
| RS014339 | 2.0 | 5.7 | 9.4E-25 | -0.9 | 5.7 | 8.0E-05 | -2.8 | 5.7 | 6.7E-47 | hypothetical protein |
| RS014350 | 2.3 | 2.0 | 3.7E-12 | -0.6 | 2.0 | 3.6E-01 | -2.9 | 2.0 | 1.8E-16 | matrix metalloproteinase-16 |
| RS014367 | 1.3 | 7.3 | 6.5E-10 | 0.0 | 7.3 | 9.7E-01 | -1.3 | 7.3 | 4.4E-10 | sec14-like protein 2 |
| RS014469 | 3.3 | 0.2 | 2.1E-06 | 0.4 | 0.2 | 8.4E-01 | -2.9 | 0.2 | 4.9E-05 | |
| RS014471 | 1.4 | 2.7 | 4.3E-09 | 0.2 | 2.7 | 7.2E-01 | -1.2 | 2.7 | 9.6E-07 | rna-binding protein 24-like |
| RS014587 | 1.8 | 4.4 | 5.7E-11 | 0.5 | 4.4 | 2.7E-01 | -1.3 | 4.4 | 4.8E-06 | lysyl oxidase homolog 3 |
| RS014615 | 1.6 | 2.1 | 3.7E-09 | 0.6 | 2.1 | 1.9E-01 | -1.0 | 2.1 | 2.6E-04 | protein rhomboid |
| RS014699 | 1.2 | 5.0 | 3.2E-16 | 0.0 | 5.0 | 9.4E-01 | -1.2 | 5.0 | 5.3E-15 | inositol-pentakisphosphate 2-kinase |
| RS014713 | 1.2 | 5.0 | 1.8E-13 | -0.2 | 5.0 | 4.2E-01 | -1.5 | 5.0 | 3.4E-18 | secreted frizzled-related protein |
| RS014715 | 1.7 | 6.9 | 1.5E-29 | 0.5 | 6.9 | 6.5E-03 | -1.1 | 6.9 | 1.0E-14 | |
| RS014818 | 2.6 | 3.9 | 3.8E-06 | 0.6 | 3.9 | 6.5E-01 | -2.0 | 3.9 | 4.5E-04 | |
| RS014819 | 1.9 | 3.9 | 6.0E-05 | 0.1 | 3.9 | 9.7E-01 | -1.9 | 3.9 | 1.8E-04 | |
| RS014858 | 2.5 | 3.6 | 8.4E-09 | 0.6 | 3.6 | 4.0E-01 | -1.9 | 3.6 | 2.4E-05 | cysteine-rich secretory protein 1-like |
| RS014862 | 2.3 | 12.2 | 3.6E-36 | 0.8 | 12.2 | 5.3E-05 | -1.5 | 12.2 | 5.4E-16 | alpha- sarcomeric |
| RS014873 | 2.0 | 3.7 | 3.5E-03 | -0.3 | 3.7 | 8.7E-01 | -2.3 | 3.7 | 2.4E-03 | serine protease |
| RS015064 | 1.6 | 7.7 | 2.9E-18 | 0.6 | 7.7 | 1.2E-02 | -1.0 | 7.7 | 5.2E-08 | beta-hexosaminidase subunit beta-like |
| RS015184 | 1.9 | 3.6 | 1.9E-10 | 0.2 | 3.6 | 8.1E-01 | -1.7 | 3.6 | 1.4E-08 | ras-related and estrogen-regulated growth inhibitor-like |
| RS015297 | 1.2 | 2.4 | 1.6E-04 | 0.2 | 2.4 | 8.2E-01 | -1.0 | 2.4 | 2.4E-03 | frizzled-2-like |
| RS015356 | 2.0 | 3.3 | 1.5E-15 | 0.4 | 3.3 | 3.4E-01 | -1.5 | 3.3 | 4.8E-10 | |
| RS015395 | 2.0 | 6.9 | 3.2E-21 | -0.2 | 6.9 | 6.1E-01 | -2.2 | 6.9 | 2.4E-25 | ryanodine receptor 44f |
| RS015396 | 1.9 | 7.2 | 9.4E-12 | -0.2 | 7.2 | 7.8E-01 | -2.1 | 7.2 | 1.1E-13 | ryanodine receptor |
| RS015405 | 1.6 | 4.8 | 1.2E-20 | 0.0 | 4.8 | 9.2E-01 | -1.5 | 4.8 | 3.4E-19 | |
| RS015406 | 1.6 | 5.5 | 3.3E-37 | 0.1 | 5.5 | 8.0E-01 | -1.6 | 5.5 | 2.0E-33 | sodium potassium-transporting atpase subunit beta-2 |
| RS015410 | 1.3 | 6.9 | 7.7E-23 | 0.2 | 6.9 | 3.8E-01 | -1.1 | 6.9 | 2.2E-16 | protein slowmo |
| RS015450 | 4.2 | 3.9 | 7.7E-54 | 1.2 | 3.9 | 4.5E-04 | -3.0 | 3.9 | 1.2E-32 | growth differentiation factor 11 |
| RS015498 | 1.5 | 3.0 | 4.1E-10 | 0.0 | 3.0 | 1.0E+00 | -1.5 | 3.0 | 1.4E-09 | calcium uniporter mitochondrial |
| RS015510 | 1.7 | 3.7 | 2.5E-07 | 0.6 | 3.7 | 2.4E-01 | -1.1 | 3.7 | 1.8E-03 | protein apcdd1 |
| RS015520 | 2.2 | 6.9 | 4.0E-83 | 0.2 | 6.9 | 2.7E-01 | -2.0 | 6.9 | 8.6E-69 | dcn1-like protein 5 |
| RS100004 | 4.8 | 4.5 | 5.2E-09 | 1.8 | 4.5 | 2.9E-01 | -3.0 | 4.5 | 3.7E-05 | lysozyme-like |
| RS100012 | 4.7 | 5.2 | 1.2E-08 | -0.4 | 5.2 | 8.0E-01 | -5.1 | 5.2 | 1.3E-09 | geranylgeranyl pyrophosphate synthase |
| RS100014 | 2.3 | 0.4 | 1.9E-05 | -0.4 | 0.4 | 8.4E-01 | -2.6 | 0.4 | 7.5E-06 | geranylgeranyl pyrophosphate synthase |
| RS100015 | 5.8 | 7.4 | 4.9E-35 | 0.8 | 7.4 | 2.4E-01 | -5.1 | 7.4 | 1.1E-28 | geranylgeranyl pyrophosphate synthase |

| | | | | | | | | | | |
|---|---|---|---|---|---|---|---|---|---|---|
| RS100016 | 2.9 | 8.9 | 3.6E-14 | 0.7 | 8.9 | 1.8E-01 | -2.2 | 8.9 | 1.8E-08 | geranylgeranyl pyrophosphate synthase |

SI_Dataset_1
Caste-biased genes: Soldier (thorax + abdomen)

| Gene ID | Thorax + abdomen Reproductive v.s. Soldier | | | Thorax + abdomen Reproductive v.s. Worker | | | Thorax + abdomen Soldier v.s. Worker | | | Gene annotation |
|---|---|---|---|---|---|---|---|---|---|---|
| | logFC | logCPM | FDR | logFC | logCPM | FDR | logFC | logCPM | FDR | |
| RS000043 | 3.3 | 3.2 | 7.1E-20 | 2.2 | 3.2 | 8.1E-09 | -1.1 | 3.2 | 2.6E-03 | |
| RS000044 | 2.8 | 2.0 | 1.6E-12 | 1.7 | 2.0 | 9.7E-05 | -1.1 | 2.0 | 7.7E-03 | enzymatic polyprotein endonuclease reverse |
| RS000208 | 2.2 | 4.6 | 1.2E-08 | -0.5 | 4.6 | 3.1E-01 | -2.7 | 4.6 | 1.4E-11 | |
| RS000213 | 1.5 | 3.2 | 9.7E-07 | 0.0 | 3.2 | 9.5E-01 | -1.6 | 3.2 | 2.4E-06 | innexin shaking-b |
| RS000247 | 1.9 | 5.6 | 1.8E-14 | 0.3 | 5.6 | 3.8E-01 | -1.6 | 5.6 | 1.6E-10 | exonuclease gor |
| RS000274 | 1.1 | 5.2 | 3.9E-07 | 0.1 | 5.2 | 7.6E-01 | -1.0 | 5.2 | 1.5E-05 | |
| RS000365 | 1.2 | 12.6 | 1.4E-06 | -0.5 | 12.6 | 8.1E-02 | -1.7 | 12.6 | 1.2E-11 | calcium-transporting atpase sarcoplasmic endoplasmic reticulum type |
| RS000368 | 1.2 | 5.7 | 2.6E-10 | -0.2 | 5.7 | 5.5E-01 | -1.4 | 5.7 | 5.3E-12 | carboxypeptidase m |
| RS000842 | 2.6 | 2.9 | 1.8E-08 | -0.5 | 2.9 | 5.6E-01 | -3.1 | 2.9 | 2.6E-09 | cytochrome p450 4c1-like |
| RS000843 | 3.7 | 3.6 | 3.5E-11 | 0.7 | 3.6 | 4.0E-01 | -3.0 | 3.6 | 9.1E-08 | cytochrome p450-like protein |
| RS000849 | 2.1 | 3.9 | 1.0E-12 | 0.9 | 3.9 | 6.6E-03 | -1.2 | 3.9 | 1.0E-04 | monocarboxylate transporter 12 |
| RS000906 | 1.9 | 5.2 | 3.2E-08 | 0.7 | 5.2 | 9.9E-02 | -1.3 | 5.2 | 7.5E-04 | |
| RS000908 | 1.7 | 3.2 | 1.4E-04 | 0.4 | 3.2 | 5.2E-01 | -1.3 | 3.2 | 6.3E-03 | |
| RS000936 | 5.2 | 9.1 | 1.4E-110 | 1.6 | 9.1 | 4.4E-13 | -3.6 | 9.1 | 3.0E-63 | |
| RS000947 | 1.4 | 9.3 | 4.8E-11 | -0.1 | 9.3 | 7.4E-01 | -1.5 | 9.3 | 4.1E-12 | protein disulfide-isomerase |
| RS000948 | 1.2 | 7.1 | 5.7E-08 | 0.0 | 7.1 | 9.8E-01 | -1.2 | 7.1 | 2.0E-07 | protein disulfide-isomerase a5 |
| RS000962 | 4.4 | 5.6 | 1.9E-08 | 1.7 | 5.6 | 2.5E-02 | -2.6 | 5.6 | 7.0E-04 | lipase 3-like |
| RS001146 | 1.7 | 3.2 | 6.9E-05 | -0.9 | 3.2 | 5.3E-02 | -2.6 | 3.2 | 2.7E-09 | fatty acyl- reductase cg5065 |
| RS001263 | 1.9 | 6.1 | 4.8E-10 | 0.0 | 6.1 | 9.9E-01 | -1.9 | 6.1 | 9.3E-10 | elongation of very long chain fatty acids protein 7-like |
| RS001264 | 3.0 | 1.1 | 3.7E-11 | 0.5 | 1.1 | 5.0E-01 | -2.5 | 1.1 | 7.9E-08 | elongation of very long chain fatty acids protein 7-like |
| RS001373 | 1.6 | 1.8 | 3.3E-04 | -0.2 | 1.8 | 7.7E-01 | -1.8 | 1.8 | 1.5E-04 | facilitated trehalose transporter tret1-2 homolog |
| RS001531 | 2.6 | 2.7 | 1.5E-09 | -0.8 | 2.7 | 2.3E-01 | -3.4 | 2.7 | 1.4E-12 | clavesin-2 |
| RS001585 | 2.5 | 0.5 | 2.7E-09 | 0.3 | 0.5 | 7.4E-01 | -2.2 | 0.5 | 5.1E-07 | fatty acyl- reductase cg5065 |
| RS001586 | 2.1 | 5.7 | 1.2E-14 | 0.0 | 5.7 | 9.8E-01 | -2.1 | 5.7 | 5.5E-14 | fatty acyl- reductase cg5065 |
| RS001595 | 1.2 | 7.0 | 5.2E-11 | -0.1 | 7.0 | 8.3E-01 | -1.3 | 7.0 | 1.1E-11 | acyl- delta desaturase |
| RS001613 | 1.1 | 5.5 | 3.6E-08 | 0.0 | 5.5 | 8.9E-01 | -1.2 | 5.5 | 4.7E-08 | |
| RS001705 | 1.9 | 5.8 | 1.3E-15 | 0.1 | 5.8 | 8.2E-01 | -1.8 | 5.8 | 8.3E-14 | tubulointerstitial nephritis antigen-like |
| RS001887 | 3.6 | 4.9 | 6.9E-29 | 0.7 | 4.9 | 8.9E-02 | -2.9 | 4.9 | 8.8E-20 | protein croquemort-like |
| RS002051 | 2.1 | 10.3 | 1.6E-22 | 0.1 | 10.3 | 8.5E-01 | -2.1 | 10.3 | 7.6E-21 | troponin |
| RS002056 | 1.0 | 2.4 | 1.6E-03 | 0.0 | 2.4 | 9.8E-01 | -1.0 | 2.4 | 3.8E-03 | |
| RS002180 | 1.3 | 3.9 | 6.2E-07 | -0.3 | 3.9 | 4.8E-01 | -1.6 | 3.9 | 2.3E-08 | serine threonine-protein phosphatase rdgc |
| RS002238 | 1.1 | 4.1 | 1.1E-05 | -0.3 | 4.1 | 3.3E-01 | -1.4 | 4.1 | 4.2E-08 | alpha-tocopherol transfer |
| RS002240 | 2.0 | 6.3 | 4.8E-23 | -0.2 | 6.3 | 4.0E-01 | -2.2 | 6.3 | 8.1E-27 | probable pyruvate dehydrogenase e1 component subunit mitochondrial |
| RS002356 | 1.9 | 4.8 | 3.8E-14 | -0.1 | 4.8 | 7.0E-01 | -2.0 | 4.8 | 2.2E-15 | fatty acyl- reductase cg5065 |
| RS002357 | 1.9 | 2.8 | 2.8E-13 | -0.1 | 2.8 | 7.6E-01 | -2.1 | 2.8 | 8.8E-14 | |
| RS002448 | 1.8 | 3.4 | 6.1E-04 | 0.0 | 3.4 | 9.9E-01 | -1.8 | 3.4 | 7.0E-04 | fatty acyl- reductase cg5065 |
| RS002487 | 2.6 | 8.5 | 5.8E-09 | 0.9 | 8.5 | 7.8E-02 | -1.7 | 8.5 | 1.8E-04 | defensin |
| RS002546 | 1.7 | 6.7 | 1.9E-10 | 0.4 | 6.7 | 1.9E-01 | -1.3 | 6.7 | 4.7E-06 | scavenger receptor class b member 1 |

| | | | | | | | | | | |
|---|---|---|---|---|---|---|---|---|---|---|
| RS002695 | 2.5 | 6.6 | 1.2E-35 | 0.0 | 6.6 | 9.1E-01 | -2.6 | 6.6 | 8.2E-36 | |
| RS002790 | 3.2 | 1.3 | 2.9E-10 | -0.9 | 1.3 | 3.8E-01 | -4.1 | 1.3 | 1.1E-11 | serine protease 33-like |
| RS002791 | 1.7 | 6.1 | 9.7E-14 | 0.6 | 6.1 | 3.8E-02 | -1.2 | 6.1 | 1.4E-06 | |
| RS002800 | 4.6 | 1.4 | 1.1E-09 | 1.3 | 1.4 | 1.7E-01 | -3.3 | 1.4 | 8.2E-06 | hypothetical protein |
| RS002801 | 4.4 | 4.2 | 2.2E-17 | 1.5 | 4.2 | 5.0E-03 | -2.9 | 4.2 | 5.8E-09 | enzymatic polyprotein endonuclease reverse |
| RS002816 | 1.6 | 6.6 | 2.0E-19 | 0.3 | 6.6 | 1.1E-01 | -1.3 | 6.6 | 3.7E-12 | multidrug resistance protein 1 |
| RS002930 | 1.4 | 3.3 | 3.0E-05 | 0.3 | 3.3 | 6.2E-01 | -1.1 | 3.3 | 1.8E-03 | acetylcholine receptor subunit alpha-like 2 |
| RS003129 | 2.3 | 2.9 | 1.5E-10 | -0.6 | 2.9 | 2.7E-01 | -2.8 | 2.9 | 1.2E-13 | organic cation transporter protein |
| RS003189 | 2.1 | 0.5 | 1.2E-03 | -0.1 | 0.5 | 9.3E-01 | -2.2 | 0.5 | 2.7E-03 | zinc metalloproteinase nas-15 |
| RS003197 | 1.1 | 6.9 | 1.5E-06 | -0.4 | 6.9 | 9.9E-02 | -1.5 | 6.9 | 2.5E-11 | 17-beta-hydroxysteroid dehydrogenase type 6 |
| RS003292 | 3.5 | 12.6 | 3.7E-14 | 2.0 | 12.6 | 1.7E-05 | -1.6 | 12.6 | 1.5E-03 | ejaculatory bulb-specific protein 3 |
| RS003343 | 1.0 | 3.0 | 5.7E-05 | -0.3 | 3.0 | 4.7E-01 | -1.3 | 3.0 | 2.2E-06 | tyrosine-protein kinase receptor torso |
| RS003367 | 1.2 | 3.2 | 1.3E-04 | 0.1 | 3.2 | 7.3E-01 | -1.0 | 3.2 | 1.9E-03 | |
| RS003372 | 1.7 | 2.5 | 2.2E-06 | 0.2 | 2.5 | 7.6E-01 | -1.5 | 2.5 | 6.7E-05 | udp-glucuronosyltransferase 2c1-like |
| RS003425 | 3.1 | 5.7 | 4.1E-13 | 1.7 | 5.7 | 6.4E-05 | -1.4 | 5.7 | 3.1E-03 | transposon ty3-g gag-pol polyprotein |
| RS003523 | 1.3 | 3.3 | 2.8E-04 | 0.1 | 3.3 | 8.4E-01 | -1.2 | 3.3 | 1.6E-03 | proton-associated sugar transporter a-like |
| RS003552 | 1.9 | 5.2 | 4.7E-15 | 0.7 | 5.2 | 1.0E-02 | -1.2 | 5.2 | 2.2E-06 | xanthine dehydrogenase oxidase-like |
| RS003709 | 6.9 | 5.9 | 1.0E-60 | -0.9 | 5.9 | 1.9E-01 | -7.8 | 5.9 | 7.6E-65 | cytochrome p450 4v2 |
| RS003997 | 1.9 | 7.2 | 1.1E-21 | 0.6 | 7.2 | 9.9E-03 | -1.3 | 7.2 | 6.7E-11 | abc transporter g family member 23 |
| RS004058 | 1.2 | 5.2 | 9.5E-10 | -0.4 | 5.2 | 1.7E-01 | -1.6 | 5.2 | 5.7E-14 | sh3 and cysteine-rich domain-containing protein 2 |
| RS004116 | 1.3 | 8.5 | 1.9E-18 | -0.2 | 8.5 | 4.1E-01 | -1.5 | 8.5 | 2.2E-22 | #NAME? |
| RS004171 | 1.5 | 4.3 | 2.9E-06 | 0.4 | 4.3 | 2.7E-01 | -1.1 | 4.3 | 2.0E-03 | cuticular protein analogous to peritrophins 1-b |
| RS004288 | 1.1 | 12.4 | 1.4E-06 | -1.5 | 12.4 | 6.7E-12 | -2.5 | 12.4 | 1.2E-31 | |
| RS004330 | 1.3 | 7.6 | 1.1E-10 | 0.0 | 7.6 | 9.5E-01 | -1.3 | 7.6 | 6.3E-10 | zinc finger ccch domain-containing protein 13 |
| RS004386 | 3.4 | 8.1 | 4.7E-21 | 2.3 | 8.1 | 2.0E-10 | -1.1 | 8.1 | 3.8E-03 | |
| RS004547 | 7.5 | 3.9 | 9.6E-111 | 0.6 | 3.9 | 6.3E-01 | -6.9 | 3.9 | 2.0E-95 | structural constituent of |
| RS004548 | 1.3 | 3.3 | 1.6E-03 | -0.3 | 3.3 | 5.8E-01 | -1.6 | 3.3 | 2.3E-04 | cuticle protein 8 |
| RS004558 | 5.5 | 9.2 | 4.9E-08 | 2.7 | 9.2 | 5.4E-03 | -2.8 | 9.2 | 4.1E-03 | cuticle protein 19 |
| RS004890 | 1.5 | 5.6 | 7.2E-11 | -1.9 | 5.6 | 9.2E-13 | -3.4 | 5.6 | 2.7E-40 | |
| RS004891 | 1.5 | 3.3 | 2.5E-06 | -2.5 | 3.3 | 1.8E-08 | -4.0 | 3.3 | 1.5E-23 | |
| RS005001 | 1.7 | 6.3 | 1.6E-08 | 0.7 | 6.3 | 5.1E-02 | -1.1 | 6.3 | 1.2E-03 | probable atp-dependent rna helicase ddx60 |
| RS005033 | 4.3 | 6.2 | 1.9E-45 | 0.8 | 6.2 | 1.6E-02 | -3.5 | 6.2 | 2.3E-32 | hydroxymethylglutaryl- synthase 1 |
| RS005034 | 4.5 | 6.1 | 3.3E-62 | 0.7 | 6.1 | 4.6E-02 | -3.8 | 6.1 | 2.9E-48 | |
| RS005088 | 3.0 | 2.0 | 1.6E-08 | 0.7 | 2.0 | 4.2E-01 | -2.3 | 2.0 | 2.5E-05 | |
| RS005096 | 3.8 | 4.4 | 1.3E-36 | 0.8 | 4.4 | 5.2E-02 | -3.0 | 4.4 | 9.6E-25 | |
| RS005507 | 4.9 | 3.6 | 2.1E-18 | 3.2 | 3.6 | 1.3E-08 | -1.8 | 3.6 | 1.1E-03 | aael014316 |
| RS005522 | 1.0 | 7.6 | 6.6E-08 | -0.1 | 7.6 | 6.9E-01 | -1.1 | 7.6 | 6.5E-09 | cartilage oligomeric matrix protein |
| RS005715 | 1.1 | 3.1 | 1.8E-04 | 0.1 | 3.1 | 8.9E-01 | -1.1 | 3.1 | 9.7E-04 | |

| | | | | | | | | | | |
|---|---|---|---|---|---|---|---|---|---|---|
| RS006262 | 1.5 | 4.9 | 3.0E-08 | -0.3 | 4.9 | 3.0E-01 | -1.9 | 4.9 | 1.8E-11 | fatty acyl- reductase cg5065 |
| RS006263 | 1.2 | 3.1 | 3.1E-03 | 0.0 | 3.1 | 9.7E-01 | -1.2 | 3.1 | 5.7E-03 | fatty acyl- reductase cg5065 |
| RS006270 | 3.2 | 3.2 | 1.7E-14 | 1.3 | 3.2 | 1.6E-02 | -1.9 | 3.2 | 3.8E-06 | organic cation transporter 1 |
| RS006393 | 2.0 | 6.3 | 2.4E-34 | -1.1 | 6.3 | 1.5E-07 | -3.1 | 6.3 | 1.3E-64 | alpha-tocopherol transfer |
| RS006415 | 2.8 | 5.4 | 1.7E-20 | 1.8 | 5.4 | 4.9E-08 | -1.1 | 5.4 | 6.4E-04 | glucose dehydrogenase |
| RS006595 | 1.3 | 2.2 | 8.9E-04 | 0.2 | 2.2 | 7.8E-01 | -1.1 | 2.2 | 8.1E-03 | non-ltr retrotransposon cats |
| RS006676 | 1.9 | 5.7 | 3.6E-23 | 0.1 | 5.7 | 7.1E-01 | -1.7 | 5.7 | 1.5E-19 | adenylate kinase isoenzyme 1 |
| RS006819 | 1.2 | 7.0 | 3.9E-07 | -0.1 | 7.0 | 6.8E-01 | -1.3 | 7.0 | 3.3E-08 | hemocyte protein-glutamine gamma-glutamyltransferase-like |
| RS006905 | 2.0 | 4.6 | 1.6E-05 | -0.5 | 4.6 | 4.3E-01 | -2.5 | 4.6 | 2.6E-07 | pancreatic lipase-related protein 2-like |
| RS006967 | 1.4 | 7.2 | 1.9E-05 | 0.2 | 7.2 | 6.5E-01 | -1.2 | 7.2 | 4.8E-04 | |
| RS007350 | 1.1 | 4.5 | 6.1E-04 | -0.5 | 4.5 | 1.8E-01 | -1.6 | 4.5 | 9.2E-07 | ankyrin repeat protein |
| RS007457 | 1.0 | 10.6 | 5.8E-09 | -0.2 | 10.6 | 3.2E-01 | -1.3 | 10.6 | 2.3E-12 | hypothetical protein |
| RS007480 | 4.8 | 4.7 | 3.0E-18 | -2.3 | 4.7 | 2.8E-03 | -7.2 | 4.7 | 2.9E-27 | geranylgeranyl pyrophosphate synthase |
| RS007482 | 2.5 | 5.5 | 5.3E-07 | -1.8 | 5.5 | 2.5E-03 | -4.3 | 5.5 | 9.0E-15 | geranylgeranyl pyrophosphate synthase |
| RS007483 | 4.9 | 4.1 | 9.2E-27 | 1.3 | 4.1 | 3.1E-02 | -3.6 | 4.1 | 3.9E-17 | geranylgeranyl pyrophosphate synthase |
| RS007516 | 1.4 | 15.4 | 8.9E-07 | 0.1 | 15.4 | 7.7E-01 | -1.3 | 15.4 | 1.4E-05 | muscle |
| RS007592 | 3.1 | 3.5 | 3.6E-13 | 0.8 | 3.5 | 8.6E-02 | -2.2 | 3.5 | 2.0E-07 | |
| RS007594 | 2.1 | 8.5 | 4.5E-20 | 0.1 | 8.5 | 7.9E-01 | -2.0 | 8.5 | 3.9E-18 | fatty acid synthase |
| RS007595 | 2.3 | 5.8 | 2.4E-16 | 0.2 | 5.8 | 6.4E-01 | -2.1 | 5.8 | 1.2E-13 | fatty acid synthase-like |
| RS007596 | 2.7 | 4.9 | 1.5E-13 | 0.7 | 4.9 | 1.1E-01 | -2.0 | 4.9 | 5.5E-08 | fatty acid synthase |
| RS007880 | 1.3 | 6.5 | 3.1E-19 | 0.1 | 6.5 | 7.9E-01 | -1.2 | 6.5 | 2.2E-16 | |
| RS007945 | 4.7 | 8.2 | 6.3E-10 | 0.3 | 8.2 | 7.1E-01 | -4.4 | 8.2 | 1.4E-08 | |
| RS007963 | 2.5 | 0.7 | 5.1E-06 | -0.1 | 0.7 | 9.2E-01 | -2.6 | 0.7 | 1.4E-05 | |
| RS007964 | 1.1 | 7.1 | 4.3E-08 | -0.1 | 7.1 | 8.0E-01 | -1.1 | 7.1 | 8.7E-09 | |
| RS008055 | 2.9 | 3.9 | 2.6E-33 | 1.8 | 3.9 | 3.4E-12 | -1.0 | 3.9 | 7.3E-06 | udp-glucuronosyltransferase 2b15-like |
| RS008107 | 1.6 | 5.8 | 1.3E-08 | -0.8 | 5.8 | 1.1E-02 | -2.4 | 5.8 | 1.2E-16 | nose resistant to fluoxetine protein 6 |
| RS008150 | 4.4 | 2.6 | 1.3E-08 | 1.5 | 2.6 | 1.4E-01 | -2.9 | 2.6 | 9.4E-05 | |
| RS008284 | 2.2 | 6.6 | 2.2E-30 | 0.6 | 6.6 | 1.5E-02 | -1.6 | 6.6 | 5.0E-17 | protein |
| RS008380 | 1.9 | 6.1 | 9.4E-10 | 0.6 | 6.1 | 6.1E-02 | -1.2 | 6.1 | 1.4E-04 | |
| RS008497 | 1.0 | 4.7 | 1.6E-03 | -0.8 | 4.7 | 1.7E-02 | -1.8 | 4.7 | 1.5E-08 | alpha-tocopherol transfer |
| RS008617 | 1.8 | 4.9 | 3.2E-22 | 0.5 | 4.9 | 4.9E-02 | -1.3 | 4.9 | 2.8E-12 | proton-coupled amino acid transporter 1 |
| RS008618 | 1.9 | 4.8 | 2.5E-18 | 0.8 | 4.8 | 4.0E-03 | -1.1 | 4.8 | 3.8E-07 | proton-coupled amino acid transporter 1-like |
| RS008630 | 4.3 | 4.6 | 1.2E-28 | 3.1 | 4.6 | 1.1E-14 | -1.2 | 4.6 | 8.2E-04 | |
| RS008823 | 5.6 | 5.1 | 3.1E-82 | 2.0 | 5.1 | 6.2E-06 | -3.6 | 5.1 | 2.9E-49 | |
| RS008824 | 6.2 | 1.0 | 2.0E-12 | 4.3 | 1.0 | 4.2E-04 | -1.9 | 1.0 | 1.4E-03 | |
| RS009030 | 1.8 | 8.0 | 6.8E-16 | 0.7 | 8.0 | 3.8E-03 | -1.1 | 8.0 | 2.6E-06 | |
| RS009112 | 3.3 | 1.6 | 6.3E-04 | 0.5 | 1.6 | 7.0E-01 | -2.8 | 1.6 | 5.0E-03 | |
| RS009627 | 1.1 | 11.8 | 6.0E-05 | 0.0 | 11.8 | 9.6E-01 | -1.1 | 11.8 | 1.5E-04 | heat shock 70 kda protein cognate 4 |
| RS009746 | 1.6 | 8.1 | 1.1E-21 | 0.2 | 8.1 | 2.3E-01 | -1.3 | 8.1 | 3.3E-15 | sphingomyelin phosphodiesterase-like |
| RS009878 | 1.0 | 4.9 | 1.1E-05 | -0.5 | 4.9 | 7.6E-02 | -1.5 | 4.9 | 1.8E-10 | |
| RS009887 | 1.0 | 9.5 | 6.4E-04 | -1.5 | 9.5 | 8.5E-08 | -2.6 | 9.5 | 8.4E-19 | |
| RS009917 | 3.0 | 5.9 | 2.1E-26 | 1.8 | 5.9 | 3.8E-10 | -1.2 | 5.9 | 3.4E-05 | phosphoenolpyruvate carboxykinase |
| RS009933 | 1.1 | 9.6 | 2.4E-09 | -0.2 | 9.6 | 3.7E-01 | -1.3 | 9.6 | 2.1E-12 | |

| | | | | | | | | | | |
|---|---|---|---|---|---|---|---|---|---|---|
| RS009934 | 1.5 | 7.6 | 4.8E-19 | 0.2 | 7.6 | 4.8E-01 | -1.3 | 7.6 | 1.3E-14 | |
| RS009935 | 1.5 | 8.4 | 5.0E-13 | -0.2 | 8.4 | 5.0E-01 | -1.7 | 8.4 | 2.1E-15 | pdz and lim domain protein 3 |
| RS010007 | 2.0 | 4.3 | 3.3E-20 | 0.8 | 4.3 | 4.9E-04 | -1.1 | 4.3 | 2.8E-07 | monocarboxylate transporter 12 |
| RS010105 | 1.7 | 6.6 | 1.3E-07 | 0.3 | 6.6 | 4.7E-01 | -1.4 | 6.6 | 3.0E-05 | elongation of very long chain fatty acids protein 6 |
| RS010158 | 6.8 | 4.0 | 3.4E-13 | 3.0 | 4.0 | 7.0E-04 | -3.8 | 4.0 | 8.4E-06 | |
| RS010570 | 1.7 | 9.4 | 3.4E-19 | 0.2 | 9.4 | 4.3E-01 | -1.5 | 9.4 | 6.8E-15 | sarcalumenin |
| RS010590 | 1.9 | 7.3 | 4.2E-18 | -0.2 | 7.3 | 4.0E-01 | -2.1 | 7.3 | 6.6E-22 | acyl- synthetase short-chain family member mitochondrial |
| RS010617 | 4.8 | 11.3 | 3.7E-07 | 1.8 | 11.3 | 4.7E-02 | -3.0 | 11.3 | 1.7E-03 | |
| RS010842 | 3.2 | 6.9 | 4.1E-10 | 0.4 | 6.9 | 4.8E-01 | -2.8 | 6.9 | 1.8E-07 | |
| RS010843 | 1.7 | 12.2 | 1.2E-08 | 0.5 | 12.2 | 1.8E-01 | -1.2 | 12.2 | 8.9E-05 | |
| RS010847 | 2.2 | 2.1 | 9.6E-09 | -0.1 | 2.1 | 9.5E-01 | -2.3 | 2.1 | 5.8E-08 | helix-loop-helix protein delilah |
| RS011058 | 1.2 | 3.4 | 7.7E-06 | -0.1 | 3.4 | 7.4E-01 | -1.4 | 3.4 | 3.5E-06 | serpin b12 |
| RS011174 | 1.2 | 7.0 | 2.2E-14 | -0.1 | 7.0 | 8.3E-01 | -1.3 | 7.0 | 9.0E-15 | haloacid dehalogenase-like hydrolase domain-containing protein 2 |
| RS011290 | 1.3 | 3.1 | 1.6E-04 | -0.6 | 3.1 | 1.6E-01 | -2.0 | 3.1 | 1.3E-07 | |
| RS011326 | 2.1 | 2.4 | 1.4E-06 | 0.7 | 2.4 | 1.9E-01 | -1.4 | 2.4 | 2.9E-03 | fatty acyl- reductase cg5065 |
| RS011333 | 9.0 | 8.7 | 1.3E-74 | 7.6 | 8.7 | 2.6E-58 | -1.4 | 8.7 | 6.2E-04 | hemolymph lipopolysaccharide-binding |
| RS011335 | 2.0 | 3.4 | 5.3E-07 | 0.7 | 3.4 | 2.0E-01 | -1.3 | 3.4 | 2.2E-03 | neuromedin-b receptor |
| RS011410 | 1.9 | 6.9 | 4.2E-33 | 0.7 | 6.9 | 1.4E-04 | -1.2 | 6.9 | 1.6E-13 | protein |
| RS011411 | 2.0 | 6.3 | 1.5E-20 | 0.0 | 6.3 | 9.5E-01 | -2.0 | 6.3 | 6.3E-20 | protein isoform b-like |
| RS011412 | 2.2 | 5.0 | 1.2E-24 | -0.3 | 5.0 | 4.4E-01 | -2.4 | 5.0 | 5.5E-27 | endonuclease-reverse transcriptase |
| RS011521 | 1.8 | 8.4 | 5.3E-03 | 0.0 | 8.4 | 9.6E-01 | -1.8 | 8.4 | 6.2E-03 | retinol dehydrogenase 14 |
| RS011651 | 1.6 | 8.9 | 6.9E-18 | -0.8 | 8.9 | 3.7E-05 | -2.4 | 8.9 | 2.5E-37 | fatty acid synthase |
| RS011668 | 3.0 | 4.9 | 9.5E-12 | 1.7 | 4.9 | 2.3E-04 | -1.3 | 4.9 | 5.1E-03 | |
| RS011688 | 2.1 | 7.7 | 3.7E-19 | 1.0 | 7.7 | 3.1E-05 | -1.1 | 7.7 | 1.5E-05 | alpha-tocopherol transfer |
| RS011689 | 10.1 | 7.6 | 2.4E-79 | 0.0 | 7.6 | 9.7E-01 | -10.1 | 7.6 | 1.2E-76 | alpha-tocopherol transfer |
| RS011693 | 3.6 | 6.4 | 7.9E-26 | 1.1 | 6.4 | 1.8E-03 | -2.5 | 6.4 | 1.2E-13 | alpha-tocopherol transfer |
| RS011713 | 6.3 | 3.6 | 2.1E-20 | 4.1 | 3.6 | 1.1E-07 | -2.2 | 3.6 | 2.4E-05 | glucose dehydrogenase |
| RS011716 | 3.4 | 1.3 | 6.5E-06 | 0.9 | 1.3 | 3.8E-01 | -2.5 | 1.3 | 1.2E-03 | glucose dehydrogenase |
| RS011839 | 1.2 | 3.1 | 3.4E-03 | -0.2 | 3.1 | 7.8E-01 | -1.4 | 3.1 | 2.2E-03 | |
| RS011966 | 1.2 | 6.0 | 9.9E-12 | -0.4 | 6.0 | 1.3E-01 | -1.6 | 6.0 | 6.1E-17 | set and mynd domain-containing protein 4-like |
| RS011982 | 2.6 | 3.8 | 2.0E-16 | -0.1 | 3.8 | 8.9E-01 | -2.7 | 3.8 | 1.2E-15 | non-ltr retrotransposon cats |
| RS011988 | 1.7 | 7.0 | 2.0E-15 | 0.5 | 7.0 | 4.1E-02 | -1.2 | 7.0 | 8.2E-08 | aquaporin-11 |
| RS012024 | 1.1 | 3.8 | 4.0E-03 | -0.6 | 3.8 | 1.6E-01 | -1.7 | 3.8 | 6.3E-06 | anosmin-1 |
| RS012035 | 4.6 | 2.2 | 2.1E-09 | 2.7 | 2.2 | 9.1E-04 | -1.9 | 2.2 | 9.8E-03 | |
| RS012245 | 1.4 | 5.1 | 3.1E-05 | 0.1 | 5.1 | 7.7E-01 | -1.2 | 5.1 | 4.1E-04 | hypothetical protein |
| RS012278 | 2.2 | 3.8 | 1.2E-09 | 1.0 | 3.8 | 1.6E-02 | -1.3 | 3.8 | 1.2E-03 | radical s-adenosyl methionine domain-containing protein 2 |
| RS012481 | 2.8 | 5.2 | 6.9E-17 | 0.2 | 5.2 | 6.0E-01 | -2.5 | 5.2 | 8.8E-14 | protein giant-lens |
| RS012544 | 2.2 | 6.6 | 2.4E-36 | 0.6 | 6.6 | 1.5E-03 | -1.6 | 6.6 | 1.4E-19 | sec23-interacting |
| RS012583 | 1.1 | 5.9 | 1.5E-06 | -0.1 | 5.9 | 6.9E-01 | -1.2 | 5.9 | 1.9E-07 | |
| RS012627 | 1.3 | 7.6 | 6.3E-09 | 0.2 | 7.6 | 5.3E-01 | -1.1 | 7.6 | 1.8E-06 | |
| RS012628 | 1.2 | 9.4 | 6.4E-11 | 0.1 | 9.4 | 7.4E-01 | -1.1 | 9.4 | 3.9E-09 | sterile alpha and tir motif-containing protein 1 |
| RS012664 | 2.1 | 4.0 | 3.1E-05 | 0.7 | 4.0 | 3.6E-01 | -1.4 | 4.0 | 7.3E-03 | 17-beta-hydroxysteroid dehydrogenase 13-like |
| RS012781 | 1.3 | 5.4 | 2.9E-06 | -1.6 | 5.4 | 5.7E-08 | -2.9 | 5.4 | 9.7E-24 | transforming growth factor-beta-induced protein ig-h3-like |

| | | | | | | | | | | |
|---|---|---|---|---|---|---|---|---|---|---|
| RS012784 | 1.1 | 7.0 | 1.2E-11 | 0.0 | 7.0 | 9.9E-01 | -1.1 | 7.0 | 7.5E-11 | excitatory amino acid transporter-like |
| RS012944 | 2.1 | 5.9 | 4.2E-32 | 0.4 | 5.9 | 3.9E-02 | -1.6 | 5.9 | 4.0E-20 | isopentenyl-diphosphate delta-isomerase 1 |
| RS012955 | 1.6 | 9.6 | 6.4E-07 | 0.1 | 9.6 | 7.7E-01 | -1.4 | 9.6 | 1.1E-05 | flightin |
| RS013098 | 2.8 | 6.1 | 1.7E-23 | 1.6 | 6.1 | 4.3E-08 | -1.2 | 6.1 | 2.1E-05 | xanthine dehydrogenase-like |
| RS013244 | 1.8 | 3.0 | 3.1E-11 | -0.3 | 3.0 | 4.2E-01 | -2.1 | 3.0 | 1.9E-13 | acyl- delta desaturase-like |
| RS013246 | 1.7 | 2.4 | 3.4E-05 | 0.1 | 2.4 | 8.6E-01 | -1.6 | 2.4 | 4.3E-04 | acyl- delta desaturase-like |
| RS013269 | 1.4 | 1.2 | 4.0E-04 | -0.4 | 1.2 | 5.9E-01 | -1.8 | 1.2 | 9.7E-05 | agap003468-pa-like protein |
| RS013361 | 1.3 | 3.8 | 4.0E-07 | -0.6 | 3.8 | 9.6E-02 | -1.9 | 3.8 | 9.6E-12 | |
| RS013376 | 1.6 | 4.4 | 4.5E-12 | 0.2 | 4.4 | 4.8E-01 | -1.4 | 4.4 | 1.4E-08 | |
| RS013379 | 3.2 | 5.7 | 1.7E-15 | 1.4 | 5.7 | 1.0E-03 | -1.8 | 5.7 | 8.0E-06 | |
| RS013610 | 3.1 | 6.3 | 9.6E-27 | 1.4 | 6.3 | 6.1E-06 | -1.7 | 6.3 | 3.5E-09 | glucose dehydrogenase |
| RS013626 | 1.2 | 3.9 | 3.3E-03 | 0.0 | 3.9 | 9.5E-01 | -1.2 | 3.9 | 4.2E-03 | |
| RS013635 | 7.8 | 4.9 | 5.0E-52 | -0.6 | 4.9 | 6.6E-01 | -8.4 | 4.9 | 4.8E-51 | cytochrome p450 4c1 |
| RS013637 | 1.5 | 5.4 | 3.2E-08 | -0.4 | 5.4 | 1.8E-01 | -1.9 | 5.4 | 3.5E-12 | hypothetical protein |
| RS013686 | 2.1 | 7.7 | 6.9E-18 | 0.3 | 7.7 | 3.2E-01 | -1.8 | 7.7 | 4.1E-13 | acyl- delta desaturase-like |
| RS013687 | 1.7 | 3.9 | 2.7E-11 | -0.4 | 3.9 | 2.4E-01 | -2.1 | 3.9 | 4.2E-15 | acyl- delta desaturase |
| RS013688 | 6.4 | 8.2 | 8.8E-43 | 4.9 | 8.2 | 4.6E-28 | -1.6 | 8.2 | 3.0E-04 | hypothetical protein |
| RS013837 | 1.1 | 4.1 | 2.5E-05 | 0.0 | 4.1 | 9.7E-01 | -1.1 | 4.1 | 7.5E-05 | leucine-rich repeat-containing protein 24-like |
| RS014029 | 1.1 | 3.7 | 6.3E-04 | 0.0 | 3.7 | 9.8E-01 | -1.1 | 3.7 | 1.8E-03 | |
| RS014071 | 5.5 | 1.3 | 4.4E-13 | 2.1 | 1.3 | 1.1E-01 | -3.4 | 1.3 | 1.9E-07 | |
| RS014220 | 1.4 | 10.2 | 1.1E-08 | 0.3 | 10.2 | 3.3E-01 | -1.1 | 10.2 | 1.6E-05 | alaserpin |
| RS014541 | 5.4 | 2.9 | 1.8E-16 | 3.3 | 2.9 | 5.3E-07 | -2.1 | 2.9 | 4.5E-04 | aael014316 |
| RS014553 | 1.6 | 1.8 | 3.6E-05 | -0.9 | 1.8 | 8.2E-02 | -2.5 | 1.8 | 3.7E-09 | fatty acid synthase-like |
| RS014585 | 5.6 | 7.3 | 2.1E-19 | 3.3 | 7.3 | 4.1E-06 | -2.4 | 7.3 | 6.7E-06 | hypothetical protein |
| RS014597 | 8.5 | 7.2 | 5.9E-07 | 3.7 | 7.2 | 2.0E-02 | -4.8 | 7.2 | 2.7E-03 | |
| RS014599 | 6.1 | 8.7 | 1.4E-04 | 1.0 | 8.7 | 6.0E-01 | -5.1 | 8.7 | 2.6E-03 | |
| RS014620 | 2.8 | 6.5 | 3.1E-16 | 1.3 | 6.5 | 3.5E-04 | -1.5 | 6.5 | 2.3E-05 | glutamate synthase |
| RS014656 | 1.5 | 3.6 | 3.9E-04 | 0.3 | 3.6 | 7.0E-01 | -1.2 | 3.6 | 6.9E-03 | neurotrimin |
| RS014698 | 5.2 | 8.4 | 3.0E-10 | 2.5 | 8.4 | 1.4E-03 | -2.6 | 8.4 | 1.0E-03 | lysozyme-like |
| RS014749 | 2.2 | 2.3 | 4.8E-05 | 0.1 | 2.3 | 9.5E-01 | -2.2 | 2.3 | 3.8E-04 | tigger transposable element-derived protein 6-like |
| RS014956 | 3.1 | 3.6 | 7.5E-13 | 1.2 | 3.6 | 9.8E-03 | -1.9 | 3.6 | 1.6E-05 | |
| RS015219 | 2.5 | 4.0 | 2.9E-15 | 1.1 | 4.0 | 3.5E-03 | -1.4 | 4.0 | 1.0E-05 | probable cytochrome p450 6a13 |
| RS015220 | 3.2 | 3.5 | 2.1E-24 | 1.5 | 3.5 | 1.5E-04 | -1.6 | 3.5 | 9.9E-09 | cytochrome p450 6k1-like |
| RS015221 | 2.0 | 4.8 | 1.2E-26 | 0.3 | 4.8 | 3.8E-01 | -1.8 | 4.8 | 1.1E-19 | cytochrome p450 |
| RS015308 | 1.7 | 4.0 | 8.0E-08 | 0.4 | 4.0 | 3.6E-01 | -1.3 | 4.0 | 7.9E-05 | |
| RS015310 | 1.2 | 5.3 | 6.2E-08 | -0.1 | 5.3 | 6.5E-01 | -1.4 | 5.3 | 4.6E-09 | atp-binding cassette sub-family g member 1 |
| RS015365 | 1.9 | 1.8 | 1.1E-04 | -4.2 | 1.8 | 1.8E-08 | -6.1 | 1.8 | 3.7E-19 | |
| RS100004 | 3.0 | 4.5 | 2.1E-12 | 1.4 | 4.5 | 2.3E-03 | -1.7 | 4.5 | 2.3E-04 | lysozyme-like |
| RS100006 | 1.5 | 8.0 | 1.1E-06 | 0.5 | 8.0 | 1.8E-01 | -1.0 | 8.0 | 1.8E-03 | beta-glucosidase |
| RS100010 | 3.9 | 4.8 | 2.0E-11 | -0.6 | 4.8 | 4.3E-01 | -4.5 | 4.8 | 2.0E-13 | geranylgeranyl pyrophosphate synthase |
| RS100012 | 3.6 | 5.2 | 9.6E-06 | -0.1 | 5.2 | 9.6E-01 | -3.6 | 5.2 | 1.3E-05 | geranylgeranyl pyrophosphate synthase |
| RS100013 | 2.8 | 4.2 | 7.0E-05 | -0.1 | 4.2 | 9.2E-01 | -2.9 | 4.2 | 3.6E-04 | geranylgeranyl pyrophosphate synthase |
| RS100015 | 9.9 | 7.4 | 1.0E-60 | 1.6 | 7.4 | 2.9E-02 | -8.3 | 7.4 | 7.0E-51 | geranylgeranyl pyrophosphate synthase |
| RS100016 | 8.1 | 8.9 | 1.1E-63 | 3.6 | 8.9 | 3.7E-17 | -4.6 | 8.9 | 1.1E-28 | geranylgeranyl pyrophosphate synthase |

| | | | | | | | | | | |
|---|---|---|---|---|---|---|---|---|---|---|
| RS100017 | 7.0 | 4.2 | 4.7E-24 | -0.9 | 4.2 | 5.9E-01 | -8.0 | 4.2 | 1.1E-23 | geranylgeranyl pyrophosphate synthase |
| RS100019 | 8.5 | 4.9 | 1.7E-28 | 5.0 | 4.9 | 4.3E-12 | -3.5 | 4.9 | 1.5E-08 | peptidoglycan-recognition protein precursor |
| RS100021 | 7.1 | 6.8 | 4.1E-15 | 3.7 | 6.8 | 8.9E-06 | -3.4 | 6.8 | 3.7E-05 | c-type lysozyme-2 |
| RS100022 | 5.8 | 7.4 | 1.3E-14 | 2.8 | 7.4 | 6.5E-05 | -3.0 | 7.4 | 2.9E-05 | lysozyme |
| RS100023 | 5.2 | 6.4 | 1.4E-10 | 2.6 | 6.4 | 8.3E-04 | -2.7 | 6.4 | 8.8E-04 | lysozyme-like |

SI_Dataset_1
Caste-biased genes: Worker (head)

| Gene ID | Head Reproductive v.s. Soldier | | | Head Reproductive v.s. Worker | | | Head Soldier v.s. Worker | | | Gene annotation |
|---|---|---|---|---|---|---|---|---|---|---|
| | logFC | logCPM | FDR | logFC | logCPM | FDR | logFC | logCPM | FDR | |
| RS000318 | 0.4 | 5.8 | 1.5E-01 | 1.5 | 5.8 | 1.9E-08 | 1.0 | 5.8 | 1.4E-04 | alpha-aminoadipic semialdehyde dehydrogenase |
| RS000709 | 0.5 | 0.5 | 7.7E-01 | 3.0 | 0.5 | 3.4E-04 | 2.5 | 0.5 | 3.3E-03 | aldose reductase-like |
| RS000846 | 0.2 | 11.1 | 9.2E-01 | 5.4 | 11.1 | 1.9E-05 | 5.2 | 11.1 | 6.2E-06 | hemocyanin subunit type 1 precursor |
| RS000959 | -0.2 | 7.0 | 9.4E-01 | 2.7 | 7.0 | 1.5E-03 | 2.9 | 7.0 | 1.6E-03 | lipase 3-like |
| RS001104 | 0.5 | 0.7 | 6.9E-01 | 2.7 | 0.7 | 1.8E-04 | 2.2 | 0.7 | 3.4E-03 | |
| RS001196 | 4.4 | 10.9 | 6.7E-19 | 9.4 | 10.9 | 2.4E-53 | 5.0 | 10.9 | 1.2E-22 | |
| RS001197 | 2.4 | 13.0 | 7.8E-15 | 3.7 | 13.0 | 2.9E-31 | 1.4 | 13.0 | 1.6E-05 | |
| RS001244 | -0.7 | 1.0 | 1.6E-01 | 1.3 | 1.0 | 1.3E-05 | 2.0 | 1.0 | 6.3E-09 | ef-hand calcium-binding domain-containing protein 1 |
| RS001265 | -0.2 | 2.9 | 9.0E-01 | 3.1 | 2.9 | 1.1E-03 | 3.3 | 2.9 | 1.8E-03 | elongation of very long chain fatty acids protein 7-like |
| RS001343 | -0.2 | 3.5 | 8.2E-01 | 1.5 | 3.5 | 8.4E-03 | 1.8 | 3.5 | 4.4E-03 | sodium potassium calcium exchanger 4 |
| RS001416 | -0.9 | 6.4 | 2.1E-03 | 1.6 | 6.4 | 7.4E-09 | 2.4 | 6.4 | 3.1E-20 | acyl- synthetase family member mitochondrial-like |
| RS001417 | -0.6 | 5.0 | 1.2E-01 | 1.1 | 5.0 | 1.9E-03 | 1.7 | 5.0 | 2.6E-07 | fatty acid-- ligase |
| RS001419 | -0.2 | 1.3 | 9.1E-01 | 1.9 | 1.3 | 8.4E-03 | 2.1 | 1.3 | 7.3E-03 | acyl- synthetase family member mitochondrial |
| RS001420 | -2.0 | 4.6 | 2.1E-03 | 1.9 | 4.6 | 3.2E-04 | 4.0 | 4.6 | 1.0E-11 | acyl- synthetase family member mitochondrial-like |
| RS001498 | 0.3 | 0.8 | 8.6E-01 | 3.1 | 0.8 | 5.3E-05 | 2.8 | 0.8 | 3.1E-04 | hypothetical protein |
| RS001693 | 0.3 | 4.2 | 3.6E-01 | 1.5 | 4.2 | 1.6E-07 | 1.2 | 4.2 | 6.0E-05 | lipase member h-like |
| RS001763 | -0.3 | 7.6 | 4.4E-01 | 1.0 | 7.6 | 1.6E-03 | 1.3 | 7.6 | 9.7E-06 | dehydrogenase reductase sdr family member 11 |
| RS001819 | -0.4 | 4.1 | 7.4E-02 | 1.7 | 4.1 | 3.5E-22 | 2.1 | 4.1 | 2.5E-31 | |
| RS001853 | 0.2 | 2.7 | 7.4E-01 | 1.9 | 2.7 | 5.5E-10 | 1.7 | 2.7 | 4.2E-08 | achaete-scute complex protein t3-like |
| RS001896 | 3.1 | 6.1 | 9.3E-05 | 7.0 | 6.1 | 2.7E-25 | 3.9 | 6.1 | 6.5E-13 | |
| RS002141 | 0.5 | 4.6 | 3.0E-01 | 1.8 | 4.6 | 9.7E-06 | 1.3 | 4.6 | 1.6E-03 | |
| RS002475 | -1.8 | 5.8 | 4.8E-04 | 2.0 | 5.8 | 3.7E-07 | 3.8 | 5.8 | 1.9E-17 | |
| RS002521 | -0.1 | 7.6 | 7.3E-01 | 1.1 | 7.6 | 9.7E-09 | 1.1 | 7.6 | 1.6E-10 | dehydrogenase reductase sdr family member 11 |
| RS003152 | -1.1 | 6.7 | 1.1E-04 | 1.1 | 6.7 | 3.5E-04 | 2.2 | 6.7 | 2.0E-15 | probable cytochrome p450 6a14 |
| RS003155 | 1.1 | 5.7 | 8.3E-04 | 2.4 | 5.7 | 9.3E-14 | 1.3 | 5.7 | 1.2E-04 | probable cytochrome p450 6a14 |
| RS003156 | 0.8 | 7.6 | 1.9E-02 | 2.1 | 7.6 | 1.3E-09 | 1.2 | 7.6 | 5.7E-04 | cytochrome p450 |
| RS003157 | 0.5 | 7.0 | 1.6E-01 | 1.9 | 7.0 | 2.0E-08 | 1.3 | 7.0 | 8.4E-05 | cytochrome p450 |
| RS003185 | -0.7 | 0.3 | 6.9E-01 | 3.3 | 0.3 | 8.4E-07 | 3.9 | 0.3 | 1.7E-06 | -related lipid transfer protein 3 |
| RS003282 | 0.2 | 3.5 | 8.0E-01 | 2.5 | 3.5 | 5.3E-11 | 2.3 | 3.5 | 5.8E-09 | aldose reductase-like |
| RS003347 | 1.0 | 1.9 | 1.1E-01 | 2.5 | 1.9 | 1.4E-06 | 1.6 | 1.9 | 3.6E-03 | a disintegrin and metalloproteinase with thrombospondin motifs 18 |
| RS003352 | -0.5 | 2.3 | 1.1E-01 | 1.1 | 2.3 | 5.3E-05 | 1.6 | 2.3 | 1.8E-09 | |
| RS003636 | -0.3 | 5.5 | 1.9E-01 | 1.2 | 5.5 | 4.7E-12 | 1.5 | 5.5 | 1.8E-17 | atp-binding cassette sub-family g member 4 |
| RS003788 | 0.5 | 7.2 | 1.4E-01 | 1.5 | 7.2 | 9.0E-08 | 1.0 | 7.2 | 4.1E-04 | probable cytochrome p450 304a1 |
| RS003831 | 0.0 | 6.3 | 9.9E-01 | 2.4 | 6.3 | 1.7E-03 | 2.4 | 6.3 | 9.6E-04 | |

| | | | | | | | | | | |
|---|---|---|---|---|---|---|---|---|---|---|
| RS004001 | -0.1 | 1.3 | 8.5E-01 | 1.9 | 1.3 | 2.1E-08 | 2.1 | 1.3 | 4.6E-08 | estrogen sulfotransferase |
| RS004129 | 0.8 | 6.3 | 3.4E-03 | 2.7 | 6.3 | 1.5E-27 | 1.9 | 6.3 | 4.3E-15 | venom carboxylesterase-6-like |
| RS004377 | 0.0 | 3.8 | 9.7E-01 | 1.1 | 3.8 | 8.3E-05 | 1.2 | 3.8 | 5.5E-05 | |
| RS004378 | -0.3 | 0.8 | 8.4E-01 | 2.9 | 0.8 | 7.3E-04 | 3.2 | 0.8 | 6.0E-04 | |
| RS004427 | -0.1 | 4.0 | 8.7E-01 | 1.0 | 4.0 | 1.4E-03 | 1.1 | 4.0 | 4.3E-04 | arylsulfatase b |
| RS004435 | -0.6 | 6.6 | 1.2E-04 | 1.2 | 6.6 | 3.0E-16 | 1.8 | 6.6 | 2.0E-35 | monocarboxylate transporter 13 |
| RS004535 | 0.6 | 2.3 | 1.1E-01 | 1.7 | 2.3 | 2.0E-11 | 1.2 | 2.3 | 1.4E-05 | c- |
| RS004557 | 1.2 | 0.6 | 2.1E-01 | 3.2 | 0.6 | 1.2E-05 | 2.0 | 0.6 | 5.9E-03 | cuticle protein 19 |
| RS004657 | 1.7 | 9.2 | 1.1E-05 | 3.0 | 9.2 | 8.7E-15 | 1.3 | 9.2 | 8.4E-04 | apolipoprotein d |
| RS004767 | -1.1 | 6.1 | 2.9E-02 | 1.6 | 6.1 | 1.2E-03 | 2.7 | 6.1 | 3.9E-09 | |
| RS005263 | 1.7 | 4.0 | 4.3E-04 | 4.6 | 4.0 | 1.6E-23 | 2.9 | 4.0 | 8.3E-12 | |
| RS005687 | -0.1 | 1.3 | 9.4E-01 | 1.6 | 1.3 | 9.7E-03 | 1.7 | 1.3 | 7.0E-03 | endochitinase |
| RS005959 | 1.2 | 1.7 | 7.4E-02 | 3.8 | 1.7 | 3.2E-15 | 2.6 | 1.7 | 1.9E-08 | transcription factor glial cells missing |
| RS006066 | 0.8 | 5.0 | 2.8E-02 | 2.4 | 5.0 | 3.8E-17 | 1.6 | 5.0 | 3.4E-08 | d-arabinitol dehydrogenase 1-like |
| RS006091 | 0.6 | 4.4 | 3.1E-01 | 2.3 | 4.4 | 1.1E-08 | 1.7 | 4.4 | 2.3E-05 | |
| RS006132 | -0.4 | 1.3 | 6.4E-01 | 1.6 | 1.3 | 4.9E-03 | 1.9 | 1.3 | 4.0E-04 | |
| RS006136 | -0.7 | 5.4 | 6.2E-01 | 2.2 | 5.4 | 5.6E-03 | 2.9 | 5.4 | 9.2E-04 | alpha-amylase |
| RS006137 | 0.3 | 7.1 | 7.2E-01 | 3.3 | 7.1 | 2.6E-16 | 3.0 | 7.1 | 2.1E-12 | alpha-amylase |
| RS006197 | 3.8 | 8.6 | 2.5E-12 | 5.8 | 8.6 | 1.7E-29 | 2.1 | 8.6 | 7.4E-07 | maltase a2 |
| RS006264 | 0.0 | 7.4 | 9.7E-01 | 2.1 | 7.4 | 2.5E-03 | 2.1 | 7.4 | 1.0E-03 | fatty acyl- reductase 1-like |
| RS006303 | 0.7 | 2.6 | 2.0E-01 | 1.9 | 2.6 | 9.8E-07 | 1.2 | 2.6 | 2.7E-03 | female reproductive tract protease |
| RS006537 | 1.3 | 5.4 | 1.6E-06 | 3.2 | 5.4 | 2.0E-32 | 1.8 | 5.4 | 7.1E-13 | venom carboxylesterase-6-like |
| RS006538 | 1.3 | 4.5 | 1.2E-03 | 3.2 | 4.5 | 4.5E-18 | 1.9 | 4.5 | 8.9E-08 | venom carboxylesterase-6-like |
| RS006664 | -0.6 | 5.0 | 1.9E-01 | 2.3 | 5.0 | 4.3E-10 | 2.9 | 5.0 | 4.3E-15 | transforming growth factor-beta-induced protein ig-h3 |
| RS006870 | 1.3 | 2.0 | 1.5E-02 | 3.0 | 2.0 | 6.9E-14 | 1.7 | 2.0 | 8.6E-06 | vitamin k-dependent protein c-like |
| RS007008 | -1.6 | 1.4 | 3.0E-01 | 6.6 | 1.4 | 2.2E-20 | 8.1 | 1.4 | 1.2E-20 | udp-glucuronosyltransferase 2c1 |
| RS007081 | -0.2 | 3.9 | 5.5E-01 | 1.1 | 3.9 | 8.5E-07 | 1.3 | 3.9 | 6.5E-09 | e3 ubiquitin-protein ligase sinat3 |
| RS007163 | 1.7 | 8.2 | 3.9E-06 | 3.1 | 8.2 | 1.2E-17 | 1.4 | 8.2 | 8.7E-05 | adenosine deaminase |
| RS007214 | 0.3 | 7.7 | 6.1E-01 | 2.8 | 7.7 | 2.3E-10 | 2.5 | 7.7 | 7.8E-09 | |
| RS007302 | 0.6 | 1.6 | 5.1E-01 | 3.4 | 1.6 | 6.5E-08 | 2.8 | 1.6 | 6.8E-06 | zinc metalloproteinase nas |
| RS007323 | 0.8 | 1.3 | 1.4E-01 | 2.0 | 1.3 | 4.3E-07 | 1.2 | 1.3 | 3.1E-03 | |
| RS007488 | -0.2 | 9.7 | 4.0E-01 | 1.3 | 9.7 | 2.3E-11 | 1.5 | 9.7 | 1.5E-15 | glutathione s-transferase 1-1 |
| RS007834 | 0.2 | 5.0 | 7.9E-01 | 2.1 | 5.0 | 1.7E-04 | 1.9 | 5.0 | 5.8E-04 | |
| RS007982 | 0.4 | 5.2 | 3.8E-01 | 1.4 | 5.2 | 1.7E-04 | 1.0 | 5.2 | 9.6E-03 | |
| RS008098 | 0.8 | 5.6 | 2.5E-02 | 2.4 | 5.6 | 1.6E-22 | 1.7 | 5.6 | 1.4E-11 | |
| RS008121 | -0.5 | 1.0 | 4.7E-01 | 1.9 | 1.0 | 1.1E-05 | 2.4 | 1.0 | 1.7E-07 | endocuticle structural glycoprotein bd-2 |
| RS008126 | -1.1 | 6.4 | 1.9E-01 | 2.2 | 6.4 | 2.8E-06 | 3.3 | 6.4 | 3.3E-09 | endocuticle structural glycoprotein bd-2 |
| RS008129 | 0.2 | 2.0 | 8.7E-01 | 4.7 | 2.0 | 5.8E-11 | 4.5 | 2.0 | 9.0E-10 | endocuticle structural glycoprotein bd-2 |
| RS008250 | 0.0 | 6.2 | 1.0E+00 | 1.7 | 6.2 | 8.3E-14 | 1.7 | 6.2 | 1.0E-13 | dehydrogenase reductase sdr family member 11 |
| RS008251 | 0.0 | 5.8 | 9.5E-01 | 1.9 | 5.8 | 4.0E-14 | 1.9 | 5.8 | 1.3E-14 | dehydrogenase reductase sdr family member 11-like |
| RS008266 | -2.2 | 2.6 | 5.9E-09 | 1.4 | 2.6 | 1.2E-05 | 3.6 | 2.6 | 2.9E-24 | protein |
| RS008334 | -1.0 | 10.0 | 3.1E-11 | 1.5 | 10.0 | 4.8E-23 | 2.5 | 10.0 | 1.3E-61 | alcohol dehydrogenase |
| RS009343 | 0.1 | 5.9 | 8.3E-01 | 2.6 | 5.9 | 3.0E-15 | 2.5 | 5.9 | 5.8E-13 | pyrimidodiazepine synthase-like |
| RS009491 | 0.7 | 3.9 | 1.3E-01 | 2.6 | 3.9 | 2.9E-15 | 1.9 | 3.9 | 1.1E-08 | |

| | | | | | | | | | | |
|---|---|---|---|---|---|---|---|---|---|---|
| RS009630 | -0.3 | 3.2 | 6.8E-01 | 2.0 | 3.2 | 9.7E-04 | 2.3 | 3.2 | 3.7E-05 | protein lethal malignant blood neoplasm 1 |
| RS009651 | -0.2 | 6.3 | 6.2E-01 | 1.1 | 6.3 | 6.8E-04 | 1.3 | 6.3 | 1.8E-05 | maltase a1 |
| RS009664 | -0.1 | 10.8 | 6.3E-01 | 1.4 | 10.8 | 2.9E-14 | 1.5 | 10.8 | 3.0E-17 | |
| RS009950 | -0.1 | 1.9 | 8.7E-01 | 1.2 | 1.9 | 2.4E-04 | 1.3 | 1.9 | 1.4E-04 | pancreatic triacylglycerol lipase-like |
| RS009969 | 3.2 | 4.5 | 1.2E-02 | 8.4 | 4.5 | 3.5E-17 | 5.1 | 4.5 | 9.8E-11 | aldose reductase-like |
| RS010002 | -0.3 | 5.2 | 4.9E-01 | 1.0 | 5.2 | 1.9E-03 | 1.3 | 5.2 | 2.2E-05 | epoxide hydrolase 4-like |
| RS010059 | 1.4 | 4.4 | 2.9E-02 | 3.2 | 4.4 | 4.7E-07 | 1.7 | 4.4 | 6.7E-03 | |
| RS010065 | -1.7 | 2.7 | 1.4E-02 | 2.0 | 2.7 | 1.9E-03 | 3.7 | 2.7 | 7.5E-09 | dentin sialophospho |
| RS010159 | 0.6 | 4.8 | 1.6E-01 | 1.7 | 4.8 | 1.3E-05 | 1.1 | 4.8 | 8.5E-03 | delta-1-pyrroline-5-carboxylate synthase |
| RS010163 | -0.6 | 7.0 | 3.6E-04 | 1.5 | 7.0 | 1.9E-24 | 2.1 | 7.0 | 4.3E-45 | cytochrome p450 6k1-like |
| RS010164 | -1.2 | 7.2 | 4.0E-09 | 1.4 | 7.2 | 2.3E-12 | 2.5 | 7.2 | 6.9E-39 | cytochrome p450 6k1-like |
| RS010241 | -0.2 | 7.3 | 6.1E-01 | 1.1 | 7.3 | 1.1E-06 | 1.2 | 7.3 | 8.4E-09 | 15-hydroxyprostaglandin dehydrogenase |
| RS010272 | 0.9 | 6.9 | 4.1E-02 | 3.4 | 6.9 | 4.5E-22 | 2.5 | 6.9 | 6.2E-14 | glucose dehydrogenase |
| RS010441 | -0.2 | -0.1 | 9.1E-01 | 4.6 | -0.1 | 1.3E-05 | 4.8 | -0.1 | 2.1E-05 | |
| RS010442 | -2.5 | 8.3 | 1.5E-12 | 1.6 | 8.3 | 2.6E-05 | 4.1 | 8.3 | 2.1E-28 | |
| RS010443 | 0.0 | 3.3 | 1.0E+00 | 2.5 | 3.3 | 7.0E-19 | 2.5 | 3.3 | 2.0E-18 | pickpocket protein 28 |
| RS010594 | 1.1 | 5.0 | 7.9E-04 | 2.8 | 5.0 | 3.3E-19 | 1.6 | 5.0 | 7.0E-08 | |
| RS010778 | -2.5 | 5.6 | 6.1E-30 | 1.9 | 5.6 | 5.2E-21 | 4.4 | 5.6 | 2.2E-85 | lipase 3-like |
| RS010849 | -0.6 | 5.8 | 2.6E-01 | 2.6 | 5.8 | 4.2E-08 | 3.2 | 5.8 | 3.2E-12 | glycine n-methyltransferase |
| RS011205 | -0.9 | 11.6 | 2.7E-01 | 4.3 | 11.6 | 1.2E-09 | 5.2 | 11.6 | 1.8E-14 | larval serum protein 1 beta chain |
| RS011571 | -1.1 | 9.6 | 2.5E-04 | 2.7 | 9.6 | 2.7E-18 | 3.8 | 9.6 | 1.8E-34 | |
| RS011572 | -0.4 | 4.8 | 4.1E-01 | 1.5 | 4.8 | 6.2E-05 | 1.8 | 4.8 | 1.6E-07 | |
| RS011591 | -1.4 | 1.0 | 6.9E-02 | 1.7 | 1.0 | 8.6E-03 | 3.0 | 1.0 | 1.8E-06 | |
| RS011592 | -0.1 | 4.8 | 6.8E-01 | 1.6 | 4.8 | 8.3E-20 | 1.7 | 4.8 | 3.5E-22 | uncharacterized protein |
| RS011717 | 1.0 | 2.7 | 1.1E-01 | 3.0 | 2.7 | 1.7E-08 | 2.0 | 2.7 | 2.2E-04 | glucose dehydrogenase |
| RS011719 | 1.1 | 3.1 | 8.0E-04 | 2.3 | 3.1 | 1.0E-14 | 1.1 | 3.1 | 1.4E-04 | glucose dehydrogenase |
| RS011736 | -0.2 | 10.7 | 5.9E-01 | 1.3 | 10.7 | 3.7E-07 | 1.5 | 10.7 | 1.6E-09 | apolipophorin-iii |
| RS011787 | 0.4 | 4.8 | 1.2E-01 | 1.5 | 4.8 | 5.1E-10 | 1.0 | 4.8 | 2.1E-05 | zinc finger and btb domain-containing protein 49 |
| RS012436 | -0.5 | 5.0 | 2.1E-01 | 3.5 | 5.0 | 1.0E-31 | 4.0 | 5.0 | 7.7E-39 | beta-glucosidase |
| RS012437 | 0.2 | 4.2 | 8.2E-01 | 2.0 | 4.2 | 2.6E-07 | 1.8 | 4.2 | 3.8E-06 | beta-glucosidase |
| RS012471 | -0.8 | 1.8 | 4.4E-02 | 1.2 | 1.8 | 1.2E-04 | 2.1 | 1.8 | 7.7E-10 | |
| RS013528 | 0.8 | 1.5 | 6.2E-01 | 5.4 | 1.5 | 3.2E-07 | 4.6 | 1.5 | 1.1E-05 | |
| RS013529 | -0.2 | 1.6 | 1.0E+00 | 7.9 | 1.6 | 1.8E-04 | 8.2 | 1.6 | 2.4E-04 | pro-resilin |
| RS013530 | -0.2 | -0.1 | 1.0E+00 | 4.9 | -0.1 | 4.7E-03 | 5.2 | -0.1 | 4.9E-03 | pro-resilin |
| RS013531 | 3.2 | 1.2 | 3.1E-02 | 7.2 | 1.2 | 2.2E-07 | 4.0 | 1.2 | 2.1E-03 | pro-resilin |
| RS013532 | -1.0 | 1.6 | 7.0E-01 | 6.0 | 1.6 | 1.8E-04 | 7.0 | 1.6 | 5.7E-06 | pro-resilin |
| RS013606 | 0.0 | 0.6 | 1.0E+00 | 5.8 | 0.6 | 7.7E-05 | 5.8 | 0.6 | 1.9E-04 | glucose dehydrogenase |
| RS013629 | 0.2 | 1.1 | 8.0E-01 | 2.3 | 1.1 | 2.1E-04 | 2.1 | 1.1 | 9.6E-04 | hypothetical protein |
| RS013912 | 1.3 | 3.3 | 1.7E-01 | 6.3 | 3.3 | 2.1E-13 | 4.9 | 3.3 | 7.1E-10 | |
| RS013959 | -1.1 | 5.8 | 6.3E-03 | 1.5 | 5.8 | 5.0E-05 | 2.5 | 5.8 | 5.6E-13 | d-arabinitol dehydrogenase 1-like |
| RS014104 | -0.1 | 7.6 | 9.8E-01 | 3.1 | 7.6 | 2.3E-03 | 3.2 | 7.6 | 1.6E-03 | |
| RS014105 | 0.3 | 8.3 | 8.6E-01 | 3.3 | 8.3 | 1.0E-05 | 3.1 | 8.3 | 7.8E-05 | |
| RS014107 | 0.7 | 12.1 | 5.0E-01 | 3.1 | 12.1 | 8.6E-04 | 2.4 | 12.1 | 7.3E-03 | |
| RS014109 | 0.8 | 11.9 | 3.6E-01 | 2.9 | 11.9 | 1.9E-04 | 2.1 | 11.9 | 6.1E-03 | |
| RS014276 | -0.2 | 3.5 | 5.5E-01 | 1.3 | 3.5 | 3.0E-07 | 1.5 | 3.5 | 2.4E-09 | hypothetical protein |
| RS014291 | 0.6 | 7.0 | 2.3E-04 | 1.8 | 7.0 | 4.2E-30 | 1.2 | 7.0 | 1.6E-13 | cytochrome p450 cyp12a2-like |
| RS014292 | -0.4 | 6.9 | 3.5E-02 | 1.3 | 6.9 | 4.3E-14 | 1.7 | 6.9 | 1.6E-23 | cytochrome p450 cyp12a2-like |
| RS014293 | 0.9 | 2.5 | 1.6E-01 | 2.6 | 2.5 | 1.7E-09 | 1.7 | 2.5 | 7.7E-05 | cytochrome p450 |
| RS014433 | 0.4 | 4.5 | 2.1E-01 | 1.5 | 4.5 | 1.1E-08 | 1.1 | 4.5 | 4.3E-05 | |

| | | | | | | | | | | |
|---|---|---|---|---|---|---|---|---|---|---|
| RS014444 | 0.0 | 4.8 | 9.4E-01 | 1.3 | 4.8 | 1.9E-12 | 1.3 | 4.8 | 1.3E-12 | heparin cofactor 2 |
| RS014493 | -1.9 | 8.0 | 2.2E-06 | 2.1 | 8.0 | 2.5E-08 | 4.0 | 8.0 | 2.7E-24 | |
| RS014494 | -1.4 | 5.7 | 9.4E-06 | 1.8 | 5.7 | 1.0E-08 | 3.2 | 5.7 | 1.7E-24 | dehydrogenase reductase sdr family member 11 |
| RS014669 | 0.0 | 2.9 | 9.9E-01 | 1.4 | 2.9 | 5.1E-07 | 1.4 | 2.9 | 6.1E-07 | frizzled-9-like |
| RS014922 | 0.2 | 6.2 | 6.4E-01 | 2.0 | 6.2 | 2.4E-11 | 1.8 | 6.2 | 1.1E-09 | 15-hydroxyprostaglandin dehydrogenase |
| RS015383 | -0.5 | 1.4 | 4.7E-01 | 1.5 | 1.4 | 6.3E-03 | 2.0 | 1.4 | 1.4E-04 | mucin-19-like |
| RS100005 | 0.3 | 4.3 | 6.1E-01 | 1.9 | 4.3 | 5.0E-07 | 1.6 | 4.3 | 5.0E-05 | myrosinase 1-like |

SI_Dataset_1
Caste-biased genes: Worker (thorax + abdomen)

| Gene ID | Thorax + abdomen Reproductive v.s. Soldier | | | Thorax + abdomen Reproductive v.s. Worker | | | Thorax + abdomen Soldier v.s. Worker | | | Gene annotation |
|---|---|---|---|---|---|---|---|---|---|---|
| | logFC | logCPM | FDR | logFC | logCPM | FDR | logFC | logCPM | FDR | |
| RS000494 | 2.0 | 0.2 | 1.2E-02 | 3.7 | 0.2 | 7.7E-10 | 1.7 | 0.2 | 1.5E-03 | |
| RS000777 | 0.0 | 1.2 | 9.6E-01 | 1.6 | 1.2 | 2.1E-03 | 1.6 | 1.2 | 4.5E-03 | |
| RS000778 | 0.9 | 3.4 | 1.8E-02 | 1.9 | 3.4 | 1.7E-08 | 1.0 | 3.4 | 4.9E-03 | |
| RS000846 | -1.8 | 11.1 | 1.8E-01 | 4.0 | 11.1 | 1.2E-03 | 5.8 | 11.1 | 1.4E-06 | hemocyanin subunit type 1 precursor |
| RS000956 | 0.1 | 3.5 | 7.8E-01 | 1.2 | 3.5 | 2.4E-09 | 1.2 | 3.5 | 8.3E-08 | nose resistant to fluoxetine protein 6-like |
| RS001085 | 0.4 | 6.2 | 1.7E-01 | 1.5 | 6.2 | 9.5E-09 | 1.1 | 6.2 | 1.1E-04 | endoplasmic reticulum resident protein 29 |
| RS001256 | -0.1 | 6.8 | 7.2E-01 | 1.1 | 6.8 | 4.7E-05 | 1.2 | 6.8 | 9.3E-06 | protein disulfide-isomerase a6 |
| RS001294 | 1.4 | 4.5 | 1.8E-03 | 2.8 | 4.5 | 4.8E-11 | 1.4 | 4.5 | 1.4E-03 | |
| RS001358 | -0.9 | 6.8 | 1.9E-03 | 1.2 | 6.8 | 1.8E-05 | 2.1 | 6.8 | 1.9E-14 | c-5 sterol desaturase erg32-like |
| RS001433 | 0.3 | 6.4 | 4.1E-01 | 1.8 | 6.4 | 1.1E-08 | 1.4 | 6.4 | 4.5E-06 | niemann-pick c1 protein |
| RS001583 | 0.8 | 4.1 | 1.7E-02 | 2.9 | 4.1 | 8.2E-22 | 2.1 | 4.1 | 2.7E-12 | superoxide dismutase |
| RS001625 | 0.6 | 8.2 | 2.6E-01 | 2.3 | 8.2 | 8.6E-08 | 1.8 | 8.2 | 1.1E-04 | |
| RS001803 | 0.0 | 6.6 | 9.0E-01 | 1.3 | 6.6 | 1.2E-07 | 1.3 | 6.6 | 5.7E-08 | glucosyl glucuronosyl transferase |
| RS001896 | 2.8 | 6.1 | 1.3E-08 | 5.1 | 6.1 | 4.6E-23 | 2.3 | 6.1 | 1.9E-06 | |
| RS001996 | 1.0 | 4.5 | 6.5E-03 | 2.3 | 4.5 | 4.8E-13 | 1.4 | 4.5 | 3.9E-05 | |
| RS002006 | 1.7 | 6.6 | 4.3E-18 | 2.7 | 6.6 | 8.2E-45 | 1.1 | 6.6 | 6.4E-08 | atp-binding cassette sub-family g member 4 |
| RS002050 | -0.8 | 7.8 | 1.3E-02 | 1.2 | 7.8 | 1.7E-04 | 2.0 | 7.8 | 9.2E-11 | laccase-like multicopper oxidase 1 |
| RS002148 | 3.9 | 6.0 | 2.7E-22 | 5.2 | 6.0 | 5.3E-35 | 1.3 | 6.0 | 1.5E-03 | |
| RS002475 | -0.1 | 5.8 | 8.8E-01 | 1.4 | 5.8 | 4.7E-05 | 1.5 | 5.8 | 3.4E-05 | |
| RS002636 | -0.1 | 3.2 | 9.3E-01 | 1.7 | 3.2 | 7.0E-05 | 1.8 | 3.2 | 9.7E-05 | kazal-type serine protease inhibitor |
| RS002821 | 1.9 | 5.5 | 7.4E-03 | 4.0 | 5.5 | 1.9E-08 | 2.1 | 5.5 | 3.0E-03 | cathepsin l |
| RS002847 | -1.5 | 8.3 | 6.9E-04 | 2.1 | 8.3 | 9.2E-07 | 3.5 | 8.3 | 4.8E-16 | gram negative bacteria binding protein 2 |
| RS002922 | -1.3 | 0.1 | 3.4E-01 | 2.9 | 0.1 | 1.3E-03 | 4.2 | 0.1 | 4.9E-05 | |
| RS003028 | -0.1 | 6.1 | 8.0E-01 | 1.3 | 6.1 | 1.0E-06 | 1.4 | 6.1 | 2.6E-07 | carbohydrate binding |
| RS003029 | 1.4 | 1.4 | 5.5E-04 | 2.5 | 1.4 | 3.2E-14 | 1.2 | 1.4 | 3.5E-04 | |
| RS003030 | 1.6 | 1.6 | 1.8E-04 | 3.3 | 1.6 | 9.8E-19 | 1.6 | 1.6 | 2.5E-06 | alpha-galactosidase |
| RS003054 | -1.7 | 0.9 | 5.5E-02 | 2.1 | 0.9 | 3.0E-03 | 3.8 | 0.9 | 1.9E-06 | hypothetical protein |
| RS003144 | 0.6 | 6.7 | 6.7E-03 | 1.9 | 6.7 | 6.7E-22 | 1.3 | 6.7 | 1.1E-10 | ejaculatory bulb-specific protein 3 |
| RS003171 | -0.5 | 4.1 | 4.7E-02 | 1.1 | 4.1 | 1.1E-06 | 1.7 | 4.1 | 1.3E-12 | |
| RS003220 | 0.1 | 6.4 | 8.1E-01 | 1.1 | 6.4 | 4.4E-06 | 1.0 | 6.4 | 5.0E-05 | probable peroxisomal acyl-coenzyme a oxidase 1 |
| RS003282 | -0.4 | 3.5 | 2.1E-01 | 1.1 | 3.5 | 1.1E-04 | 1.6 | 3.5 | 1.4E-07 | aldose reductase-like |
| RS003495 | 0.3 | 4.7 | 4.9E-01 | 1.3 | 4.7 | 3.0E-06 | 1.1 | 4.7 | 3.7E-04 | oxidative stress-induced growth inhibitor 2-like |
| RS003509 | 0.2 | 4.3 | 7.3E-01 | 1.2 | 4.3 | 9.2E-05 | 1.1 | 4.3 | 1.3E-03 | a disintegrin and metalloproteinase with thrombospondin motifs 18 |
| RS003522 | -0.3 | 8.9 | 3.6E-01 | 1.2 | 8.9 | 2.3E-05 | 1.5 | 8.9 | 1.1E-07 | aminopeptidase n |
| RS003602 | -1.2 | 1.4 | 1.1E-01 | 1.6 | 1.4 | 7.7E-03 | 2.8 | 1.4 | 1.8E-05 | solute carrier family 28 member 3 |

| | | | | | | | | | | |
|---|---|---|---|---|---|---|---|---|---|---|
| RS003662 | 0.2 | 3.9 | 5.8E-01 | 1.5 | 3.9 | 3.5E-07 | 1.2 | 3.9 | 4.2E-05 | natterin-3-like |
| RS003684 | -0.2 | 5.7 | 4.1E-01 | 1.1 | 5.7 | 6.1E-09 | 1.3 | 5.7 | 7.5E-12 | ankyrin repeat domain-containing protein 11 |
| RS004063 | 0.2 | 8.4 | 5.9E-01 | 1.4 | 8.4 | 7.9E-07 | 1.2 | 8.4 | 5.5E-05 | endoplasmin |
| RS004077 | 0.3 | 11.7 | 6.2E-01 | 1.8 | 11.7 | 1.1E-03 | 1.4 | 11.7 | 8.0E-03 | transferrin |
| RS004088 | 1.7 | 5.7 | 7.6E-11 | 2.7 | 5.7 | 4.7E-26 | 1.0 | 5.7 | 1.5E-04 | |
| RS004129 | 2.1 | 6.3 | 1.2E-17 | 3.9 | 6.3 | 1.6E-51 | 1.7 | 6.3 | 4.7E-13 | venom carboxylesterase-6-like |
| RS004146 | 0.3 | 4.4 | 4.0E-01 | 1.5 | 4.4 | 1.6E-06 | 1.2 | 4.4 | 4.2E-04 | beta-glucosidase |
| RS004767 | -1.0 | 6.1 | 4.0E-02 | 1.5 | 6.1 | 1.1E-03 | 2.4 | 6.1 | 5.5E-08 | |
| RS005168 | 0.6 | 4.2 | 1.6E-01 | 2.4 | 4.2 | 9.2E-10 | 1.7 | 4.2 | 1.4E-05 | ankyrin repeat protein |
| RS005263 | 0.8 | 4.0 | 1.5E-01 | 3.1 | 4.0 | 1.5E-11 | 2.3 | 4.0 | 7.2E-07 | |
| RS005409 | 0.6 | 2.4 | 6.4E-01 | 5.6 | 2.4 | 4.0E-08 | 4.9 | 2.4 | 1.4E-06 | apolipoprotein d-like |
| RS005544 | 0.8 | 7.5 | 2.0E-02 | 2.0 | 7.5 | 4.8E-10 | 1.2 | 7.5 | 5.2E-04 | beta-ureidopropionase |
| RS005818 | 1.8 | 5.9 | 2.8E-13 | 2.9 | 5.9 | 1.3E-30 | 1.1 | 5.9 | 4.3E-05 | |
| RS006066 | 0.0 | 5.0 | 9.6E-01 | 1.2 | 5.0 | 3.6E-08 | 1.2 | 5.0 | 4.7E-08 | d-arabinitol dehydrogenase 1-like |
| RS006091 | 1.6 | 4.4 | 4.4E-06 | 3.2 | 4.4 | 1.2E-21 | 1.6 | 4.4 | 1.4E-06 | |
| RS006134 | 6.0 | 2.4 | 1.8E-16 | 7.6 | 2.4 | 6.1E-27 | 1.6 | 2.4 | 2.1E-03 | alpha-amylase |
| RS006135 | 4.6 | 3.5 | 2.2E-17 | 6.2 | 3.5 | 6.6E-30 | 1.6 | 3.5 | 4.3E-04 | alpha |
| RS006136 | -0.1 | 5.4 | 7.5E-01 | 1.5 | 5.4 | 1.5E-10 | 1.6 | 5.4 | 1.1E-11 | alpha-amylase |
| RS006137 | -0.1 | 7.1 | 7.3E-01 | 1.3 | 7.1 | 9.1E-12 | 1.3 | 7.1 | 4.7E-13 | alpha-amylase |
| RS006303 | 0.6 | 2.6 | 1.2E-01 | 1.6 | 2.6 | 3.1E-08 | 1.1 | 2.6 | 5.5E-04 | female reproductive tract protease |
| RS006304 | 0.3 | 6.9 | 1.2E-01 | 1.7 | 6.9 | 3.6E-21 | 1.3 | 6.9 | 5.9E-14 | retinol dehydrogenase 14 |
| RS006394 | 0.1 | 5.2 | 6.7E-01 | 1.5 | 5.2 | 1.9E-14 | 1.4 | 5.2 | 3.6E-12 | alpha-tocopherol transfer |
| RS006520 | 0.1 | 5.3 | 8.3E-01 | 1.2 | 5.3 | 1.3E-03 | 1.1 | 5.3 | 6.0E-03 | |
| RS006537 | 3.1 | 5.4 | 6.2E-28 | 4.9 | 5.4 | 6.9E-64 | 1.8 | 5.4 | 2.7E-12 | venom carboxylesterase-6-like |
| RS006538 | 3.1 | 4.5 | 1.5E-15 | 4.8 | 4.5 | 4.1E-34 | 1.7 | 4.5 | 3.3E-06 | venom carboxylesterase-6-like |
| RS006718 | 2.2 | -0.3 | 1.1E-01 | 4.4 | -0.3 | 6.0E-06 | 2.2 | -0.3 | 9.5E-03 | |
| RS006883 | -0.2 | 3.6 | 6.8E-01 | 1.6 | 3.6 | 2.8E-09 | 1.7 | 3.6 | 1.8E-10 | crustin-like antimicrobial peptide |
| RS006953 | -0.5 | 7.2 | 2.4E-01 | 1.9 | 7.2 | 8.4E-07 | 2.5 | 7.2 | 6.3E-10 | termicin |
| RS007008 | -1.2 | 1.4 | 4.3E-01 | 5.2 | 1.4 | 2.5E-09 | 6.5 | 1.4 | 9.2E-10 | udp-glucuronosyltransferase 2c1 |
| RS007059 | -1.4 | 5.1 | 2.5E-09 | 1.2 | 5.1 | 9.4E-08 | 2.6 | 5.1 | 1.1E-29 | pi-plc x domain-containing protein 1-like |
| RS007302 | 0.5 | 1.6 | 6.1E-01 | 3.0 | 1.6 | 1.5E-05 | 2.5 | 1.6 | 5.2E-04 | zinc metalloproteinase nas |
| RS007323 | 1.2 | 1.3 | 1.3E-01 | 2.9 | 1.3 | 7.1E-08 | 1.8 | 1.3 | 7.4E-04 | |
| RS007834 | -0.8 | 5.0 | 1.8E-01 | 1.6 | 5.0 | 2.4E-03 | 2.4 | 5.0 | 1.0E-05 | |
| RS007835 | -1.6 | 2.1 | 5.6E-02 | 2.1 | 2.1 | 4.3E-03 | 3.8 | 2.1 | 3.3E-06 | |
| RS007931 | -1.2 | 0.9 | 5.9E-01 | 4.9 | 0.9 | 3.6E-03 | 6.1 | 0.9 | 1.2E-03 | |
| RS008098 | -0.1 | 5.6 | 8.3E-01 | 1.4 | 5.6 | 2.4E-14 | 1.4 | 5.6 | 4.0E-15 | |
| RS008121 | 1.2 | 1.0 | 2.8E-01 | 3.2 | 1.0 | 3.1E-05 | 1.9 | 1.0 | 8.5E-03 | endocuticle structural glycoprotein bd-2 |
| RS008124 | 3.4 | 3.3 | 1.0E-06 | 6.1 | 3.3 | 1.0E-17 | 2.7 | 3.3 | 2.7E-05 | endocuticle structural glycoprotein bd-1-like |
| RS008129 | 3.6 | 2.0 | 1.5E-04 | 6.9 | 2.0 | 3.3E-17 | 3.2 | 2.0 | 3.9E-07 | endocuticle structural glycoprotein bd-2 |
| RS008165 | 1.1 | 7.0 | 2.6E-07 | 2.2 | 7.0 | 1.4E-25 | 1.1 | 7.0 | 6.4E-07 | |
| RS008597 | 0.4 | 0.6 | 6.4E-01 | 2.6 | 0.6 | 1.1E-05 | 2.2 | 0.6 | 4.1E-04 | insulin-like growth factor i |
| RS008750 | -0.2 | 10.2 | 6.2E-01 | 1.9 | 10.2 | 9.2E-10 | 2.1 | 10.2 | 1.2E-11 | protein g12 |
| RS009348 | -0.4 | 7.5 | 7.8E-02 | 1.2 | 7.5 | 1.3E-08 | 1.7 | 7.5 | 1.3E-14 | zinc carboxypeptidase-like |
| RS009610 | -2.0 | 4.3 | 2.2E-01 | 2.8 | 4.3 | 2.9E-03 | 4.8 | 4.3 | 1.0E-04 | |
| RS009611 | -0.1 | 3.4 | 1.0E+00 | 4.0 | 3.4 | 1.7E-03 | 4.0 | 3.4 | 3.8E-03 | chemosensory protein |
| RS009668 | 0.1 | 3.9 | 9.2E-01 | 1.5 | 3.9 | 8.5E-05 | 1.5 | 3.9 | 2.5E-04 | serine proteases 1 2-like |
| RS009735 | -0.2 | 1.6 | 7.4E-01 | 1.4 | 1.6 | 4.2E-05 | 1.5 | 1.6 | 1.4E-05 | homeobox protein orthopedia |
| RS010059 | 1.2 | 4.4 | 6.9E-02 | 3.8 | 4.4 | 1.5E-09 | 2.6 | 4.4 | 5.3E-05 | |

| | | | | | | | | | | |
|---|---|---|---|---|---|---|---|---|---|---|
| RS010352 | 0.5 | 1.1 | 5.2E-01 | 2.9 | 1.1 | 2.1E-06 | 2.3 | 1.1 | 1.4E-04 | |
| RS010353 | 0.8 | 1.9 | 2.4E-01 | 3.1 | 1.9 | 1.2E-07 | 2.2 | 1.9 | 1.4E-04 | lipase member h-like |
| RS010442 | -0.9 | 8.3 | 3.3E-02 | 2.1 | 8.3 | 1.5E-08 | 3.0 | 8.3 | 7.3E-15 | |
| RS010594 | 0.9 | 5.0 | 6.2E-03 | 2.3 | 5.0 | 2.1E-15 | 1.5 | 5.0 | 1.7E-06 | |
| RS010767 | -0.1 | 1.9 | 9.3E-01 | 2.0 | 1.9 | 2.5E-03 | 2.1 | 1.9 | 2.8E-03 | proclotting enzyme |
| RS010778 | -0.2 | 5.6 | 5.4E-01 | 1.6 | 5.6 | 3.7E-14 | 1.8 | 5.6 | 1.5E-16 | lipase 3-like |
| RS010849 | 0.3 | 5.8 | 6.6E-01 | 3.1 | 5.8 | 6.2E-12 | 2.8 | 5.8 | 3.7E-10 | glycine n-methyltransferase |
| RS011169 | 0.1 | 5.5 | 6.6E-01 | 1.1 | 5.5 | 2.4E-11 | 1.0 | 5.5 | 4.0E-09 | insulin-like growth factor 2 mrna-binding protein 1 |
| RS011205 | 1.2 | 11.6 | 7.9E-02 | 6.0 | 11.6 | 1.9E-17 | 4.8 | 11.6 | 1.8E-12 | larval serum protein 1 beta chain |
| RS011571 | 0.2 | 9.6 | 5.6E-01 | 2.5 | 9.6 | 1.4E-16 | 2.3 | 9.6 | 1.2E-13 | |
| RS011587 | 1.1 | 3.5 | 7.2E-02 | 3.4 | 3.5 | 1.7E-09 | 2.3 | 3.5 | 7.0E-05 | collagen alpha-2 chain |
| RS011590 | 0.5 | 2.5 | 5.2E-01 | 2.3 | 2.5 | 2.0E-04 | 1.7 | 2.5 | 7.1E-03 | |
| RS011706 | 0.4 | 8.2 | 8.9E-02 | 1.5 | 8.2 | 3.5E-14 | 1.1 | 8.2 | 6.8E-08 | probable fatty acid-binding protein |
| RS012048 | 0.9 | 3.9 | 6.4E-04 | 1.9 | 3.9 | 5.4E-16 | 1.0 | 3.9 | 2.9E-05 | sodium-independent sulfate anion transporter-like |
| RS012179 | -0.8 | 6.7 | 5.8E-02 | 3.2 | 6.7 | 5.9E-16 | 4.0 | 6.7 | 2.3E-23 | multiple inositol polyphosphate phosphatase 1-like |
| RS012436 | 1.9 | 5.0 | 2.6E-05 | 5.2 | 5.0 | 2.5E-43 | 3.3 | 5.0 | 2.8E-23 | beta-glucosidase |
| RS012687 | -5.0 | 10.1 | 2.9E-61 | 1.2 | 10.1 | 1.0E-05 | 6.2 | 10.1 | 6.0E-85 | endo-beta- -glucanase |
| RS013025 | 0.6 | 6.6 | 3.5E-02 | 1.9 | 6.6 | 3.5E-13 | 1.3 | 6.6 | 2.5E-06 | ankyrin repeat domain protein |
| RS013380 | -0.5 | 4.2 | 1.6E-01 | 1.4 | 4.2 | 9.8E-08 | 1.9 | 4.2 | 7.7E-12 | |
| RS013381 | -1.1 | 0.3 | 2.4E-01 | 1.8 | 0.3 | 6.7E-03 | 3.0 | 0.3 | 1.1E-04 | |
| RS013424 | 0.1 | 6.1 | 9.6E-01 | 4.1 | 6.1 | 8.9E-08 | 4.0 | 6.1 | 6.7E-07 | chemosensory protein |
| RS013433 | -0.6 | 2.0 | 2.7E-01 | 1.2 | 2.0 | 4.5E-03 | 1.9 | 2.0 | 6.5E-05 | |
| RS013912 | 0.0 | 3.3 | 9.9E-01 | 3.0 | 3.3 | 2.2E-04 | 3.0 | 3.3 | 3.7E-04 | |
| RS014005 | 2.7 | 6.7 | 4.3E-08 | 4.7 | 6.7 | 2.7E-19 | 2.0 | 6.7 | 9.2E-05 | cationic trypsin-3-like |
| RS014047 | 1.0 | 2.1 | 8.8E-02 | 2.4 | 2.1 | 1.7E-06 | 1.4 | 2.1 | 7.9E-03 | zinc transporter zip1-like |
| RS014293 | 1.0 | 2.5 | 5.7E-03 | 2.2 | 2.5 | 5.2E-11 | 1.1 | 2.5 | 9.5E-04 | cytochrome p450 |
| RS014496 | -0.1 | 5.2 | 7.8E-01 | 1.1 | 5.2 | 1.7E-04 | 1.2 | 5.2 | 5.8E-05 | dehydrogenase reductase sdr family member 11 |
| RS014890 | 1.2 | 1.0 | 2.2E-01 | 3.4 | 1.0 | 3.9E-05 | 2.2 | 1.0 | 1.0E-02 | secretin receptor-like |
| RS015152 | -0.2 | 1.5 | 7.0E-01 | 1.2 | 1.5 | 2.1E-03 | 1.4 | 1.5 | 6.3E-04 | homeobox protein nkx- |
| RS015197 | 0.2 | 8.3 | 4.4E-01 | 1.8 | 8.3 | 1.1E-13 | 1.5 | 8.3 | 2.2E-10 | clavesin-1 |
| RS015368 | -1.2 | 8.1 | 8.1E-05 | 1.7 | 8.1 | 2.2E-09 | 2.9 | 8.1 | 1.8E-23 | |
| RS015525 | 0.6 | 2.2 | 7.8E-02 | 1.7 | 2.2 | 3.3E-08 | 1.0 | 2.2 | 1.2E-03 | e3 ubiquitin-protein ligase siah1-like |
| RS100005 | 0.4 | 4.3 | 3.1E-01 | 1.9 | 4.3 | 7.4E-11 | 1.5 | 4.3 | 3.4E-07 | myrosinase 1-like |
| RS100018 | -2.9 | 8.6 | 3.3E-12 | 2.1 | 8.6 | 1.7E-07 | 5.0 | 8.6 | 1.2E-29 | gram negative bacteria-binding protein |
| RS100025 | 2.7 | 2.0 | 4.3E-06 | 4.3 | 2.0 | 8.3E-14 | 1.6 | 2.0 | 3.7E-03 | |
| RS100026 | -1.7 | 4.2 | 8.2E-02 | 4.3 | 4.2 | 1.4E-06 | 6.0 | 4.2 | 1.6E-10 | |
